# Structure-based design of selective salt-inducible kinase (SIK) inhibitors

**DOI:** 10.1101/2021.04.08.439011

**Authors:** Roberta Tesch, Marcel Rak, Monika Raab, Lena M. Berger, Thales Kronenberger, Andreas C. Joerger, Benedict-Tilman Berger, Ismahan Abdi, Thomas Hanke, Antti Poso, Klaus Strebhardt, Mourad Sanhaji, Stefan Knapp

**Author notes:** Correspondence: Stefan Knapp. These authors contributed equally to the work (Roberta Tesch, Marcel Rak).

## Abstract

Salt-inducible kinases (SIKs) are key metabolic regulators. Imbalance of SIK function is associated with the development of diverse cancers, including breast, gastric and ovarian cancer. Chemical tools to clarify the roles of SIK in different diseases are, however, sparse and are generally characterized by poor kinome-wide selectivity. Here, we have adapted the pyrido[2,3-*d*]pyrimidin-7-one-based PAK inhibitor G-5555 for the targeting of SIK, by exploiting differences in the back-pocket region of these kinases. Optimization was supported by high-resolution crystal structures of G-5555 bound to the known off-targets MST3 and MST4, leading to a chemical probe, MRIA9, with dual SIK/PAK activity and excellent selectivity over other kinases. Furthermore, we show that MRIA9 sensitizes ovarian cancer cells to treatment with the mitotic agent paclitaxel, confirming earlier data from genetic knockdown studies and suggesting a combination therapy with SIK inhibitors and paclitaxel for the treatment of paclitaxel-resistant ovarian cancer.

## INTRODUCTION

Salt-inducible kinases (SIK1-3) are members of the AMP-activated protein kinase (AMPK) sub-family of the calcium/calmodulin-dependent kinase (CaMK) group. These serine/threonine kinases act as regulators of energy homeostasis and metabolic stress. The SIK family member SIK2, for example, is activated in cells recovering from starvation, leading to phosphorylation and hence activation of the transcription factor cAMP response element-binding protein (CREB1). However, SIK family members also repress CREB1-mediated gene expression by modulating the activity of the associated transducer of regulated CREB1 (TORC).^1–3^ In addition to the key function of SIK in regulating metabolism, imbalance of SIK has been observed in the context of several diseases, especially in cancer, with both tumor promoting and tumor suppressive roles being reported.^4^ SIK2 is often deleted in breast cancer, and downregulation of SIK1 has been linked to a tumor suppressor role but also the development of metastasis by promoting p53-dependent anoikis.^5^ In contrast, SIK3 is highly expressed in breast cancer patients, with an effect on the inflammation process in this type of cancer,^6^ while SIK2 is often amplified in diffuse large B cell lymphoma (DLBCL), increasing cell survival and tumor progression.^7^ In leukemia, SIK3 is critical for cell proliferation by regulating subcellular localization and activity of HDAC4.^8,9^ In ovarian cancer, overexpression of SIK2 has been observed in adipocyte-rich environment of metastatic deposits, where it enhances lipid synthesis, resulting in poor prognosis.^10–12^ Depletion of SIK2 in ovarian cancer delays mitotic progression and decreases PI3K and AKT phosphorylation levels.^13^ Interestingly, SIK2 knockdown re-sensitizes ovarian cancer cells to taxane-based chemotherapy.^14^

We have recently shown that in samples from patients with gastric cancer, there were increased levels of SIK2 mRNA and protein in advanced stages of the tumor compared with lower grade tumor, independent of the metastatic stage.^15^ Overall, these data highlight the complex roles of SIK family proteins in different types of cancer, making them important therapeutic targets. To clarify the multifaceted roles of these kinases in disease and normal physiology, chemical tools are urgently needed. A number of potent SIK inhibitors have been reported and have been used to investigate the roles of SIK kinases in cancer (Figure 1). Among those inhibitors are the tyrosine kinase inhibitors dasatinib (**1**) and bosutinib (**2**), which are currently used for the treatment of chronic myelogenous leukemia by targeting ABL and SRC family kinases, which all bear a threonine residue at the gatekeeper position.^16^ SIK kinases also harbor a threonine gatekeeper and are therefore inhibited by these and many other tyrosine kinase inhibitors. HG-9-91-01 (3) is a pan-SIK inhibitor with IC_50_ values of 0.92 nM, 6.6 nM and 9.6 nM for SIK1, SIK2 and SIK3, respectively, but it also inhibits several other kinases with important roles in tumorigenesis, including SRC, YES, BTK, FGFR1 and ephrin receptors.^17^ The inhibitor YKL-05-099 (**4**) has been designed based on HG-9-91-01 as a more selective chemical tool to investigate the roles of SIK2 in vivo, which highlighted the possibility of a therapeutic window for SIK inhibitors to achieve immunomodulatory effects without misbalancing the normal glucose metabolism.^18^ However, **4** is still not sufficiently selective for mechanism-based studies because it inhibited more than 60 kinases by more than 90% at 1 µM in a panel of 468 kinases.^18^ Recently, compound ARN-3236 (**5**) has been reported as a panSIK inhibitor with modest selectivity towards SIK2 over SIK1 and SIK3, with IC_50_ values of less than 1 nM, 21 nM and 6 nM, respectively.^19^ Treatment of ovarian cancer cells with ARN-3236 prevented centrosome separation and led to sensitization to paclitaxel treatment.^14^ However, selectivity data for **5** is only available against 74 kinases, at a single concentration. This rather limited selectivity screen has identified significant activity of this compound (more than 80% inhibition at 0.5 µM) against the kinases JAK2, LCK, NUAK2, SRPK1, and VEGFR2 (**Supplementary Figure 1**).^19^ Thus, all currently available SIK inhibitors display significant off-target activity, limiting their use as chemical probes.

**Figure 1.**
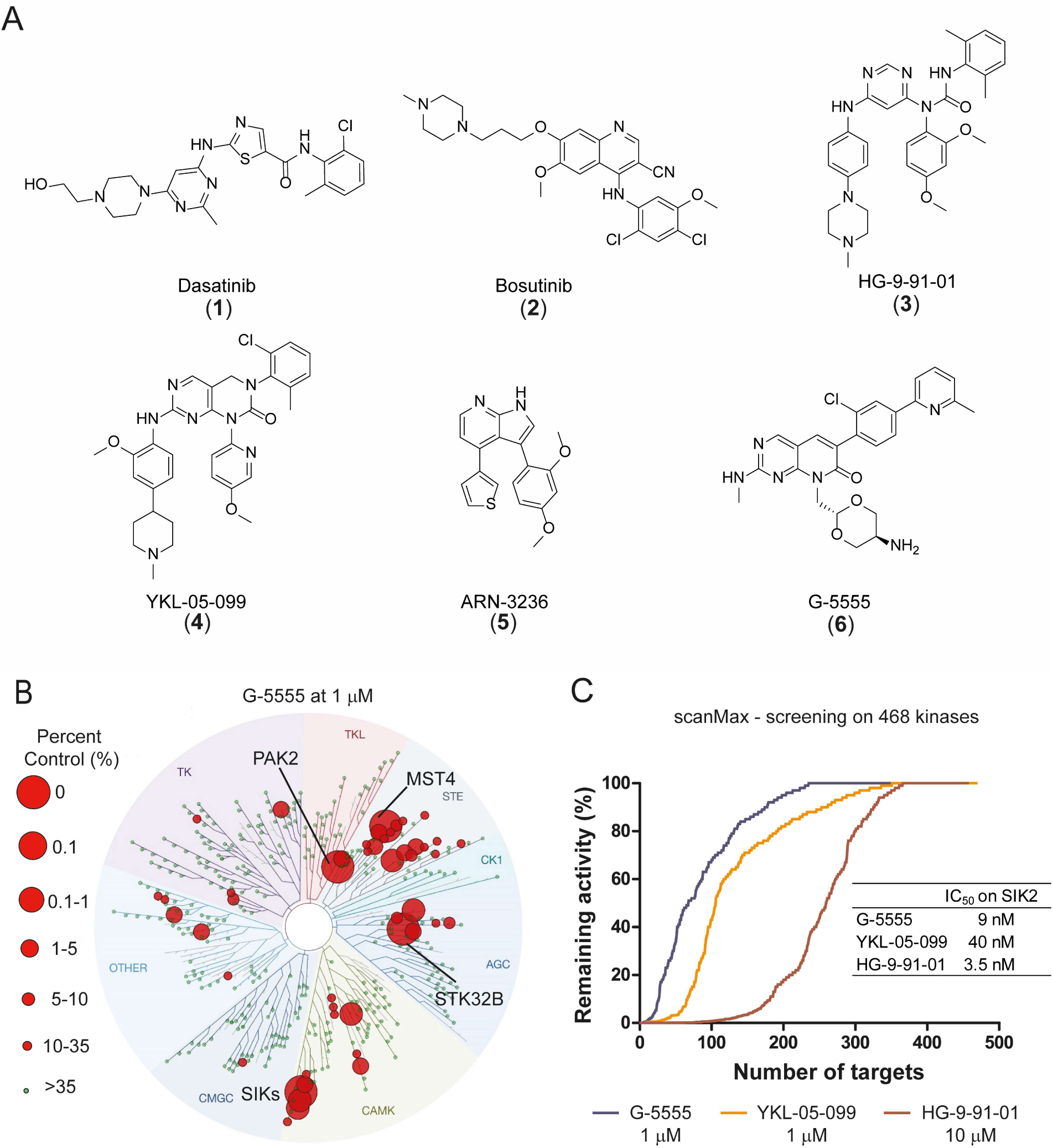
Selectivity profile of SIK inhibitors. (A) Chemical structures of known SIK inhibitors. (B) Selectivity profile of G-5555 at a concentration of 1 µM, using the scanMAX℠ kinase assay panel of 468 kinases (*Eurofins Scientific*). The data is shown on a kinase phylogenetic tree, provided by *Eurofins Scientific*. Inhibited targets are highlighted by red circles, with the circle radius representing % activity compared with untreated control. Activities on mutant kinases are not shown. Data summarized in this figure can be found in **Supplementary Table 2** (C) Comparison of the selectivity profile of three SIK inhibitors using the KINOMEscan^®^ assay platform (*Eurofins Scientific*) performed for 468 kinases. Data is reported as percent of activity remaining in the presence of inhibitor relative to activity measured in absence of inhibitor (100%).

We therefore aimed to develop a well-characterized chemical tool for SIK family members that lacks activity on tyrosine kinases and other off-targets. In addition, sequence conservation within the ATP binding site suggests that isoform selectivity would be difficult to achieve using ATP competitive inhibitors. Searching the literature for a chemical starting point for the development of SIK inhibitors with improved selectivity, we focused on three criteria: (i) a small number of reported off-targets, (ii) sufficiently comprehensive selectivity profiling on a large set of kinases and (iii) a synthetically easily accessible scaffold. Of particular interest to us was the PAK inhibitor G-5555 (**6**), which has been developed by Ndubaku and co-workers.^20^ In a comprehensive selectivity panel of 235 kinases, G-5555 (**6**) showed inhibition of PAK family members, related STE kinases (KHS1, MST3, MST4, YSK1) and the tyrosine kinase LCK, but also potent activity against SIK2.^20^ The dual activity of G-5555 against STE kinases as well as SIK is intriguing, considering that these kinases share very little sequence homology, including key residue changes such as the gatekeeper. We therefore hypothesized that selectivity between the main known G5555 targets within the STE family and SIK can be achieved. Another PAK1 inhibitor structurally related to G-5555 was developed by Rudolph and co-workers, but this inhibitor was profiled against a comparatively small panel of 96 kinases^21^, which motivated us selecting G-5555 (**6**) as a chemical starting point for this study. Here, we report the development and biological evaluation of a suitable chemical tool compound targeting SIK based on scaffold **6**. Improved potency and selectivity were achieved by structural variation of the back-pocket binding motif of compound **6**, which was guided by crystallographic data on STE off-targets and molecular dynamics simulations.

## RESULTS AND DISCUSSION

### Extended selectivity profile of SIK inhibitor G-5555

To validate compound G-5555 (6) as a suitable starting scaffold for the development of selective SIK inhibitors, we decided profiling the selectivity of G-5555 more broadly using the scanMAX^℠^ kinase assay panel of 468 kinases (*Eurofins Scientific*), revealing a few additional off-targets (**Supplementary Table 1-2**). Comparing the screening data for **4** and **6** at 1 µM compound concentration, compound **6** showed a significantly improved selectivity profile (Figure 1). In addition, screening compound **6** at lower concentration (100 nM) revealed only few off-targets (**Supplementary Figure 2**). Thus, given the sequence and structural diversity of SIK compared with STE kinases, compound **6** represented an attractive starting point for the development of selective SIK inhibitors.

### Strategy for designing selective SIK pyrido[2,3-*d*]pyrimidin-7-one inhibitors

Crystallizing any isoform of SIK has been challenging so far due to poor expression yields in insect cell expression systems and instability of the protein. To provide a structural basis for the rational design of compounds with high selectivity for SIK kinases, we therefore determined crystal structures of G-5555 bound to the off-targets MST3 and MST4 (PDB codes 7B30 and 7B36) (Figure 2A-B and **Supplementary Figure 3**). The resolution of the structures was 2.1 Å in both cases. These structures then served as surrogate models for rational modifications and as a template for generating a SIK2 family homology model. We chose SIK2 as a representative family member. However, all specific SIK2 model residues discussed in the manuscript and considered for the inhibitor design are conserved among SIK family proteins. The binding mode of G-5555 in both of these highly conserved kinase domains revealed (i) that the chlorinated aromatic ring system is perpendicular to the pyrido[2,3-*d*]pyrimidin-7-one and the 6-methylpyridine ring system, positioning the pyridine ring near the catalytic residue Lys65 (MST3 residue numbering), (ii) a hydrogen-bond interaction of the 5-amino-1,3-dioxane group with Asn161, and most interestingly, (iii) the formation of a water-mediated hydrogen-bond network linking the glycine-rich loop (P-loop) residue Ser46, the catalytic Lys65, the pyridine ring of G-5555 and the carbonyl group of the pyrido[2,3-*d*]-pyrimidin-7-one ring. Notably, in the reported complex of G-5555 with PAK1 (PDB code 5DEY), the glycine-rich loop has moved out of the ATP-binding pocket, and the pyridine ring is packed between Val342 and Glu315. Importantly, MST3 and MST4 harbor a bulkier isoleucine (Ile109 and Ile97, respectively) instead of the valine (Val342) present in PAK1 (**Supplementary Figure 4A**). When comparing the complexes G-5555-MST3/4 and G-5555/PAK1 with the SIK family homology model, a number of differences in the back-pocket region and other ATP-binding site residues became apparent (Figure 2C). Most strikingly, SIK family members bear a threonine residue at the gatekeeper position and a methionine at position +4 after the conserved glutamic acid on the αC helix. In contrast, both MSTs and PAK1 have a methionine in the gatekeeper position, and either a leucine (Leu86 in MST3) or a methionine (Met319 in PAK1) at position +4 on the αC helix. The pyrido[2,3-*d*]-pyrimidin-7-one ring of G-5555 forms hydrogen-bond interactions with the hinge region in a similar way in both MST structures and the PAK1 complex.

**Figure 2.**
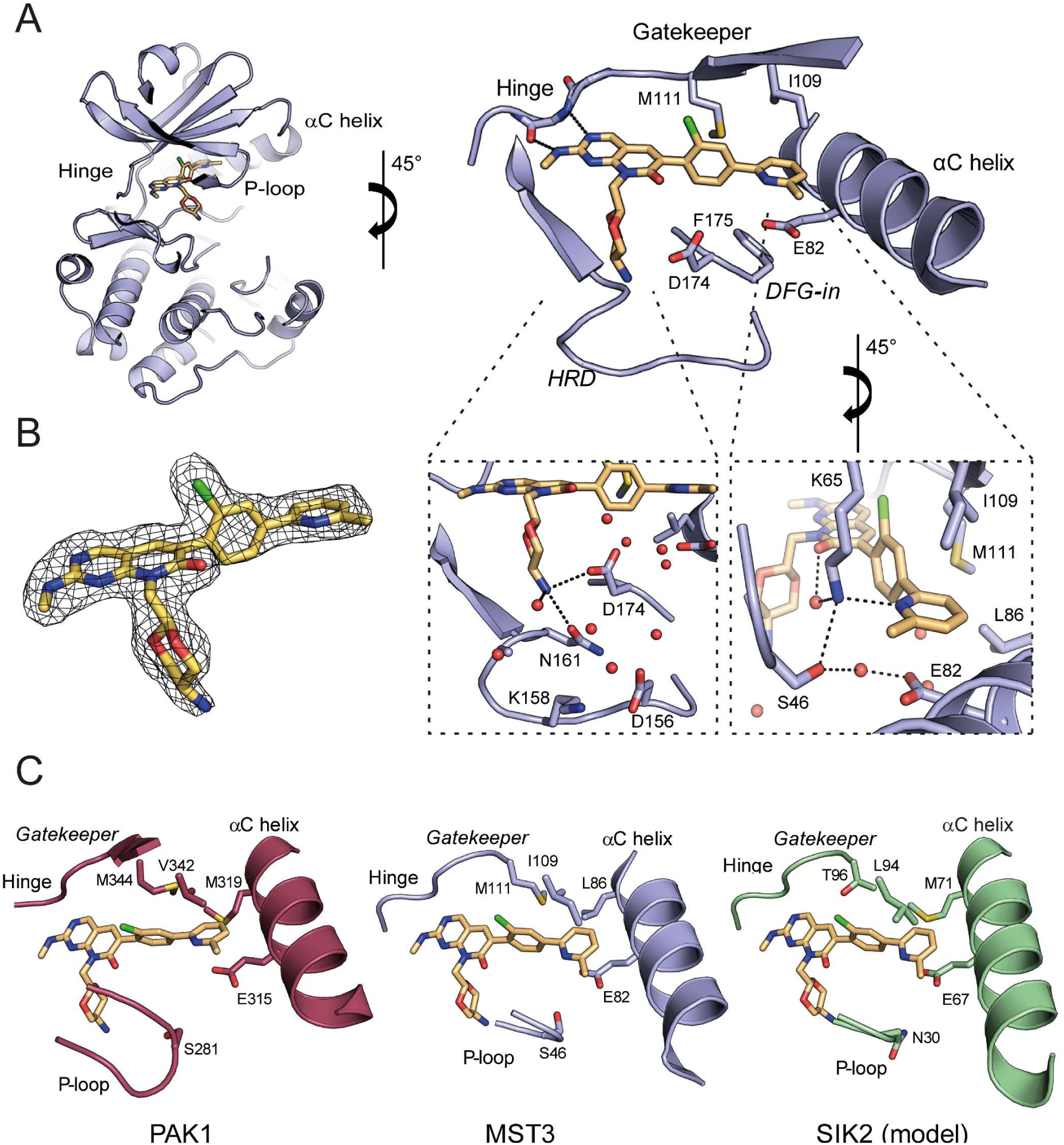
Crystal structure of MST3 in complex with G-5555. (A) Binding-mode analysis of G-5555 in MST3. The insets show the water network around the amino group of the 1,3-dioxane moiety in the HRD region (left) and the interactions of the 6-methylpyridine moiety in the back-pocket region (right). Selected hydrogen bonds are highlighted by dashed lines. (B) 2F_o_-F_c_ omit electron density map around the ligand shown at a contour level of 2.0 σ. (C) Comparison of the back-pocket region of PAK1 (PDB code 5DEY), MST3 (PDB code 7B30), and SIK2 (homology model).

To explore whether the water network observed was not due to crystal packing effects, we determined the water thermodynamic profile with a WaterMap simulation. The water profile was calculated from MD simulations with explicit waters and analyzed as a cluster of hydration sites for calculation of entropy, enthalpy, and free energy properties. We identified conserved hydration sites that matched the position of the water molecules in the crystal structure MST3/G-5555 between Ser46 and Glu82 (hydration site 5) and between Lys65 and the carbonyl group of the pyrido[2,3-*d*]-pyrimidin-7-one ring (hydration site 4) (Figure 3A). These enthalpically favorable hydration sites suggest that the water network observed in the back-pocket region is not due to crystal packing but comprises stable structural water molecules that contribute to stabilizing the binding mode of G-5555. In addition, we analyzed the binding site without the bound compound and observed unfavorable hydration sites that overlap with the binding region of G-5555. Optimal geometry and space fill of the inhibitor is therefore important for efficiently displacing those unfavorable water molecules from the binding pocket.

**Figure 3.**
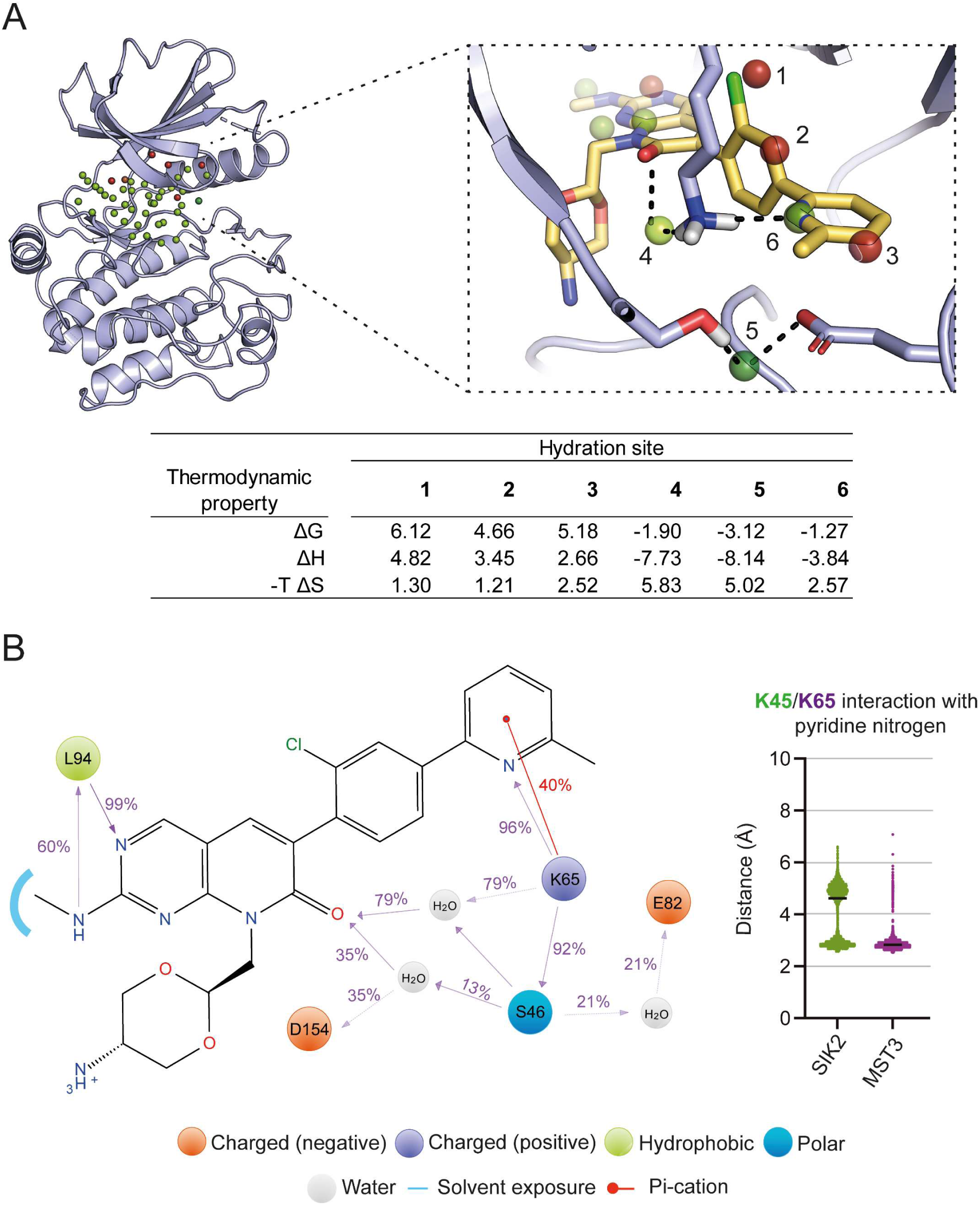
Stabilization of G-5555 during MD simulation and water thermodynamic profile. (A) WaterMap simulation of the G-5555 binding pocket in unbound MST3 highlighting favorable (green) and unfavorable (red) hydration sites during the simulation and their location relative to bound G-5555. (B) Frequency of interaction (in percent) between the inhibitor and a particular residue during the MD simulation and distance profile between the catalytic lysine and the pyridine moiety of G-5555 in both SIK2 (green) and MST3 (purple).

To provide additional structural insights for inhibitor optimization, we next carried out molecular dynamics (MD) simulations on the complex of MST3/G-5555 and the model SIK2/G-5555 (4 µs). Our simulation on the MST3 complex showed a stable interaction between Ser46 and the pyridyl moiety of G-5555 mediated by Lys65 (Figure 3B). Through the course of the simulation, the interaction between Lys65 and the pyridyl moiety, and Lys65 and Ser46 occurred in 96% and 92% of the snapshots along the 4 µs simulation, respectively, suggesting a major role of the MST3 glycine-rich loop in G-5555 stabilization upon binding. Interestingly, the interaction between the G-5555 pyridyl moiety and the equivalent lysine (Lys45) in the SIK2 complex was more flexible during the simulation (Figure 3B). The overall movement of the P-loop in the SIK2 model was analyzed by monitoring the distance between Asn30 (conserved in all three SIK family members) and the back pocket (**Supplementary Figure 5A**). To assess flexibility in the back pocket, we monitored the distance between the gatekeeper residues and the αC helix in both SIK2 and MST3, showing that the dynamics of this region are more constrained in MST3 than in SIK2 (**Supplementary Figure 5B**) and suggesting that certain modifications of the pyridyl moiety may impair binding to MST3 but not SIK family proteins. On the basis of this MD analysis and the characteristic amino acid variations in the pyridyl moiety binding pocket, we hypothesized that modifications at the ortho position of the pyridyl moiety may destabilize the interaction with the back pocket of MST but not SIK kinases.

Based on the crystallographic data on the G-5555 binding mode and the *in silico* calculations, we designed a first series of compounds in order to explore the potential of the pyrido[2,3-*d*]pyrimidin-7-one scaffold for the development of selective SIK inhibitors. For the optimization of back-pocket interactions, we focused on substitution at positions 3, 4 and 6 of the pendant pyridyl moiety (R1), while modification in R2 addressed optimization of the solvent-exposed region (Figure 4).

**Figure 4.**
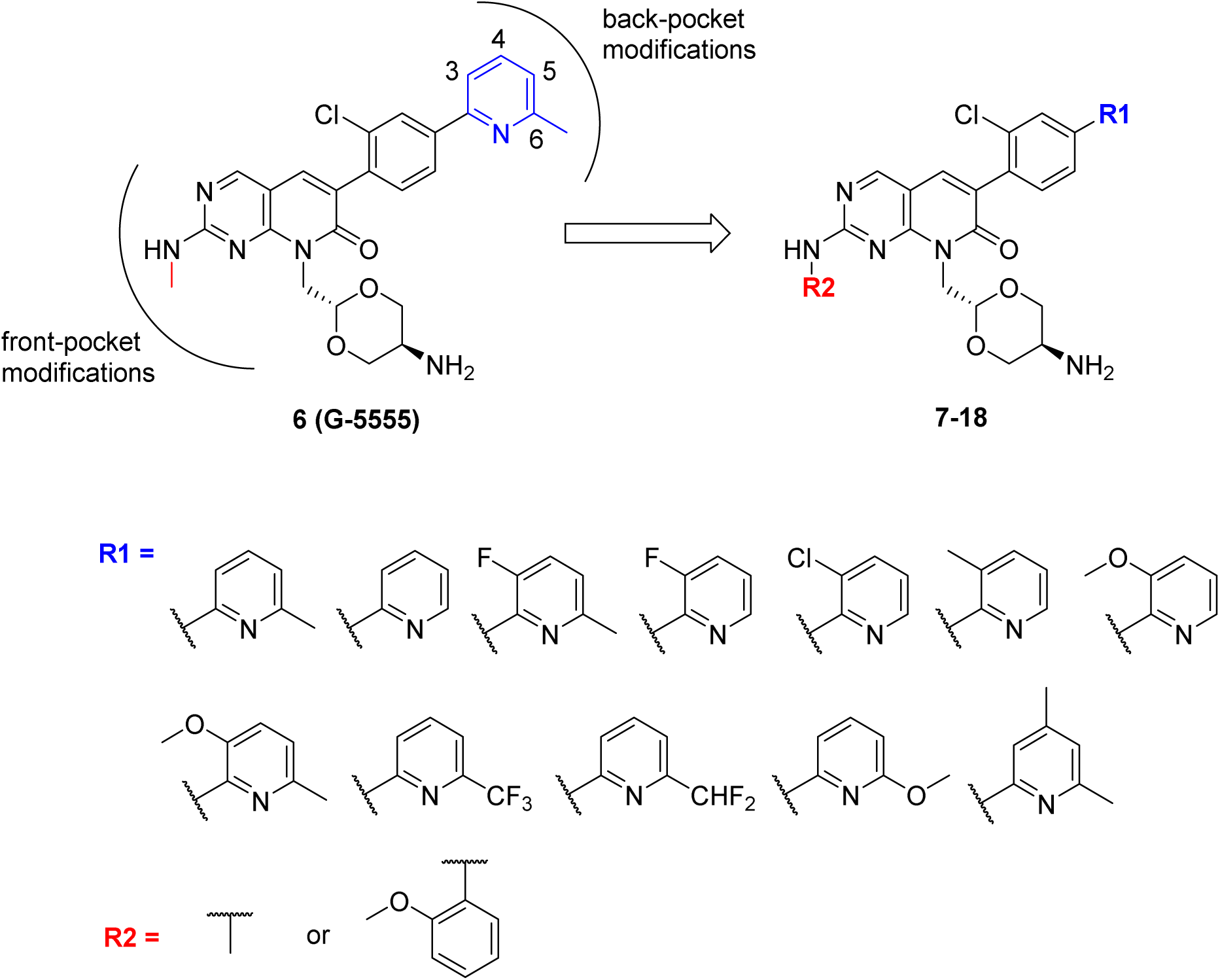
Design of a focused series of pyrido[2,3-*d*]pyrimidin-7-one analogues of PAK1 inhibitor G-5555. Based on the back-pocket interactions of G-5555 in both PAK1 and MST3/4, the pyridyl moiety was substituted at position 3, 4 and 6 with different electron withdrawing and donating groups.

### Synthesis of G-5555 derivatives

Compounds **7-18** were synthesized using a nine-step synthetic route, based on the synthetic strategy published by Rudolph *et al*.^21^, as outlined in Scheme 1 and 2. The synthesis started with the esterification of commercially available 2-(4-bromo-2-chlorophenyl)acetic acid (**19**) with ethanol to obtain ethyl 2-(4-bromo-2-chloro-phenyl)acetate (**20**) (step a, Scheme 1). A ring-closure reaction of 20 with 4-amino-2-(methylthio)pyrimidine-5-carbaldehyde under basic conditions (step b, Scheme 1) led to the formation of the pyrido[2,3-*d*]pyrimidine scaffold **21**. In the following Miyaura borylation reaction (step c, Scheme 1), the bromo substituent of **21** was replaced by a boronic acid pinacol ester to obtain **22**. The 5-amino-1,3-dioxane moiety, required for the subsequent nucleophilic substitution, was synthesized in a two-step reaction. The first step was the protection of the amine group of serinol (**23**) with a phthalimide protecting group (step a, Scheme 2), leading to **24**. In the following ring-closure reaction (step b, Scheme 2), **24** was treated with 2-bromo-1,1’-diethoxyethane and p-toluenesulfonic acid in toluene to obtain the 5-amino-1,3-dioxane ring **25**. A Suzuki cross-coupling reaction (step c, Scheme 2) enabled the derivatization of the lead structure by substituting the boronic acid pinacol ester of **22** with different pyridine derivatives, leading to the formation of the corresponding compounds **26-37**. **25** was used in the subsequent nucleophilic substitution reaction (step d, Scheme 2) with compounds **26-37** to obtain the corresponding compounds **38-49**. The last step of the synthetic route started with an oxidation of the methylthioether of **38-49** with *m-*CPBA (step e, Scheme 2), leading to a mixture of the corresponding sulfoxide and sulfone product. The mixture was used in the following nucleophilic aromatic substitution without further purification. Compound 7 was synthesized under acidic conditions by treatment of the corresponding mixture of sulfoxide/sulfone product with o-anisidine (step f, Scheme 2), and compounds **8-18** were synthesized under basic conditions by treatment with methylamine (step g, Scheme 2).

**Scheme 1.**
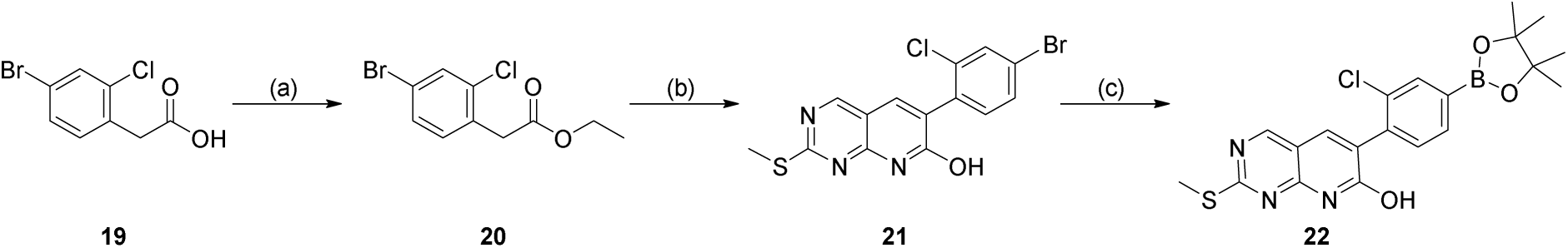
Synthesis of intermediate **22**^a^. ^a^ Reagents and conditions: (a) H_2_SO_4_, ethanol, 15 h, 100 °C; (b) 4-amino-2-(methylthio)-pyrimidine-5-carbaldehyde, K_2_CO_3_, DMF, 16 h, 120 °C; (c) bis(pinacolato)diboron, [1,1’-bis(diphenylphosphino)-ferrocene]dichloropalladium(II) (Pd(dppf)Cl_2_), KOAc, dioxane, 18 h, 115 °C.

**Scheme 2.**
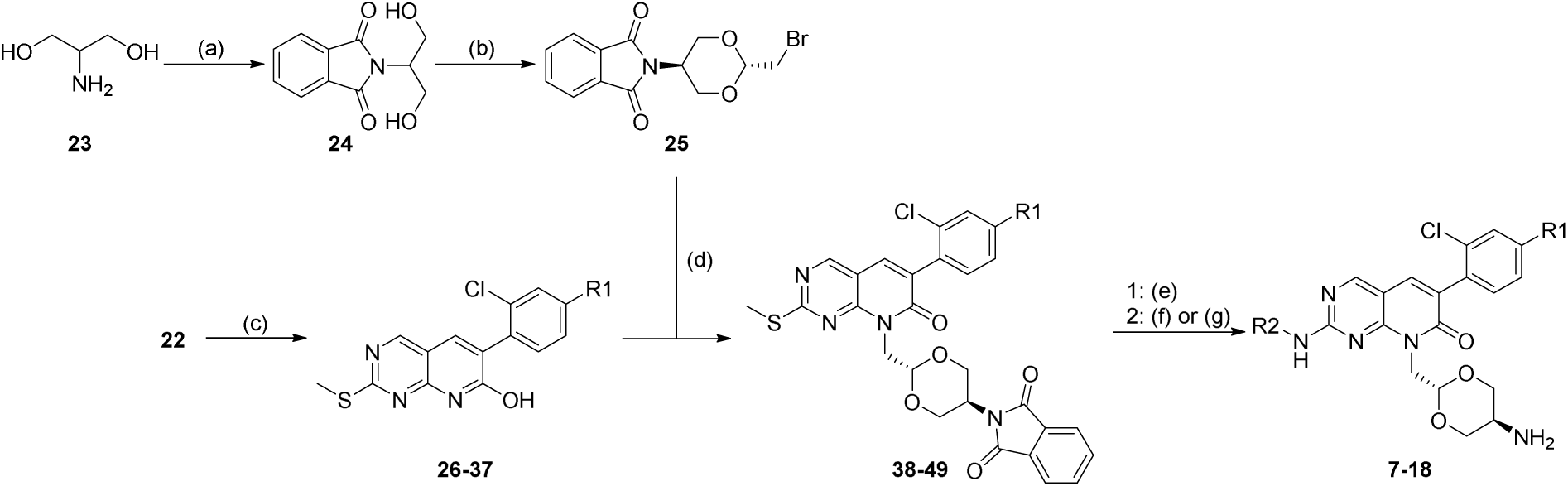
Synthesis of pyrido[2,3-*d*]pyrimidin-7-one derivatives (**7-18**)^a^. ^a^ Reagents and conditions: (a) phthalic anhydride, DMF, 4 h, 90 °C; (b) 2-bromo-1,1-diethoxyethane, p-toluenesulfonic acid, toluene, 24 h, 115 °C; (c) 2-bromopyridines, Pd(dppf)Cl_2_, KOAc, dioxane/H_2_O (2:1), 18 h, 107 °C; (d) Cs_2_CO_3_, DMF, 18 h, 100 °C; (e) 3-chloroperbenzoic acid (*m-*CPBA), DCM, 3 h, RT; (f) *o*-anisidine, HCl, ethanol, 17 h, 100 °C; (g) methylamine, *N,N*’-diisopropylethylamine (DIPEA), ethanol, 17 h, 85 °C. For R1 and R2 see Table 1.

### Structure-activity relationship of the G-5555-based compound series

We evaluated the activity of the compounds against SIK2 and SIK3 using the cellular target engagement assay NanoBRET^22,23^ and compared these data with the off-target profile on MSTs and PAK1 based on melting temperature shift (Δ*T*_m_) from a differential scanning fluorimetry (DSF) assay.^24^ In DSF assays, the melting point of a protein in the presence and absence of a compound was measured. In this assay format, the thermal denaturation of a protein is measured with the help of a fluorescent dye that changes its fluorescence properties upon binding to hydrophobic surfaces exposed in unfolded proteins. The relative binding strength of a ligand is measured by an increase in protein melting temperature (Δ*T*_m_) compared to the unbound protein. Comparison with binding affinities measured in solution shows that Δ*T*_m_ values of a target protein linearly correlate with affinity of ligands with conserved binding modes.^23^ The pan-SIK inhibitors **3** and **4** were used as a positive control together with G-5555 (**6**). The assay data correlated well when comparing the melting temperature profiles on off-targets with the IC_50_ values reported in the literature. G-5555 showed a melting temperature shift (Δ*T*_m_) of 9.9 °C, 6.6 °C and 6.6 °C for MST3, MST4 and PAK1, respectively. Reported IC_50_ values for these targets are 43 nM, 20 nM, and 3.7 nM for MST3, MST4, and PAK1, respectively.^20^ Our first attempt was to develop a hybrid of G-5555 (**6**) and YKL-05-099 (**4**) because the ortho-methoxyphenyl moiety present in 4 had been highlighted as an important moiety for achieving selectivity for MSTs and PAK1.^18^ This effort led to compound **7**, which was equipotent towards SIK2 and 4 times more potent towards SIK3 in NanoBRET^TM^ assay compared with G-5555 (**6**) and YKL-05-099 (**4**), but it still potently inhibited MST3, MST4 and PAK1 as shown by Δ*T*_m_ values of 9.5 °C, 7.8 °C and 5.0 °C (Table 1). We confirmed the lack of selectivity of compound **7** using the scanMAX^℠^ kinase assay panel of 468 kinases (*Eurofins Scientific*) at a concentration of 1 µM, obtaining an S-score (35) of 0.18, a value that represents the count of data points with higher than 35% inhibition divided by the total number of data points (excluding mutants) (**Supplementary Figure 6A-B**, **Supplementary Table 3**).

**Table 1.**
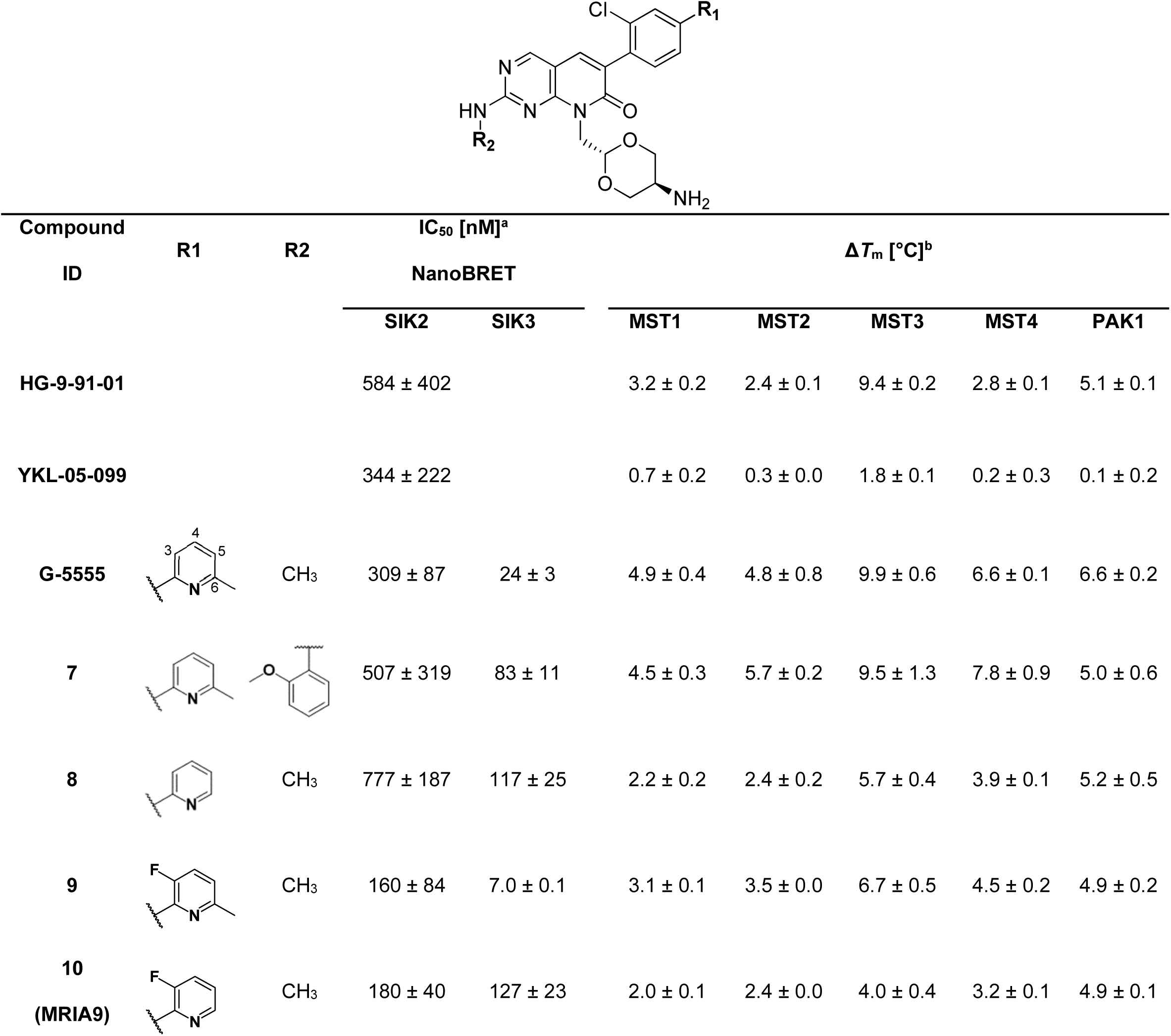
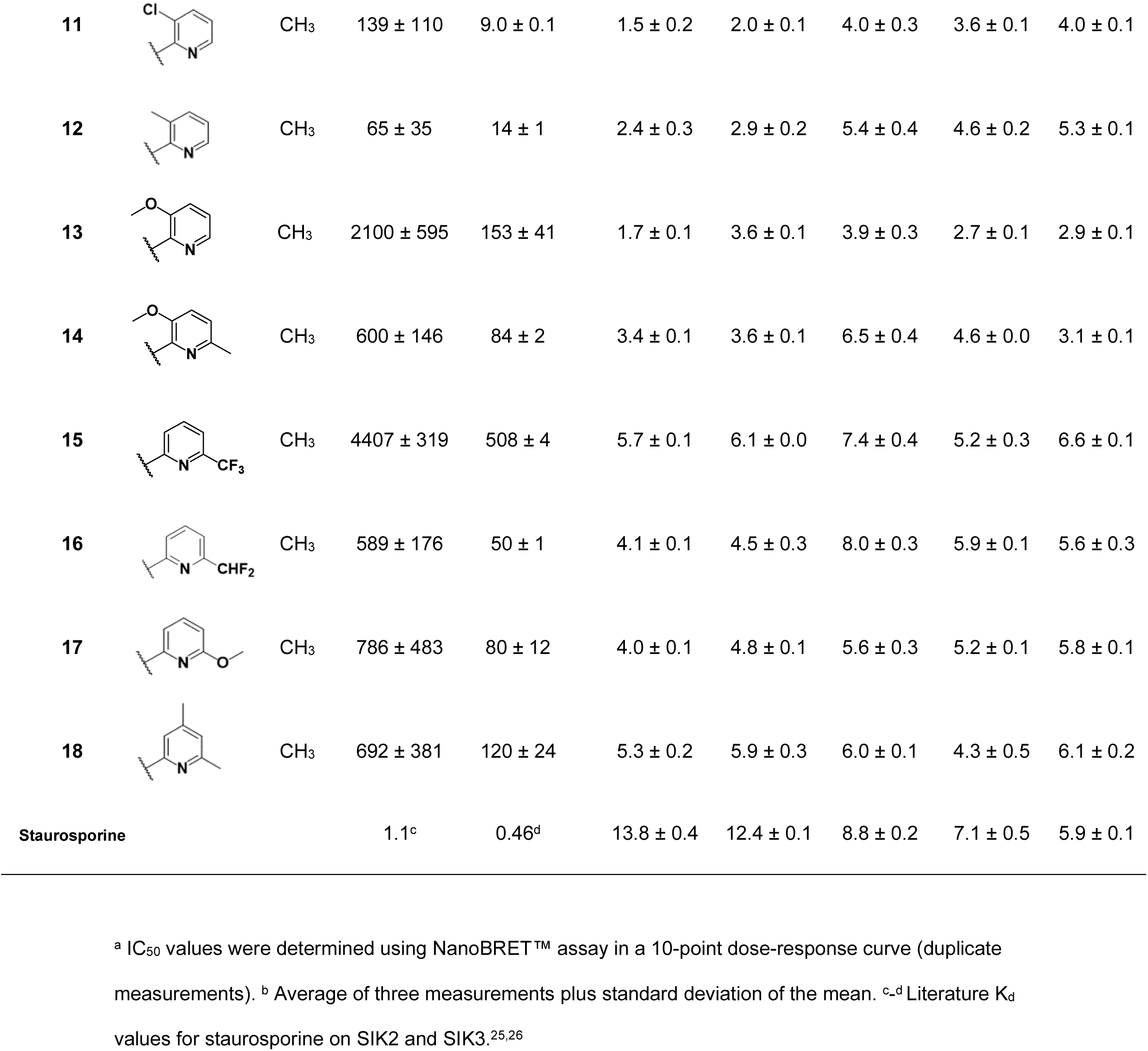
Selectivity profile of pyrido[2,3-*d*]pyrimidin-7-one series **7-18** as SIK2 inhibitors.

| Compound ID | R1 | R2 | IC<sub>50</sub> [nM]<sup>a</sup> |  | ΔT<sub>m</sub> [°C]<sup>b</sup> |  |  |  |  |
| --- | --- | --- | --- | --- | --- | --- | --- | --- | --- |
|  |  |  | NanoBRET |  |  |  |  |  |  |
|  |  |  | SIK2 | SIK3 | MST1 | MST2 | MST3 | MST4 | PAK1 |
| HG-9-91-01 |  |  | 584 ± 402 |  | 3.2 ± 0.2 | 2.4 ± 0.1 | 9.4 ± 0.2 | 2.8 ± 0.1 | 5.1 ± 0.1 |
| YKL-05-099 |  |  | 344 ± 222 |  | 0.7 ± 0.2 | 0.3 ± 0.0 | 1.8 ± 0.1 | 0.2 ± 0.3 | 0.1 ± 0.2 |
| G-5555 |  | CH<sub>3</sub> | 309 ± 87 | 24 ± 3 | 4.9 ± 0.4 | 4.8 ± 0.8 | 9.9 ± 0.6 | 6.6 ± 0.1 | 6.6 ± 0.2 |
| 7 |  |  | 507 ± 319 | 83 ± 11 | 4.5 ± 0.3 | 5.7 ± 0.2 | 9.5 ± 1.3 | 7.8 ± 0.9 | 5.0 ± 0.6 |
| 8 |  | CH<sub>3</sub> | 777 ± 187 | 117 ± 25 | 2.2 ± 0.2 | 2.4 ± 0.2 | 5.7 ± 0.4 | 3.9 ± 0.1 | 5.2 ± 0.5 |
| 9 |  | CH<sub>3</sub> | 160 ± 84 | 7.0 ± 0.1 | 3.1 ± 0.1 | 3.5 ± 0.0 | 6.7 ± 0.5 | 4.5 ± 0.2 | 4.9 ± 0.2 |
| 10<br>(MRIA9) |  | CH<sub>3</sub> | 180 ± 40 | 127 ± 23 | 2.0 ± 0.1 | 2.4 ± 0.0 | 4.0 ± 0.4 | 3.2 ± 0.1 | 4.9 ± 0.1 |
| 11 |  | CH<sub>3</sub> | 139 ± 110 | 9.0 ± 0.1 | 1.5 ± 0.2 | 2.0 ± 0.1 | 4.0 ± 0.3 | 3.6 ± 0.1 | 4.0 ± 0.1 |
| 12 |  | CH<sub>3</sub> | 65 ± 35 | 14 ± 1 | 2.4 ± 0.3 | 2.9 ± 0.2 | 5.4 ± 0.4 | 4.6 ± 0.2 | 5.3 ± 0.1 |
| 13 |  | CH<sub>3</sub> | 2100 ± 595 | 153 ± 41 | 1.7 ± 0.1 | 3.6 ± 0.1 | 3.9 ± 0.3 | 2.7 ± 0.1 | 2.9 ± 0.1 |
| 14 |  | CH<sub>3</sub> | 600 ± 146 | 84 ± 2 | 3.4 ± 0.1 | 3.6 ± 0.1 | 6.5 ± 0.4 | 4.6 ± 0.0 | 3.1 ± 0.1 |
| 15 |  | CH<sub>3</sub> | 4407 ± 319 | 508 ± 4 | 5.7 ± 0.1 | 6.1 ± 0.0 | 7.4 ± 0.4 | 5.2 ± 0.3 | 6.6 ± 0.1 |
| 16 |  | CH<sub>3</sub> | 589 ± 176 | 50 ± 1 | 4.1 ± 0.1 | 4.5 ± 0.3 | 8.0 ± 0.3 | 5.9 ± 0.1 | 5.6 ± 0.3 |
| 17 |  | CH<sub>3</sub> | 786 ± 483 | 80 ± 12 | 4.0 ± 0.1 | 4.8 ± 0.1 | 5.6 ± 0.3 | 5.2 ± 0.1 | 5.8 ± 0.1 |
| 18 |  | CH<sub>3</sub> | 692 ± 381 | 120 ± 24 | 5.3 ± 0.2 | 5.9 ± 0.3 | 6.0 ± 0.1 | 4.3 ± 0.5 | 6.1 ± 0.2 |
| Staurosporine |  |  | 1.1<sup>c</sup> | 0.46<sup>d</sup> | 13.8 ± 0.4 | 12.4 ± 0.1 | 8.8 ± 0.2 | 7.1 ± 0.5 | 5.9 ± 0.1 |

<sup>a</sup> IC<sub>50</sub> values were determined using NanoBRET™ assay in a 10-point dose-response curve (duplicate measurements). <sup>b</sup> Average of three measurements plus standard deviation of the mean. <sup>c-d</sup> Literature K<sub>d</sub> values for staurosporine on SIK2 and SIK3.<sup>25,26</sup>

We also determined the crystal structure of compound 7 with MST3, revealing that the ortho-methoxyphenyl ring resulted in a small induced-fit movement of the side chain of Tyr113 by about 1.0 Å (PDB code 7B32) (**Supplementary Figure 6C**). Since substitutions in the solvent-exposed area of the ligand did not improve selectivity, we focused on the methyl group at position 6 of the pyridyl moiety. Removing this methyl group in compound **8** resulted in lower potency against SIK2 (IC_50_ = 777 ± 187 nM) and SIK3 (IC_50_ = 117 ± 25 nM), and a Δ*T*_m_ of 5.7 °C and 3.9 °C for MST3 and MST4, respectively, indicating that the methyl group is important for MST binding. The addition of a fluorine atom at position 3 in compound **9** increased the cellular potency against SIK2 and SIK3, but **9** still retained considerable activity against MST3/4 and PAK1 as judged by Δ*T*_m_ data. The selectivity profile of compound **9** was very similar to that of **6** (G-5555), showing no improved selectivity (**Supplementary Figure 7B-C** and **Supplementary Table 4**). Consistent with the data for compound **8** which demonstrated the importance of the methyl group at position **6** of the pyridine ring for potency against MST3/4, the non-methylated analogue 10 (MRIA9) showed reduced Δ*T*_m_ values for MST3/4 by 5.9 °C and 3.4 °C compared with G-5555. To our surprise, the fluorine substitution at position 3 seemed to compensate for the lack of the methyl group at position **6**, retaining high potency towards SIK2 (IC_50_ = 180 ± 40 nM) and SIK3 (IC_50_ = 127 ± 23 nM). The chlorine analogue **11** had a similar cellular potency in NanoBRET^TM^ assay (IC_50_ of 139 ± 110 nM), confirming that halogen substitution at position 3 can account for high potency towards both SIK2 and SIK3. The comparison of the crystal structures of MST3 in complex with **9** (PDB code 7B34) and MST3 in complex with **10** (MRIA9) (PDB code 7B31) revealed the molecular basis for the importance of the methyl group for MST3/4 inhibition potency. In the MST complexes, the methyl group of the ligand points towards Phe47 in the glycine-rich loop, forming additional hydrophobic interactions. In addition, the methyl group in position 6 increases the electron-donating properties of the pyridine nitrogen, thereby strengthening the hydrogen-bond network with Lys65 and Ser46 (**Supplementary Figure 7A**).

When introducing a methyl group in position 3 (compound **12**), the potency towards SIK2 and SIK3 increased to 65 ± 35 nM and 14 ±1 nM, respectively, but also the melting temperatures for MST4 and PAK1 increased, suggesting poorer selectivity. Considering the environment of the back pocket of SIK2, PAK1 and MST3/4, we expected that the addition of a bulkier group at position 3 might impair the activity for PAK1, which was confirmed with the 3-methoxy analogues **13** and **14**, which induced melting temperature shifts, Δ*T*_m_ s, of only 2.9 °C and 3.1 °C. However, only one of those two compounds with a 3-methoxy modification, **14**, retained potent SIK2 activity, with an IC_50_ of 600 ± 146 nM compared with an IC_50_ > 2 µM for **13**, again highlighting the importance of the methyl group at position 6 for the activity against both SIK2 and MST3/4. To our surprise, compound **13** maintained considerable potency against SIK3 but presented a similar kinome profile than G-5555 (data not shown). The crystal structure of MST3 in complex with **14** (PDB code 7B35) showed that the methoxy group of **14** is accommodated in the back pocket next to Leu86. Such an arrangement would result in steric hindrance upon binding to PAK1, which has a larger methionine residue at the corresponding position in the back pocket (**Supplementary Figure 8**).

These data highlight the crucial role of substitutions at positions 3 and 6 of the G-5555 core scaffold for modulating SIK2 and SIK3 activity. Interestingly, substitution of the methyl group at position 6 with a trifluoromethyl group (CF_3_) in compound **15** abrogated activity for SIK2 (IC_50_ > 4 µM) and lowered the potency for SIK3 by a factor of 20 (IC_50_ = 508 ± 4 nM), without affecting the thermal shifts for MST3/4 and PAK1. The STE off-target profile of the difluoromethyl analogue (CHF_2_), **16**, was similar to that of the parent molecule G-5555, with however the affinity for both SIK2 and SIK3 reduced about 2-fold, while maintaining potency for MST3/4 and PAK1. The co-crystal structure of 16 with MST3 (PDB code 7B33) revealed that the acidic hydrogen of the CHF_2_ group interacted with the side-chain hydroxyl of Ser46 in the glycine-rich loop (calculated distance of 2.4 Å between the hydroxyl oxygen and the acidic hydrogen of the CHF_2_ moiety). Ser46 in MST3 is replaced by an asparagine (Asn30) in SIK2, which may explain the differential effect of the CHF_2_ and CF_3_ moieties on binding affinity in the two kinases (**Supplementary Figure 9**).

### Selectivity profile of the panSIK inhibitor MRIA9

With our focused series of SIK inhibitors based on the G-5555 scaffold, we were able to significantly improve the cellular potency of the core scaffold against SIK2 and SIK3 but, importantly, also the selectivity over the main known off-targets of the parent compound. MRIA9 showed the best balance of SIK2/3 inhibition potency and activity against STE kinases, and we therefore screened this compound at 1 µM against 443 kinases in the radiometric ^33^PanQinase^®^ activity assay (*Reaction Biology*) (**Supplementary Table 5**), which returned an S-Score (35) of 0.018. In comparison, selectivity data of G-5555 measured at the same concentration resulted in an S-Score (35) of 0.12, confirming that our strategy was successful in designing a more selective SIK inhibitor (Figure 5B). Interestingly, a favorable balance between potency and selectivity was achieved not only by the addition of a group at position 3 of the pyridine ring, but also by the removal of the methyl group at position 6, which seems key for the increased selectivity of this series.

**Figure 5.**
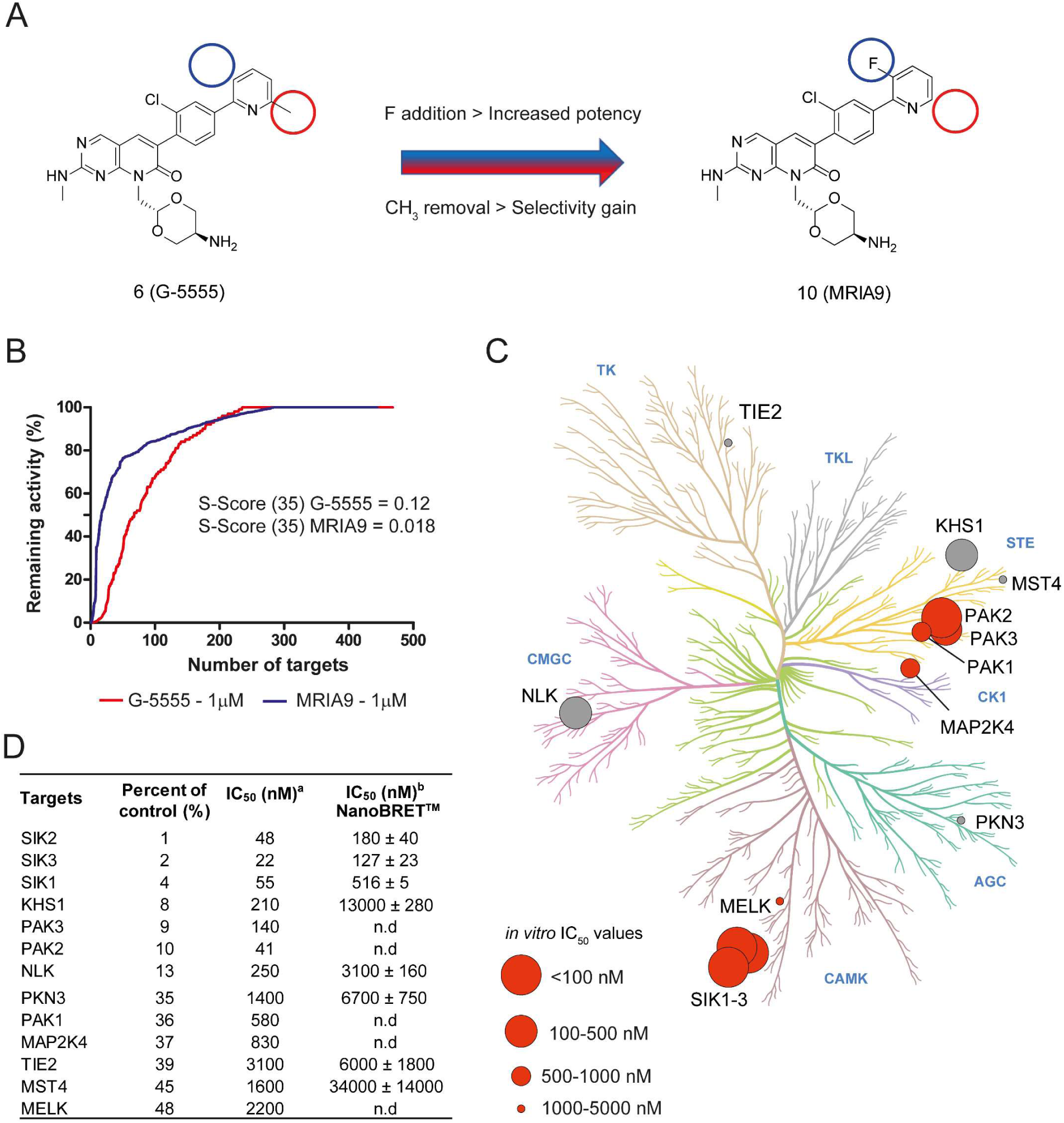
Selectivity profile of MRIA9 (10). (A) Optimization of pyrido[2,3-*d*]pyrimidin-7-one compound **6** (G-5555) by altering the substitution pattern on the pendant pyridine ring yielded compound **10** (MRIA9) with increased cellular potency and selectivity for SIK. (B) Comparison of selectivity data of MRIA9 (**10**) and G-5555 (**6**) against 443 kinases, including kinase mutants, as percentage of remaining activity. (C) Mapping of kinases inhibited by MRIA9 (10), shown on the kinase phylogenetic tree illustrated with the web-based tool CORAL.^28^ Inhibited targets are highlighted by circles, with the circle radius representing the *in vitro* IC_50_ values. The circles colored in gray highlight the off-targets that were only weakly inhibited in the NanoBRET™ assay. Activities on mutant kinases are not shown. Data displayed in this figure are included in **Supplementary Table 5**. (D) Results from a screening of 10 against 443 kinases (as percent of control < 50% at 1 μM) as well as *in vitro* and *in cellulo* IC_50_ values for the main kinase targets. ^a^IC_50_ values were determined by radiometric protein kinase assay (^33^PanQinase^®^ activity assay, by Reaction Biology). ^b^IC_50_ values were determined using NanoBRET™ assay in a 10-point dose-response curve (duplicate measurements).

Next, we followed up the kinome-wide screening by determining IC_50_ values in enzyme kinetic assays for all off-targets identified with inhibition values higher than 50% (Figure 5D). MRIA9 was equipotent towards SIK1-3 and PAK2, whereas it was 20 times less potent against PAK1, with an *in vitro* IC_50_ of only 580 nM. In addition, MRIA9 was only 5 times less potent against KHS1 and NLK compared with SIK inhibition *in vitro*. However, when comparing the results from cellular NanoBRET™ assays, MRIA9 had an IC_50_ of 516 ± 5 nM, 180 ± 40 nM and 127 ± 23 nM on SIK1, SIK2 and SIK3, respectively, and it did not target KHS1 and NLK in the cells. We also achieved selectivity over MST4 *in vitro* and *in cellulo* as expected based on our profiling data.

Unfortunately, we were unable to assess cellular PAK activity in the NanoBRET^TM^ assay due to a lack of suitable tracer molecules for this kinase. Nevertheless, our results show that MRIA9 is a highly potent dual pan-SIK and PAK2/3 inhibitor with excellent cellular activity, especially against SIK3. Together with a selective PAK inhibitor as a control, such as the highly selective allosteric inhibitor NVS-PAK1^27^, MRIA9 (**10**) can be used as a chemical probe for elucidating the complex biology of SIK kinases.

### Low concentration of MRIA9 inhibits the auto-phosphorylation activity of SIK2 *in vitro*

To further validate the specificity and the potency of MRIA9 over G-5555 in inhibiting SIK2, we performed an *in vitro* kinase auto-phosphorylation assay. Purified GST-tagged SIK2 was incubated with increasing concentrations of both inhibitors in the presence of radioactive [γ-^32^P]ATP. The auto-phosphorylation of SIK2 was monitored as an indicator of its activity. Exposing SIK2 to increasing concentrations of MRIA9 led to a gradual decrease in auto-phosphorylation activity. The half-maximal inhibitory concentration of SIK2 (IC_50_) was reached at a concentration of 0.3 nM of MRIA9 (10). G-5555 (6) did not show the same potency in this SIK2 auto-phosphorylation assay, and the highest inhibitor concentration tested (5 nM) (6) did only inhibit 57% of the activity of SIK2 (IC_50_ = 4.4 nM) (**Supplementary Figure 10**). These data confirmed the high potency of MRIA9 (10) in inhibiting SIK2 auto-phosphorylation activity.

### MRIA9 sensitizes ovarian cancer cells to paclitaxel treatment

After the characterization of MRIA9 as a potent SIK2 inhibitor with the lowest off-target profile characterized so far, we further validated the compound as a potential chemical tool for mechanistic and phenotypic studies. One intriguing observation for translational research of SIK inhibitors was that SIK2 depletion by RNAi sensitizes ovarian cancer cells to paclitaxel treatment.^13,14^ Based on these data, we next sought to investigate the therapeutic potential of MRIA9 (10) on ovarian cancer cell survival and Taxol sensitivity. To this end, the growth of the human ovarian cancer cell line SKOV3 was monitored in a long-term experiment (9 days) upon treatment with increasing concentrations of MRIA9 (0.5 µM to 5 µM) in the presence and absence of paclitaxel. Low concentrations of MRIA9 (0.5 and 1 µM) showed no significant effect on cell viability, and only high concentration (5 µM) reduced the viability somewhat compared with control cells (Figure 6A). Remarkably, a strong reduction in cell viability was observed when paclitaxel was combined with MRIA9 (Figure 6A). These results are in accordance with the profile of MRIA9 on the NCI-60 panel, which did not show a strong cytotoxic effect alone (**Supplementary Figure 11**). To investigate the apoptotic response of cells that is induced by the single treatment with MRIA9 or by the combinatorial treatment with paclitaxel and MRIA9, we monitored caspase 3/7 activation (Figure 6B). A moderate caspase-3/7 activation was observed in cells treated with paclitaxel alone. As expected, based on the limited effect of MRIA9 on cell survival, only the highest concentration used (5 µM) induced caspase activation. Caspase 3/7 activation was significantly enhanced, however, in combination treatment with paclitaxel and MRIA9 (Figure 6B).

**Figure 6.**
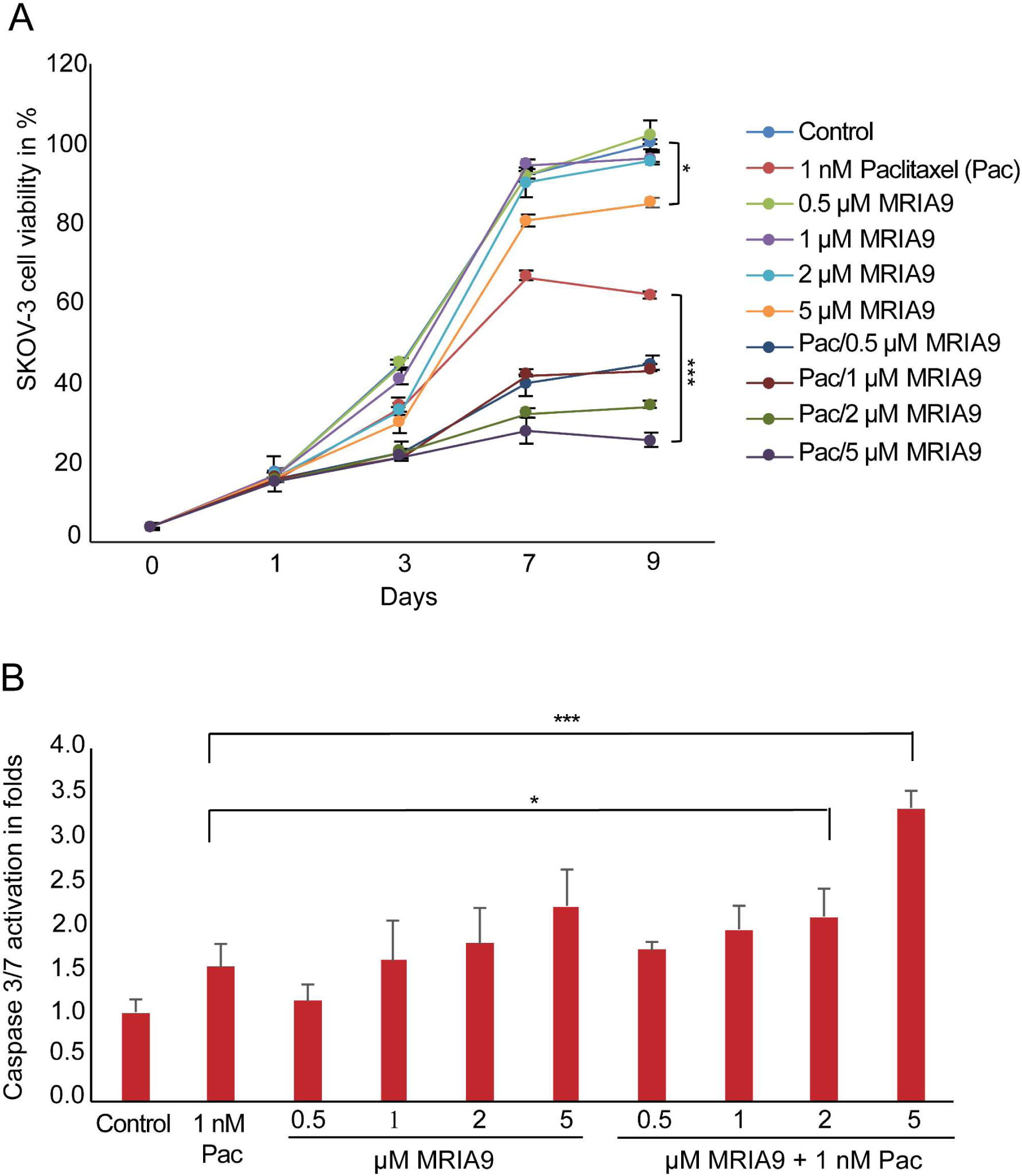
Incubation with MRIA9 sensitizes SKOV3 cells to paclitaxel treatment through inducing pronounced apoptosis. (A) SKOV3 cells were either treated with 1 nM paclitaxel, or increasing concentrations of MRIA9, or their combinations for 9 days. Cell viability was determined using the Cell-Titer-Blue Cell Viability Assay. Measurements were statistically significant by the Student’s t-test (* p ≤ 0.05; ** p ≤ 0.01). (B) Caspase-3/7-activity in whole lysates of SKOV3 cells treated with 1 nM paclitaxel, increasing concentrations of MRIA9 or their combinations was measured 48 h post-treatment using the Caspase-Glo® 3/7 Assay. Each bar graph represents the mean value ± SD (* p ≤ 0.05; *** p ≤ 0.001).

The 3D-spheroid system is widely used in studies involving the efficacy assessment of new small-molecule inhibitors *in vitro*. Therefore, we decided to investigate the effect of MRIA9-dependent inhibition of SIK2 on a 3D-spheroid model of HeLa cells. We evaluated the viability of HeLa cells derived spheroids by using the live/dead cell viability fluorescence-based assay upon 48 h-treatment with paclitaxel (2 nM) or MRIA9 (10) (5 µM) or the combination 2 nM paclitaxel / 5 µM MRIA9 (Figure 7). The inhibition of SIK2 using MRIA9 as a single treatment increased the fraction of dead cells compared with the depletion of SIK2 or with paclitaxel as a single agent (Figure 7). Remarkably, the combination paclitaxel/MRIA9 induced a robust increase of the fraction of dead HeLa cells compared with paclitaxel alone or even when compared with the combination paclitaxel and SIK2 siRNA after 48 h of treatment (p ≤0.001 for the combination paclitaxel (2 nM) and MRIA9 (5 µM)). These results indicate that the MRIA9-dependent inhibition of SIK2 enhanced the cytotoxicity of paclitaxel and argue for this combination therapy as already demonstrated in studies using less selective inhibitors with SIK activity and in genetic knockdown studies.^14^

**Figure 7.**
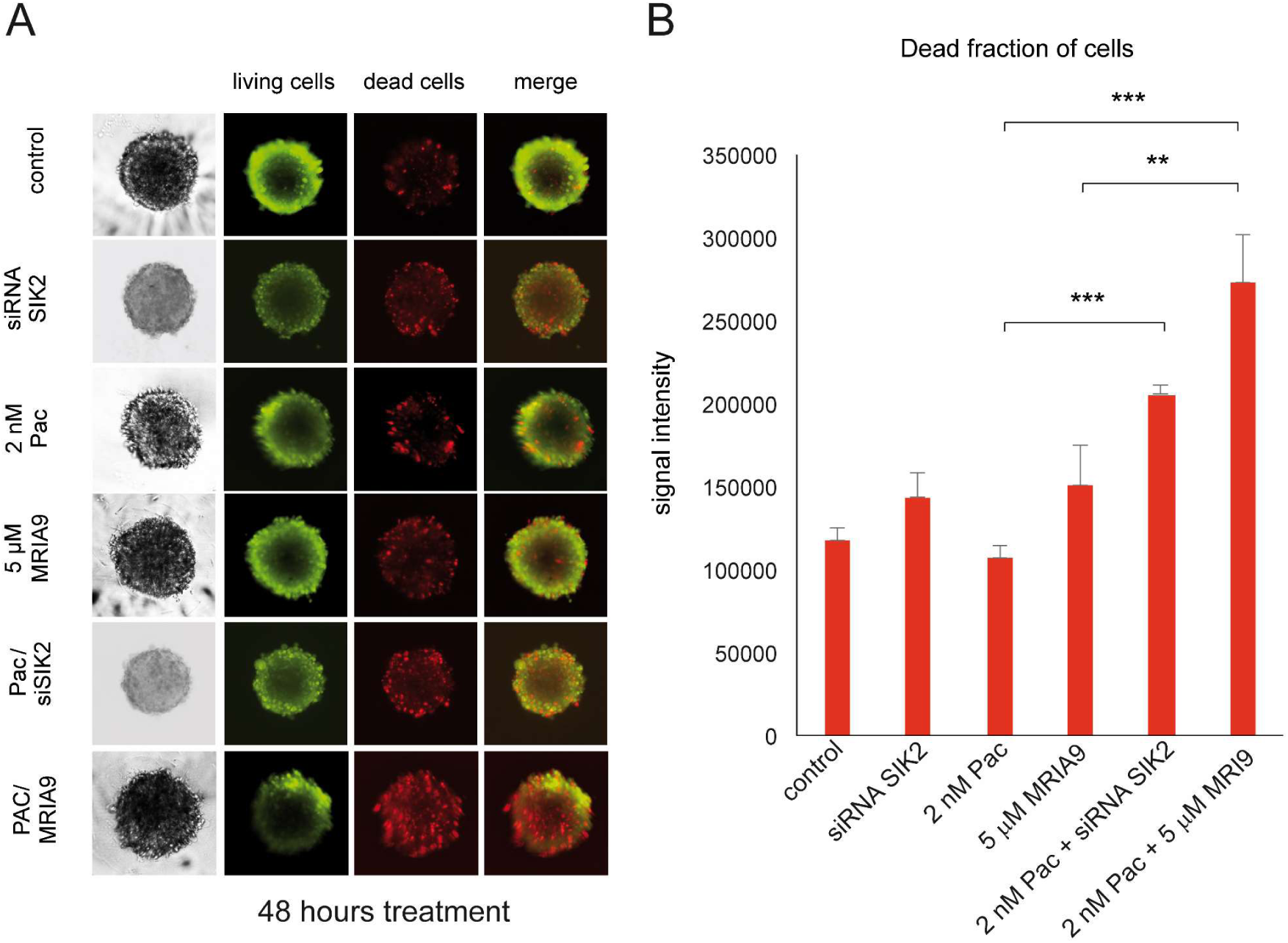
The combination of the SIK inhibitor MRIA9 with paclitaxel significantly enhances cell death in HeLa cells. (A) Spheroid cultures grown from HeLa cells were SIK2 depleted, treated with paclitaxel (2 nM), treated with the SIK inhibitor MRIA9 (5 µM), or treated with the combination paclitaxel/ 5 µM MRIA9 for 48 h. The spheroids were stained for immunofluorescence analysis with the live/dead viability/cytotoxicity kit. (B) The fraction of dead cells was quantified and presented as a bar graph. The results are presented as mean ± SD and statistically analyzed. ***p<0.001, **p<0.01

### Metabolic stability of MRIA9 and pharmacokinetic profile prediction

As a first evaluation of the pharmacokinetic properties of MRIA9 for further use in vivo, we analyzed the metabolic stability against an activated microsome mix derived from the liver of Sprague-Dawley rats as previously reported.^29^ MRIA9 (10) was incubated at 37 °C for 15, 30 and 60 min, and the amount of unmetabolized compound was determined by HPLC at these time points, including time zero. MRIA9 showed excellent metabolic stability after 1h incubation, with more than 90% of the compound detected (**Supplementary Figure 12**) and is therefore a good candidate for more detailed PK studies in vivo. **Supplementary Figure 12** also lists the results from different pharmacokinetics predictors for MRIA9 and the parent compound G-5555. According to these predictions, both G-5555 and MRIA9 are expected to show good cell permeability in Caco-2 and MDCK cells as well as high oral bioavailability (F = 75-80%). The pharmacokinetic behavior of G-5555 was experimentally assessed by Ndubaku et al., revealing high oral bioavailability in mice (F = 80%) and monkeys (F = 72%), which is in agreement with our predictions.

## CONCLUSION

The lack of available crystal structures of SIK proteins has impeded the development of potent and selective SIK inhibitors. We have therefore utilized an off-target approach to develop a new series of pyrido[2,3-*d*]pyrimidin-7-one inhibitors with SIK kinase activity. Starting from compound 6 (G-5555), we analyzed the binding mode of this inhibitor in off-targets, combining crystallographic data on MST3/4 and MD simulations, to rationally design a series of derivatives that exploit differences in the back-pocket region of SIK family proteins. From this approach, compound 10 (MRIA9) was the most promising chemical tool candidate, with both cellular potency and selectivity optimized. PAK2 was the only remaining off-target inhibited with the same potency *in vitro*. Further assessment of the cellular potency of MRIA9 towards the PAK subfamily will help delineate the usage of this new chemical tool in different phenotypic assays. Importantly, we have shown the potency of MRIA9 in sensitizing ovarian cancer cells to paclitaxel, suggesting that a combination therapy with paclitaxel and SIK inhibitors may result in improved therapeutic effects.

## EXPERIMENTAL SECTION

### Chemistry

The synthesis of compounds **7-18** will be explained in the following. Further information regarding the analytical data for compounds **7-37** can be found in the supporting information. All commercial chemicals were purchased from common suppliers in reagent grade and used without further purification. Reactions were proceeded under argon atmosphere. For compound purification by flash chromatography, a puriFlash^®^ XS 420 device with a UV-VIS multi-wave detector (200-400 nm) from *Interchim* with prepacked normal phase PF-SIHP silica columns with particle sizes of 30 µm (*Interchim*) was used. All compounds synthesized were characterized by ^1^H and ^13^C NMR and mass spectrometry (ESI +/-). In addition, final compounds were identified by HRMS and their purity was evaluated by HPLC. ^1^H and ^13^C NMR spectra were measured on a DPX250, an AV400 or an AV500 HD AvanceIII spectrometer from *Bruker Corporation*. Chemical shifts (*δ*) are reported in parts per million (ppm). DMSO-d_6_ was used as a solvent, and the spectra were calibrated to the solvent signal: 2.50 ppm (^1^H NMR) or 39.52 ppm (^13^C NMR). Mass spectra (ESI +/-) were measured on a Surveyor MSQ device from *ThermoFisher*. High resolution mass spectra (HRMS) were obtained on a MALDI LTQ Orbitrap XL from *ThermoScientific*. Preparative purification by HPLC was carried out on an *Agilent* 1260 Infinity II device using an Eclipse XDB-C18 (*Agilent*, 21.2 x 250mm, 7µm) reversed-phase column. A suitable gradient (flow rate 21 ml/min.) was used, with 0.1% TFA in water (A) and 0.1% TFA in acetonitrile (B) as a mobile phase. Determination of the compound purity by HPLC was carried out on the same *Agilent* 1260 Infinity II device, together with HPLC-MS measurements using an LC/MSD device (G6125B, ESI pos. 100-1000). The compounds were analyzed on an Eclipse XDB-C18 (*Agilent*, 4.6 x 250mm, 5µm) reversed-phase column, using 0.1% TFA in water (A) and 0.1% TFA in acetonitrile (B) as a mobile phase. The following gradient was used: 0 min 2% B - 2 min 2% B - 10 min 98% B – 15 min 98% B - 17 min 2% B - 19 min 2% B (flow rate of 1 mL/min.). UV-detection was performed at 254 and 280 nm. All final compounds showed > 95% purity.

### General Procedure I for Suzuki cross-coupling reaction

6-(2-chloro-4-(4,4,5,5-tetramethyl-1,3,2-dioxaborolan-2-yl)phenyl)-2-(methylthio)pyrido[2,3-*d*]pyrimidin-7-ol (22) (1.0 eq), potassium acetate (2.0 eq), [1,1′-Bis(diphenylphosphino)ferrocene] dichloropalladium(II) (0.1 eq) and the corresponding pyridine derivative (1.0 eq or 1.1 eq) were dissolved in a mixture of dioxane/water (2:1; 12 mL) and stirred at 107 °C for 18 hours. The solvent was evaporated under reduced pressure and the remaining residue was purified by column chromatography on silica gel using cyclohexane/ethyl acetate as an eluent.

### General procedure II for nucleophilic substitution

The corresponding 6-[2-chloro-4-(3-R1-4-R2-6-R3-pyridin-2-yl)phenyl]2-(methylthio)pyrido[2,3-*d*]pyrimidin-7-ol (1.0 eq), obtained from general procedure I, 2-((2r,5r)-2-(bromomethyl)-1,3-dioxan-5-yl)isoindoline-1,3-dione (1.5 eq) and cesium carbonate (3.0 eq) were dissolved in anhydrous *N,N*-dimethylformamide (DMF) (10 mL) and stirred at 100 °C for 18 hours. The solvent was evaporated under reduced pressure. To remove unreacted starting material, a column chromatography on silica gel was performed using DCM/MeOH as an eluent. A mixture of the corresponding 2-((2r,5r)-2-((6-(2-chloro-4-(3-R1-4-R2-6-R3-pyridin-2-yl)phenyl)-2-(methylthio)-7-oxopyrido[2,3-*d*]pyrimidin-8(7H)-yl)methyl)-1,3-dioxan-5-yl)isoindoline-1,3-dione and 2-(((2r,5r)-2-((6-(2-chloro-4-(3-R1-4-R2-6-R3-pyridin-2-yl)phenyl)-2-(methylthio)-7-oxopyrido[2,3-*d*]pyrimidin-8(7H)-yl)methyl)-1,3-dioxan-5-yl)-carbamoyl)benzoic acid was obtained and used in the next step without further separation.

### General procedure III for oxidation

The corresponding mixture of (2-((2r,5r)-2-((6-(2-chloro-4-(3-R1-4-R2-6-R3-pyridin-2-yl)phenyl)-2-(methylthio)-7-oxopyrido[2,3-*d*]pyrimidin-8(7H)-yl)methyl)-1,3-dioxan-5-yl)isoindoline-1,3-dione and 2-(((2r,5r)-2-((6-(2-chloro-4-(3-R1-4-R2-6-R3-pyridin-2-yl)phenyl)-2-(methylthio)-7-oxopyrido[2,3-*d*]pyrimidin-8(7H)-yl)methyl)-1,3-dioxan-5-yl)-carbamoyl)benzoic acid, obtained from general procedure II (1.0 eq) was dissolved in anhydrous dichloromethane (DCM) (10 mL). M-chloroperoxybenzoic acid (*m-*CPBA) (≤ 77%, 2.1 eq) was added in one portion, and the reaction solution was stirred at room temperature for three hours. The remaining *m-*CPBA was quenched by adding saturated NaHCO_3_-solution (10 mL). The aqueous layer was extracted with a mixture of DCM/MeOH (4:1; 3 x 15 mL) and DCM (15 mL). The combined organic layers were dried over Na_2_SO_4_ and the solvent was evaporated under reduced pressure. The remaining residue, consisting the corresponding sulfoxide and sulfone product, was used in the next step without further purification.

### General procedure IV for nucleophilic aromatic substitution

The remaining residue of the corresponding product of general procedure III (1.0 eq) was diluted in ethanol (10 mL), and a solution of methylamine in THF (2M, 4.0 eq) and *N,N*-diisopropylethylamine (DIPEA) (5.0 eq) were added. The reaction solution was stirred at 85 °C for 17 hours. After the reaction was finished, the solvent was evaporated under reduced pressure. The remaining residue was purified by HPLC chromatography on silica gel (H_2_O/ACN; 0.1% TFA) to obtain the corresponding 8-(((2r,5r)-5-amino-1,3-dioxan-2-yl)methyl)-6-(2-chloro-4-(3-R1-4-R2-6-R3-pyridin-2-yl)phenyl)-2-(methylamino)pyrido[2,3-*d*]pyrimidin-7(8*H*)-one product as a TFA salt.

#### Ethyl 2-(4-bromo-2-chloro-phenyl)acetate (**20**)

2-(4-bromo-2-chloro-phenyl)-acetic acid (**19**; 4.00 g, 16.0 mmol) was dissolved in ethanol (80 mL) and cooled to 0 °C. Sulfuric acid (1.57 g, 16.0 mmol) was added, and the reaction solution was allowed to warm to room temperature. The reaction was stirred at 100 °C for 15 hours. Afterwards the solvent was evaporated under reduced pressure, and the remaining residue was dissolved in ethyl acetate (30 mL). The organic layer was washed with saturated NaHCO_3_-solution (2 x 20 mL) and brine (20 mL). The aqueous layer was extracted with ethyl acetate (20 mL), and the combined organic layers were dried over MgSO_4_. The solvent was evaporated under reduced pressure to afford **20** as a yellow liquid with a yield of 93% (4.15 g). ^1^H NMR (500 MHz, DMSO-d_6_, 300K): *δ* = 7.73 (d, *J* = 2.0 Hz, 1H), 7.54 (dd, *J* = 8.2, 2.0 Hz, 1H), 7.38 (d, *J* = 8.2 Hz, 1H), 4.09 (q, *J* = 7.1 Hz, 2H), 3.79 (s, 2H), 1.18 (t, *J* = 7.1 Hz, 3H) ppm. ^13^C NMR (126 MHz, DMSO-d_6_, 300K): *δ* = 169.6, 134.9, 133.7, 132.4, 131.2, 130.3, 120.7, 60.5, 38.0, 14.0 ppm. MS-ESI+ (m/z) [fragment+H]^+^; calculated: 205.0, found: 205.0.

#### 6-(4-Bromo-2-chlorophenyl)-2-(methylthio)pyrido[2,3-*d*]pyrimidin-7-ol (**21**)

4-amino-2-(methylthio)pyrimidine-5-carbaldehyde (1.70 g, 10.1 mmol), **20** (3.07 g, 11.1 mmol) and potassium carbonate (4.17 g, 30.1 mmol) were dissolved in anhydrous DMF (70 mL) and stirred at 120 °C for 16 hours. Water (300 mL) was added to the reaction solution, leading to the precipitation of **21** as a pale-yellow solid. After filtration, the aqueous layer was extracted with ethyl acetate (3 x 30 mL), and the combined organic layers were dried over MgSO_4_. The solvent was evaporated under reduced pressure, and the remaining residue was recrystallized from acetone. A combined yield of 93% (3.57 g) was achieved. ^1^H NMR (500 MHz, DMSO-d_6_, 300K): *δ* = 12.69 (s, 1H), 8.88 (s, 1H), 7.95 (s, 1H), 7.83 (d, *J* = 1.9 Hz, 1H), 7.64 (dd, *J* = 8.2, 2.0 Hz, 1H), 7.38 (d, *J* = 8.2 Hz, 1H), 2.58 (s, 3H) ppm. ^13^C NMR (126 MHz, DMSO-d_6_, 300K): *δ* = 172.2, 161.5, 157.0, 154.5, 136.6, 134.3, 134.1, 133.3, 131.5, 130.7, 130.2, 121.9, 108.8, 13.7 ppm.

#### 6-(2-Chloro-4-(4,4,5,5-tetramethyl-1,3,2-dioxaborolan-2-yl)phenyl)-2-(methylthio)-pyrido[2,3-*d*]pyrimidin-7-ol (**22**)

**21** (200 mg, 0.52 mmol), bis(pinacolato)diboron (173 mg, 0.68 mmol), potassium acetate (154 mg, 1.57 mmol) and [1,1’-bis(diphenylphosphino) ferrocene]dichloropalladium(II) in complex with DCM (30 mg, 0.04 mmol) were dissolved in anhydrous dioxane (10 mL) and stirred at 115 °C for 18 hours. The reaction solution was filtered over a pad of Celite^®^. Ethyl acetate (20 mL) was added, and the organic layer was washed with brine (20 mL). The aqueous layer was extracted with ethyl acetate (2 x 20 mL), and the combined organic layers were dried over Na_2_SO_4_. The solvent was evaporated under reduced pressure, and the remaining residue was purified by flash chromatography on silica gel (cyclohexane/ethyl acetate 8:1 - 2:1). Recrystallization from DCM/cyclohexane was performed to obtain **21** as a pale-yellow solid with a yield of 89% (200 mg). ^1^H NMR (500 MHz, DMSO-d_6_, 300K): *δ* = 12.67 (s, 1H), 8.91 (s, 1H), 7.96 (s, 1H), 7.71 (s, 1H), 7.67 (dd, *J* = 7.5, 1.0 Hz, 1H), 7.44 (d, *J* = 7.5 Hz, 1H), 2.59 (s, 3H), 1.32 (s, 12H) ppm. ^13^C NMR (126 MHz, DMSO-d_6_, 300K): *δ* = 172.2, 161.2, 157.0, 154.2, 137.7, 136.5, 134.5, 132.8, 132.7, 131.6, 131.5, 130.7, 108.8, 84.2, 24.6, 13.7 ppm. MS-ESI+ (m/z) [M+H]^+^; calculated: 430.1, found: 430.6.

#### 2-(1,3-Dihydroxypropan-2-yl)isoindoline-1,3-dione (**24**)

Serinol (**23**) (500 mg, 5.49 mmol) and phthalic anhydride (800 mg, 5.49 mmol) were dissolved in anhydrous DMF (5 mL) and stirred at 90 °C for 4 hours. The solvent was evaporated under reduced pressure, leading to the formation of a yellow solid. The solid was recrystallized from 2-propanol/cyclohexane to obtain **24** as a white solid with a yield of 87% (1.05 g). ^1^H NMR (500 MHz, DMSO-d_6_, 300K): *δ* = 7.86-7.82 (m, 4H), 4.87 (t, *J* = 6.0 Hz, 2H), 4.27-4.21 (m, 1H), 3.83-3.77 (m, 2H), 3.68-3.64 (m, 2H) ppm. ^13^C NMR (126 MHz, DMSO-d_6_, 300K): *δ* = 168.4, 134.2, 131.7, 122.8, 58.3, 56.5 ppm. MS-ESI+ (m/z) [M+Na]^+^; calculated: 244.1, found: 244.1.

#### 2-((2r,5r)-2-(Bromomethyl)-1,3-dioxan-5-yl)isoindoline-1,3-dione (**25**)

**24** (2.50 g, 11.3 mmol), 2-bromo-1,1’-diethoxy-ethane (1.20 g, 13.6 mmol) and p-toluenesulfonic acid monohydrate (0.43 g, 2.3 mmol) were dissolved in 60 mL toluene and stirred at 115 °C for 24 hours. The organic layer was washed with saturated NaHCO_3_ solution (2 x 50 mL) and brine (50 mL). The aqueous layer was extracted with ethyl acetate (3 x 20 mL), and the combined organic layers were dried over Na_2_SO_4_. The solvent was evaporated under reduced pressure, and the remaining residue was purified by flash chromatography on silica gel (cyclohexane/ethyl acetate 6:1 - 2:1) to obtain **25** as a white solid with a yield of 73% (2.70 g). ^1^H NMR (400 MHz, DMSO-d_6_, 300K): *δ* = 7.88-7.84 (m, 4H), 4.84 (t, *J* = 4.0 Hz, 1H), 4.31-4.30 (m, 3H), 4.11-4.09 (m, 2H), 3.51 (d, *J* = 4.1 Hz, 2H) ppm. ^13^C NMR (100 MHz, DMSO-d_6_, 300K): *δ* = 167.5, 134.5, 131.4, 123.1, 98.5, 65.8, 43.6, 32.2 ppm. MS-ESI+ (m/z) [fragment+K]^+^; calculated: 382.0, found: 382.3.

#### 6-(2-Chloro-4-(6-methylpyridin-2-yl)phenyl)-2-(methylthio)pyrido[2,3-*d*]pyrimidin-7-ol (**26**)

Compound **26** was prepared according to general procedure I, using 2-bromo-6-methyl-pyridine (1.0 eq). **26** was obtained as a white solid with a yield of 80% (88 mg). ^1^H NMR (500 MHz, DMSO-d_6_, 300K): *δ* = 12.69 (s, 1H), 8.92 (s, 1H), 8.24 (d, *J* = 1.7 Hz, 1H), 8.11 (dd, *J* = 8.0, 1.7 Hz, 1H), 8.02 (s, 1H), 7.88 (d, *J* = 7.8 Hz, 1H), 7.82 (t, *J* = 7.7 Hz, 1H), 7.53 (d, *J* = 8.0 Hz, 1H), 7.29 (d, *J* = 7.6 Hz, 1H), 2.59 (s, 3H), 2.57 (s, 3H) ppm. ^13^C NMR (126 MHz, DMSO-d_6_, 300K): *δ* = 172.2, 161.4, 158.1, 157.0, 154.2, 153.3, 140.4, 137.7, 136.6, 135.0, 133.4, 132.1, 131.4, 126.9, 124.9, 122.8, 117.7, 108.8, 24.3, 13.7 ppm. MS-ESI+ (m/z) [M+H]^+^; calculated: 395.1, found: 395.0.

#### 6-(2-Chloro-4-(pyridin-2-yl)phenyl)-2-(methylthio)pyrido[2,3-*d*]pyrimidin-7-ol (**27**)

Compound **27** was prepared according to general procedure I, using 2-bromo-pyridine (1.0 eq). **27** was obtained as a white solid with a yield of 20% (117 mg). ^1^H NMR (500 MHz, DMSO-d_6_, 300K): *δ* = 12.69 (s, 1H), 8.92 (s, 1H), 8.71 (dd, *J* = 4.7, 0.8 Hz, 1H), 8.26 (d, *J* = 1.7 Hz, 1H), 8.13 (dd, *J* = 8.0, 1.7 Hz, 1H), 8.09 (d, *J* = 8.0 Hz, 1H), 8.02 (s, 1H), 7.94 (dt, *J* = 7.7, 1.8 Hz, 1H), 7.55 (d, *J* = 8.0 Hz, 1H), 7.44-7.41 (m, 1H), 2.59 (s, 3H) ppm. ^13^C NMR (126 MHz, DMSO-d_6_, 300K): *δ* = 172.2, 161.4, 157.0, 154.2, 154.0, 149.7, 140.3, 137.5, 136.6, 135.2, 133.5, 132.2, 131.4, 127.0, 124.9, 123.4, 120.7, 108.8, 13.7 ppm. MS-ESI+ (m/z) [M+H]^+^; calculated: 381.1, found: 381.0.

#### 6-(2-Chloro-4-(3-fluoro-6-methylpyridin-2-yl)phenyl)-2-(methylthio)-pyrido[2,3-*d*]-pyrimidin-7-ol (**28**)

Compound **28** was prepared according to general procedure I, using 2-bromo-3-fluoro-6-methyl-pyridine (1.0 eq). **28** was obtained as a white solid with a yield of 60% (173 mg). ^1^H NMR (500 MHz, DMSO-d_6_, 300K): *δ* = 12.70 (s, 1H), 8.92 (s, 1H), 8.04 (s, 2H), 7.95 (s, 1H), 7.77 (s, 1H), 7.57 (s, 1H), 7.39 (s, 1H), 2.59 (s, 3H), 2.55 (s, 3H) ppm. ^13^C NMR (126 MHz, DMSO-d_6_, 300K): *δ* = 172.2, 161.3, 159.4, 157.0, 154.8, 154.3, 141.4, 136.6, 135.4, 133.1, 131.9, 131.3, 128.8, 127.0, 125.4, 125.2, 124.7, 108.8, 23.5, 13.7 ppm. MS-ESI+ (m/z) [M+H]^+^; calculated: 413.1, found: 413.0.

#### 6-(2-Chloro-4-(3-fluoropyridin-2-yl)phenyl)-2-(methylthio)pyrido[2,3-*d*]pyrimidin-7-ol (**29**)

Compound **29** was prepared according to general procedure I, using 2-bromo-3-fluoro-pyridine (1.0 eq). **29** was obtained as a white solid with a yield of 94% (260 mg). ^1^H NMR (500 MHz, DMSO-d_6_, 300K): *δ* = 12.70 (s, 1H), 8.92 (s, 1H), 8.60 (d, *J* = 1.7 Hz, 1H), 8.06 (s, 1H), 8.04 (s, 1H), 7.96 (d, *J* = 8.0 Hz, 1H), 7.93-7.87 (m, 1H), 7.60-7.53 (m, 2H), 2.59 (s, 3H) ppm. ^13^C NMR (126 MHz, DMSO-d_6_, 300K): *δ* = 172.7, 161.8, 158.8, 157.5, 156.7, 154.7, 146.5, 143.3, 137.1, 136.9, 136.0, 133.6, 132.4, 131.7, 129.3, 127.5, 125.8, 125.6, 125.4, 109.3, 14.2 ppm. MS-ESI-(m/z) [M-H]^-^; calculated: 397.0, found: 397.2.

#### 6-(2-Chloro-4-(3-chloropyridin-2-yl)phenyl)-2-(methylthio)pyrido[2,3-*d*]pyrimidin-7-ol (**30**)

Compound **30** was prepared according to general procedure I, using 2-bromo-3-chloro-pyridine (1.0 eq). **30** was obtained as a white solid with a yield of 69% (200 mg). ^1^H NMR (500 MHz, DMSO-d_6_, 300K): *δ* = 12.71 (s, 1H), 8.91 (s, 1H), 8.68 (dd, *J* = 4.7, 1.1 Hz, 1H), 8.11 (d, *J* = 8.1 Hz, 1H), 8.05 (s, 1H), 7.84 (d, *J* = 1.3 Hz, 1H), 7.75 (dd, *J* = 7.9, 1.4 Hz, 1H), 7.55 (d, *J* = 7.9 Hz, 1H), 7.51 (dd, *J* = 8.1, 4.6 Hz, 1H), 2.59 (s, 3H) ppm. ^13^C NMR (126 MHz, DMSO-d_6_, 300K): *δ* = 172.2, 161.3, 157.0, 154.2, 153.6, 148.2, 139.4, 138.6, 136.7, 135.2, 132.6, 131.4, 131.3, 129.8, 129.3, 127.9, 124.6, 108.8, 13.7 ppm. MS-ESI+ (m/z) [M+Na]^+^; calculated: 437.0, found: 437.0.

#### 6-(2-Chloro-4-(3-methylpyridin-2-yl)phenyl)-2-(methylthio)pyrido[2,3-*d*]pyrimidin-7-ol (**31**)

Compound **31** was prepared according to general procedure I, using 2-bromo-3-methyl-pyridine (1.0 eq). **31** was obtained as a white solid with a yield of 73% (200 mg). ^1^H NMR (500 MHz, DMSO-d_6_, 300K): *δ* = 12.68 (s, 1H), 8.92 (s, 1H), 8.53 (dd, *J* = 4.7, 1.1 Hz, 1H), 8.04 (s, 1H), 7.77 (dd, *J* = 7.7, 0.8 Hz, 1H), 7.72 (d, *J* = 1.7 Hz, 1H), 7.62 (dd, *J* = 7.9, 4.7 Hz, 1H), 7.52 (d, *J* = 7.9 Hz, 1H), 7.35 (dd, *J* = 7.7, 4.7 Hz, 1H), 2.60 (s, 3H), 2.40 (s, 3H) ppm. ^13^C NMR (126 MHz, DMSO-d_6_, 300K): *δ* = 172.1, 161.4, 156.9, 155.7, 154.2, 147.0, 141.9, 138.9, 136.6, 134.2, 132.6, 131.4, 131.3, 130.8, 129.6, 127.6, 122.9, 108.8, 19.6, 13.7 ppm. MS-ESI+ (m/z) [M+H]^+^; calculated: 395.1, found: 395.1.

#### 6-(2-Chloro-4-(3-methoxypyridin-2-yl)phenyl)-2-(methylthio)pyrido[2,3-*d*]pyrimidin-7-ol (**32**)

Compound **32** was prepared according to general procedure I, using 2-bromo-3-methoxy-pyridine (1.0 eq). **32** was obtained as a white solid with a yield of 70% (200 mg). ^1^H NMR (500 MHz, DMSO-d_6_, 300K): *δ* = 12.68 (s, 1H), 8.92 (s, 1H), 8.31 (dd, *J* = 4.6, 0.9 Hz, 1H), 8.06-8.00 (m, 2H), 7.94 (dd, *J* = 8.0, 1.5 Hz, 1H), 7.67 (d, *J* = 8.3 Hz, 1H), 7.50 (d, *J* = 8.0 Hz, 1H), 7.48-7.45 (m, 1H), 3.92 (s, 3H), 2.59 (s, 3H) ppm. ^13^C NMR (126 MHz, DMSO-d_6_, 300K): *δ* = 172.1, 161.4, 156.9, 154.2, 153.8, 144.0, 140.8, 138.8, 136.5, 134.5, 132.4, 131.5, 131.3, 129.5, 127.7, 124.5, 120.3, 108.8, 55.9, 13.7 ppm. MS-ESI+ (m/z) [M+H]^+^; calculated: 411.1, found: 411.0.

#### 6-(2-Chloro-4-(3-methoxy-6-methylpyridin-2-yl)phenyl)-2-(methylthio)pyrido[2,3-*d*]-pyrimidin-7-ol (**33**)

Compound **33** was prepared according to general procedure I, using 2-bromo-3-methoxy-6-methyl-pyridine (1.1 eq). **33** was obtained as a white solid with a yield of 78% (230 mg). ^1^H NMR (500 MHz, DMSO-d_6_, 300K): *δ* = 12.67 (s, 1H), 8.92 (s, 1H), 8.01 (d, *J* = 1.6 Hz, 1H), 8.00 (s, 1H), 7.93 (dd, *J* = 8.0, 1.7 Hz, 1H), 7.53 (d, *J* = 8.5 Hz, 1H), 7.47 (d, *J* = 8.0 Hz, 1H), 7.27 (d, *J* = 8.5 Hz, 1H), 3.87 (s, 3H), 2.59 (s, 3H), 2.48 (s, 3H) ppm. ^13^C NMR (126 MHz, DMSO-d_6_, 300K): *δ* = 172.1, 161.4, 156.9, 154.2, 151.7, 149.1, 143.0, 139.3, 136.4, 134.2, 132.3, 131.6, 131.2, 129.3, 127.6, 123.5, 120.7, 108.8, 55.8, 23.1, 13.7 ppm. MS-ESI+(m/z) [M+H]^+^; calculated: 425.1, found: 425.0.

#### 6-(2-Chloro-4-(6-(trifluoromethyl)pyridin-2-yl)phenyl)-2-(methylthio)pyrido[2,3-*d*]-pyrimidin-7-ol (**34**)

Compound **34** was prepared according to general procedure I, using 2-bromo-6-trifluoromethyl-pyridine (1.1 eq). **34** was obtained as a yellow solid with a yield of 96% (300 mg). ^1^H NMR (500 MHz, DMSO-d_6_, 300K): *δ* = 12.70 (s, 1H), 8.93 (s, 1H), 8.43 (d, *J* = 8.0 Hz, 1H), 8.28 (d, *J* = 1.7 Hz, 1H), 8.24 (t, *J* = 7.9 Hz, 1H), 8.18 (dd, *J* = 8.0, 1.7 Hz, 1H), 8.03 (s, 1H), 7.93 (d, *J* = 7.7 Hz, 1H), 7.61 (d, *J* = 8.0 Hz, 1H), 2.59 (s, 3H) ppm. ^13^C NMR (126 MHz, DMSO-d_6_, 300K): *δ* = 172.3, 161.3, 157.0, 154.8, 154.3, 146.9, 146.7, 146.4, 146.1, 134.0, 138.6, 136.7, 136.2, 133.7, 132.4, 131.2, 127.3, 125.4, 124.8, 124.3, 122.6, 120.5, 120.0, 118.3, 108.8, 13.7 ppm. MS-ESI-(m/z) [M-H]^-^; calculated: 447.0, found: 447.0.

#### 6-(2-Chloro-4-(6-(difluoromethyl)pyridin-2-yl)phenyl)-2-(methylthio)pyrido[2,3-*d*]pyrimidin-7-ol (**35**)

Compound **35** was prepared according to general procedure I, using 2-bromo-6-difluoromethyl-pyridine (1.1 eq). **35** was obtained as a pale yellow solid with a yield of 87% (260 mg). ^1^H NMR (500 MHz, DMSO-d_6_, 300K): *δ* = 12.70 (s, 1H), 8.93 (s, 1H), 8.29-8.28 (m, 2H), 8.18-8.13 (m, 2H), 8.03 (s, 1H), 7.73 (d, *J* = 7.7 Hz, 1H), 7.59 (d, *J* = 8.0 Hz, 1H), 7.07 (t, *J* = 54.9 Hz, 1H), 2.59 (s, 3H) ppm. ^13^C NMR (126 MHz, DMSO-d_6_, 300K): *δ* = 172.2, 161.3, 157.0, 154.2, 154.2, 152.3, 152.1, 151.9, 139.4, 139.2, 136.6, 135.8, 133.7, 132.3, 131.3, 127.2, 125.2, 122.7, 119.8, 115.7, 113.8, 111.9, 108.8, 13.7 ppm. MS-ESI+ (m/z) [M+K]^+^; calculated: 469.0, found: 468.8.

#### 6-(2-Chloro-4-(6-methoxypyridin-2-yl)phenyl)-2-(methylthio)pyrido[2,3-*d*]pyrimidin-7-ol (**36**)

Compound **36** was prepared according to general procedure I, using 2-bromo-6-methoxy-pyridine (1.0 eq). (**36**) was obtained as a white solid with a yield of 98% (295 mg). ^1^H NMR (500 MHz, DMSO-d_6_, 300K): *δ* = 12.67 (s, 1H), 8.92 (s, 1H), 8.24 (d, *J* = 1.7 Hz, 1H), 8.12 (dd, *J* = 8.0, 1.7 Hz, 1H), 8.01 (s, 1H), 7.85-7.82 (m, 1H), 7.68 (d, *J* = 7.4 Hz, 1H), 7.54 (d, *J* = 8.0 Hz, 1H), 6.85 (d, *J* = 8.2 Hz, 1H), 3.98 (s, 3H), 2.59 (s, 3H) ppm. ^13^C NMR (126 MHz, DMSO-d_6_, 300K): *δ* = 172.2, 163.3, 161.3, 156.9, 154.2, 151.7, 140.2, 140.0, 136.5, 135.1, 133.4, 132.1, 131.4, 126.8, 124.9, 113.5, 110.3, 108.8, 53.0, 13.7 ppm. MS-ESI+ (m/z) [M+Na]^+^; calculated: 433.1, found: 433.1.

#### 6-(2-Chloro-4-(4,6-dimethylpyridin-2-yl)phenyl)-2-(methylthio)pyrido[2,3-*d*]pyrimidin-7-ol (**37**)

Compound **37** was prepared according to general procedure I, using 2-bromo-4,6-dimethylpyridine (1.0 eq). (**37**) was obtained as a white solid with a yield of 74% (210 mg). ^1^H NMR (500 MHz, DMSO-d_6_, 300K): *δ* = 12.66 (s, 1H), 8.91 (s, 1H), 8.23 (d, *J* = 1.4 Hz, 1H), 8.10 (dd, *J* = 8.0, 1.5 Hz, 1H), 8.01 (s, 1H), 7.74 (s, 1H), 7.52 (d, *J* = 8.0 Hz, 1H), 7.12 (s, 1H), 2.59 (s, 3H), 2.52 (s, 3H), 2.37 (s, 3H) ppm. ^13^C NMR (126 MHz, DMSO-d_6_, 300K): *δ* = 172.1, 161.4, 157.7, 156.9, 154.3, 153.2, 148.3, 140.5, 136.5, 134.9, 133.3, 132.0, 131.4, 126.9, 124.9, 123.5, 118.6, 108.8, 24.1, 20.5, 13.7 ppm. MS-ESI+ (m/z) [M+H]^+^; calculated: 409.1, found: 409.1.

#### 8-(((2r,5r)-5-Amino-1,3-dioxan-2-yl)methyl)-6-(2-chloro-4-(6-methylpyridin-2-yl)phenyl)-2-((2-methoxyphenyl)amino)pyrido[2,3-*d*]pyrimidin-7(8H)-one (**7**)

Compound **7** was prepared according to general procedures II and III. The remaining residue of general procedure III containing the corresponding sulfoxide/sulfone product (1.0 eq) and o-anisidine (4.0 eq) were dissolved in ethanol (5 mL). HCl (1M, 0.1 mL) was added, and the reaction solution was stirred at 100 °C for 17 hours. Water (5 mL) was added to the reaction solution, and the aqueous layer was extracted with a mixture of DCM/MeOH (4:1, 3 x 10 mL) and DCM (10 mL). The combined organic layers were dried over Na_2_SO_4_, and the solvent was evaporated under reduced pressure. The remaining residue was purified by HPLC chromatography on silica gel (H_2_O/ACN; 0.1% TFA) to obtain 7 as an orange solid with a yield of 13% (20 mg) over 3 steps. ^1^H NMR (500 MHz, DMSO-d_6_, 300K): *δ* = 8.84 (d, *J* = 5.3 Hz, 1H), 8.24 (d, *J* = 1.7 Hz, 1H), 8.10 (dd, *J* = 8.0, 1.7 Hz, 2H), 8.02 (s, 1H), 7.99 (s, 2H), 7.97 (s, 1H), 7.89 (d, *J* = 7.8 Hz, 1H), 7.83 (t, *J* = 7.7 Hz, 1H), 7.53 (d, *J* = 8.0 Hz, 1H), 7.30 (d, *J* = 7.5 Hz, 1H), 7.16-7.11 (m, 2H), 7.01 (t, *J* = 7.5 Hz, 1H), 5.05 (t, *J* = 5.3 Hz, 1H), 4.49 (d, *J* = 5.1 Hz, 2H), 4.19 (dd, *J* = 11.0, 4.4 Hz, 2H), 3.88 (s, 3H), 3.54 (t, *J* = 10.7 Hz, 2H), 3.38 (s, 1H), 2.58 (s, 3H) ppm. ^13^C NMR (126 MHz, DMSO-d_6_, 300K): *δ* = 160.9, 159.5, 159.0, 158.0, 155.1, 153.2, 150.5, 140.0, 137.9, 136.6, 135.6, 133.6, 133.0, 132.3, 127.5, 127.0, 126.0, 124.9, 124.5, 122.8, 120.2, 117.8, 111.3, 105.8, 97.3, 66.3, 55.8, 42.5, 42.4, 24.2 ppm. HRMS (FTMS +p MALDI): m/z calculated for C_31_H_30_ClN_6_O_4_ [M+H]^+^: 585.2012, found: 585.2019. HPLC: t_R_ = 11.49 min.; purity ≥ 95% (UV: 254/280 nm). LCMS-ESI+ (m/z) [M+H]^+^; calculated: 585.2, found: 585.0.

#### 8-(((2r,5r)-5-Amino-1,3-dioxan-2-yl)methyl)-6-(2-chloro-4-(pyridin-2-yl)phenyl)-2-(methylamino)pyrido[2,3-*d*]pyrimidin-7(8H)-one (**8**)

Compound **8** was prepared according to general procedures II-IV and was obtained as a white solid with a yield of 15% (31 mg) over 3 steps. ^1^H NMR (500 MHz, DMSO-d_6_, 300K): *δ* = 8.73-8.66 (m, 2H), 8.24 (d, *J* = 1.7 Hz, 1H), 8.13-8.09 (m, 2H), 8.06-8.05 (m, 3H), 7.99 (td, *J* = 7.8, 1.7 Hz, 1H), 7.88 (s, 1H), 7.53 (d, *J* = 8.0 Hz, 1H), 7.48-7.46 (m, 1H), 5.13-5.12 (m, 1H), 4.55-4.49 (m, 2H), 4.22-4.19 (m, 2H), 3.59-3.55 (m, 2H), 3.39 (s, 1H), 2.92 (d, *J* = 2.2 Hz, 3H) ppm. ^13^C NMR (126 MHz, DMSO-d_6_, 300K): *δ* = 163.1, 161.6, 161.0, 159.3, 155.4, 153.8, 149.3, 139.4, 138.2, 137.0, 136.3, 133.8, 132.4, 127.1, 125.0, 124.2, 123.6, 121.0, 97.4, 66.3, 42.4, 42.1, 27.9 ppm. HRMS (FTMS +p MALDI): m/z calculated for C_24_H_24_ClN_6_O_3_ [M+H]^+^: 479.1593, found: 479.1613. HPLC: t_R_ = 10.74 min.; purity ≥ 95% (UV: 254/280 nm). LCMS-ESI+ (m/z) [M+H]^+^; calculated: 479.2, found: 479.0.

#### 8-(((2r,5r)-5-Amino-1,3-dioxan-2-yl)methyl)-6-(2-chloro-4-(3-fluoro-6-methylpyridin-2-yl)-phenyl)-2-(methylamino)pyrido[2,3-*d*]pyrimidin-7(8H)-one (**9**)

Compound **9** was prepared according to general procedures II-IV and was obtained as a yellow solid with a yield of 11% (21 mg) over 3 steps. ^1^H NMR (500 MHz, DMSO-d_6_, 300K): *δ* = 8.74-8.66 (m, 1H), 8.07-8.03 (m, 4H), 7.98-7.97 (m, 1H), 7.93 (d, *J* = 8.1 Hz, 1H), 7.88 (s, 1H), 7.77 (dd, *J* = 11.3, 8.5 Hz, 1H), 7.54 (d, *J* = 8.0 Hz, 1H), 7.38 (dd, *J* = 8.4, 3.3 Hz, 1H), 5.14-5.06 (m, 1H), 4.56-4.49 (m, 2H), 4.22-4.20 (m, 2H), 3.59-3.55 (m, 2H), 3.40 (s, 1H), 2.92 (d, *J* = 4.5 Hz, 3H), 2.53 (s, 3H) ppm. ^13^C NMR (126 MHz, DMSO-d_6_, 300K): *δ* = 161.8, 160.9, 159.5, 156.8, 155.3, 154.8, 154.3, 154.2, 141.5, 141.4, 137.0, 136.2, 136.1, 133.3, 132.1, 128.8, 128.7, 126.9, 125.4, 125.2, 124.7, 124.6, 124.1, 103.9, 97.4, 66.2, 42.3, 42.1, 27.9, 23.5 ppm. HRMS (FTMS +p MALDI): m/z calculated for C_25_H_25_ClFN_6_O_3_ [M+H]^+^: 511.1655, found: 511.1647. HPLC: t_R_ = 11.62 min.; purity ≥ 95% (UV: 254/280 nm). LCMS-ESI+ (m/z) [M+H]^+^; calculated: 511.2, found: 511.1.

#### 8-(((2r,5r)-5-Amino-1,3-dioxan-2-yl)methyl)-6-(2-chloro-4-(3-fluoropyridin-2-yl)phenyl)-2-(methylamino)pyrido[2,3-*d*]pyrimidin-7(8H)-one (**10**)

Compound **10** was prepared according to general procedures II-IV and obtained as a yellow solid with a yield of 23% (35 mg) over 3 steps. ^1^H NMR (500 MHz, DMSO-d_6_, 300K): *δ* = 8.73-8.66 (m, 1H), 8.60-8.59 (m, 1H), 8.09 (s, 3H), 8.05 (s, 1H), 7.98-7.97 (m, 1H), 7.95-7.94 (m, 1H), 7.92-7.88 (m, 2H), 7.57-7.54 (m, 2H), 5.14-5.06 (m, 1H), 4.56-4.49 (m, 2H), 4.22-4.20 (m, 2H), 3.60-3.56 (m, 2H), 3.39 (s, 1H), 2.91 (d, *J* = 4.5 Hz, 3H) ppm. ^13^C NMR (126 MHz, DMSO-d_6_, 300K): *δ* = 170.0, 161.8, 160.9, 159.5, 158.3, 156.3, 155.3, 146.0, 142.9, 137.0, 136.4, 136.0, 133.4, 132.2, 128.9, 127.0, 125.3, 125.1, 125.0, 124.0, 103.9, 97.4, 66.3, 42.3, 42.1, 27.9 ppm. HRMS (FTMS +p MALDI): m/z calculated for C_24_H_23_ClFN_6_O_3_ [M+H]^+^: 497.1499, found: 497.1491. HPLC: t_R_ = 11.71 min.; purity ≥ 95% (UV: 254/280 nm). LCMS-ESI+ (m/z) [M+H]^+^; calculated: 497.1, found: 497.0.

#### 8-(((2r,5r)-5-Amino-1,3-dioxan-2-yl)methyl)-6-(2-chloro-4-(3-chloropyridin-2-yl)phenyl)-2-(methylamino)pyrido[2,3-*d*]pyrimidin-7(8H)-one (**11**)

Compound **11** was prepared according to general procedures II-IV and obtained as a yellow solid with a yield of 15% (30 mg) over 3 steps. ^1^H NMR (500 MHz, DMSO-d_6_, 300K): *δ* = 8.73-8.65 (m, 2H), 8.11-8.08 (m, 4H), 7.97 (s, 1H), 7.90 (s, 1H), 7.82 (d, *J* = 1.7 Hz, 1H), 7.73 (dd, *J* = 7.9, 1.7 Hz, 1H), 7.53-7.49 (m, 2H), 5.14-5.06 (m, 1H), 4.56-4.50 (m, 2H), 4.22-4.20 (m, 2H), 3.60-3.56 (m, 2H), 3.39 (s, 1H), 2.92 (d, *J* = 4.5 Hz, 3H) ppm. ^13^C NMR (126 MHz, DMSO-d_6_, 300K): *δ* = 166.5, 162.3, 159.9, 155.8, 154.1, 148.6, 139.1, 137.5, 133.8, 133.3, 132.1, 131.1, 130.3, 129.8, 129.3, 128.4, 125.1, 104.4, 97.8, 66.7, 42.8, 42.5, 28.4 ppm. HRMS (FTMS +p MALDI): m/z calculated for C_24_H_23_Cl_2_N_6_O_3_ [M+H]^+^: 513.1203, found: 513.1202. HPLC: t_R_ = 11.73 min.; purity ≥ 95% (UV: 254/280 nm). LCMS-ESI + (m/z) [M+H]^+^; calculated: 513.1, found: 513.1.

#### 8-(((2r,5r)-5-Amino-1,3-dioxan-2-yl)methyl)-6-(2-chloro-4-(3-methylpyridin-2-yl)phenyl)-2-(methylamino)pyrido[2,3-*d*]pyrimidin-7(8H)-one (**12**)

Compound **12** was prepared according to general procedures II-IV and obtained as a white solid with a yield of 27% (42 mg) over 3 steps. ^1^H NMR (500 MHz, DMSO-d_6_, 300K): *δ* = 8.77-8.69 (m, 2H), 8.36 (d, *J* = 7.8 Hz, 1H), 8.17 (s, 3H), 8.07 (s, 1H), 7.91 (s, 1H), 7.87 (d, *J* = 1.7 Hz, 1H), 7.85 (dd, *J* = 7.9, 5.5 Hz, 1H), 7.69 (dd, *J* = 7.9, 1.7 Hz, 1H), 7.62 (d, *J* = 7.9 Hz, 1H), 5.13-5.07 (m, 1H), 4.55-4.50 (m, 2H), 4.23-4.20 (m, 2H), 3.61-3.57 (m, 2H), 3.39 (s, 1H), 2.93 (s, 3H), 2.45 (s, 3H) ppm. ^13^C NMR (126 MHz, DMSO-d_6_, 300K): *δ* = 161.4, 161.0, 159.0, 155.6, 151.5, 145.3, 142.1, 137.3, 137.2, 135.0, 134.9, 133.5, 132.2, 130.2, 128.1, 125.2, 124.1, 104.0, 97.5, 66.4, 42.5, 42.3, 28.0, 18.9 ppm. HRMS (FTMS +p MALDI): m/z calculated for C_25_H_26_ClN_6_O_3_ [M+H]^+^: 493.1749, found: 493.1752. HPLC: t_R_ = 10.60 min.; purity ≥ 95% (UV: 254/280 nm). LCMS-ESI+ (m/z) [M+H]^+^; calculated: 493.2, found: 493.2.

#### 8-(((2r,5r)-5-Amino-1,3-dioxan-2-yl)methyl)-6-(2-chloro-4-(3-methoxypyridin-2-yl)phenyl)-2-(methylamino)pyrido[2,3-*d*]pyrimidin-7(8H)-one (**13**)

Compound **13** was prepared according to general procedures II-IV and obtained as a yellow solid with a yield of 33% (41 mg) over 3 steps. ^1^H NMR (500 MHz, DMSO-d_6_, 300K): *δ* = 8.73-8.66 (m, 1H), 8.31 (d, *J* = 4.4 Hz, 1H), 8.05 (s, 3H), 8.01 (s, 1H), 7.97-7.96 (m, 1H), 7.92 (d, *J* = 8.0 Hz, 1H), 7.87 (s, 1H), 7.66 (d, *J* = 8.4 Hz, 1H), 7.48-7.44 (m, 2H), 5.13-5.06 (m, 1H), 4.56-4.49 (m, 2H), 4.22-4.21 (m, 2H), 3.91 (s, 3H), 3.59-3.55 (m, 2H), 3.39 (s, 1H), 2.92 (d, *J* = 3.8 Hz, 3H) ppm. ^13^C NMR (126 MHz, DMSO-d_6_, 300K): *δ* = 161.7, 161.0, 159.6, 159.4, 155.3, 153.7, 144.2, 140.9, 138.4, 136.8, 135.3, 132.7, 131.4, 129.5, 127.6, 124.4, 120.2, 103.9, 97.4, 66.3, 55.8, 42.3, 42.1, 27.9 ppm. HRMS (FTMS +p MALDI): m/z calculated for C_25_H_26_ClN_6_O_4_ [M+H]^+^: 509.1699, found: 509.1712. HPLC: t_R_ = 10.68 min.; purity ≥ 95% (UV: 254/280 nm). LCMS-ESI+ (m/z) [M+H]^+^; calculated: 509.2, found: 509.0.

#### 8-(((2r,5r)-5-Amino-1,3-dioxan-2-yl)methyl)-6-(2-chloro-4-(3-methoxy-6-methylpyridin-2-yl)phenyl)-2-(methylamino)pyrido[2,3-*d*]pyrimidin-7(8H)-one (**14**)

Compound **14** was prepared according to general procedures II-IV and obtained as a yellow solid with a yield of 12% (17 mg) over 3 steps. ^1^H NMR (500 MHz, DMSO-d_6_, 300K): *δ* = 8.73-8.66 (m, 1H), 8.08-8.04 (m, 3H), 7.99 (d, *J* = 1.7 Hz, 1H), 7.97-7.96 (m, 1H), 7.91 (dd, *J* = 8.0, 1.7 Hz, 1H), 7.86 (s, 1H), 7.59 (d, *J* = 8.5 Hz, 1H), 7.46 (d, *J* = 8.0 Hz, 1H), 7.32 (d, *J* = 8.5 Hz, 1H), 5.14-5.06 (m, 1H), 4.56-4.49 (m, 2H), 4.22-4.20 (m, 2H), 3.88 (s, 3H), 3.59-3.55 (m, 2H), 3.40 (s, 1H), 2.92 (d, *J* = 4.5 Hz, 3H), 2.49 (s, 3H) ppm. ^13^C NMR (126 MHz, DMSO-d_6_, 300K): *δ* 161.8, 161.0, 159.5, 155.3, 151.8, 148.8, 142.8, 138.2, 136.8, 135.3, 132.6, 131.4, 129.4, 127.6, 124.4, 123.8, 121.3, 103.9, 97.4, 66.2, 56.0, 42.3, 42.1, 27.9, 22.8 ppm. HRMS (FTMS +p MALDI): m/z calculated for C_26_H_28_ClN_6_O_4_: 523.1855 [M+H]^+^, found: 523.1846 [M+H]^+^. HPLC: t_R_ = 10.66 min.; purity ≥ 95% (UV: 254/280 nm). LCMS-ESI+ (m/z) [M+H]^+^; calculated: 523.2, found: 523.0.

#### 8-(((2r,5r)-5-Amino-1,3-dioxan-2-yl)methyl)-6-(2-chloro-4-(6-(trifluoromethyl)pyridin-2-yl)phenyl)-2-(methylamino)pyrido[2,3-*d*]pyrimidin-7(8H)-one (**15**)

Compound **15** was prepared according to general procedures II-IV and obtained as a yellow solid with a yield of 34% (40 mg) over 3 steps. ^1^H NMR (500 MHz, DMSO-d_6_, 300K): *δ* = 8.73-8.66 (m, 1H), 8.41 (d, *J* = 8.0 Hz, 1H), 8.26 (s, 1H), 8.23 (t, *J* = 7.9 Hz, 1H), 8.16-8.12 (m, 4H), 7.98 (s, 1H), 7.92 (d, *J* = 7.7 Hz, 1H), 7.87 (s, 1H), 7.58 (d, *J* = 8.0 Hz, 1H), 5.13-5.06 (m, 1H), 4.54-4.49 (m, 2H), 4.22-4.20 (m, 2H), 3.60-3.56 (m, 2H), 3.39 (s, 1H), 2.91 (d, *J* = 2.8 Hz, 3H) ppm. ^13^C NMR (126 MHz, DMSO-d_6_, 300K): *δ* = 161.8, 160.0, 155.5, 155.4, 154.9, 147.0, 146.7, 146.4, 146.2, 140.0, 138.2, 137.1, 137.0, 134.0, 132.7, 127.3, 125.3, 124.8, 124.2, 124.0, 122.7, 120.5, 119.9, 118.3, 103.9, 97.4, 66.3, 42.3, 42.1, 27.9 ppm. HRMS (FTMS +p MALDI): m/z calculated for C_25_H_23_ClF_3_N_6_O_3_: 547.1467 [M+H]^+^, found: 547.1457 [M+H]^+^. HPLC: t_R_ = 12.35 min.; purity ≥ 95% (UV: 254/280 nm). LCMS-ESI+ (m/z) [M+H]^+^; calculated: 547.1, found: 547.0.

#### 8-(((2r,5r)-5-Amino-1,3-dioxan-2-yl)methyl)-6-(2-chloro-4-(6-(difluoromethyl)pyridin-2-yl)phenyl)-2-(methylamino)pyrido[2,3-*d*]pyrimidin-7(8H)-one (**16**)

Compound **16** was prepared according to general procedures II-IV and obtained as a yellow solid with a yield of 30% (30 mg) over 3 steps. ^1^H NMR (500 MHz, DMSO-d_6_, 300K): *δ* = 8.73-8.66 (m, 1H), 8.28-8.27 (m, 2H), 8.16-8.10 (m, 5H), 7.98 (s, 1H), 7.88 (s, 1H), 7.72 (d, *J* = 7.7 Hz, 1H), 7.56 (d, *J* = 8.0 Hz, 1H), 7.06 (t, *J* = 55.0 Hz, 1H), 5.13-5.06 (m, 1H), 4.55-4.49 (m, 2H), 4.22-4.21 (m, 2H), 3.60-3.56 (m, 2H), 3.40 (s, 1H), 2.91 (d, *J* = 4.1 Hz, 3H) ppm. ^13^C NMR (126 MHz, DMSO-d_6_, 300K): *δ* = 161.8, 160.9, 159.5, 155.3, 154.3, 152.3, 152.1, 151.9, 139.3, 138.7, 137.0, 136.7, 133.9, 132.5, 127.1, 125.1, 124.0, 122.6, 119.7, 115.7, 113.8, 111.9, 103.9, 97.4, 66.3, 42.3, 42.1, 27.9 ppm. HRMS (FTMS +p MALDI): m/z calculated for C_25_H_24_ClF_2_N_6_O_3_: 529.1561 [M+H]^+^, found: 529.1553 [M+H]^+^. HPLC: t_R_ = 12.06 min.; purity ≥ 95% (UV: 254/280 nm). LCMS-ESI+ (m/z) [M+H]^+^; calculated: 529.2, found: 529.0.

#### 8-(((2r,5r)-5-Amino-1,3-dioxan-2-yl)methyl)-6-(2-chloro-4-(6-methoxypyridin-2-yl)phenyl)-2-(methylamino)pyrido[2,3-*d*]pyrimidin-7(8H)-one (**17**)

Compound **17** was prepared according to general procedures II-IV and obtained as a yellow solid with a yield of 28% (43 mg) over 3 steps. ^1^H NMR (500 MHz, DMSO-d_6_, 300K): *δ* = 8.72-8.67 (m, 1H), 8.21 (d, *J* = 1.7 Hz, 1H), 8.11-8.08 (m, 4H), 8.05 (s, 1H), 7.85 (s, 1H), 7.82-7.79 (m, 1H), 7.65 (d, *J* = 7.4 Hz, 1H), 7.50 (d, *J* = 8.0 Hz, 1H), 6.83 (d, *J* = 8.2 Hz, 1H), 5.13-5.07 (m, 1H), 4.54-4.50 (m, 2H), 4.23-4.20 (m, 2H), 3.97 (s, 3H), 3.60-3.56 (m, 2H), 3.40 (s, 1H), 2.92 (s, 3H) ppm. ^13^C NMR (126 MHz, DMSO-d_6_, 300K): *δ* = 163.4, 162.4, 161.1, 158.6, 155.6, 151.9, 140.3, 139.7, 136.8, 136.0, 133.8, 132.3, 126.9, 124.9, 124.7, 113.4, 110.3, 104.0, 97.4, 66.4, 53.0, 42.4, 42.2, 28.0 ppm. HRMS (FTMS +p MALDI): m/z calculated for C_25_H_26_ClN_6_O_4_: 509.1699 [M+H]^+^, found: 509.1698 [M+H]^+^. HPLC: t_R_ = 12.27 min.; purity ≥ 95% (UV: 254/280 nm). LCMS-ESI+ (m/z) [M+H]^+^; calculated: 509.2, found: 509.2.

#### 8-(((2r,5r)-5-Amino-1,3-dioxan-2-yl)methyl)-6-(2-chloro-4-(4,6-(dimethyl)pyridin-2-yl)phenyl)-2-(methylamino)pyrido[2,3-*d*]pyrimidin-7(8H)-one (**18**)

Compound **18** was prepared according to general procedures II-IV and obtained as a white solid with a yield of 22% (34 mg) over 3 steps. ^1^H NMR (500 MHz, DMSO-d_6_, 300K): *δ* = 8.74-8.67 (m, 1H), 8.20 (d, *J* = 1.8 Hz, 1H), 8.07-8.03 (m, 4H), 7.99 (s, 1H), 7.90 (s, 1H), 7.89 (s, 1H), 7.58 (d, *J* = 8.0 Hz, 1H), 7.39 (s, 1H), 5.14-5.06 (m, 1H), 4.55-4.49 (m, 2H), 4.22-4.19 (m, 2H), 3.60-3.55 (m, 2H), 3.39 (s, 1H), 2.92 (d, *J* = 4.1 Hz, 3H), 2.62 (s, 3H), 2.47 (s, 3H) ppm. ^13^C NMR (126 MHz, DMSO-d_6_, 300K): *δ* = 161.7, 161.0, 160.9, 159.7, 159.5, 156.1, 155.3, 151.5, 137.0, 136.9, 133.7, 132.4, 127.8, 125.7, 124.8, 123.9, 120.8, 103.9, 97.4, 66.3, 42.3, 42.1, 27.9, 22.2, 20.9 ppm. HRMS (FTMS +p MALDI): m/z calculated for C_26_H_28_ClFN_6_O_3_: 507.1906 [M+H]^+^, found: 507.1898 [M+H]^+^. HPLC: t_R_ = 10.68 min.; purity ≥ 95% (UV: 254/280 nm). LCMS-ESI+ (m/z) [M+H]^+^; calculated: 507.2, found: 507.1.

### Homology modeling and molecular docking

System preparation and docking calculations were performed using the Schrödinger Drug Discovery suite for molecular modeling (version 2019.4). Protein-ligand complexes were prepared with the protein preparation wizard^30^ to fix protonation states of amino acids, add hydrogens and also fix missing side-chain atoms. The missing activation loop between Ala174 and Thr194 of MST3 was modelled based on the MST3 (PDB 3A7J) coordinates and optimized using Prime^31^, and Thr178 was kept phosphorylated. The homology model of the human SIK2 kinase domain (UniProt: Q9H0K1, residues 20-271) was generated based on a stable frame from the MST3 MD simulation (described below) using Prime^32^. All ligands for docking were drawn using Maestro and prepared using LigPrep^33^ to generate the three-dimensional conformation, adjust the protonation state to physiological pH (7.4), and calculate the partial atomic charges, with the force field OPLS3e.^34^ Docking studies with the prepared ligands were performed using Glide (Glide V7.7)^35,36^ with the flexible modality of induced-fit docking with extra precision (XP), followed by a side-chain minimization step using Prime. Ligands were docked within a grid around 12 Å from the centroid of the co-crystallized ligand, generating ten poses per ligand.

### Molecular dynamics simulation

Selected docking poses were further validated by molecular dynamics simulation, where ligand stability within the proposed pocket and its interactions were evaluated. MD simulations were carried out using Desmond^37^ engine with the OPLS3e force-field,^34,38^ which leads to improved performance in predicting protein-ligand binding affinities. The protein-ligand systems were placed in a cubic box with 13 Å from the box edges to any atom of the protein, using PBC conditions, and filled with TIP3P^39^ water. Then, all systems were equilibrated by short simulations under the NPT ensemble for 5 ns implementing the Berendsen thermostat and barostat methods. A constant temperature of 310 K and 1 atm of pressure was used throughout the simulation using the Nose-Hoover thermostat algorithm and Martyna-Tobias-Klein Barostat algorithm, respectively. After minimization and relaxing steps, production steps of at least 1 µs were performed. All MD simulations were performed in at least three independent runs with randomly generated seeds. Trajectories and interaction data are available from the Zenodo repository (accession code: 10.5281/zenodo.4288340). The stability of the MD simulations was monitored by analyzing root mean square deviation (RMSD) and root mean square fluctuation (RMSF) of the ligand and protein along the trajectory. Variation in the RMSD values stabilized after 150 ns for all the ligands, which was related to the stabilization of the activation loop in the case of MST3, or the relaxation of the homology model, in the case of SIK2. For principal component analyses, the backbone of each frame was extracted and aligned using trj_selection_dl.py and trj_align.py scripts from Schrodinger. Individual simulations from all the runs were merged using trj_merge.py into a final trajectory and CMS file, which was further used for the generation of the principal components. Principal components of protein C-alpha atoms were calculated using the trj_essential_dynamics.py script. Most of the large movements were captured in the first two components, with the first component having 27% and the second component having 17% of the total motion protein-ligand interactions. Protein conformational changes were analyzed using the Simulation Interaction Diagram tool (SID).

### WaterMap calculation

WaterMap calculations^40^ were performed using Maestro, and the system was solvated in TIP3P water box extending at least 10 Å beyond the truncated protein in all directions. 5 ns molecular dynamics simulation was performed, following a standard relaxation protocol, and the water molecule trajectories were then clustered into distinct hydration sites. Entropy and enthalpy values for each hydration site were calculated using inhomogeneous solvation theory.

### Statistical analyses and figures

Structural images were generated using PyMol 2.4.3,^41^ and graphs were plotted using either GraphPad Prism version 8.4, *GraphPad Software* (San Diego, California USA, www.graphpad.com) or the Seaborn package implemented in Python 3.8.

### *In vitro* kinase assay

To measure kinase activity of SIK2, *in vitro* kinase assays were performed as described in Matthess et al.^42^ Samples were resolved by SDS-PAGE and subjected to autoradiography. Recombinant human active SIK2 was purchased from ProQinase Reaction Biology, product No. 1371-0000-1.

### Proliferation assays

To measure the cell proliferation using the Cell Titer-Blue^®^ Cell Viability Assay, 2,500 cells per well were seeded in 96-well plates and incubated with the different single or combination treatments. Cells were incubated with 20 µl substrate of the Cell Titer-Blue^®^ Cell Viability Assay for 4 h, and the light absorbance was measured at 540 nm (Victor X4, Perkin Elmer). 

### Caspase3/7 activity

The activity of Caspase 3/7 was determined using the Caspase-Glo^®^ 3/7 Assay (*Promega*). Upon incubation of the cells with the different treatment regimens, 20 µl substrate were added to cell lysates. After 30 minutes shaking at room temperature in the dark, luminescence was detected (Victor X4, Perkin Elmer).

### Spheroid (3D) cultures

A cell suspension of 1,500 cells/50 µl was prepared and pipetted from the topside into a 96 well Perfect 3D Hanging Drop plate (BioTrend). Plates were incubated at 37 ^°^C for several days until hanging drops had developed. The 3D culture was harvested on a 96 well plate covered with 1% agarose by low spin centrifugation. Treatment of cells with the different combinations was performed as indicated. Cells were stained with the LIVE/DEAD viability/cytotoxicity kit (Molecular Probes/Thermofisher) for 30 min and inspected using a fluorescence microscope AxioObserver.Z1 microscope with a HCX PL APO CS 10 × 1.4 UV objective (Zeiss, Göttingen). While the polyanionic dye calcein is retained in live cells, producing an intense uniform green fluorescence, EthD-1 enters cells with damaged membranes, thereby producing a bright red fluorescence upon binding to nucleic acids in dead cells. The ratios of viable/dead cells were calculated with the software ImageJ Fiji.

### NanoBRET™ assay

The assay was performed as described previously.^22,23^ In brief: full-length kinases were obtained as plasmids cloned in frame with a terminal NanoLuc-fusion (*Promega*) as specified in **Supplementary Table 6**. Plasmids were transfected into HEK293T cells using FuGENE HD (*Promega*, E2312), and proteins were allowed to express for 20 h. Serially diluted inhibitor and NanoBRET™ Kinase Tracer K10 (*Promega*) at a concentration determined previously as the Tracer K10 IC_50_ (**Supplementary Table 6**) were pipetted into white 384-well plates (Greiner 781207) using an Echo acoustic dispenser (*Labcyte*). The corresponding protein-transfected cells were added and reseeded at a density of 2 x 10^5^ cells/mL after trypsinization and resuspending in Opti-MEM without phenol red (*Life Technologies*). The system was allowed to equilibrate for 2 hours at 37 °C/5% CO_2_ prior to BRET measurements. To measure BRET, NanoBRET™ NanoGlo Substrate + Extracellular NanoLuc Inhibitor (*Promega*, N2540) was added as per the manufacturer’s protocol, and filtered luminescence was measured on a PHERAstar plate reader (BMG Labtech) equipped with a luminescence filter pair (450 nm BP filter (donor) and 610 nm LP filter (acceptor)). Competitive displacement data were then graphed using GraphPad Prism 8 software using a normalized 3-parameter curve fit with the following equation: Y=100/(1+10^(X-LogIC50)).

### Differential scanning fluorometry (DSF) assay

DSF assays were performed as previously described by Fedorov et al.^24^ Differences in melting temperature are given as ΔT_m_ values in °C. Kinome-wide selectivity profile. Compounds 6, 7 and 9 were tested against a panel of 468 kinases in the scanMAX^℠^ kinase assay performed by *Eurofins Scientific*. Compound 6 was tested at a concentration of 100 nM and 1 µM, while compounds 7 and 9 were tested at 1 µM. Compound 10 was tested against a panel of 443 kinases in the radiometric ^33^PanQinase^®^ Activity Assay at a concentration of 1 µM. The Assay was performed by *Reaction Biology*.

### Protein expression and purification

MST3 was expressed using a construct containing residues R4-D301 followed by a C-terminal hexahistidine tag in vector pNIC-CH. MST4 was expressed as a fusion protein with an N-terminal hexahistidine tag, followed by a TEV cleavage site and MST4 residues M1-S300 in vector pNIC28-Bsa4. The expression plasmids were transformed into BL21(D3)-R3-pRARE2 cells, and cells were grown in 2x LB media containing 50 µg ml^−1^ kanamycin and 34 µg ml^−1^ chloramphenicol and cultured overnight at 37 °C. 10-12 ml of the overnight culture were transferred into 1L of TB media containing 50 µg ml^−1^ kanamycin, and cultures were grown at 37 °C with shaking until an OD_600_ of 1.5-1.8 was reached. The temperature was reduced to 18 °C, and IPTG was added at a final concentration of 0.5 mM. The cells were grown overnight and harvested on the next day by centrifugation at 5,000 rpm for 10 minutes. The cell pellet was resuspended in lysis buffer containing 50 mM HEPES pH 7.5, 500 mM NaCl, 30 mM imidazole, 5% glycerol, and 0.5 mM TCEP. Protease inhibitor cocktail from Sigma was added at a dilution of 1:5000. The resuspended cells were lysed by sonication and cleared of DNA by addition of 0.15% polyethylenimine, followed by centrifugation at 23,000 rpm for 30 minutes. The supernatant was loaded onto a gravity column containing 5 ml of 50% Ni-NTA resin (Qiagen) pre-equilibrated with 40 ml lysis buffer. The column was washed with 100 ml lysis buffer, and MST3 and MST4 were then eluted with lysis buffer containing 50 mM, 100 mM, 200 mM, and 300 mM imidazole. The fractions containing MST3 were pooled and concentrated using a 10 kDa cut-off ultrafiltration device and further purified on a size exclusion chromatography (SEC; HiLoad 16/600 Superdex 200) pre-equilibrated with SEC buffer containing 25 mM HEPES pH 7.5, 200 mM NaCl, 5% glycerol and 0.5 mM TCEP. In the case of MST4, the eluted fractions from the Ni-NTA column containing MST4 were dialyzed in SEC buffer and submitted to cleavage of the tag by TEV protease overnight. The cleaved MST4 sample was then loaded onto a gravity column containing 2 ml of 50% Ni-NTA, and the flow through was collected, concentrated and purified by SEC using a Hiload 16/600 Superdex 200 column (same buffer as for MST3). Protein identity was confirmed by electrospray ionization mass spectrometry (MST3: expected 34,507.7 Da, observed 34,506.9 Da, MST4: expected 33,817.9 Da, observed 33,818.7 Da).

### Crystallization and structure determination

Inhibitors were added to the protein solution (10 mg/ml MST3 or 12 mg/ml to MST4) at a final concentration of 1 mM and incubated for at least 1 h at 4 °C. The samples were then centrifuged to remove any insoluble material, and crystals were obtained using the sitting drop vapor diffusion method. Crystallization buffers are given in **Supplementary Table 7**. The crystals were cryoprotected with crystallization buffer complemented with 25% ethylene glycol and flash frozen in liquid nitrogen. Diffraction data were collected at 100 K at the Swiss Light Source (beamline X06SA). The data were processed with XDS^43^ and scaled with AIMLESS^44^, and the structures were solved by molecular replacement with the program Phaser^45^ implemented in the CCP4 suite^46^ using the structure of MST3 (PDB ID 3ZHP) as a search model for MST3 and subsequently the new G-5555/MST3 complex as a search model for MST4.The structures were then refined using iterative cycles of model building in COOT^47^ and refinement with REFMAC^48^ or PHENIX^49^. Ligand dictionary files were obtained from the GRADE server (http://grade.globalphasing.org). Data collection and refinement statistics are summarized in **Supplementary Table 7**.

## Accession Codes

Coordinates and structure factors of the MST3/4-inhibitor complexes are available in the Protein Data Bank (PDB) under accession codes 7B30 (MST3-**6** complex), 7B31 (MST3-**10** complex), 7B32 (MST3-**7** complex), 7B33 (MST3-**16** complex), 7B34 (MST3-**9** complex), 7B35 (MST3-**14** complex) and 7B36 (MST4-**6** complex).

## ACKNOWLEDGMENT

R.T. is supported by the German Research Foundation (DFG) grant 397659447. M. Rak is grateful for the support by the Dr. Hilmer-Stiftung. We are also grateful for support by the SGC, a registered charity (no. 1097737) that receives funds from; AbbVie, Bayer AG, Boehringer Ingelheim, the Canada Foundation for Innovation, Eshelman Institute for Innovation, Genentech, Genome Canada through Ontario Genomics Institute [OGI-196], EU/EFPIA/OICR/McGill/KTH/Diamond, Innovative Medicines Initiative 2 Joint Undertaking [EUbOPEN grant 875510], Janssen, Merck KGaA (aka EMD in Canada and US), Merck & Co (aka MSD outside Canada and US), Pfizer, the São Paulo Research Foundation-FAPESP and Takeda as well as support from the German translational cancer network DKTK and the Frankfurt Cancer Institute (FCI). A.C.J. is supported by German Research Foundation (DFG) grant JO 1473/1-3. The data collection at SLS was supported by funding from the European Union’s Horizon 2020 research and innovation program grant agreement number 730872, project CALIPSOplus. We thank the staff at beamline X06SA of the Swiss Light Source for assistance during data collection. We thank the CSC-Finland for the generous computational resources provided. We thank Amelie Tjaden for help with the cell-based assays and Astrid Kaiser for performing the microsomal stability assays.

## ABBREVIATIONS

ABL: Abelson murine leukemia viral oncogene homolog 1
AKT: protein kinase B
AMPK: AMP-activated protein kinase
BTK: Bruton’s tyrosine kinase
CaMK: calcium calmodulin kinase
CREB: cAMP response element-binding protein
DLBCL: diffuse large B-cell lymphoma
DSF: differential scanning fluorometry
FGF: fibroblast growth factor
HDAC: histone deacetylase
JAK2: Janus kinase 2
KHS1: mitogen-activated protein kinase kinase kinase kinase 5
LCK: lymphocyte-specific protein tyrosine kinase
MST: mammalian STE20-like protein kinase
NUAK2: SNF1/AMP kinase-related kinase
PAK: p21-activated kinase
PI3K: phosphoinositide 3-kinase
SIK: salt-inducible kinase
SRPK1: serine/arginine protein kinase 1
STE: homologous kinases to the yeast proteins STE20, STE11, and STE7
TORC: transducer of regulated CREB1
VEGFR2: vascular endothelial growth factor receptor 2
YSK1: serine/threonine-protein kinase 25

