## Supplementary Information for "Structure-based design of selective salt-inducible kinase (SIK) inhibitors"

**Table of Content**


Supplementary Figures S1-S12 ........................................................................ S3-S14

Table S1: Selectivity profile of G-5555 (100 nM)................................................ S15-S33

Table S2: Selectivity profile of G-5555 (1 µM).................................................... S34-S50

Table S3: Selectivity profile of **7** (1 µM)................................................................ S51-S68

Table S4: Selectivity profile of **9** (1 µM)................................................................ S69-S86

Table S5: Selectivity profile of **10** (1 µM).......................................................... S87-S102

Table S6: NanoBRET™ assay information ..............................................................S103

Table S7: Data collection and refinement statistics................. ................................S104

Analytical data for compounds **7-38**................. ............................................. S105-S181

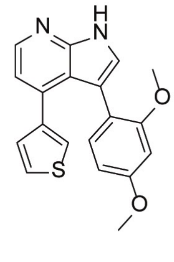

ARN-3236
(**5**)

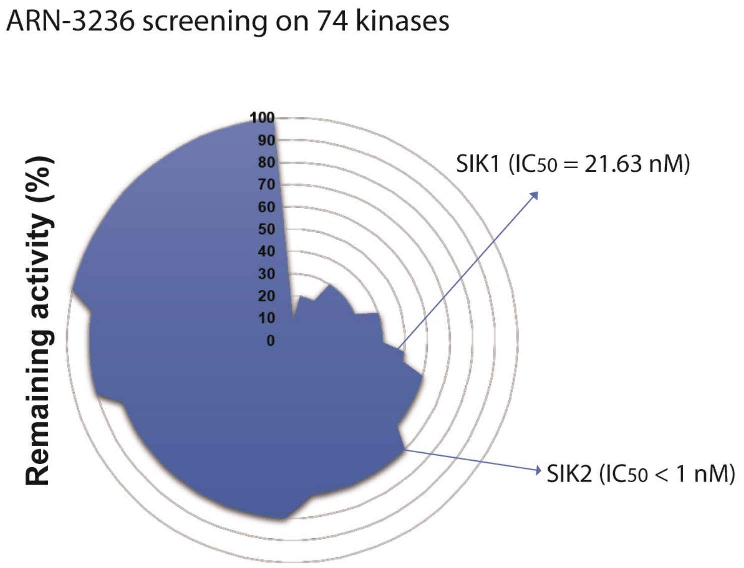


**Figure S1.** Selectivity profile of SIK2 inhibitor ARN-3236 (5) at a concentration of 500 nM against a panel of 74 kinases. Data report as percent of activity remaining in presence of inhibitor relative to activity measured in absence of inhibitor (100%). Graph created from the information on the supplementary material from Lombardi, M. S.; Gilliéron, C.; Dietrich, D.; Gabay, C. SIK inhibition in human myeloid cells modulates TLR and IL-1R signaling and induces an anti-inflammatory phenotype. *Journal of leukocyte biology* **2016**, *99* (5), 711–721.

A

0.1 μM

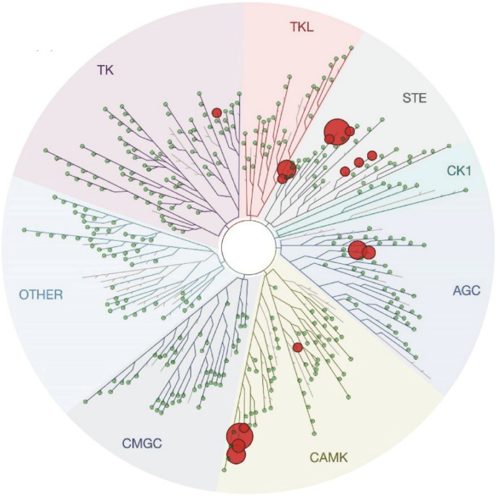


B

1 μM

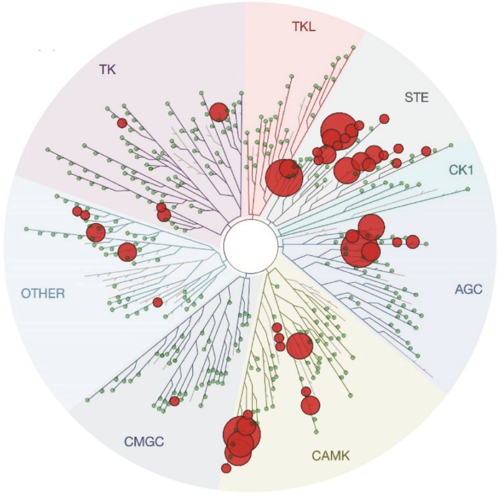


C

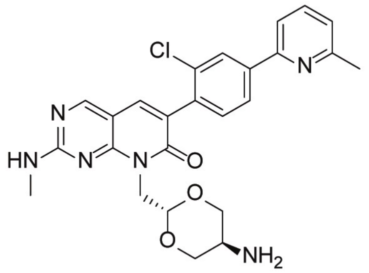

| | remaining activity (%) | |
|---|---|---|
| Targets | 0.1 μM | 1 μM |
| KIAA0999 (SIK3) | 0.2 | 0 |
| MST4 | 0.5 | 0 |
| SIK1 | 1.4 | 0.6 |
| STK32B | 3.5 | 3.7 |
| PAK1 | 4.4 | 4.5 |
| STK32A | 5.3 | 3.7 |
| SIK2 | 6.5 | 0.8 |
| CDPK1 | 9.1 | 0.4 |
| MAP2K5 | 13 | 0 |
| PAK2 | 13 | 0 |
| STK24 | 15 | 1.7 |
| MAP4K5 | 18 | 5.3 |
| NUAK2 | 19 | 2 |
| MAP2K1 | 24 | 1 |
| LTK | 28 | 4.1 |
| DCLK3 | 29 | 0.5 |
| MAP2K2 | 33 | 1.4 |
| STK36 | 36 | 2.9 |
| AURKC | 37 | 20 |
| STK10 | 37 | 2.1 |
| STK25 | 40 | 36 |
| CAMK1D | 43 | 2.5 |
| MARK3 | 45 | 18 |
| MAST1 | 47 | 0.4 |
| MYO3B | 48 | 57 |
| PIP5K1A | 48 | 67 |
| MAP2K4 | 49 | 9.6 |

**Figure S2.** Selectivity profile of G-5555. (A) Kinome tree of G-5555 screened at a concentration of 0.1 μM using scanMAX$^{SM}$ kinase assay panel of 468 kinases (*Eurofins Scientific*). (B) Kinome tree of G-5555 screened at a concentration of 1 μM using scanMAX$^{SM}$ kinase assay panel of 468 kinases (*Eurofins Scientific*). (C) Top off-targets at 0.1 μM and 1 μM.

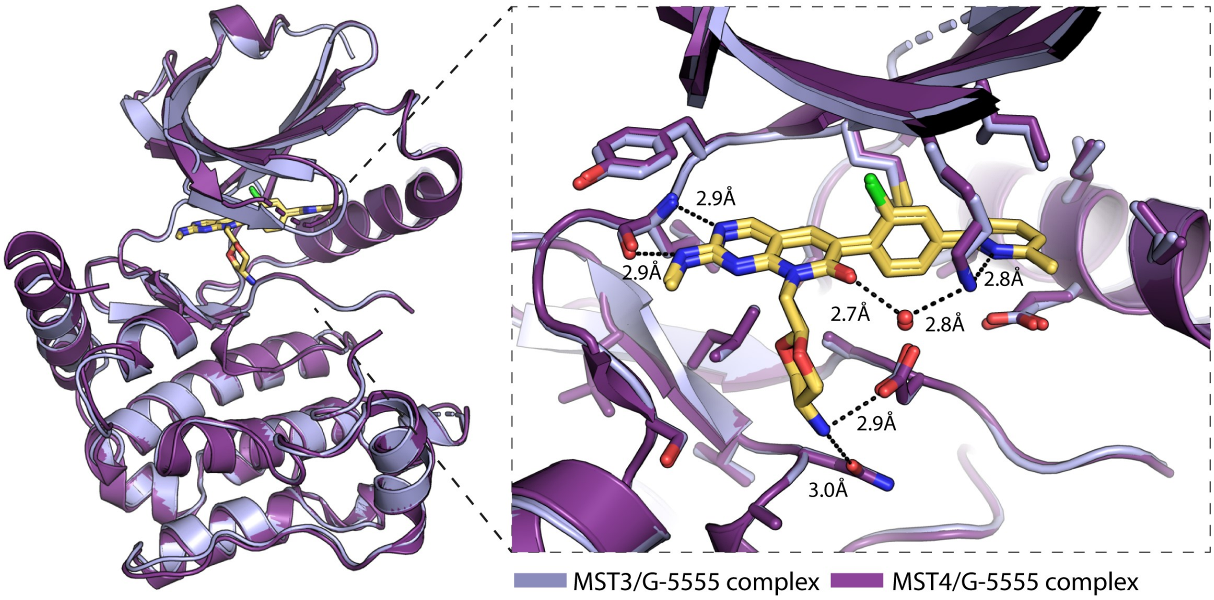


**Figure S3.** Comparison of the crystal structures of MST3 (light purple) and MST4 (violet) highlighting the conservation of amino acids in the binding site, interactions with G-5555 and conservation of a water molecule in the binding site.

A

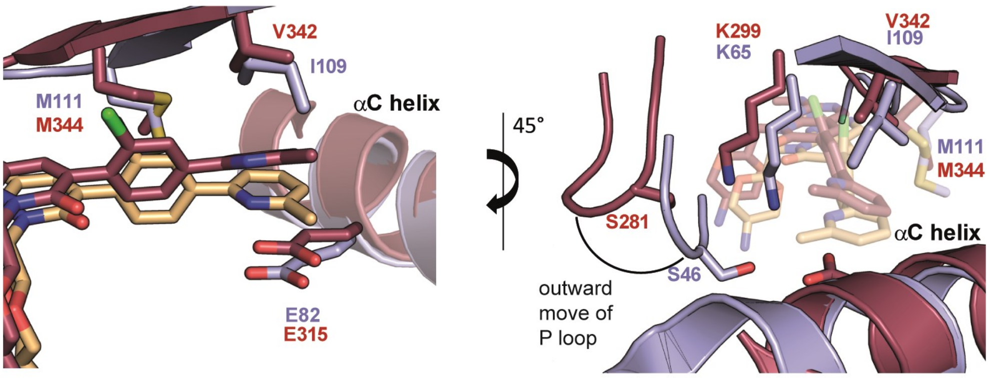


B

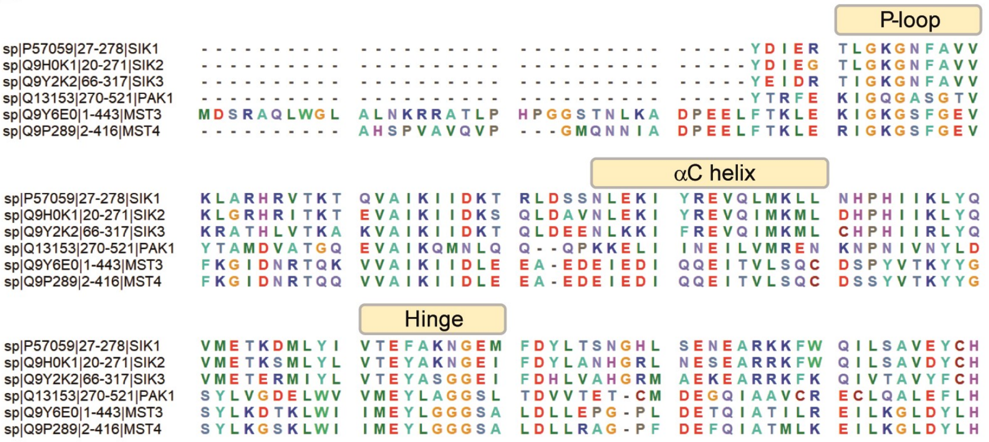


**Figure S4.** (A) Comparison of the crystal structures of MST3 (purple) and PAK1 (dark red) highlighting the main differences between the binding mode of G-5555 in the back-pocket region. (B) Sequence alignment of SIKs and main off-targets of G-5555, MSTs and PAK1 (BioEdit 7.2).

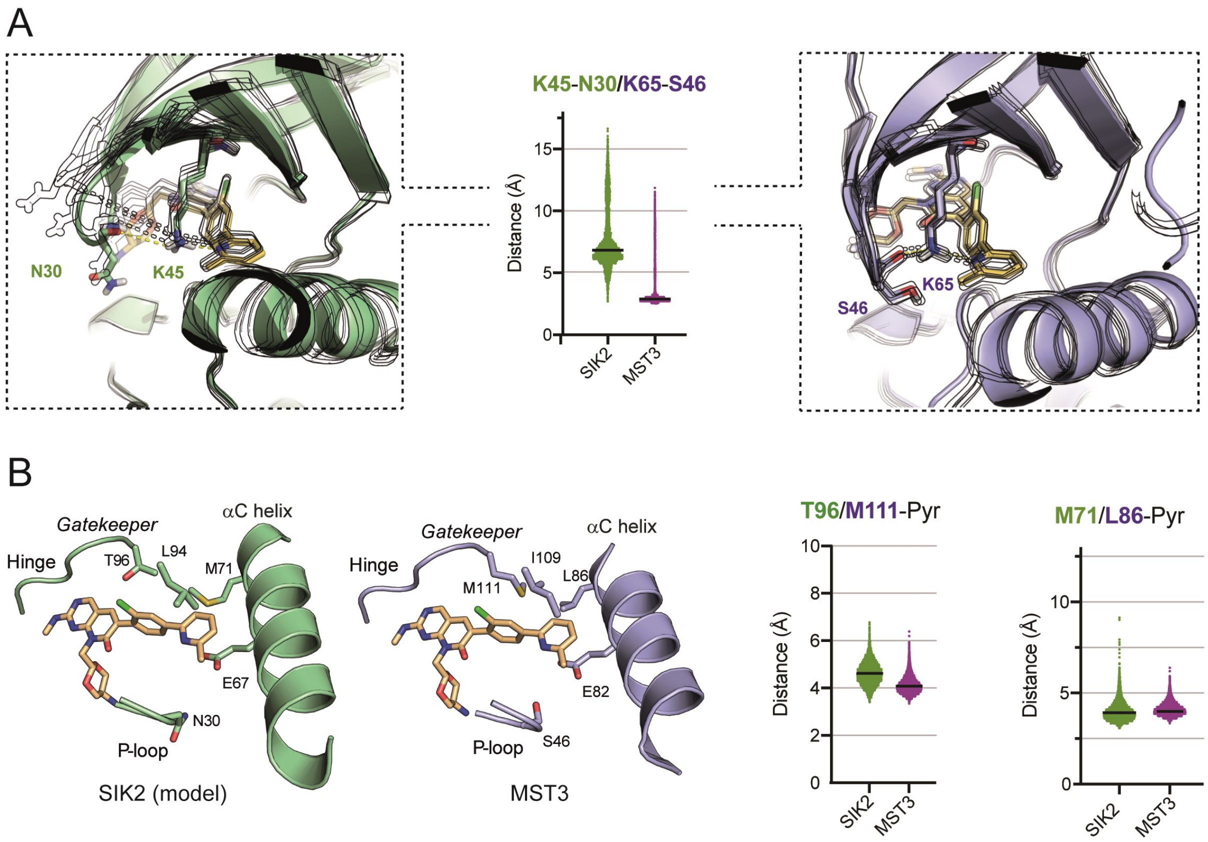


**Figure S5.** Binding pocket fluctuation in MD simulations. (A) Comparison of the movement of the P-loop region relative to the catalytic lysine in SIK2 and MST3. The distance from Ser46 in MST3 and Asn30 in SIK2 to the catalytic lysine was calculated for snapshots of the MD trajectory to assess the flexibility of this region. The oxygen from the carbonyl group of the main chain of Ser36 and Asn30 and the nitrogen from the catalytic lysine were chosen for the distance calculation. (B) Distribution of the distance between back-pocket residues of SIK2 and MST3 and the pyridine moiety of G-5555 along the MD trajectory. The first atom of the side chain of the back pockets was chosen for distance calculation with the center of mass of the pyridine ring.

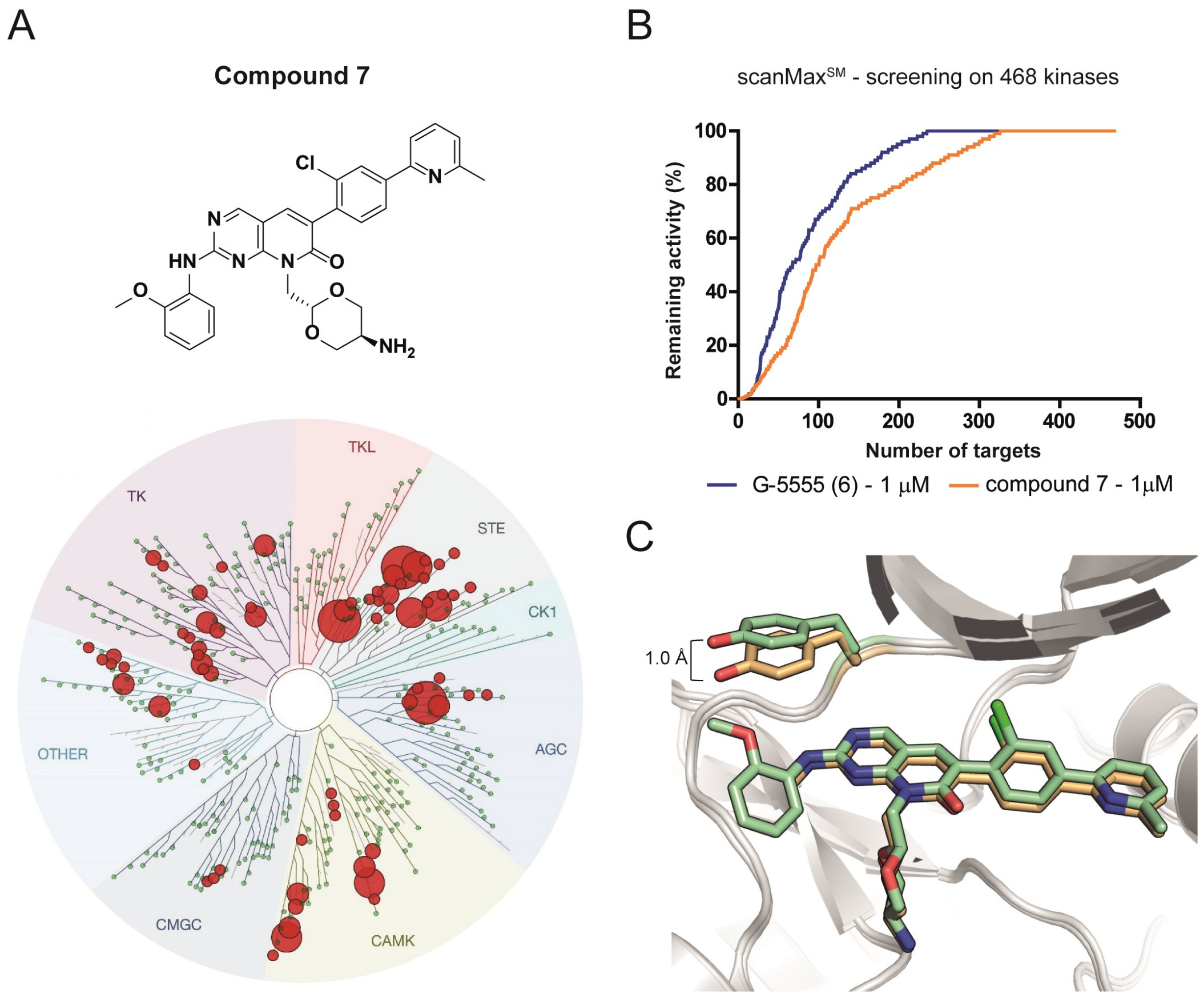


**Figure S6.** Comparison of the selectivity profile of G-5555 and compound **7**. (A) Kinome tree of compound **7** screened at 1 µM using scanMAX$^{SM}$ kinase assay panel of 468 kinases (*Eurofins Scientific*). (B) Comparison of selectivity profile of G-5555 and compound **7**. (C) Comparison of the binding mode of G-5555 in yellow (PDB code 7B30) and compound **7** in green (PDB code 7B32).

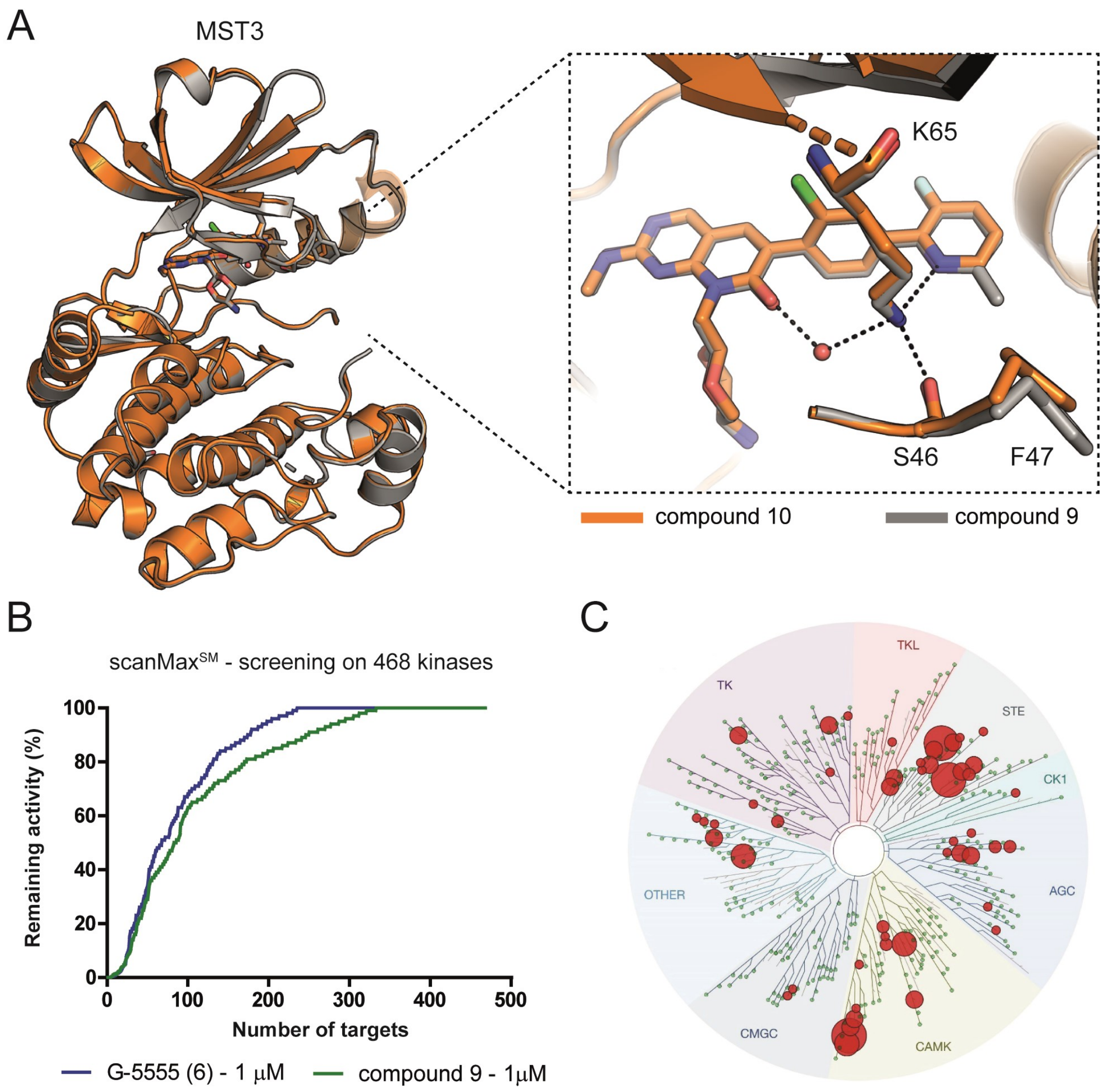


**Figure S7.** Binding mode of compounds **9** and **10** (MRIA9) in MST3. (A) Comparison of the binding mode of compound **9** (gray, PDB code 7B34) and compound **10** (MRIA9) (orange, PDB code 7B31). (B) Selectivity profile of G-5555 and compound **9** using scanMAX$^{SM}$ kinase assay panel of 468 kinases (*Eurofins Scientific*). (C) Kinome tree of compound **9** screened at 1 μM in scanMAX$^{SM}$ kinase assay (*Eurofins Scientific*).

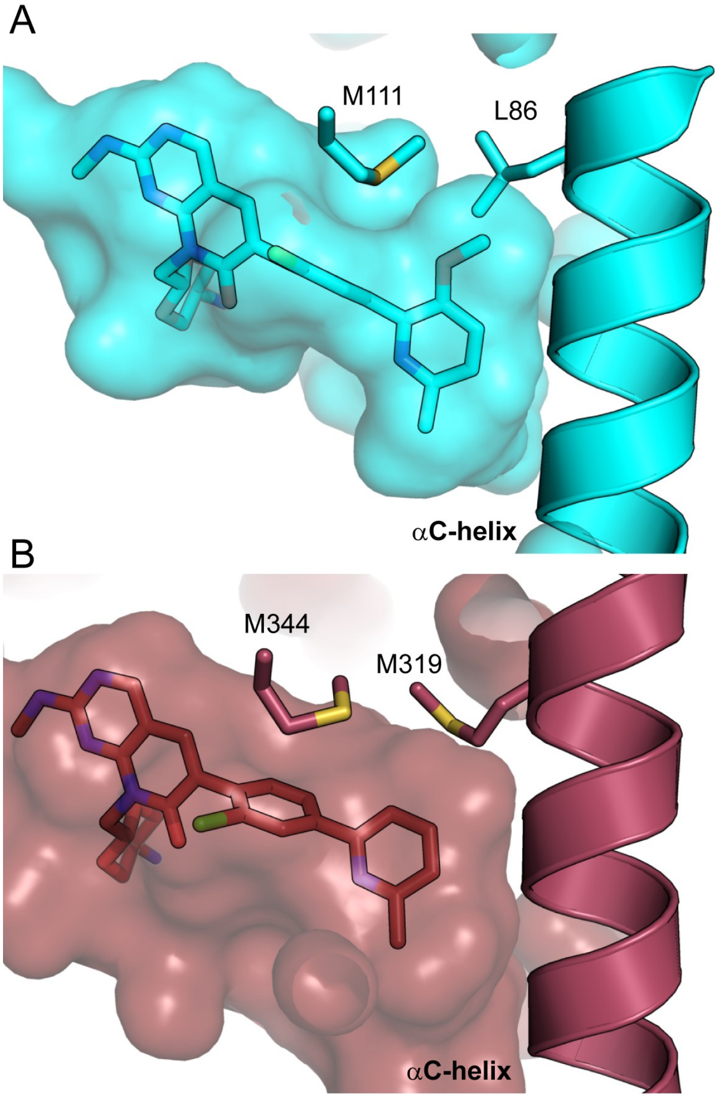


**Figure S8.** Comparison of the MST3 and PAK1 back pocket. (A) Binding mode of compound **14** in MST3 (cyan) and accommodation of the methoxy group in the back pocket (PDB code 7B35). (B) Binding mode of compound G-5555 in PAK1 (dark red), highlighting the back-pocket region (PDB code 5DEY).

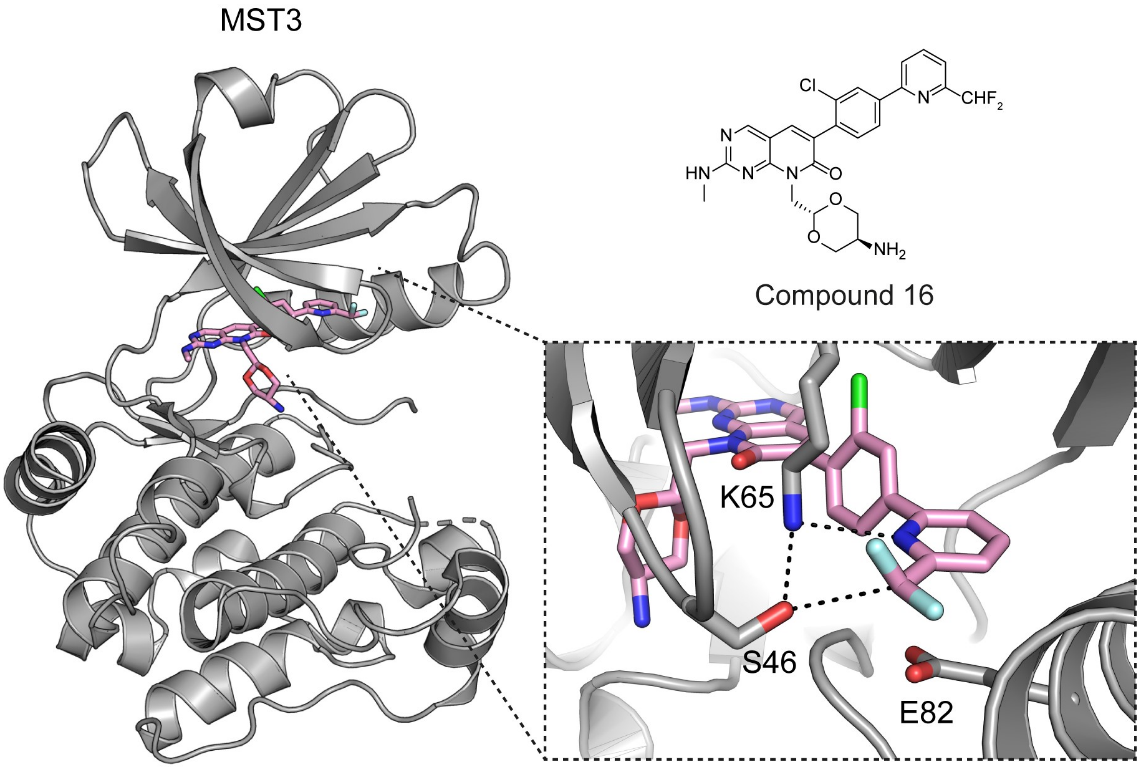


**Figure S9.** Binding mode of compound **16** with MST3 (PDB 7B33) and main interactions in the back pocket. The $CHF_2$ moiety contributes to binding to MST3 through direct interaction with Ser46.

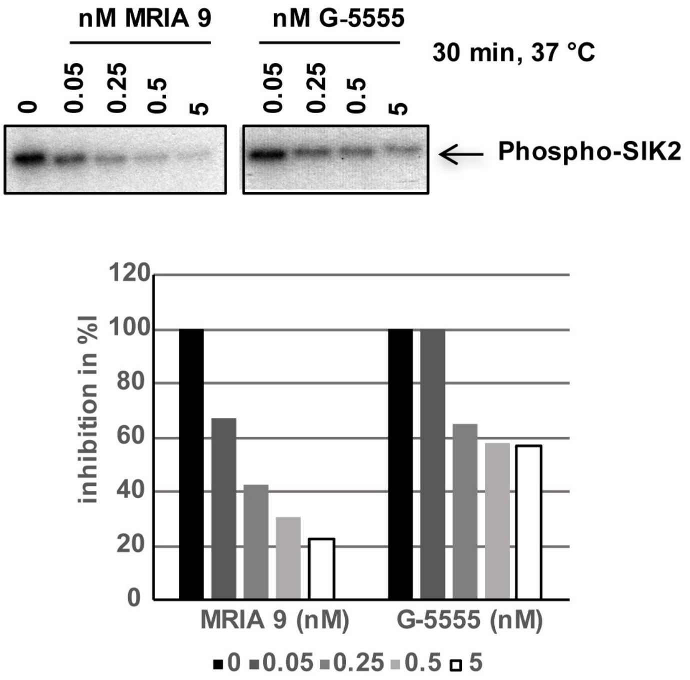


**Figure S10.** MRIA9 potently inhibits SIK2 *in vitro*. Representative *in vitro* kinase assay in which the autophosphorylation of SIK2 has been assessed for increasing concentrations of MRIA9 and G-5555. Band intensities were quantified and are represented as a bar graph.

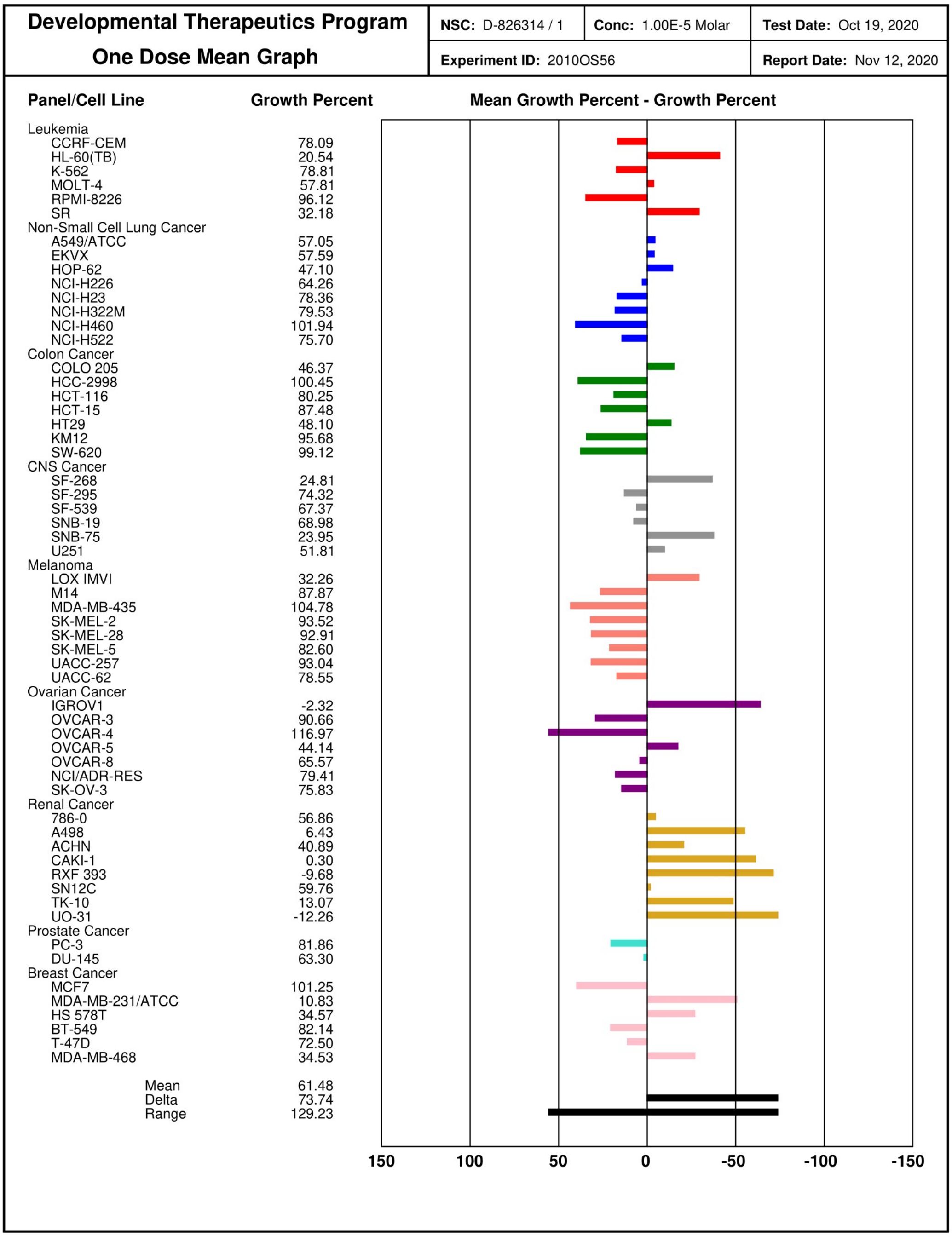


**Figure S11.** Results of compound **10** (MRIA9) tested against 60 cancer cell lines in the NCI-60 Human Tumor Cell Lines Screen at a single-dose concentration of 1 μM. Growth inhibition (values between 0 and 100) and lethality (values between 0 and -100) are shown as a bar chart.

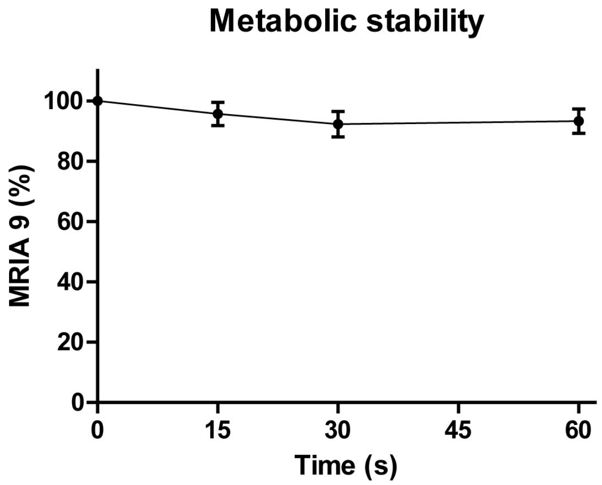


| Descriptor | Interpretation | Limit Values | G-5555[+] | G-5555 | MRIA9[+] | MRIA9 |
|---|---|---|---|---|---|---|
| PCaco | Predicted apparent Caco-2 cell permeability in nm/sec[a]. Caco-2 cells are a model for the gut-blood barrier | <25 poor, >500 great | 110 | 140.1 | 93.9 | 92.2 |
| logBB | Predicted brain/blood partition coefficient[b] | -3.0 – 1.2 | -0.8 | -0.8 | -0.8 | -0.8 |
| PPMDCK | Predicted apparent MDCK cell permeability in nm/sec[a]. MDCK cells are considered to be a good mimic for the blood-brain barrier | <25 poor, >500 great | 95.2 | 125.4 | 123.2 | 121.6 |
| logKp | Predicted skin permeability | -8.0 – -1.0 | -4.5 | -4.4 | -4.6 | -4.7 |
| logK'hsa | Prediction of binding to human serum albumin | -1.5 – 1.5 | 0.1 | 0.1 | 0.0 | 0.0 |
| Oral Abs | Predicted human oral absorption on 0 to 100% scale. The prediction is based on a quantitative multiple linear regression model | >80% is high <25% is poor | 80 | 81 | 77 | 76 |

**Figure S12.** Metabolism of MRIA9. The compound was treated with activated microsomes and measured at 0, 15, 30 and 60 minutes via HPLC. Data are given as mean ± S.E.M. obtained in 3 independent experiments. Predicted ADME properties for the selected compounds. Predictions were generated using QikProp (Schrödinger, 2020) and compared with literature values. [a]Predictions are for non-active transport. [b]Predictions are for orally delivered drugs so, for example, dopamine and serotonin are CNS negative because they are too polar to cross the blood-brain barrier.

**Table S1**. Selectivity screening results of G-5555 (**6**) at 100 nM.

| DiscoveRx Gene Symbol | Entrez Gene Symbol | Percent Control | Compound Concentration (nM) |
|---|---|---|---|
| QSK | KIAA0999 | 0.2 | 100 |
| MST4 | MST4 | 0.5 | 100 |
| SIK | SIK1 | 1.4 | 100 |
| YANK2 | STK32B | 3.5 | 100 |
| PAK1 | PAK1 | 4,4 | 100 |
| YANK1 | STK32A | 5.3 | 100 |
| SIK2 | SIK2 | 6.5 | 100 |
| PFCDPK1 (*P. falciparum*) | CDPK1 | 9.1 | 100 |
| MEK5 | MAP2K5 | 13 | 100 |
| PAK2 | PAK2 | 13 | 100 |
| MST3 | STK24 | 15 | 100 |
| MAP4K5 | MAP4K5 | 18 | 100 |
| SNARK | NUAK2 | 19 | 100 |
| MEK1 | MAP2K1 | 24 | 100 |
| LTK | LTK | 28 | 100 |
| DCAMKL3 | DCLK3 | 29 | 100 |
| MEK2 | MAP2K2 | 33 | 100 |
| STK36 | STK36 | 36 | 100 |
| AURKC | AURKC | 37 | 100 |
| LOK | STK10 | 37 | 100 |
| YSK1 | STK25 | 40 | 100 |
| CAMK1D | CAMK1D | 43 | 100 |
| MARK3 | MARK3 | 45 | 100 |
| MAST1 | MAST1 | 47 | 100 |

| | | | |
|---|---|---|---|
| MYO3B | MYO3B | 48 | 100 |
| PIP5K1A | PIP5K1A | 48 | 100 |
| MEK4 | MAP2K4 | 49 | 100 |
| GAK | GAK | 51 | 100 |
| MELK | MELK | 51 | 100 |
| PIK3CD | PIK3CD | 53 | 100 |
| FLT3(ITD,F691L) | FLT3 | 56 | 100 |
| SgK110 | SgK110 | 56 | 100 |
| LATS1 | LATS1 | 57 | 100 |
| LATS2 | LATS2 | 58 | 100 |
| LCK | LCK | 58 | 100 |
| DYRK1B | DYRK1B | 61 | 100 |
| EPHB6 | EPHB6 | 61 | 100 |
| MAP4K2 | MAP4K2 | 63 | 100 |
| EPHA1 | EPHA1 | 64 | 100 |
| MST2 | STK3 | 64 | 100 |
| DRAK2 | STK17B | 66 | 100 |
| PIP5K2B | PIP4K2B | 68 | 100 |
| PRKD1 | PRKD1 | 68 | 100 |
| YSK4 | MAP3K19 | 70 | 100 |
| AURKB | AURKB | 72 | 100 |
| MAP3K15 | MAP3K15 | 72 | 100 |
| MRCKB | CDC42BPB | 73 | 100 |
| MST1R | MST1R | 73 | 100 |
| NLK | NLK | 73 | 100 |
| PIK3CA(H1047Y) | PIK3CA | 73 | 100 |
| MLCK | MYLK3 | 74 | 100 |

| | | | |
|---|---|---|---|
| MEK3 | MAP2K3 | 75 | 100 |
| MYLK | MYLK | 75 | 100 |
| PLK2 | PLK2 | 75 | 100 |
| GCN2(Kin.Dom.2,S808G) | EIF2AK4 | 76 | 100 |
| MET(M1250T) | MET | 76 | 100 |
| PIK3CA(M1043I) | PIK3CA | 76 | 100 |
| ERBB3 | ERBB3 | 77 | 100 |
| KIT(V559D,V654A) | KIT | 77 | 100 |
| MINK | MINK1 | 77 | 100 |
| TRKA | NTRK1 | 77 | 100 |
| HPK1 | MAP4K1 | 78 | 100 |
| ABL1(H396P)-nonphosphorylated | ABL1 | 79 | 100 |
| CAMK2A | CAMK2A | 79 | 100 |
| CDC2L5 | CDK13 | 79 | 100 |
| DRAK1 | STK17A | 79 | 100 |
| PIP5K2C | PIP4K2C | 79 | 100 |
| RSK1(Kin.Dom.1-N-terminal) | RPS6KA1 | 79 | 100 |
| DCAMKL1 | DCLK1 | 80 | 100 |
| JNK1 | MAPK8 | 80 | 100 |
| MEK6 | MAP2K6 | 80 | 100 |
| PIK3CA(C420R) | PIK3CA | 80 | 100 |
| KIT(A829P) | KIT | 81 | 100 |
| MAP3K1 | MAP3K1 | 81 | 100 |
| PRKCQ | PRKCQ | 81 | 100 |
| CDKL5 | CDKL5 | 82 | 100 |
| KIT-autoinhibited | KIT | 82 | 100 |

| | | | |
|---|---|---|---|
| PLK1 | PLK1 | 82 | 100 |
| PRKCH | PRKCH | 82 | 100 |
| GRK3 | ADRBK2 | 83 | 100 |
| LRRK2 | LRRK2 | 83 | 100 |
| MTOR | MTOR | 83 | 100 |
| ROCK1 | ROCK1 | 83 | 100 |
| RSK2(Kin.Dom.2-C-terminal) | RPS6KA3 | 83 | 100 |
| TESK1 | TESK1 | 83 | 100 |
| MAP3K4 | MAP3K4 | 84 | 100 |
| PLK4 | PLK4 | 84 | 100 |
| SNRK | SNRK | 84 | 100 |
| TRKC | NTRK3 | 84 | 100 |
| TYK2(JH2domain-pseudokinase) | TYK2 | 84 | 100 |
| FLT3(D835V) | FLT3 | 85 | 100 |
| MAP4K3 | MAP4K3 | 85 | 100 |
| PIK3CA(E545K) | PIK3CA | 85 | 100 |
| PIK3CA(Q546K) | PIK3CA | 85 | 100 |
| RIOK3 | RIOK3 | 85 | 100 |
| ABL1(Q252H)-nonphosphorylated | ABL1 | 86 | 100 |
| HIPK3 | HIPK3 | 86 | 100 |
| MAP3K3 | MAP3K3 | 86 | 100 |
| MKK7 | MAP2K7 | 86 | 100 |
| p38-delta | MAPK13 | 86 | 100 |
| PLK3 | PLK3 | 86 | 100 |
| PRKG2 | PRKG2 | 86 | 100 |
| RET(V804M) | RET | 86 | 100 |

| | | | |
|---|---|---|---|
| VPS34 | PIK3C3 | 86 | 100 |
| ABL1(F317I)-phosphorylated | ABL1 | 87 | 100 |
| ARK5 | NUAK1 | 87 | 100 |
| BTK | BTK | 87 | 100 |
| CDK11 | CDK19 | 87 | 100 |
| HASPIN | GSG2 | 87 | 100 |
| PIK3CA(I800L) | PIK3CA | 87 | 100 |
| PIK3CG | PIK3CG | 87 | 100 |
| TIE2 | TEK | 87 | 100 |
| CSNK2A1 | CSNK2A1 | 88 | 100 |
| DDR2 | DDR2 | 88 | 100 |
| FLT3(ITD,D835V) | FLT3 | 88 | 100 |
| PIK3C2B | PIK3C2B | 88 | 100 |
| PIK3CA(E545A) | PIK3CA | 88 | 100 |
| ALK | ALK | 89 | 100 |
| CDKL3 | CDKL3 | 89 | 100 |
| CSF1R | CSF1R | 89 | 100 |
| DMPK2 | CDC42BPG | 89 | 100 |
| EGFR(E746-A750del) | EGFR | 89 | 100 |
| FLT3(D835H) | FLT3 | 89 | 100 |
| GRK7 | GRK7 | 89 | 100 |
| MET | MET | 89 | 100 |
| NIM1 | MGC42105 | 89 | 100 |
| PIK3CA(E542K) | PIK3CA | 89 | 100 |
| SLK | SLK | 89 | 100 |
| ABL1(T315I)-nonphosphorylated | ABL1 | 90 | 100 |

| | | | |
|---|---|---|---|
| BMPR1A | BMPR1A | 90 | 100 |
| CSNK1D | CSNK1D | 90 | 100 |
| MAK | MAK | 90 | 100 |
| MST1 | STK4 | 90 | 100 |
| PFPK5 (*P. falciparum*) | MAL13P1.279 | 90 | 100 |
| RIPK5 | DSTYK | 90 | 100 |
| ULK3 | ULK3 | 90 | 100 |
| ANKK1 | ANKK1 | 91 | 100 |
| NEK10 | NEK10 | 91 | 100 |
| ERN1 | ERN1 | 92 | 100 |
| IKK-alpha | CHUK | 92 | 100 |
| JAK1(JH2domain-pseudokinase) | JAK1 | 92 | 100 |
| JNK3 | MAPK10 | 92 | 100 |
| PAK6 | PAK6 | 92 | 100 |
| PRKD2 | PRKD2 | 92 | 100 |
| S6K1 | RPS6KB1 | 92 | 100 |
| TNK2 | TNK2 | 92 | 100 |
| AMPK-alpha2 | PRKAA2 | 93 | 100 |
| IKK-beta | IKBKB | 93 | 100 |
| LIMK1 | LIMK1 | 93 | 100 |
| MYLK2 | MYLK2 | 93 | 100 |
| NEK3 | NEK3 | 93 | 100 |
| PRP4 | PRPF4B | 93 | 100 |
| SGK3 | SGK3 | 93 | 100 |
| ABL1-nonphosphorylated | ABL1 | 94 | 100 |
| FLT1 | FLT1 | 94 | 100 |
| FLT3(D835Y) | FLT3 | 94 | 100 |

| | | | |
|---|---|---|---|
| FLT3-autoinhibited | FLT3 | 94 | 100 |
| PIK3CA | PIK3CA | 94 | 100 |
| RET(V804L) | RET | 94 | 100 |
| ROCK2 | ROCK2 | 94 | 100 |
| RSK2(Kin.Dom.1-N-terminal) | RPS6KA3 | 94 | 100 |
| SRPK1 | SRPK1 | 94 | 100 |
| TYK2(JH1domain-catalytic) | TYK2 | 94 | 100 |
| VRK2 | VRK2 | 94 | 100 |
| ABL1(F317L)-phosphorylated | ABL1 | 95 | 100 |
| ABL1-phosphorylated | ABL1 | 95 | 100 |
| AKT3 | AKT3 | 95 | 100 |
| AMPK-alpha1 | PRKAA1 | 95 | 100 |
| BMPR1B | BMPR1B | 95 | 100 |
| HIPK4 | HIPK4 | 95 | 100 |
| INSR | INSR | 95 | 100 |
| IRAK3 | IRAK3 | 95 | 100 |
| IRAK4 | IRAK4 | 95 | 100 |
| MERTK | MERTK | 95 | 100 |
| PIK3C2G | PIK3C2G | 95 | 100 |
| PRKCE | PRKCE | 95 | 100 |
| PRKD3 | PRKD3 | 95 | 100 |
| CASK | CASK | 96 | 100 |
| CDK2 | CDK2 | 96 | 100 |
| CSNK2A2 | CSNK2A2 | 96 | 100 |
| FLT3(R834Q) | FLT3 | 96 | 100 |
| RET | RET | 96 | 100 |

| | | | |
|---|---|---|---|
| TYRO3 | TYRO3 | 96 | 100 |
| WNK4 | WNK4 | 96 | 100 |
| ABL1(H396P)-phosphorylated | ABL1 | 97 | 100 |
| ABL1(Q252H)-phosphorylated | ABL1 | 97 | 100 |
| ABL1(Y253F)-phosphorylated | ABL1 | 97 | 100 |
| ACVR1 | ACVR1 | 97 | 100 |
| CAMK1B | PNCK | 97 | 100 |
| CLK4 | CLK4 | 97 | 100 |
| CSF1R-autoinhibited | CSF1R | 97 | 100 |
| EIF2AK1 | EIF2AK1 | 97 | 100 |
| GRK2 | ADRBK1 | 97 | 100 |
| GSK3B | GSK3B | 97 | 100 |
| IRAK1 | IRAK1 | 97 | 100 |
| MAP3K2 | MAP3K2 | 97 | 100 |
| NDR1 | STK38 | 97 | 100 |
| p38-beta | MAPK11 | 97 | 100 |
| PDGFRB | PDGFRB | 97 | 100 |
| PIK4CB | PI4KB | 97 | 100 |
| PRKCI | PRKCI | 97 | 100 |
| RIPK4 | RIPK4 | 97 | 100 |
| RSK4(Kin.Dom.1-N-terminal) | RPS6KA6 | 97 | 100 |
| ZAK | ZAK | 97 | 100 |
| AURKA | AURKA | 98 | 100 |
| ERK5 | MAPK7 | 98 | 100 |
| FAK | PTK2 | 98 | 100 |

| | | | |
|---|---|---|---|
| KIT(V559D,T670I) | KIT | 98 | 100 |
| SBK1 | SBK1 | 98 | 100 |
| SRPK2 | SRPK2 | 98 | 100 |
| TSSK1B | TSSK1B | 98 | 100 |
| VEGFR2 | KDR | 98 | 100 |
| WEE1 | WEE1 | 98 | 100 |
| ADCK4 | ADCK4 | 99 | 100 |
| ASK1 | MAP3K5 | 99 | 100 |
| EPHA3 | EPHA3 | 99 | 100 |
| FGFR3(G697C) | FGFR3 | 99 | 100 |
| JAK2(JH1domain-catalytic) | JAK2 | 99 | 100 |
| KIT(V559D) | KIT | 99 | 100 |
| OSR1 | OXSR1 | 99 | 100 |
| PDPK1 | PDPK1 | 99 | 100 |
| SGK | SGK1 | 99 | 100 |
| SRMS | SRMS | 99 | 100 |
| TAOK2 | TAOK2 | 99 | 100 |
| TGFBR1 | TGFBR1 | 99 | 100 |
| YANK3 | STK32C | 99 | 100 |
| AAK1 | AAK1 | 100 | 100 |
| ABL1(E255K)-phosphorylated | ABL1 | 100 | 100 |
| ABL1(F317I)-nonphosphorylated | ABL1 | 100 | 100 |
| ABL1(F317L)-nonphosphorylated | ABL1 | 100 | 100 |
| ABL1(M351T)-phosphorylated | ABL1 | 100 | 100 |
| ABL1(T315I)- | ABL1 | 100 | 100 |

| | | | |
|---|---|---|---|
| phosphorylated | | | |
| ABL2 | ABL2 | 100 | 100 |
| ACVR1B | ACVR1B | 100 | 100 |
| ACVR2A | ACVR2A | 100 | 100 |
| ACVR2B | ACVR2B | 100 | 100 |
| ACVRL1 | ACVRL1 | 100 | 100 |
| ADCK3 | CABC1 | 100 | 100 |
| AKT1 | AKT1 | 100 | 100 |
| AKT2 | AKT2 | 100 | 100 |
| ALK(C1156Y) | ALK | 100 | 100 |
| ALK(L1196M) | ALK | 100 | 100 |
| ASK2 | MAP3K6 | 100 | 100 |
| AXL | AXL | 100 | 100 |
| BIKE | BMP2K | 100 | 100 |
| BLK | BLK | 100 | 100 |
| BMPR2 | BMPR2 | 100 | 100 |
| BMX | BMX | 100 | 100 |
| BRAF | BRAF | 100 | 100 |
| BRAF(V600E) | BRAF | 100 | 100 |
| BRK | PTK6 | 100 | 100 |
| BRSK1 | BRSK1 | 100 | 100 |
| BRSK2 | BRSK2 | 100 | 100 |
| BUB1 | BUB1 | 100 | 100 |
| CAMK1 | CAMK1 | 100 | 100 |
| CAMK1G | CAMK1G | 100 | 100 |
| CAMK2B | CAMK2B | 100 | 100 |
| CAMK2D | CAMK2D | 100 | 100 |

| | | | |
|---|---|---|---|
| CAMK2G | CAMK2G | 100 | 100 |
| CAMK4 | CAMK4 | 100 | 100 |
| CAMKK1 | CAMKK1 | 100 | 100 |
| CAMKK2 | CAMKK2 | 100 | 100 |
| CDC2L1 | CDK11B | 100 | 100 |
| CDC2L2 | CDC2L2 | 100 | 100 |
| CDK3 | CDK3 | 100 | 100 |
| CDK4 | CDK4 | 100 | 100 |
| CDK4-cyclinD1 | CDK4 | 100 | 100 |
| CDK4-cyclinD3 | CDK4 | 100 | 100 |
| CDK5 | CDK5 | 100 | 100 |
| CDK7 | CDK7 | 100 | 100 |
| CDK8 | CDK8 | 100 | 100 |
| CDK9 | CDK9 | 100 | 100 |
| CDKL1 | CDKL1 | 100 | 100 |
| CDKL2 | CDKL2 | 100 | 100 |
| CHEK1 | CHEK1 | 100 | 100 |
| CHEK2 | CHEK2 | 100 | 100 |
| CIT | CIT | 100 | 100 |
| CLK1 | CLK1 | 100 | 100 |
| CLK2 | CLK2 | 100 | 100 |
| CLK3 | CLK3 | 100 | 100 |
| CSK | CSK | 100 | 100 |
| CSNK1A1 | CSNK1A1 | 100 | 100 |
| CSNK1A1L | CSNK1A1L | 100 | 100 |
| CSNK1E | CSNK1E | 100 | 100 |
| CSNK1G1 | CSNK1G1 | 100 | 100 |

| | | | |
|---|---|---|---|
| CSNK1G2 | CSNK1G2 | 100 | 100 |
| CSNK1G3 | CSNK1G3 | 100 | 100 |
| CTK | MATK | 100 | 100 |
| DAPK1 | DAPK1 | 100 | 100 |
| DAPK2 | DAPK2 | 100 | 100 |
| DAPK3 | DAPK3 | 100 | 100 |
| DCAMKL2 | DCLK2 | 100 | 100 |
| DDR1 | DDR1 | 100 | 100 |
| DLK | MAP3K12 | 100 | 100 |
| DMPK | DMPK | 100 | 100 |
| DYRK1A | DYRK1A | 100 | 100 |
| DYRK2 | DYRK2 | 100 | 100 |
| EGFR | EGFR | 100 | 100 |
| EGFR(G719C) | EGFR | 100 | 100 |
| EGFR(G719S) | EGFR | 100 | 100 |
| EGFR(L747-E749del, A750P) | EGFR | 100 | 100 |
| EGFR(L747-S752del, P753S) | EGFR | 100 | 100 |
| EGFR(L747-T751del,Sins) | EGFR | 100 | 100 |
| EGFR(L858R) | EGFR | 100 | 100 |
| EGFR(L858R,T790M) | EGFR | 100 | 100 |
| EGFR(L861Q) | EGFR | 100 | 100 |
| EGFR(S752-I759del) | EGFR | 100 | 100 |
| EGFR(T790M) | EGFR | 100 | 100 |
| EPHA2 | EPHA2 | 100 | 100 |
| EPHA4 | EPHA4 | 100 | 100 |
| EPHA5 | EPHA5 | 100 | 100 |

| | | | |
|---|---|---|---|
| EPHA6 | EPHA6 | 100 | 100 |
| EPHA7 | EPHA7 | 100 | 100 |
| EPHA8 | EPHA8 | 100 | 100 |
| EPHB1 | EPHB1 | 100 | 100 |
| EPHB2 | EPHB2 | 100 | 100 |
| EPHB3 | EPHB3 | 100 | 100 |
| EPHB4 | EPHB4 | 100 | 100 |
| ERBB2 | ERBB2 | 100 | 100 |
| ERBB4 | ERBB4 | 100 | 100 |
| ERK1 | MAPK3 | 100 | 100 |
| ERK2 | MAPK1 | 100 | 100 |
| ERK3 | MAPK6 | 100 | 100 |
| ERK4 | MAPK4 | 100 | 100 |
| ERK8 | MAPK15 | 100 | 100 |
| FER | FER | 100 | 100 |
| FES | FES | 100 | 100 |
| FGFR1 | FGFR1 | 100 | 100 |
| FGFR2 | FGFR2 | 100 | 100 |
| FGFR3 | FGFR3 | 100 | 100 |
| FGFR4 | FGFR4 | 100 | 100 |
| FGR | FGR | 100 | 100 |
| FLT3 | FLT3 | 100 | 100 |
| FLT3(ITD) | FLT3 | 100 | 100 |
| FLT3(K663Q) | FLT3 | 100 | 100 |
| FLT3(N841I) | FLT3 | 100 | 100 |
| FLT4 | FLT4 | 100 | 100 |
| FRK | FRK | 100 | 100 |

| | | | |
|---|---|---|---|
| FYN | FYN | 100 | 100 |
| GRK1 | GRK1 | 100 | 100 |
| GRK4 | GRK4 | 100 | 100 |
| GSK3A | GSK3A | 100 | 100 |
| HCK | HCK | 100 | 100 |
| HIPK1 | HIPK1 | 100 | 100 |
| HIPK2 | HIPK2 | 100 | 100 |
| HUNK | HUNK | 100 | 100 |
| ICK | ICK | 100 | 100 |
| IGF1R | IGF1R | 100 | 100 |
| IKK-epsilon | IKBKE | 100 | 100 |
| INSRR | INSRR | 100 | 100 |
| ITK | ITK | 100 | 100 |
| JAK1(JH1domain-catalytic) | JAK1 | 100 | 100 |
| JAK3(JH1domain-catalytic) | JAK3 | 100 | 100 |
| JNK2 | MAPK9 | 100 | 100 |
| KIT | KIT | 100 | 100 |
| KIT(D816H) | KIT | 100 | 100 |
| KIT(D816V) | KIT | 100 | 100 |
| KIT(L576P) | KIT | 100 | 100 |
| LIMK2 | LIMK2 | 100 | 100 |
| LKB1 | STK11 | 100 | 100 |
| LRRK2(G2019S) | LRRK2 | 100 | 100 |
| LYN | LYN | 100 | 100 |
| LZK | MAP3K13 | 100 | 100 |
| MAP4K4 | MAP4K4 | 100 | 100 |

| | | | |
|---|---|---|---|
| MAPKAPK2 | MAPKAPK2 | 100 | 100 |
| MAPKAPK5 | MAPKAPK5 | 100 | 100 |
| MARK1 | MARK1 | 100 | 100 |
| MARK2 | MARK2 | 100 | 100 |
| MARK4 | MARK4 | 100 | 100 |
| MET(Y1235D) | MET | 100 | 100 |
| MKNK1 | MKNK1 | 100 | 100 |
| MKNK2 | MKNK2 | 100 | 100 |
| MLK1 | MAP3K9 | 100 | 100 |
| MLK2 | MAP3K10 | 100 | 100 |
| MLK3 | MAP3K11 | 100 | 100 |
| MRCKA | CDC42BPA | 100 | 100 |
| MUSK | MUSK | 100 | 100 |
| MYLK4 | MYLK4 | 100 | 100 |
| MYO3A | MYO3A | 100 | 100 |
| NDR2 | STK38L | 100 | 100 |
| NEK1 | NEK1 | 100 | 100 |
| NEK11 | NEK11 | 100 | 100 |
| NEK2 | NEK2 | 100 | 100 |
| NEK4 | NEK4 | 100 | 100 |
| NEK5 | NEK5 | 100 | 100 |
| NEK6 | NEK6 | 100 | 100 |
| NEK7 | NEK7 | 100 | 100 |
| NEK9 | NEK9 | 100 | 100 |
| NIK | MAP3K14 | 100 | 100 |
| p38-alpha | MAPK14 | 100 | 100 |
| p38-gamma | MAPK12 | 100 | 100 |

| | | | |
|---|---|---|---|
| PAK3 | PAK3 | 100 | 100 |
| PAK4 | PAK4 | 100 | 100 |
| PAK7 | PAK7 | 100 | 100 |
| PCTK1 | CDK16 | 100 | 100 |
| PCTK2 | CDK17 | 100 | 100 |
| PCTK3 | CDK18 | 100 | 100 |
| PDGFRA | PDGFRA | 100 | 100 |
| PFTAIRE2 | CDK15 | 100 | 100 |
| PFTK1 | CDK14 | 100 | 100 |
| PHKG1 | PHKG1 | 100 | 100 |
| PHKG2 | PHKG2 | 100 | 100 |
| PIK3CA(H1047L) | PIK3CA | 100 | 100 |
| PIK3CB | PIK3CB | 100 | 100 |
| PIKFYVE | PIKFYVE | 100 | 100 |
| PIM1 | PIM1 | 100 | 100 |
| PIM2 | PIM2 | 100 | 100 |
| PIM3 | PIM3 | 100 | 100 |
| PIP5K1C | PIP5K1C | 100 | 100 |
| PKAC-alpha | PRKACA | 100 | 100 |
| PKAC-beta | PRKACB | 100 | 100 |
| PKMYT1 | PKMYT1 | 100 | 100 |
| PKN1 | PKN1 | 100 | 100 |
| PKN2 | PKN2 | 100 | 100 |
| PKNB (*M. tuberculosis*) | pknB | 100 | 100 |
| PRKCD | PRKCD | 100 | 100 |
| PRKG1 | PRKG1 | 100 | 100 |
| PRKR | EIF2AK2 | 100 | 100 |

| | | | |
|---|---|---|---|
| PRKX | PRKX | 100 | 100 |
| PYK2 | PTK2B | 100 | 100 |
| RAF1 | RAF1 | 100 | 100 |
| RET(M918T) | RET | 100 | 100 |
| RIOK1 | RIOK1 | 100 | 100 |
| RIOK2 | RIOK2 | 100 | 100 |
| RIPK1 | RIPK1 | 100 | 100 |
| RIPK2 | RIPK2 | 100 | 100 |
| ROS1 | ROS1 | 100 | 100 |
| RPS6KA4(Kin.Dom.1-N-terminal) | RPS6KA4 | 100 | 100 |
| RPS6KA4(Kin.Dom.2-C-terminal) | RPS6KA4 | 100 | 100 |
| RPS6KA5(Kin.Dom.1-N-terminal) | RPS6KA5 | 100 | 100 |
| RPS6KA5(Kin.Dom.2-C-terminal) | RPS6KA5 | 100 | 100 |
| RSK1(Kin.Dom.2-C-terminal) | RPS6KA1 | 100 | 100 |
| RSK3(Kin.Dom.1-N-terminal) | RPS6KA2 | 100 | 100 |
| RSK3(Kin.Dom.2-C-terminal) | RPS6KA2 | 100 | 100 |
| RSK4(Kin.Dom.2-C-terminal) | RPS6KA6 | 100 | 100 |
| SGK2 | SGK2 | 100 | 100 |
| SRC | SRC | 100 | 100 |
| SRPK3 | SRPK3 | 100 | 100 |
| STK16 | STK16 | 100 | 100 |
| STK33 | STK33 | 100 | 100 |
| STK35 | STK35 | 100 | 100 |

| | | | |
|---|---|---|---|
| STK39 | STK39 | 100 | 100 |
| SYK | SYK | 100 | 100 |
| TAK1 | MAP3K7 | 100 | 100 |
| TAOK1 | TAOK1 | 100 | 100 |
| TAOK3 | TAOK3 | 100 | 100 |
| TBK1 | TBK1 | 100 | 100 |
| TEC | TEC | 100 | 100 |
| TGFBR2 | TGFBR2 | 100 | 100 |
| TIE1 | TIE1 | 100 | 100 |
| TLK1 | TLK1 | 100 | 100 |
| TLK2 | TLK2 | 100 | 100 |
| TNIK | TNIK | 100 | 100 |
| TNK1 | TNK1 | 100 | 100 |
| TNNI3K | TNNI3K | 100 | 100 |
| TRKB | NTRK2 | 100 | 100 |
| TRPM6 | TRPM6 | 100 | 100 |
| TSSK3 | TSSK3 | 100 | 100 |
| TTK | TTK | 100 | 100 |
| TXK | TXK | 100 | 100 |
| ULK1 | ULK1 | 100 | 100 |
| ULK2 | ULK2 | 100 | 100 |
| WEE2 | WEE2 | 100 | 100 |
| WNK1 | WNK1 | 100 | 100 |
| WNK2 | WNK2 | 100 | 100 |
| WNK3 | WNK3 | 100 | 100 |
| YES | YES1 | 100 | 100 |
| ZAP70 | ZAP70 | 100 | 100 |

**Table S2**. Selectivity screening results of G-5555 (**6**) at 1 μM.

| DiscoveRx Gene Symbol | Entrez Gene Symbol | Percent Control | Compound Concentration (nM) |
|---|---|---|---|
| QSK | KIAA0999 | 0 | 1000 |
| MST4 | MST4 | 0 | 1000 |
| YANK2 | STK32B | 0 | 1000 |
| PAK2 | PAK2 | 0 | 1000 |
| MEK5 | MAP2K5 | 0.2 | 1000 |
| PFCDPK1 (*P. falciparum*) | CDPK1 | 0.4 | 1000 |
| MAST1 | MAST1 | 0.4 | 1000 |
| DCAMKL3 | DCLK3 | 0.5 | 1000 |
| SIK | SIK1 | 0.6 | 1000 |
| SIK2 | SIK2 | 0.8 | 1000 |
| MEK1 | MAP2K1 | 1 | 1000 |
| MEK2 | MAP2K2 | 1.4 | 1000 |
| MST3 | STK24 | 1.7 | 1000 |
| GAK | GAK | 1.7 | 1000 |
| SNARK | NUAK2 | 2 | 1000 |
| LOK | STK10 | 2.1 | 1000 |
| CAMK1D | CAMK1D | 2.5 | 1000 |
| STK36 | STK36 | 2.9 | 1000 |
| YANK1 | STK32A | 3.7 | 1000 |
| LTK | LTK | 4.1 | 1000 |
| PAK1 | PAK1 | 4.5 | 1000 |
| MAP4K5 | MAP4K5 | 5.3 | 1000 |
| MRCKB | CDC42BPB | 5.3 | 1000 |
| LATS2 | LATS2 | 8 | 1000 |

| | | | |
|---|---|---|---|
| EPHB6 | EPHB6 | 9 | 1000 |
| MEK4 | MAP2K4 | 9.6 | 1000 |
| LCK | LCK | 11 | 1000 |
| DMPK2 | CDC42BPG | 15 | 1000 |
| PRKD1 | PRKD1 | 17 | 1000 |
| NLK | NLK | 17 | 1000 |
| MARK3 | MARK3 | 18 | 1000 |
| WNK4 | WNK4 | 18 | 1000 |
| YSK4 | MAP3K19 | 19 | 1000 |
| AURKC | AURKC | 20 | 1000 |
| PRKD3 | PRKD3 | 20 | 1000 |
| AURKB | AURKB | 23 | 1000 |
| MAP4K3 | MAP4K3 | 23 | 1000 |
| MAP3K3 | MAP3K3 | 23 | 1000 |
| ARK5 | NUAK1 | 24 | 1000 |
| MINK | MINK1 | 26 | 1000 |
| HPK1 | MAP4K1 | 26 | 1000 |
| PIP5K2C | PIP4K2C | 26 | 1000 |
| MST2 | STK3 | 27 | 1000 |
| PRKD2 | PRKD2 | 27 | 1000 |
| MEK3 | MAP2K3 | 29 | 1000 |
| MEK6 | MAP2K6 | 30 | 1000 |
| CAMK1B | PNCK | 30 | 1000 |
| MAP4K2 | MAP4K2 | 32 | 1000 |
| SLK | SLK | 33 | 1000 |
| EPHA1 | EPHA1 | 34 | 1000 |
| YSK1 | STK25 | 36 | 1000 |

| | | | |
|---|---|---|---|
| MET | MET | 40 | 1000 |
| AMPK-alpha1 | PRKAA1 | 40 | 1000 |
| LATS1 | LATS1 | 41 | 1000 |
| JNK1 | MAPK8 | 41 | 1000 |
| FLT3(ITD,F691L) | FLT3 | 43 | 1000 |
| AURKA | AURKA | 43 | 1000 |
| MET(M1250T) | MET | 45 | 1000 |
| PAK6 | PAK6 | 45 | 1000 |
| MELK | MELK | 47 | 1000 |
| AMPK-alpha2 | PRKAA2 | 47 | 1000 |
| MAP3K15 | MAP3K15 | 48 | 1000 |
| MST1R | MST1R | 48 | 1000 |
| AKT3 | AKT3 | 48 | 1000 |
| DYRK1B | DYRK1B | 49 | 1000 |
| IRAK1 | IRAK1 | 49 | 1000 |
| MARK2 | MARK2 | 49 | 1000 |
| MST1 | STK4 | 51 | 1000 |
| DCAMKL2 | DCLK2 | 51 | 1000 |
| PKNB (*M. tuberculosis*) | pknB | 51 | 1000 |
| TNIK | TNIK | 51 | 1000 |
| MAP3K4 | MAP3K4 | 52 | 1000 |
| TIE2 | TEK | 52 | 1000 |
| ULK3 | ULK3 | 52 | 1000 |
| ERK5 | MAPK7 | 52 | 1000 |
| CSNK1E | CSNK1E | 52 | 1000 |
| DDR1 | DDR1 | 53 | 1000 |
| PRKCQ | PRKCQ | 55 | 1000 |

| | | | |
|---|---|---|---|
| ABL1(H396P)-nonphosphorylated | ABL1 | 56 | 1000 |
| MYO3B | MYO3B | 57 | 1000 |
| RSK3(Kin.Dom.1-N-terminal) | RPS6KA2 | 57 | 1000 |
| S6K1 | RPS6KB1 | 58 | 1000 |
| CDKL2 | CDKL2 | 58 | 1000 |
| LIMK1 | LIMK1 | 59 | 1000 |
| CAMK1 | CAMK1 | 59 | 1000 |
| SGK3 | SGK3 | 60 | 1000 |
| NEK2 | NEK2 | 60 | 1000 |
| GCN2(Kin.Dom.2,S808G) | EIF2AK4 | 63 | 1000 |
| KIT(A829P) | KIT | 63 | 1000 |
| RSK2(Kin.Dom.2-C-terminal) | RPS6KA3 | 63 | 1000 |
| BTK | BTK | 63 | 1000 |
| PRKCI | PRKCI | 63 | 1000 |
| TRKA | NTRK1 | 64 | 1000 |
| RSK2(Kin.Dom.1-N-terminal) | RPS6KA3 | 65 | 1000 |
| RIPK2 | RIPK2 | 65 | 1000 |
| PIP5K1A | PIP5K1A | 67 | 1000 |
| PLK4 | PLK4 | 67 | 1000 |
| FLT3(D835V) | FLT3 | 67 | 1000 |
| MAP3K2 | MAP3K2 | 67 | 1000 |
| MARK1 | MARK1 | 68 | 1000 |
| TBK1 | TBK1 | 68 | 1000 |
| RSK1(Kin.Dom.1-N-terminal) | RPS6KA1 | 69 | 1000 |
| STK16 | STK16 | 69 | 1000 |

| | | | |
|---|---|---|---|
| WNK1 | WNK1 | 69 | 1000 |
| NDR1 | STK38 | 70 | 1000 |
| SRPK2 | SRPK2 | 70 | 1000 |
| ASK2 | MAP3K6 | 70 | 1000 |
| MRCKA | CDC42BPA | 70 | 1000 |
| SgK110 | SgK110 | 71 | 1000 |
| PRKCE | PRKCE | 71 | 1000 |
| ASK1 | MAP3K5 | 71 | 1000 |
| PDPK1 | PDPK1 | 71 | 1000 |
| FRK | FRK | 71 | 1000 |
| GRK3 | ADRBK2 | 72 | 1000 |
| NEK3 | NEK3 | 72 | 1000 |
| EPHA8 | EPHA8 | 72 | 1000 |
| FLT3(ITD,D835V) | FLT3 | 74 | 1000 |
| TYK2(JH1domain-catalytic) | TYK2 | 74 | 1000 |
| ERK1 | MAPK3 | 74 | 1000 |
| LYN | LYN | 74 | 1000 |
| ABL1(Q252H)-nonphosphorylated | ABL1 | 75 | 1000 |
| JNK2 | MAPK9 | 75 | 1000 |
| STK33 | STK33 | 75 | 1000 |
| MKK7 | MAP2K7 | 77 | 1000 |
| PAK4 | PAK4 | 77 | 1000 |
| PRKCH | PRKCH | 78 | 1000 |
| IKK-epsilon | IKBKE | 78 | 1000 |
| DCAMKL1 | DCLK1 | 79 | 1000 |
| YANK3 | STK32C | 79 | 1000 |

| | | | |
|---|---|---|---|
| EPHB2 | EPHB2 | 79 | 1000 |
| IKK-beta | IKBKB | 80 | 1000 |
| KIT-autoinhibited | KIT | 81 | 1000 |
| MYLK2 | MYLK2 | 81 | 1000 |
| FLT3(D835Y) | FLT3 | 81 | 1000 |
| RSK1(Kin.Dom.2-C-terminal) | RPS6KA1 | 81 | 1000 |
| PAK7 | PAK7 | 82 | 1000 |
| MTOR | MTOR | 83 | 1000 |
| JAK2(JH1domain-catalytic) | JAK2 | 83 | 1000 |
| CSNK1G2 | CSNK1G2 | 83 | 1000 |
| PLK2 | PLK2 | 84 | 1000 |
| ALK | ALK | 84 | 1000 |
| CSF1R-autoinhibited | CSF1R | 84 | 1000 |
| PDGFRB | PDGFRB | 84 | 1000 |
| EGFR(L747-E749del, A750P) | EGFR | 84 | 1000 |
| FES | FES | 84 | 1000 |
| TSSK3 | TSSK3 | 84 | 1000 |
| MYLK | MYLK | 85 | 1000 |
| BMPR1A | BMPR1A | 85 | 1000 |
| ABL1-nonphosphorylated | ABL1 | 85 | 1000 |
| CSK | CSK | 85 | 1000 |
| EGFR(L858R) | EGFR | 85 | 1000 |
| EGFR(T790M) | EGFR | 85 | 1000 |
| ULK1 | ULK1 | 85 | 1000 |
| PIK3CA(H1047Y) | PIK3CA | 86 | 1000 |
| ERBB3 | ERBB3 | 86 | 1000 |

| | | | |
|---|---|---|---|
| TNK2 | TNK2 | 86 | 1000 |
| CDK7 | CDK7 | 86 | 1000 |
| ERK2 | MAPK1 | 86 | 1000 |
| PIP5K2B | PIP4K2B | 87 | 1000 |
| ANKK1 | ANKK1 | 87 | 1000 |
| JNK3 | MAPK10 | 87 | 1000 |
| ABL1(F317L)-nonphosphorylated | ABL1 | 87 | 1000 |
| FYN | FYN | 87 | 1000 |
| TAOK1 | TAOK1 | 87 | 1000 |
| CAMK2A | CAMK2A | 88 | 1000 |
| EPHA3 | EPHA3 | 88 | 1000 |
| EGFR | EGFR | 88 | 1000 |
| MLK2 | MAP3K10 | 88 | 1000 |
| WNK2 | WNK2 | 88 | 1000 |
| SNRK | SNRK | 89 | 1000 |
| GRK7 | GRK7 | 89 | 1000 |
| BUB1 | BUB1 | 89 | 1000 |
| CSNK1A1 | CSNK1A1 | 89 | 1000 |
| RIOK3 | RIOK3 | 90 | 1000 |
| PIK3CA(E545A) | PIK3CA | 90 | 1000 |
| ROCK2 | ROCK2 | 90 | 1000 |
| JAK3(JH1domain-catalytic) | JAK3 | 90 | 1000 |
| RIPK5 | DSTYK | 91 | 1000 |
| CSF1R | CSF1R | 92 | 1000 |
| JAK1(JH2domain-pseudokinase) | JAK1 | 92 | 1000 |
| IRAK3 | IRAK3 | 92 | 1000 |

| | | | |
|---|---|---|---|
| TYRO3 | TYRO3 | 92 | 1000 |
| PKN1 | PKN1 | 92 | 1000 |
| RSK3(Kin.Dom.2-C-terminal) | RPS6KA2 | 92 | 1000 |
| RSK4(Kin.Dom.2-C-terminal) | RPS6KA6 | 92 | 1000 |
| TAK1 | MAP3K7 | 92 | 1000 |
| PLK3 | PLK3 | 93 | 1000 |
| PIK3CG | PIK3CG | 93 | 1000 |
| VEGFR2 | KDR | 93 | 1000 |
| BRAF(V600E) | BRAF | 93 | 1000 |
| MET(Y1235D) | MET | 93 | 1000 |
| PLK1 | PLK1 | 94 | 1000 |
| PRKG2 | PRKG2 | 94 | 1000 |
| RET | RET | 94 | 1000 |
| ABL1(H396P)-phosphorylated | ABL1 | 94 | 1000 |
| PRKR | EIF2AK2 | 94 | 1000 |
| TIE1 | TIE1 | 94 | 1000 |
| PRP4 | PRPF4B | 95 | 1000 |
| ABL1-phosphorylated | ABL1 | 95 | 1000 |
| IRAK4 | IRAK4 | 95 | 1000 |
| CSNK2A2 | CSNK2A2 | 95 | 1000 |
| RSK4(Kin.Dom.1-N-terminal) | RPS6KA6 | 95 | 1000 |
| PKAC-beta | PRKACB | 95 | 1000 |
| TRKC | NTRK3 | 96 | 1000 |
| ABL1(F317I)-phosphorylated | ABL1 | 96 | 1000 |
| BMPR1B | BMPR1B | 96 | 1000 |

| | | | |
|---|---|---|---|
| ACVR1B | ACVR1B | 96 | 1000 |
| AXL | AXL | 96 | 1000 |
| CAMK4 | CAMK4 | 96 | 1000 |
| MLK1 | MAP3K9 | 96 | 1000 |
| MYO3A | MYO3A | 96 | 1000 |
| PDGFRA | PDGFRA | 96 | 1000 |
| PIK3CD | PIK3CD | 97 | 1000 |
| CDC2L5 | CDK13 | 97 | 1000 |
| CDKL5 | CDKL5 | 97 | 1000 |
| PIK3CA(E545K) | PIK3CA | 97 | 1000 |
| INSR | INSR | 97 | 1000 |
| CDC2L1 | CDK11B | 97 | 1000 |
| FLT3(N841I) | FLT3 | 97 | 1000 |
| MKNK2 | MKNK2 | 97 | 1000 |
| PIK3CB | PIK3CB | 97 | 1000 |
| TTK | TTK | 97 | 1000 |
| MAP3K1 | MAP3K1 | 98 | 1000 |
| TYK2(JH2domain-pseudokinase) | TYK2 | 98 | 1000 |
| CSNK2A1 | CSNK2A1 | 98 | 1000 |
| IKK-alpha | CHUK | 98 | 1000 |
| SRPK1 | SRPK1 | 98 | 1000 |
| CDK5 | CDK5 | 98 | 1000 |
| TNNI3K | TNNI3K | 98 | 1000 |
| TESK1 | TESK1 | 99 | 1000 |
| FLT3(D835H) | FLT3 | 99 | 1000 |
| BLK | BLK | 99 | 1000 |
| CAMK1G | CAMK1G | 99 | 1000 |

| | | | |
|---|---|---|---|
| CDK4-cyclinD1 | CDK4 | 99 | 1000 |
| DRAK2 | STK17B | 100 | 1000 |
| MLCK | MYLK3 | 100 | 1000 |
| PIK3CA(M1043I) | PIK3CA | 100 | 1000 |
| KIT(V559D,V654A) | KIT | 100 | 1000 |
| DRAK1 | STK17A | 100 | 1000 |
| PIK3CA(C420R) | PIK3CA | 100 | 1000 |
| LRRK2 | LRRK2 | 100 | 1000 |
| ROCK1 | ROCK1 | 100 | 1000 |
| PIK3CA(Q546K) | PIK3CA | 100 | 1000 |
| HIPK3 | HIPK3 | 100 | 1000 |
| p38-delta | MAPK13 | 100 | 1000 |
| RET(V804M) | RET | 100 | 1000 |
| VPS34 | PIK3C3 | 100 | 1000 |
| CDK11 | CDK19 | 100 | 1000 |
| HASPIN | GSG2 | 100 | 1000 |
| PIK3CA(I800L) | PIK3CA | 100 | 1000 |
| DDR2 | DDR2 | 100 | 1000 |
| PIK3C2B | PIK3C2B | 100 | 1000 |
| CDKL3 | CDKL3 | 100 | 1000 |
| EGFR(E746-A750del) | EGFR | 100 | 1000 |
| NIM1 | MGC42105 | 100 | 1000 |
| PIK3CA(E542K) | PIK3CA | 100 | 1000 |
| ABL1(T315I)-nonphosphorylated | ABL1 | 100 | 1000 |
| CSNK1D | CSNK1D | 100 | 1000 |
| MAK | MAK | 100 | 1000 |
| PFPK5(*P. falciparum*) | MAL13P1.279 | 100 | 1000 |

| | | | |
|---|---|---|---|
| NEK10 | NEK10 | 100 | 1000 |
| ERN1 | ERN1 | 100 | 1000 |
| FLT1 | FLT1 | 100 | 1000 |
| FLT3-autoinhibited | FLT3 | 100 | 1000 |
| PIK3CA | PIK3CA | 100 | 1000 |
| RET(V804L) | RET | 100 | 1000 |
| VRK2 | VRK2 | 100 | 1000 |
| ABL1(F317L)-phosphorylated | ABL1 | 100 | 1000 |
| HIPK4 | HIPK4 | 100 | 1000 |
| MERTK | MERTK | 100 | 1000 |
| PIK3C2G | PIK3C2G | 100 | 1000 |
| CASK | CASK | 100 | 1000 |
| CDK2 | CDK2 | 100 | 1000 |
| FLT3(R834Q) | FLT3 | 100 | 1000 |
| ABL1(Q252H)-phosphorylated | ABL1 | 100 | 1000 |
| ABL1(Y253F)-phosphorylated | ABL1 | 100 | 1000 |
| ACVR1 | ACVR1 | 100 | 1000 |
| CLK4 | CLK4 | 100 | 1000 |
| EIF2AK1 | EIF2AK1 | 100 | 1000 |
| GRK2 | ADRBK1 | 100 | 1000 |
| GSK3B | GSK3B | 100 | 1000 |
| p38-beta | MAPK11 | 100 | 1000 |
| PIK4CB | PI4KB | 100 | 1000 |
| RIPK4 | RIPK4 | 100 | 1000 |
| ZAK | ZAK | 100 | 1000 |
| FAK | PTK2 | 100 | 1000 |

| | | | |
|---|---|---|---|
| KIT(V559D,T670I) | KIT | 100 | 1000 |
| SBK1 | SBK1 | 100 | 1000 |
| TSSK1B | TSSK1B | 100 | 1000 |
| WEE1 | WEE1 | 100 | 1000 |
| ADCK4 | ADCK4 | 100 | 1000 |
| FGFR3(G697C) | FGFR3 | 100 | 1000 |
| KIT(V559D) | KIT | 100 | 1000 |
| OSR1 | OXSR1 | 100 | 1000 |
| SGK | SGK1 | 100 | 1000 |
| SRMS | SRMS | 100 | 1000 |
| TAOK2 | TAOK2 | 100 | 1000 |
| TGFBR1 | TGFBR1 | 100 | 1000 |
| AAK1 | AAK1 | 100 | 1000 |
| ABL1(E255K)-phosphorylated | ABL1 | 100 | 1000 |
| ABL1(F317I)-nonphosphorylated | ABL1 | 100 | 1000 |
| ABL1(M351T)-phosphorylated | ABL1 | 100 | 1000 |
| ABL1(T315I)-phosphorylated | ABL1 | 100 | 1000 |
| ABL2 | ABL2 | 100 | 1000 |
| ACVR2A | ACVR2A | 100 | 1000 |
| ACVR2B | ACVR2B | 100 | 1000 |
| ACVRL1 | ACVRL1 | 100 | 1000 |
| ADCK3 | CABC1 | 100 | 1000 |
| AKT1 | AKT1 | 100 | 1000 |
| AKT2 | AKT2 | 100 | 1000 |
| ALK(C1156Y) | ALK | 100 | 1000 |

| | | | |
|---|---|---|---|
| ALK(L1196M) | ALK | 100 | 1000 |
| BIKE | BMP2K | 100 | 1000 |
| BMPR2 | BMPR2 | 100 | 1000 |
| BMX | BMX | 100 | 1000 |
| BRAF | BRAF | 100 | 1000 |
| BRK | PTK6 | 100 | 1000 |
| BRSK1 | BRSK1 | 100 | 1000 |
| BRSK2 | BRSK2 | 100 | 1000 |
| CAMK2B | CAMK2B | 100 | 1000 |
| CAMK2D | CAMK2D | 100 | 1000 |
| CAMK2G | CAMK2G | 100 | 1000 |
| CAMKK1 | CAMKK1 | 100 | 1000 |
| CAMKK2 | CAMKK2 | 100 | 1000 |
| CDC2L2 | CDC2L2 | 100 | 1000 |
| CDK3 | CDK3 | 100 | 1000 |
| CDK4 | CDK4 | 100 | 1000 |
| CDK4-cyclinD3 | CDK4 | 100 | 1000 |
| CDK8 | CDK8 | 100 | 1000 |
| CDK9 | CDK9 | 100 | 1000 |
| CDKL1 | CDKL1 | 100 | 1000 |
| CHEK1 | CHEK1 | 100 | 1000 |
| CHEK2 | CHEK2 | 100 | 1000 |
| CIT | CIT | 100 | 1000 |
| CLK1 | CLK1 | 100 | 1000 |
| CLK2 | CLK2 | 100 | 1000 |
| CLK3 | CLK3 | 100 | 1000 |
| CSNK1A1L | CSNK1A1L | 100 | 1000 |

| | | | |
|---|---|---|---|
| CSNK1G1 | CSNK1G1 | 100 | 1000 |
| CSNK1G3 | CSNK1G3 | 100 | 1000 |
| CTK | MATK | 100 | 1000 |
| DAPK1 | DAPK1 | 100 | 1000 |
| DAPK2 | DAPK2 | 100 | 1000 |
| DAPK3 | DAPK3 | 100 | 1000 |
| DLK | MAP3K12 | 100 | 1000 |
| DMPK | DMPK | 100 | 1000 |
| DYRK1A | DYRK1A | 100 | 1000 |
| DYRK2 | DYRK2 | 100 | 1000 |
| EGFR(G719C) | EGFR | 100 | 1000 |
| EGFR(G719S) | EGFR | 100 | 1000 |
| EGFR(L747-S752del, P753S) | EGFR | 100 | 1000 |
| EGFR(L747-T751del,Sins) | EGFR | 100 | 1000 |
| EGFR(L858R,T790M) | EGFR | 100 | 1000 |
| EGFR(L861Q) | EGFR | 100 | 1000 |
| EGFR(S752-I759del) | EGFR | 100 | 1000 |
| EPHA2 | EPHA2 | 100 | 1000 |
| EPHA4 | EPHA4 | 100 | 1000 |
| EPHA5 | EPHA5 | 100 | 1000 |
| EPHA6 | EPHA6 | 100 | 1000 |
| EPHA7 | EPHA7 | 100 | 1000 |
| EPHB1 | EPHB1 | 100 | 1000 |
| EPHB3 | EPHB3 | 100 | 1000 |
| EPHB4 | EPHB4 | 100 | 1000 |
| ERBB2 | ERBB2 | 100 | 1000 |

| | | | |
|---|---|---|---|
| ERBB4 | ERBB4 | 100 | 1000 |
| ERK3 | MAPK6 | 100 | 1000 |
| ERK4 | MAPK4 | 100 | 1000 |
| ERK8 | MAPK15 | 100 | 1000 |
| FER | FER | 100 | 1000 |
| FGFR1 | FGFR1 | 100 | 1000 |
| FGFR2 | FGFR2 | 100 | 1000 |
| FGFR3 | FGFR3 | 100 | 1000 |
| FGFR4 | FGFR4 | 100 | 1000 |
| FGR | FGR | 100 | 1000 |
| FLT3 | FLT3 | 100 | 1000 |
| FLT3(ITD) | FLT3 | 100 | 1000 |
| FLT3(K663Q) | FLT3 | 100 | 1000 |
| FLT4 | FLT4 | 100 | 1000 |
| GRK1 | GRK1 | 100 | 1000 |
| GRK4 | GRK4 | 100 | 1000 |
| GSK3A | GSK3A | 100 | 1000 |
| HCK | HCK | 100 | 1000 |
| HIPK1 | HIPK1 | 100 | 1000 |
| HIPK2 | HIPK2 | 100 | 1000 |
| HUNK | HUNK | 100 | 1000 |
| ICK | ICK | 100 | 1000 |
| IGF1R | IGF1R | 100 | 1000 |
| INSRR | INSRR | 100 | 1000 |
| ITK | ITK | 100 | 1000 |
| JAK1(JH1domain-catalytic) | JAK1 | 100 | 1000 |
| KIT | KIT | 100 | 1000 |

| | | | |
|---|---|---|---|
| KIT(D816H) | KIT | 100 | 1000 |
| KIT(D816V) | KIT | 100 | 1000 |
| KIT(L576P) | KIT | 100 | 1000 |
| LIMK2 | LIMK2 | 100 | 1000 |
| LKB1 | STK11 | 100 | 1000 |
| LRRK2(G2019S) | LRRK2 | 100 | 1000 |
| LZK | MAP3K13 | 100 | 1000 |
| MAP4K4 | MAP4K4 | 100 | 1000 |
| MAPKAPK2 | MAPKAPK2 | 100 | 1000 |
| MAPKAPK5 | MAPKAPK5 | 100 | 1000 |
| MARK4 | MARK4 | 100 | 1000 |
| MKNK1 | MKNK1 | 100 | 1000 |
| MLK3 | MAP3K11 | 100 | 1000 |
| MUSK | MUSK | 100 | 1000 |
| MYLK4 | MYLK4 | 100 | 1000 |
| NDR2 | STK38L | 100 | 1000 |
| NEK1 | NEK1 | 100 | 1000 |
| NEK11 | NEK11 | 100 | 1000 |
| NEK4 | NEK4 | 100 | 1000 |
| NEK5 | NEK5 | 100 | 1000 |
| NEK6 | NEK6 | 100 | 1000 |
| NEK7 | NEK7 | 100 | 1000 |
| NEK9 | NEK9 | 100 | 1000 |
| NIK | MAP3K14 | 100 | 1000 |
| p38-alpha | MAPK14 | 100 | 1000 |
| p38-gamma | MAPK12 | 100 | 1000 |
| PAK3 | PAK3 | 100 | 1000 |

| | | | |
|---|---|---|---|
| PCTK1 | CDK16 | 100 | 1000 |
| PCTK2 | CDK17 | 100 | 1000 |
| PCTK3 | CDK18 | 100 | 1000 |
| PFTAIRE2 | CDK15 | 100 | 1000 |
| PFTK1 | CDK14 | 100 | 1000 |
| PHKG1 | PHKG1 | 100 | 1000 |
| PHKG2 | PHKG2 | 100 | 1000 |
| PIK3CA(H1047L) | PIK3CA | 100 | 1000 |
| PIKFYVE | PIKFYVE | 100 | 1000 |
| PIM1 | PIM1 | 100 | 1000 |
| PIM2 | PIM2 | 100 | 1000 |
| PIM3 | PIM3 | 100 | 1000 |
| PIP5K1C | PIP5K1C | 100 | 1000 |
| PKAC-alpha | PRKACA | 100 | 1000 |
| PKMYT1 | PKMYT1 | 100 | 1000 |
| PKN2 | PKN2 | 100 | 1000 |
| PRKCD | PRKCD | 100 | 1000 |
| PRKG1 | PRKG1 | 100 | 1000 |
| PRKX | PRKX | 100 | 1000 |
| PYK2 | PTK2B | 100 | 1000 |
| RAF1 | RAF1 | 100 | 1000 |
| RET(M918T) | RET | 100 | 1000 |
| RIOK1 | RIOK1 | 100 | 1000 |
| RIOK2 | RIOK2 | 100 | 1000 |
| RIPK1 | RIPK1 | 100 | 1000 |
| ROS1 | ROS1 | 100 | 1000 |
| RPS6KA4(Kin.Dom.1-N-terminal) | RPS6KA4 | 100 | 1000 |

| | | | |
|---|---|---|---|
| RPS6KA4(Kin.Dom.2-C-terminal) | RPS6KA4 | 100 | 1000 |
| RPS6KA5(Kin.Dom.1-N-terminal) | RPS6KA5 | 100 | 1000 |
| RPS6KA5(Kin.Dom.2-C-terminal) | RPS6KA5 | 100 | 1000 |
| SGK2 | SGK2 | 100 | 1000 |
| SRC | SRC | 100 | 1000 |
| SRPK3 | SRPK3 | 100 | 1000 |
| STK35 | STK35 | 100 | 1000 |
| STK39 | STK39 | 100 | 1000 |
| SYK | SYK | 100 | 1000 |
| TAOK3 | TAOK3 | 100 | 1000 |
| TEC | TEC | 100 | 1000 |
| TGFBR2 | TGFBR2 | 100 | 1000 |
| TLK1 | TLK1 | 100 | 1000 |
| TLK2 | TLK2 | 100 | 1000 |
| TNK1 | TNK1 | 100 | 1000 |
| TRKB | NTRK2 | 100 | 1000 |
| TRPM6 | TRPM6 | 100 | 1000 |
| TXK | TXK | 100 | 1000 |
| ULK2 | ULK2 | 100 | 1000 |
| WEE2 | WEE2 | 100 | 1000 |
| WNK3 | WNK3 | 100 | 1000 |
| YES | YES1 | 100 | 1000 |
| ZAP70 | ZAP70 | 100 | 1000 |

**Table S3**. Selectivity screening results for **7** at 1μM.

| DiscoveRx Gene Symbol | Entrez Gene Symbol | Percent Control | Compound Concentration (nM) |
|---|---|---|---|
| KIT(A829P) | KIT | 0 | 1000 |
| MST4 | MST4 | 0 | 1000 |
| PAK2 | PAK2 | 0 | 1000 |
| YANK2 | STK32B | 0 | 1000 |
| MEK5 | MAP2K5 | 0.2 | 1000 |
| MEK4 | MAP2K4 | 0.3 | 1000 |
| CAMK1D | CAMK1D | 0.4 | 1000 |
| SIK | SIK1 | 0.7 | 1000 |
| MAP4K5 | MAP4K5 | 0.9 | 1000 |
| GAK | GAK | 1 | 1000 |
| SIK2 | SIK2 | 1 | 1000 |
| LOK | STK10 | 1.2 | 1000 |
| YANK1 | STK32A | 1.4 | 1000 |
| EPHB6 | EPHB6 | 1.6 | 1000 |
| MST3 | STK24 | 1.7 | 1000 |
| CAMK1B | PNCK | 2.2 | 1000 |
| LTK | LTK | 2.6 | 1000 |
| TNK2 | TNK2 | 3.5 | 1000 |
| ARK5 | NUAK1 | 3.9 | 1000 |
| PAK1 | PAK1 | 3.9 | 1000 |
| FES | FES | 4.7 | 1000 |
| STK36 | STK36 | 4.7 | 1000 |
| FER | FER | 5.2 | 1000 |
| SNARK | NUAK2 | 5.5 | 1000 |

| | | | |
|---|---|---|---|
| FRK | FRK | 5.8 | 1000 |
| DCAMKL2 | DCLK2 | 6 | 1000 |
| PFCDPK1 (*P. falciparum*) | CDPK1 | 6.6 | 1000 |
| FAK | PTK2 | 7.1 | 1000 |
| PDGFRB | PDGFRB | 8 | 1000 |
| KIT(V559D) | KIT | 8.4 | 1000 |
| YANK3 | STK32C | 8.5 | 1000 |
| PLK4 | PLK4 | 9.1 | 1000 |
| LCK | LCK | 9.3 | 1000 |
| MAP4K3 | MAP4K3 | 10 | 1000 |
| MEK1 | MAP2K1 | 11 | 1000 |
| MRCKB | CDC42BPB | 11 | 1000 |
| NLK | NLK | 11 | 1000 |
| CSF1R | CSF1R | 12 | 1000 |
| PLK3 | PLK3 | 12 | 1000 |
| PYK2 | PTK2B | 13 | 1000 |
| MEK2 | MAP2K2 | 14 | 1000 |
| MINK | MINK1 | 14 | 1000 |
| PRKD3 | PRKD3 | 14 | 1000 |
| MAP4K4 | MAP4K4 | 15 | 1000 |
| WNK4 | WNK4 | 15 | 1000 |
| JNK1 | MAPK8 | 16 | 1000 |
| KIT(L576P) | KIT | 16 | 1000 |
| MAST1 | MAST1 | 16 | 1000 |
| CAMK1 | CAMK1 | 17 | 1000 |
| EGFR(L861Q) | EGFR | 17 | 1000 |
| HPK1 | MAP4K1 | 17 | 1000 |

| | | | |
|---|---|---|---|
| MARK3 | MARK3 | 17 | 1000 |
| AURKB | AURKB | 18 | 1000 |
| SLK | SLK | 18 | 1000 |
| AURKC | AURKC | 19 | 1000 |
| EGFR(G719C) | EGFR | 19 | 1000 |
| EGFR(L858R) | EGFR | 19 | 1000 |
| ERK5 | MAPK7 | 19 | 1000 |
| KIT(D816V) | KIT | 20 | 1000 |
| PIP5K2C | PIP4K2C | 20 | 1000 |
| KIT | KIT | 22 | 1000 |
| TNK1 | TNK1 | 22 | 1000 |
| EPHA1 | EPHA1 | 23 | 1000 |
| ERBB4 | ERBB4 | 23 | 1000 |
| TNIK | TNIK | 23 | 1000 |
| MST1 | STK4 | 24 | 1000 |
| BLK | BLK | 25 | 1000 |
| MEK6 | MAP2K6 | 26 | 1000 |
| PRKD1 | PRKD1 | 26 | 1000 |
| MEK3 | MAP2K3 | 28 | 1000 |
| MST2 | STK3 | 28 | 1000 |
| DMPK2 | CDC42BPG | 29 | 1000 |
| LATS2 | LATS2 | 29 | 1000 |
| PRKD2 | PRKD2 | 31 | 1000 |
| PAK3 | PAK3 | 32 | 1000 |
| MELK | MELK | 33 | 1000 |
| MYO3A | MYO3A | 33 | 1000 |
| AURKA | AURKA | 34 | 1000 |

| | | | |
|---|---|---|---|
| MYO3B | MYO3B | 34 | 1000 |
| YSK1 | STK25 | 36 | 1000 |
| ABL1(F317L)-phosphorylated | ABL1 | 37 | 1000 |
| QSK | KIAA0999 | 38 | 1000 |
| LIMK1 | LIMK1 | 40 | 1000 |
| MLK1 | MAP3K9 | 40 | 1000 |
| HCK | HCK | 41 | 1000 |
| KIT(D816H) | KIT | 41 | 1000 |
| MAP3K3 | MAP3K3 | 42 | 1000 |
| AMPK-alpha1 | PRKAA1 | 43 | 1000 |
| DCAMKL3 | DCLK3 | 43 | 1000 |
| ERBB2 | ERBB2 | 44 | 1000 |
| SYK | SYK | 45 | 1000 |
| MAP4K2 | MAP4K2 | 46 | 1000 |
| ABL1(F317L)-nonphosphorylated | ABL1 | 48 | 1000 |
| LATS1 | LATS1 | 48 | 1000 |
| LYN | LYN | 48 | 1000 |
| PDGFRA | PDGFRA | 48 | 1000 |
| EGFR(G719S) | EGFR | 50 | 1000 |
| EGFR(T790M) | EGFR | 50 | 1000 |
| FLT3(D835Y) | FLT3 | 50 | 1000 |
| WNK2 | WNK2 | 50 | 1000 |
| FLT3(K663Q) | FLT3 | 52 | 1000 |
| ABL1(H396P)-nonphosphorylated | ABL1 | 53 | 1000 |
| DCAMKL1 | DCLK1 | 53 | 1000 |
| FLT3 | FLT3 | 53 | 1000 |

| | | | |
|---|---|---|---|
| JNK2 | MAPK9 | 53 | 1000 |
| MAP3K2 | MAP3K2 | 53 | 1000 |
| MAP3K4 | MAP3K4 | 54 | 1000 |
| MARK1 | MARK1 | 56 | 1000 |
| ABL1(F317I)-phosphorylated | ABL1 | 57 | 1000 |
| ERK1 | MAPK3 | 57 | 1000 |
| MKK7 | MAP2K7 | 57 | 1000 |
| EGFR | EGFR | 58 | 1000 |
| JNK3 | MAPK10 | 58 | 1000 |
| FLT3(D835V) | FLT3 | 59 | 1000 |
| TESK1 | TESK1 | 59 | 1000 |
| ERK2 | MAPK1 | 60 | 1000 |
| MARK2 | MARK2 | 60 | 1000 |
| VEGFR2 | KDR | 60 | 1000 |
| DDR2 | DDR2 | 61 | 1000 |
| TSSK1B | TSSK1B | 61 | 1000 |
| ULK3 | ULK3 | 61 | 1000 |
| NEK3 | NEK3 | 62 | 1000 |
| RET | RET | 62 | 1000 |
| CSNK1E | CSNK1E | 63 | 1000 |
| EGFR(L747-S752del, P753S) | EGFR | 63 | 1000 |
| RSK2(Kin.Dom.1-N-terminal) | RPS6KA3 | 63 | 1000 |
| TRKA | NTRK1 | 63 | 1000 |
| PLK2 | PLK2 | 64 | 1000 |
| SRPK2 | SRPK2 | 64 | 1000 |
| STK33 | STK33 | 64 | 1000 |

| | | | |
|---|---|---|---|
| ABL1(Y253F)-phosphorylated | ABL1 | 65 | 1000 |
| PKN2 | PKN2 | 65 | 1000 |
| CDK4 | CDK4 | 66 | 1000 |
| EGFR(E746-A750del) | EGFR | 66 | 1000 |
| FLT3(ITD,F691L) | FLT3 | 66 | 1000 |
| YSK4 | MAP3K19 | 66 | 1000 |
| TIE2 | TEK | 68 | 1000 |
| FGR | FGR | 69 | 1000 |
| PIP5K1A | PIP5K1A | 69 | 1000 |
| LRRK2 | LRRK2 | 70 | 1000 |
| EGFR(L858R,T790M) | EGFR | 71 | 1000 |
| FLT3(N841I) | FLT3 | 71 | 1000 |
| FLT3(R834Q) | FLT3 | 71 | 1000 |
| INSR | INSR | 71 | 1000 |
| KIT-autoinhibited | KIT | 71 | 1000 |
| MAP3K15 | MAP3K15 | 71 | 1000 |
| MARK4 | MARK4 | 71 | 1000 |
| NDR1 | STK38 | 71 | 1000 |
| TTK | TTK | 71 | 1000 |
| ADCK4 | ADCK4 | 72 | 1000 |
| CAMK2A | CAMK2A | 72 | 1000 |
| GRK1 | GRK1 | 72 | 1000 |
| RSK2(Kin.Dom.2-C-terminal) | RPS6KA3 | 72 | 1000 |
| BRK | PTK6 | 73 | 1000 |
| BUB1 | BUB1 | 73 | 1000 |
| FLT3-autoinhibited | FLT3 | 73 | 1000 |

| | | | |
|---|---|---|---|
| NEK10 | NEK10 | 73 | 1000 |
| ROCK2 | ROCK2 | 73 | 1000 |
| WNK1 | WNK1 | 73 | 1000 |
| ABL1(F317I)-nonphosphorylated | ABL1 | 74 | 1000 |
| BTK | BTK | 74 | 1000 |
| NEK2 | NEK2 | 74 | 1000 |
| PDPK1 | PDPK1 | 74 | 1000 |
| TSSK3 | TSSK3 | 74 | 1000 |
| ABL1(Q252H)-nonphosphorylated | ABL1 | 75 | 1000 |
| ANKK1 | ANKK1 | 75 | 1000 |
| DDR1 | DDR1 | 75 | 1000 |
| JAK1(JH2domain-pseudokinase) | JAK1 | 75 | 1000 |
| MKNK2 | MKNK2 | 75 | 1000 |
| PAK6 | PAK6 | 75 | 1000 |
| PAK7 | PAK7 | 75 | 1000 |
| SGK2 | SGK2 | 75 | 1000 |
| SNRK | SNRK | 75 | 1000 |
| ULK1 | ULK1 | 75 | 1000 |
| ABL1(M351T)-phosphorylated | ABL1 | 76 | 1000 |
| ABL2 | ABL2 | 76 | 1000 |
| HASPIN | GSG2 | 76 | 1000 |
| PIK3CA(Q546K) | PIK3CA | 76 | 1000 |
| RIPK5 | DSTYK | 76 | 1000 |
| ROCK1 | ROCK1 | 76 | 1000 |
| RSK4(Kin.Dom.1-N-terminal) | RPS6KA6 | 76 | 1000 |

| | | | |
|---|---|---|---|
| WNK3 | WNK3 | 76 | 1000 |
| AMPK-alpha2 | PRKAA2 | 77 | 1000 |
| CHEK2 | CHEK2 | 77 | 1000 |
| EGFR(L747-T751del,Sins) | EGFR | 77 | 1000 |
| IKK-alpha | CHUK | 77 | 1000 |
| PFTAIRE2 | CDK15 | 77 | 1000 |
| GRK7 | GRK7 | 78 | 1000 |
| MRCKA | CDC42BPA | 78 | 1000 |
| MYLK | MYLK | 78 | 1000 |
| PCTK2 | CDK17 | 78 | 1000 |
| ABL1-nonphosphorylated | ABL1 | 79 | 1000 |
| CHEK1 | CHEK1 | 79 | 1000 |
| JAK2(JH1domain-catalytic) | JAK2 | 79 | 1000 |
| MTOR | MTOR | 79 | 1000 |
| NEK11 | NEK11 | 79 | 1000 |
| PHKG2 | PHKG2 | 79 | 1000 |
| PLK1 | PLK1 | 79 | 1000 |
| SGK | SGK1 | 79 | 1000 |
| STK39 | STK39 | 79 | 1000 |
| VPS34 | PIK3C3 | 79 | 1000 |
| ABL1(T315I)-phosphorylated | ABL1 | 80 | 1000 |
| CAMK1G | CAMK1G | 80 | 1000 |
| ERBB3 | ERBB3 | 80 | 1000 |
| RSK3(Kin.Dom.2-C-terminal) | RPS6KA2 | 80 | 1000 |
| EPHA8 | EPHA8 | 81 | 1000 |

| | | | |
|---|---|---|---|
| FLT3(D835H) | FLT3 | 81 | 1000 |
| PFPK5 (P*. falciparum*) | MAL13P1.279 | 81 | 1000 |
| RSK3(Kin.Dom.1-N-terminal) | RPS6KA2 | 81 | 1000 |
| BMPR1B | BMPR1B | 82 | 1000 |
| NIM1 | MGC42105 | 82 | 1000 |
| PRP4 | PRPF4B | 82 | 1000 |
| RSK4(Kin.Dom.2-C-terminal) | RPS6KA6 | 82 | 1000 |
| SGK3 | SGK3 | 82 | 1000 |
| TBK1 | TBK1 | 82 | 1000 |
| VRK2 | VRK2 | 82 | 1000 |
| ABL1(T315I)-nonphosphorylated | ABL1 | 83 | 1000 |
| EIF2AK1 | EIF2AK1 | 83 | 1000 |
| PAK4 | PAK4 | 83 | 1000 |
| PIK3CA(E545K) | PIK3CA | 83 | 1000 |
| PIK3CB | PIK3CB | 83 | 1000 |
| ABL1-phosphorylated | ABL1 | 84 | 1000 |
| AXL | AXL | 84 | 1000 |
| ERK4 | MAPK4 | 84 | 1000 |
| NDR2 | STK38L | 84 | 1000 |
| S6K1 | RPS6KB1 | 84 | 1000 |
| TXK | TXK | 84 | 1000 |
| ULK2 | ULK2 | 84 | 1000 |
| YES | YES1 | 84 | 1000 |
| CSF1R-autoinhibited | CSF1R | 85 | 1000 |
| CSNK1A1L | CSNK1A1L | 85 | 1000 |
| TRKC | NTRK3 | 85 | 1000 |

| | | | |
|---|---|---|---|
| TYRO3 | TYRO3 | 85 | 1000 |
| CDC2L5 | CDK13 | 86 | 1000 |
| CDK9 | CDK9 | 86 | 1000 |
| IRAK1 | IRAK1 | 86 | 1000 |
| PIP5K1C | PIP5K1C | 86 | 1000 |
| TYK2(JH1domain-catalytic) | TYK2 | 86 | 1000 |
| ABL1(Q252H)-phosphorylated | ABL1 | 87 | 1000 |
| CSK | CSK | 87 | 1000 |
| MLCK | MYLK3 | 87 | 1000 |
| CDKL5 | CDKL5 | 88 | 1000 |
| CSNK2A1 | CSNK2A1 | 88 | 1000 |
| ERN1 | ERN1 | 88 | 1000 |
| FLT3(ITD,D835V) | FLT3 | 88 | 1000 |
| GCN2(Kin.Dom.2,S808G) | EIF2AK4 | 88 | 1000 |
| OSR1 | OXSR1 | 88 | 1000 |
| PRKG2 | PRKG2 | 88 | 1000 |
| RIOK3 | RIOK3 | 88 | 1000 |
| RSK1(Kin.Dom.2-C-terminal) | RPS6KA1 | 88 | 1000 |
| TRPM6 | TRPM6 | 88 | 1000 |
| IGF1R | IGF1R | 89 | 1000 |
| PHKG1 | PHKG1 | 89 | 1000 |
| PIK3CG | PIK3CG | 89 | 1000 |
| RET(M918T) | RET | 89 | 1000 |
| TEC | TEC | 89 | 1000 |
| DAPK2 | DAPK2 | 90 | 1000 |
| IRAK4 | IRAK4 | 90 | 1000 |

| | | | |
|---|---|---|---|
| JAK3(JH1domain-catalytic) | JAK3 | 90 | 1000 |
| MLK3 | MAP3K11 | 90 | 1000 |
| PIK3CA(I800L) | PIK3CA | 90 | 1000 |
| ABL1(E255K)-phosphorylated | ABL1 | 91 | 1000 |
| ABL1(H396P)-phosphorylated | ABL1 | 91 | 1000 |
| CDC2L1 | CDK11B | 91 | 1000 |
| EGFR(S752-I759del) | EGFR | 91 | 1000 |
| HIPK2 | HIPK2 | 91 | 1000 |
| IKK-beta | IKBKB | 91 | 1000 |
| ITK | ITK | 91 | 1000 |
| KIT(V559D,V654A) | KIT | 91 | 1000 |
| LRRK2(G2019S) | LRRK2 | 91 | 1000 |
| MET | MET | 91 | 1000 |
| MUSK | MUSK | 91 | 1000 |
| TIE1 | TIE1 | 91 | 1000 |
| CLK1 | CLK1 | 92 | 1000 |
| DYRK1A | DYRK1A | 92 | 1000 |
| EPHB1 | EPHB1 | 92 | 1000 |
| GRK3 | ADRBK2 | 92 | 1000 |
| PIK3CA(E542K) | PIK3CA | 92 | 1000 |
| CSNK1G2 | CSNK1G2 | 93 | 1000 |
| DAPK3 | DAPK3 | 93 | 1000 |
| FLT4 | FLT4 | 93 | 1000 |
| MST1R | MST1R | 93 | 1000 |
| PRKCI | PRKCI | 93 | 1000 |
| TYK2(JH2domain- | TYK2 | 93 | 1000 |

| | | | |
|---|---|---|---|
| pseudokinase) | | | |
| CAMK2D | CAMK2D | 94 | 1000 |
| CAMK2G | CAMK2G | 94 | 1000 |
| CAMKK2 | CAMKK2 | 94 | 1000 |
| LZK | MAP3K13 | 94 | 1000 |
| PIK3CA(H1047L) | PIK3CA | 94 | 1000 |
| PRKCE | PRKCE | 94 | 1000 |
| RSK1(Kin.Dom.1-N-terminal) | RPS6KA1 | 94 | 1000 |
| STK16 | STK16 | 94 | 1000 |
| TLK1 | TLK1 | 94 | 1000 |
| CDK7 | CDK7 | 95 | 1000 |
| DAPK1 | DAPK1 | 95 | 1000 |
| MET(M1250T) | MET | 95 | 1000 |
| NEK7 | NEK7 | 95 | 1000 |
| PIK3C2B | PIK3C2B | 95 | 1000 |
| RPS6KA5(Kin.Dom.2-C-terminal) | RPS6KA5 | 95 | 1000 |
| CSNK2A2 | CSNK2A2 | 96 | 1000 |
| DRAK2 | STK17B | 96 | 1000 |
| PRKX | PRKX | 96 | 1000 |
| RIPK4 | RIPK4 | 96 | 1000 |
| BMPR2 | BMPR2 | 97 | 1000 |
| CDK8 | CDK8 | 97 | 1000 |
| CSNK1A1 | CSNK1A1 | 97 | 1000 |
| DLK | MAP3K12 | 97 | 1000 |
| GRK2 | ADRBK1 | 97 | 1000 |
| IKK-epsilon | IKBKE | 97 | 1000 |

| | | | |
|---|---|---|---|
| INSRR | INSRR | 97 | 1000 |
| PIK3CA | PIK3CA | 97 | 1000 |
| RPS6KA4(Kin.Dom.2-C-terminal) | RPS6KA4 | 97 | 1000 |
| CAMKK1 | CAMKK1 | 98 | 1000 |
| FYN | FYN | 98 | 1000 |
| NEK6 | NEK6 | 98 | 1000 |
| p38-delta | MAPK13 | 98 | 1000 |
| SgK110 | SgK110 | 98 | 1000 |
| WEE2 | WEE2 | 98 | 1000 |
| CAMK2B | CAMK2B | 99 | 1000 |
| HIPK4 | HIPK4 | 99 | 1000 |
| LIMK2 | LIMK2 | 99 | 1000 |
| MAPKAPK5 | MAPKAPK5 | 99 | 1000 |
| PIK3CA(C420R) | PIK3CA | 99 | 1000 |
| PKMYT1 | PKMYT1 | 99 | 1000 |
| SRC | SRC | 99 | 1000 |
| AAK1 | AAK1 | 100 | 1000 |
| ACVR1 | ACVR1 | 100 | 1000 |
| ACVR1B | ACVR1B | 100 | 1000 |
| ACVR2A | ACVR2A | 100 | 1000 |
| ACVR2B | ACVR2B | 100 | 1000 |
| ACVRL1 | ACVRL1 | 100 | 1000 |
| ADCK3 | CABC1 | 100 | 1000 |
| AKT1 | AKT1 | 100 | 1000 |
| AKT2 | AKT2 | 100 | 1000 |
| AKT3 | AKT3 | 100 | 1000 |
| ALK | ALK | 100 | 1000 |

| | | | |
|---|---|---|---|
| ALK(C1156Y) | ALK | 100 | 1000 |
| ALK(L1196M) | ALK | 100 | 1000 |
| ASK1 | MAP3K5 | 100 | 1000 |
| ASK2 | MAP3K6 | 100 | 1000 |
| BIKE | BMP2K | 100 | 1000 |
| BMPR1A | BMPR1A | 100 | 1000 |
| BMX | BMX | 100 | 1000 |
| BRAF | BRAF | 100 | 1000 |
| BRAF(V600E) | BRAF | 100 | 1000 |
| BRSK1 | BRSK1 | 100 | 1000 |
| BRSK2 | BRSK2 | 100 | 1000 |
| CAMK4 | CAMK4 | 100 | 1000 |
| CASK | CASK | 100 | 1000 |
| CDC2L2 | CDC2L2 | 100 | 1000 |
| CDK11 | CDK19 | 100 | 1000 |
| CDK2 | CDK2 | 100 | 1000 |
| CDK3 | CDK3 | 100 | 1000 |
| CDK4-cyclinD1 | CDK4 | 100 | 1000 |
| CDK4-cyclinD3 | CDK4 | 100 | 1000 |
| CDK5 | CDK5 | 100 | 1000 |
| CDKL1 | CDKL1 | 100 | 1000 |
| CDKL2 | CDKL2 | 100 | 1000 |
| CDKL3 | CDKL3 | 100 | 1000 |
| CIT | CIT | 100 | 1000 |
| CLK2 | CLK2 | 100 | 1000 |
| CLK3 | CLK3 | 100 | 1000 |
| CLK4 | CLK4 | 100 | 1000 |

| | | | |
|---|---|---|---|
| CSNK1D | CSNK1D | 100 | 1000 |
| CSNK1G1 | CSNK1G1 | 100 | 1000 |
| CSNK1G3 | CSNK1G3 | 100 | 1000 |
| CTK | MATK | 100 | 1000 |
| DMPK | DMPK | 100 | 1000 |
| DRAK1 | STK17A | 100 | 1000 |
| DYRK1B | DYRK1B | 100 | 1000 |
| DYRK2 | DYRK2 | 100 | 1000 |
| EGFR(L747-E749del, A750P) | EGFR | 100 | 1000 |
| EPHA2 | EPHA2 | 100 | 1000 |
| EPHA3 | EPHA3 | 100 | 1000 |
| EPHA4 | EPHA4 | 100 | 1000 |
| EPHA5 | EPHA5 | 100 | 1000 |
| EPHA6 | EPHA6 | 100 | 1000 |
| EPHA7 | EPHA7 | 100 | 1000 |
| EPHB2 | EPHB2 | 100 | 1000 |
| EPHB3 | EPHB3 | 100 | 1000 |
| EPHB4 | EPHB4 | 100 | 1000 |
| ERK3 | MAPK6 | 100 | 1000 |
| ERK8 | MAPK15 | 100 | 1000 |
| FGFR1 | FGFR1 | 100 | 1000 |
| FGFR2 | FGFR2 | 100 | 1000 |
| FGFR3 | FGFR3 | 100 | 1000 |
| FGFR3(G697C) | FGFR3 | 100 | 1000 |
| FGFR4 | FGFR4 | 100 | 1000 |
| FLT1 | FLT1 | 100 | 1000 |
| FLT3(ITD) | FLT3 | 100 | 1000 |

| | | | |
|---|---|---|---|
| GRK4 | GRK4 | 100 | 1000 |
| GSK3A | GSK3A | 100 | 1000 |
| GSK3B | GSK3B | 100 | 1000 |
| HIPK1 | HIPK1 | 100 | 1000 |
| HIPK3 | HIPK3 | 100 | 1000 |
| HUNK | HUNK | 100 | 1000 |
| ICK | ICK | 100 | 1000 |
| IRAK3 | IRAK3 | 100 | 1000 |
| JAK1(JH1domain-catalytic) | JAK1 | 100 | 1000 |
| KIT(V559D,T670I) | KIT | 100 | 1000 |
| LKB1 | STK11 | 100 | 1000 |
| MAK | MAK | 100 | 1000 |
| MAP3K1 | MAP3K1 | 100 | 1000 |
| MAPKAPK2 | MAPKAPK2 | 100 | 1000 |
| MERTK | MERTK | 100 | 1000 |
| MET(Y1235D) | MET | 100 | 1000 |
| MKNK1 | MKNK1 | 100 | 1000 |
| MLK2 | MAP3K10 | 100 | 1000 |
| MYLK2 | MYLK2 | 100 | 1000 |
| MYLK4 | MYLK4 | 100 | 1000 |
| NEK1 | NEK1 | 100 | 1000 |
| NEK4 | NEK4 | 100 | 1000 |
| NEK5 | NEK5 | 100 | 1000 |
| NEK9 | NEK9 | 100 | 1000 |
| NIK | MAP3K14 | 100 | 1000 |
| p38-alpha | MAPK14 | 100 | 1000 |
| p38-beta | MAPK11 | 100 | 1000 |

| | | | |
|---|---|---|---|
| p38-gamma | MAPK12 | 100 | 1000 |
| PCTK1 | CDK16 | 100 | 1000 |
| PCTK3 | CDK18 | 100 | 1000 |
| PFTK1 | CDK14 | 100 | 1000 |
| PIK3C2G | PIK3C2G | 100 | 1000 |
| PIK3CA(E545A) | PIK3CA | 100 | 1000 |
| PIK3CA(H1047Y) | PIK3CA | 100 | 1000 |
| PIK3CA(M1043I) | PIK3CA | 100 | 1000 |
| PIK3CD | PIK3CD | 100 | 1000 |
| PIK4CB | PI4KB | 100 | 1000 |
| PIKFYVE | PIKFYVE | 100 | 1000 |
| PIM1 | PIM1 | 100 | 1000 |
| PIM2 | PIM2 | 100 | 1000 |
| PIM3 | PIM3 | 100 | 1000 |
| PIP5K2B | PIP4K2B | 100 | 1000 |
| PKAC-alpha | PRKACA | 100 | 1000 |
| PKAC-beta | PRKACB | 100 | 1000 |
| PKN1 | PKN1 | 100 | 1000 |
| PKNB (*M. tuberculosis*) | pknB | 100 | 1000 |
| PRKCD | PRKCD | 100 | 1000 |
| PRKCH | PRKCH | 100 | 1000 |
| PRKCQ | PRKCQ | 100 | 1000 |
| PRKG1 | PRKG1 | 100 | 1000 |
| PRKR | EIF2AK2 | 100 | 1000 |
| RAF1 | RAF1 | 100 | 1000 |
| RET(V804L) | RET | 100 | 1000 |
| RET(V804M) | RET | 100 | 1000 |

| | | | |
|---|---|---|---|
| RIOK1 | RIOK1 | 100 | 1000 |
| RIOK2 | RIOK2 | 100 | 1000 |
| RIPK1 | RIPK1 | 100 | 1000 |
| RIPK2 | RIPK2 | 100 | 1000 |
| ROS1 | ROS1 | 100 | 1000 |
| RPS6KA4(Kin.Dom.1-N-terminal) | RPS6KA4 | 100 | 1000 |
| RPS6KA5(Kin.Dom.1-N-terminal) | RPS6KA5 | 100 | 1000 |
| SBK1 | SBK1 | 100 | 1000 |
| SRMS | SRMS | 100 | 1000 |
| SRPK1 | SRPK1 | 100 | 1000 |
| SRPK3 | SRPK3 | 100 | 1000 |
| STK35 | STK35 | 100 | 1000 |
| TAK1 | MAP3K7 | 100 | 1000 |
| TAOK1 | TAOK1 | 100 | 1000 |
| TAOK2 | TAOK2 | 100 | 1000 |
| TAOK3 | TAOK3 | 100 | 1000 |
| TGFBR1 | TGFBR1 | 100 | 1000 |
| TGFBR2 | TGFBR2 | 100 | 1000 |
| TLK2 | TLK2 | 100 | 1000 |
| TNNI3K | TNNI3K | 100 | 1000 |
| TRKB | NTRK2 | 100 | 1000 |
| WEE1 | WEE1 | 100 | 1000 |
| ZAK | ZAK | 100 | 1000 |
| ZAP70 | ZAP70 | 100 | 1000 |

**Table S4**. Selectivity screening results for compound **9** at 1μM.

| DiscoveRx Gene Symbol | Entrez Gene Symbol | Percent Control | Compound Concentration (nM) |
|---|---|---|---|
| MEK5 | MAP2K5 | 0 | 1000 |
| MST4 | MST4 | 0 | 1000 |
| PFCDPK1 (*P. falciparum*) | CDPK1 | 0 | 1000 |
| SIK2 | SIK2 | 0 | 1000 |
| SIK | SIK1 | 0.3 | 1000 |
| DCAMKL3 | DCLK3 | 0.4 | 1000 |
| GAK | GAK | 0.7 | 1000 |
| LOK | STK10 | 1 | 1000 |
| MEK1 | MAP2K1 | 1.1 | 1000 |
| MAP4K5 | MAP4K5 | 1.4 | 1000 |
| STK36 | STK36 | 1.4 | 1000 |
| YANK2 | STK32B | 1.4 | 1000 |
| MEK2 | MAP2K2 | 1.6 | 1000 |
| YANK1 | STK32A | 2.2 | 1000 |
| SNARK | NUAK2 | 2.3 | 1000 |
| LTK | LTK | 2.5 | 1000 |
| QSK | KIAA0999 | 3.4 | 1000 |
| PAK2 | PAK2 | 3.8 | 1000 |
| MST3 | STK24 | 4 | 1000 |
| PAK1 | PAK1 | 4.3 | 1000 |
| CAMK1D | CAMK1D | 4.8 | 1000 |
| LCK | LCK | 4.9 | 1000 |
| MRCKB | CDC42BPB | 6.3 | 1000 |
| MEK4 | MAP2K4 | 7.1 | 1000 |
| EPHB6 | EPHB6 | 7.8 | 1000 |

| | | | |
|---|---|---|---|
| PRKD2 | PRKD2 | 8.4 | 1000 |
| PRKD3 | PRKD3 | 8.8 | 1000 |
| DMPK2 | CDC42BPG | 9.2 | 1000 |
| NLK | NLK | 10 | 1000 |
| MST2 | STK3 | 12 | 1000 |
| MAST1 | MAST1 | 14 | 1000 |
| SLK | SLK | 14 | 1000 |
| MELK | MELK | 15 | 1000 |
| AURKC | AURKC | 16 | 1000 |
| DDR1 | DDR1 | 16 | 1000 |
| FLT3(ITD,F691L) | FLT3 | 17 | 1000 |
| CSNK1E | CSNK1E | 20 | 1000 |
| MINK | MINK1 | 20 | 1000 |
| AKT3 | AKT3 | 21 | 1000 |
| ARK5 | NUAK1 | 22 | 1000 |
| PKN2 | PKN2 | 22 | 1000 |
| PRKD1 | PRKD1 | 22 | 1000 |
| YANK3 | STK32C | 23 | 1000 |
| ERK5 | MAPK7 | 25 | 1000 |
| MET(M1250T) | MET | 25 | 1000 |
| MAP4K3 | MAP4K3 | 26 | 1000 |
| MET | MET | 28 | 1000 |
| PIP5K2C | PIP4K2C | 28 | 1000 |
| AURKB | AURKB | 29 | 1000 |
| LIMK1 | LIMK1 | 29 | 1000 |
| AURKA | AURKA | 31 | 1000 |
| LATS2 | LATS2 | 34 | 1000 |

| | | | |
|---|---|---|---|
| EPHA8 | EPHA8 | 35 | 1000 |
| MARK3 | MARK3 | 36 | 1000 |
| STK33 | STK33 | 36 | 1000 |
| TNIK | TNIK | 37 | 1000 |
| WNK4 | WNK4 | 37 | 1000 |
| YSK4 | MAP3K19 | 37 | 1000 |
| MKNK2 | MKNK2 | 38 | 1000 |
| RIPK2 | RIPK2 | 38 | 1000 |
| HPK1 | MAP4K1 | 39 | 1000 |
| MST1R | MST1R | 39 | 1000 |
| YSK1 | STK25 | 39 | 1000 |
| EPHA1 | EPHA1 | 40 | 1000 |
| RSK1(Kin.Dom.2-C-terminal) | RPS6KA1 | 40 | 1000 |
| AMPK-alpha1 | PRKAA1 | 41 | 1000 |
| KIT(A829P) | KIT | 41 | 1000 |
| PDPK1 | PDPK1 | 41 | 1000 |
| DDR2 | DDR2 | 42 | 1000 |
| ERK2 | MAPK1 | 42 | 1000 |
| ERK1 | MAPK3 | 43 | 1000 |
| WNK1 | WNK1 | 43 | 1000 |
| AMPK-alpha2 | PRKAA2 | 44 | 1000 |
| CAMK1B | PNCK | 44 | 1000 |
| MAP4K2 | MAP4K2 | 45 | 1000 |
| MAP4K4 | MAP4K4 | 46 | 1000 |
| TRKA | NTRK1 | 46 | 1000 |
| ABL1(H396P)-nonphosphorylated | ABL1 | 47 | 1000 |

| | | | |
|---|---|---|---|
| IRAK1 | IRAK1 | 47 | 1000 |
| TIE2 | TEK | 47 | 1000 |
| PRKCI | PRKCI | 48 | 1000 |
| MAP3K3 | MAP3K3 | 49 | 1000 |
| JNK1 | MAPK8 | 50 | 1000 |
| PKNB (*M. tuberculosis*) | pknB | 50 | 1000 |
| RSK4(Kin.Dom.2-C-terminal) | RPS6KA6 | 50 | 1000 |
| JNK2 | MAPK9 | 51 | 1000 |
| MARK2 | MARK2 | 51 | 1000 |
| RSK2(Kin.Dom.2-C-terminal) | RPS6KA3 | 51 | 1000 |
| FRK | FRK | 52 | 1000 |
| ULK3 | ULK3 | 53 | 1000 |
| CDKL2 | CDKL2 | 57 | 1000 |
| TESK1 | TESK1 | 57 | 1000 |
| MEK6 | MAP2K6 | 58 | 1000 |
| SGK3 | SGK3 | 59 | 1000 |
| MRCKA | CDC42BPA | 60 | 1000 |
| NEK2 | NEK2 | 60 | 1000 |
| LATS1 | LATS1 | 61 | 1000 |
| MKK7 | MAP2K7 | 61 | 1000 |
| FLT3(ITD,D835V) | FLT3 | 62 | 1000 |
| MET(Y1235D) | MET | 63 | 1000 |
| RSK3(Kin.Dom.1-N-terminal) | RPS6KA2 | 63 | 1000 |
| S6K1 | RPS6KB1 | 63 | 1000 |
| PDGFRB | PDGFRB | 64 | 1000 |
| WNK2 | WNK2 | 64 | 1000 |

| | | | |
|---|---|---|---|
| ABL1(F317L)-phosphorylated | ABL1 | 65 | 1000 |
| CAMK2A | CAMK2A | 65 | 1000 |
| FLT3(K663Q) | FLT3 | 65 | 1000 |
| MYO3A | MYO3A | 65 | 1000 |
| PFTAIRE2 | CDK15 | 65 | 1000 |
| PIP5K1A | PIP5K1A | 65 | 1000 |
| RSK4(Kin.Dom.1-N-terminal) | RPS6KA6 | 65 | 1000 |
| CDK7 | CDK7 | 66 | 1000 |
| FLT3 (D835V) | FLT3 | 66 | 1000 |
| MAP3K4 | MAP3K4 | 66 | 1000 |
| NEK3 | NEK3 | 66 | 1000 |
| PAK6 | PAK6 | 66 | 1000 |
| RSK2 (Kin.Dom.1-N-terminal) | RPS6KA3 | 66 | 1000 |
| MAP3K2 | MAP3K2 | 67 | 1000 |
| MARK1 | MARK1 | 67 | 1000 |
| PFPK5 (*P. falciparum)* | MAL13P1.279 | 67 | 1000 |
| TAOK1 | TAOK1 | 67 | 1000 |
| BRAF(V600E) | BRAF | 68 | 1000 |
| MYO3B | MYO3B | 68 | 1000 |
| AKT1 | AKT1 | 69 | 1000 |
| FLT3(D835Y) | FLT3 | 69 | 1000 |
| MST1 | STK4 | 69 | 1000 |
| EPHA3 | EPHA3 | 70 | 1000 |
| ERBB3 | ERBB3 | 70 | 1000 |
| HASPIN | GSG2 | 70 | 1000 |
| ROCK2 | ROCK2 | 70 | 1000 |

| | | | |
|---|---|---|---|
| KIT(L576P) | KIT | 71 | 1000 |
| MYLK | MYLK | 71 | 1000 |
| CDC2L5 | CDK13 | 72 | 1000 |
| CSK | CSK | 72 | 1000 |
| PYK2 | PTK2B | 72 | 1000 |
| SGK | SGK1 | 72 | 1000 |
| SNRK | SNRK | 72 | 1000 |
| ABL1(Q252H)-nonphosphorylated | ABL1 | 73 | 1000 |
| ABL1(Y253F)-phosphorylated | ABL1 | 73 | 1000 |
| BMPR1B | BMPR1B | 73 | 1000 |
| DCAMKL2 | DCLK2 | 73 | 1000 |
| EIF2AK1 | EIF2AK1 | 73 | 1000 |
| PKN1 | PKN1 | 73 | 1000 |
| PRKCQ | PRKCQ | 73 | 1000 |
| TRKC | NTRK3 | 73 | 1000 |
| TYK2(JH1domain-catalytic) | TYK2 | 73 | 1000 |
| LZK | MAP3K13 | 74 | 1000 |
| MLK2 | MAP3K10 | 74 | 1000 |
| TSSK3 | TSSK3 | 74 | 1000 |
| LYN | LYN | 75 | 1000 |
| NEK10 | NEK10 | 75 | 1000 |
| RSK1(Kin.Dom.1-N-terminal) | RPS6KA1 | 75 | 1000 |
| ANKK1 | ANKK1 | 76 | 1000 |
| ERBB2 | ERBB2 | 76 | 1000 |
| IKK-epsilon | IKBKE | 76 | 1000 |

| | | | |
|---|---|---|---|
| JNK3 | MAPK10 | 76 | 1000 |
| PIK3CD | PIK3CD | 76 | 1000 |
| SGK2 | SGK2 | 76 | 1000 |
| CHEK2 | CHEK2 | 77 | 1000 |
| CSNK1A1L | CSNK1A1L | 77 | 1000 |
| FLT3(R834Q) | FLT3 | 77 | 1000 |
| ROCK1 | ROCK1 | 77 | 1000 |
| VPS34 | PIK3C3 | 77 | 1000 |
| CHEK1 | CHEK1 | 78 | 1000 |
| IKK-alpha | CHUK | 78 | 1000 |
| PIK3C2B | PIK3C2B | 78 | 1000 |
| IKK-beta | IKBKB | 79 | 1000 |
| IRAK3 | IRAK3 | 79 | 1000 |
| MTOR | MTOR | 79 | 1000 |
| ABL1(Q252H)-phosphorylated | ABL1 | 80 | 1000 |
| BLK | BLK | 80 | 1000 |
| PLK2 | PLK2 | 80 | 1000 |
| BRSK1 | BRSK1 | 81 | 1000 |
| BTK | BTK | 81 | 1000 |
| CDK4 | CDK4 | 81 | 1000 |
| GRK3 | ADRBK2 | 81 | 1000 |
| GRK7 | GRK7 | 81 | 1000 |
| JAK3(JH1domain-catalytic) | JAK3 | 81 | 1000 |
| PIK3CA | PIK3CA | 81 | 1000 |
| PLK4 | PLK4 | 81 | 1000 |
| PRKCH | PRKCH | 81 | 1000 |

| | | | |
|---|---|---|---|
| RSK3(Kin.Dom.2-C-terminal) | RPS6KA2 | 81 | 1000 |
| TNNI3K | TNNI3K | 81 | 1000 |
| ABL1(F317I)-phosphorylated | ABL1 | 82 | 1000 |
| ABL1(M351T)-phosphorylated | ABL1 | 82 | 1000 |
| ABL1-nonphosphorylated | ABL1 | 82 | 1000 |
| EGFR(T790M) | EGFR | 82 | 1000 |
| JAK2(JH1domain-catalytic) | JAK2 | 82 | 1000 |
| NDR1 | STK38 | 82 | 1000 |
| PAK4 | PAK4 | 82 | 1000 |
| PLK1 | PLK1 | 82 | 1000 |
| WNK3 | WNK3 | 82 | 1000 |
| ABL1-phosphorylated | ABL1 | 83 | 1000 |
| ERN1 | ERN1 | 83 | 1000 |
| NEK11 | NEK11 | 83 | 1000 |
| PIK3CG | PIK3CG | 83 | 1000 |
| PRKG2 | PRKG2 | 83 | 1000 |
| RIOK3 | RIOK3 | 83 | 1000 |
| ABL1(F317L)-nonphosphorylated | ABL1 | 84 | 1000 |
| DCAMKL1 | DCLK1 | 84 | 1000 |
| FYN | FYN | 84 | 1000 |
| PAK7 | PAK7 | 84 | 1000 |
| PIK3CA(Q546K) | PIK3CA | 84 | 1000 |
| RET(V804M) | RET | 84 | 1000 |
| WEE2 | WEE2 | 84 | 1000 |
| EPHA4 | EPHA4 | 85 | 1000 |

| | | | |
|---|---|---|---|
| FLT1 | FLT1 | 85 | 1000 |
| FLT3(D835H) | FLT3 | 85 | 1000 |
| LIMK2 | LIMK2 | 85 | 1000 |
| NIM1 | MGC42105 | 85 | 1000 |
| PIK3CA(E545K) | PIK3CA | 85 | 1000 |
| RIPK5 | DSTYK | 85 | 1000 |
| STK39 | STK39 | 85 | 1000 |
| TBK1 | TBK1 | 85 | 1000 |
| TYRO3 | TYRO3 | 85 | 1000 |
| VRK2 | VRK2 | 85 | 1000 |
| ABL1(T315I)-phosphorylated | ABL1 | 86 | 1000 |
| BMX | BMX | 86 | 1000 |
| CDKL5 | CDKL5 | 86 | 1000 |
| KIT(D816V) | KIT | 86 | 1000 |
| MYLK2 | MYLK2 | 86 | 1000 |
| SgK110 | SgK110 | 86 | 1000 |
| STK16 | STK16 | 86 | 1000 |
| ABL1(T315I)-nonphosphorylated | ABL1 | 87 | 1000 |
| AKT2 | AKT2 | 87 | 1000 |
| CTK | MATK | 87 | 1000 |
| EGFR(E746-A750del) | EGFR | 87 | 1000 |
| EPHA6 | EPHA6 | 87 | 1000 |
| FLT3(ITD) | FLT3 | 87 | 1000 |
| GCN2(Kin.Dom.2,S808G) | EIF2AK4 | 87 | 1000 |
| NEK9 | NEK9 | 87 | 1000 |
| PIP5K2B | PIP4K2B | 87 | 1000 |

| | | | |
|---|---|---|---|
| TAOK3 | TAOK3 | 87 | 1000 |
| ABL1(H396P)-phosphorylated | ABL1 | 88 | 1000 |
| BUB1 | BUB1 | 88 | 1000 |
| FES | FES | 88 | 1000 |
| MEK3 | MAP2K3 | 88 | 1000 |
| PIP5K1C | PIP5K1C | 88 | 1000 |
| TIE1 | TIE1 | 88 | 1000 |
| TRKB | NTRK2 | 88 | 1000 |
| ACVR1B | ACVR1B | 89 | 1000 |
| BRK | PTK6 | 89 | 1000 |
| CDK5 | CDK5 | 89 | 1000 |
| INSR | INSR | 89 | 1000 |
| RIPK4 | RIPK4 | 89 | 1000 |
| CLK1 | CLK1 | 90 | 1000 |
| EPHA5 | EPHA5 | 90 | 1000 |
| MAP3K1 | MAP3K1 | 90 | 1000 |
| SRC | SRC | 90 | 1000 |
| BMPR1A | BMPR1A | 91 | 1000 |
| CDK8 | CDK8 | 91 | 1000 |
| ERBB4 | ERBB4 | 91 | 1000 |
| GSK3A | GSK3A | 91 | 1000 |
| ICK | ICK | 91 | 1000 |
| KIT(V559D) | KIT | 91 | 1000 |
| LKB1 | STK11 | 91 | 1000 |
| PHKG1 | PHKG1 | 91 | 1000 |
| PIK3CA(E542K) | PIK3CA | 91 | 1000 |
| PIK3CA(I800L) | PIK3CA | 91 | 1000 |

| | | | |
|---|---|---|---|
| PRKX | PRKX | 91 | 1000 |
| CSNK1G2 | CSNK1G2 | 92 | 1000 |
| DLK | MAP3K12 | 92 | 1000 |
| EGFR(G719C) | EGFR | 92 | 1000 |
| EGFR(L858R,T790M) | EGFR | 92 | 1000 |
| EPHB2 | EPHB2 | 92 | 1000 |
| FLT3-autoinhibited | FLT3 | 92 | 1000 |
| KIT(D816H) | KIT | 92 | 1000 |
| PIK4CB | PI4KB | 92 | 1000 |
| PLK3 | PLK3 | 92 | 1000 |
| BRAF | BRAF | 93 | 1000 |
| LRRK2 | LRRK2 | 93 | 1000 |
| PDGFRA | PDGFRA | 93 | 1000 |
| RAF1 | RAF1 | 93 | 1000 |
| RET | RET | 93 | 1000 |
| RPS6KA4(Kin.Dom.2-C-terminal) | RPS6KA4 | 93 | 1000 |
| SYK | SYK | 93 | 1000 |
| AXL | AXL | 94 | 1000 |
| CDC2L2 | CDC2L2 | 94 | 1000 |
| CSNK2A2 | CSNK2A2 | 94 | 1000 |
| EGFR(L861Q) | EGFR | 94 | 1000 |
| IGF1R | IGF1R | 94 | 1000 |
| MLK1 | MAP3K9 | 94 | 1000 |
| NDR2 | STK38L | 94 | 1000 |
| PKAC-beta | PRKACB | 94 | 1000 |
| RPS6KA5(Kin.Dom.1-N-terminal) | RPS6KA5 | 94 | 1000 |

| | | | |
|---|---|---|---|
| SRPK2 | SRPK2 | 94 | 1000 |
| ALK | ALK | 95 | 1000 |
| CAMK1 | CAMK1 | 95 | 1000 |
| CDK9 | CDK9 | 95 | 1000 |
| FGR | FGR | 95 | 1000 |
| HCK | HCK | 95 | 1000 |
| IRAK4 | IRAK4 | 95 | 1000 |
| KIT | KIT | 95 | 1000 |
| PIK3CB | PIK3CB | 95 | 1000 |
| ABL1(E255K)-phosphorylated | ABL1 | 96 | 1000 |
| CDC2L1 | CDK11B | 96 | 1000 |
| CSF1R | CSF1R | 96 | 1000 |
| DYRK2 | DYRK2 | 96 | 1000 |
| EGFR(G719S) | EGFR | 96 | 1000 |
| PCTK2 | CDK17 | 96 | 1000 |
| RPS6KA5(Kin.Dom.2-C-terminal) | RPS6KA5 | 96 | 1000 |
| TNK2 | TNK2 | 96 | 1000 |
| ULK1 | ULK1 | 96 | 1000 |
| ULK2 | ULK2 | 96 | 1000 |
| CLK2 | CLK2 | 97 | 1000 |
| DAPK1 | DAPK1 | 97 | 1000 |
| DAPK2 | DAPK2 | 97 | 1000 |
| DYRK1A | DYRK1A | 97 | 1000 |
| FLT3(N841I) | FLT3 | 97 | 1000 |
| MLCK | MYLK3 | 97 | 1000 |
| p38-beta | MAPK11 | 97 | 1000 |

| | | | |
|---|---|---|---|
| ACVRL1 | ACVRL1 | 98 | 1000 |
| ERK4 | MAPK4 | 98 | 1000 |
| JAK1(JH1domain-catalytic) | JAK1 | 98 | 1000 |
| MAK | MAK | 98 | 1000 |
| MAPKAPK5 | MAPKAPK5 | 98 | 1000 |
| NEK6 | NEK6 | 98 | 1000 |
| PKAC-alpha | PRKACA | 98 | 1000 |
| PRKCE | PRKCE | 98 | 1000 |
| PRP4 | PRPF4B | 98 | 1000 |
| BIKE | BMP2K | 99 | 1000 |
| CAMK1G | CAMK1G | 99 | 1000 |
| CAMK4 | CAMK4 | 99 | 1000 |
| CLK3 | CLK3 | 99 | 1000 |
| EGFR(L858R) | EGFR | 99 | 1000 |
| EPHA2 | EPHA2 | 99 | 1000 |
| FER | FER | 99 | 1000 |
| HIPK2 | HIPK2 | 99 | 1000 |
| MAP3K15 | MAP3K15 | 99 | 1000 |
| MARK4 | MARK4 | 99 | 1000 |
| TGFBR2 | TGFBR2 | 99 | 1000 |
| TLK1 | TLK1 | 99 | 1000 |
| AAK1 | AAK1 | 100 | 1000 |
| ABL1(F317I)-nonphosphorylated | ABL1 | 100 | 1000 |
| ABL2 | ABL2 | 100 | 1000 |
| ACVR1 | ACVR1 | 100 | 1000 |
| ACVR2A | ACVR2A | 100 | 1000 |

| | | | |
|---|---|---|---|
| ACVR2B | ACVR2B | 100 | 1000 |
| ADCK3 | CABC1 | 100 | 1000 |
| ADCK4 | ADCK4 | 100 | 1000 |
| ALK(C1156Y) | ALK | 100 | 1000 |
| ALK(L1196M) | ALK | 100 | 1000 |
| ASK1 | MAP3K5 | 100 | 1000 |
| ASK2 | MAP3K6 | 100 | 1000 |
| BMPR2 | BMPR2 | 100 | 1000 |
| BRSK2 | BRSK2 | 100 | 1000 |
| CAMK2B | CAMK2B | 100 | 1000 |
| CAMK2D | CAMK2D | 100 | 1000 |
| CAMK2G | CAMK2G | 100 | 1000 |
| CAMKK1 | CAMKK1 | 100 | 1000 |
| CAMKK2 | CAMKK2 | 100 | 1000 |
| CASK | CASK | 100 | 1000 |
| CDK11 | CDK19 | 100 | 1000 |
| CDK2 | CDK2 | 100 | 1000 |
| CDK3 | CDK3 | 100 | 1000 |
| CDK4-cyclinD1 | CDK4 | 100 | 1000 |
| CDK4-cyclinD3 | CDK4 | 100 | 1000 |
| CDKL1 | CDKL1 | 100 | 1000 |
| CDKL3 | CDKL3 | 100 | 1000 |
| CIT | CIT | 100 | 1000 |
| CLK4 | CLK4 | 100 | 1000 |
| CSF1R-autoinhibited | CSF1R | 100 | 1000 |
| CSNK1A1 | CSNK1A1 | 100 | 1000 |
| CSNK1D | CSNK1D | 100 | 1000 |

| | | | |
|---|---|---|---|
| CSNK1G1 | CSNK1G1 | 100 | 1000 |
| CSNK1G3 | CSNK1G3 | 100 | 1000 |
| CSNK2A1 | CSNK2A1 | 100 | 1000 |
| DAPK3 | DAPK3 | 100 | 1000 |
| DMPK | DMPK | 100 | 1000 |
| DRAK1 | STK17A | 100 | 1000 |
| DRAK2 | STK17B | 100 | 1000 |
| DYRK1B | DYRK1B | 100 | 1000 |
| EGFR | EGFR | 100 | 1000 |
| EGFR(L747-E749del, A750P) | EGFR | 100 | 1000 |
| EGFR(L747-S752del, P753S) | EGFR | 100 | 1000 |
| EGFR(L747-T751del,Sins) | EGFR | 100 | 1000 |
| EGFR(S752-I759del) | EGFR | 100 | 1000 |
| EPHA7 | EPHA7 | 100 | 1000 |
| EPHB1 | EPHB1 | 100 | 1000 |
| EPHB3 | EPHB3 | 100 | 1000 |
| EPHB4 | EPHB4 | 100 | 1000 |
| ERK3 | MAPK6 | 100 | 1000 |
| ERK8 | MAPK15 | 100 | 1000 |
| FAK | PTK2 | 100 | 1000 |
| FGFR1 | FGFR1 | 100 | 1000 |
| FGFR2 | FGFR2 | 100 | 1000 |
| FGFR3 | FGFR3 | 100 | 1000 |
| FGFR3(G697C) | FGFR3 | 100 | 1000 |
| FGFR4 | FGFR4 | 100 | 1000 |
| FLT3 | FLT3 | 100 | 1000 |

| | | | |
|---|---|---|---|
| FLT4 | FLT4 | 100 | 1000 |
| GRK1 | GRK1 | 100 | 1000 |
| GRK2 | ADRBK1 | 100 | 1000 |
| GRK4 | GRK4 | 100 | 1000 |
| GSK3B | GSK3B | 100 | 1000 |
| HIPK1 | HIPK1 | 100 | 1000 |
| HIPK3 | HIPK3 | 100 | 1000 |
| HIPK4 | HIPK4 | 100 | 1000 |
| HUNK | HUNK | 100 | 1000 |
| INSRR | INSRR | 100 | 1000 |
| ITK | ITK | 100 | 1000 |
| JAK1(JH2domain-pseudokinase) | JAK1 | 100 | 1000 |
| KIT(V559D,T670I) | KIT | 100 | 1000 |
| KIT(V559D,V654A) | KIT | 100 | 1000 |
| KIT-autoinhibited | KIT | 100 | 1000 |
| LRRK2(G2019S) | LRRK2 | 100 | 1000 |
| MAPKAPK2 | MAPKAPK2 | 100 | 1000 |
| MERTK | MERTK | 100 | 1000 |
| MKNK1 | MKNK1 | 100 | 1000 |
| MLK3 | MAP3K11 | 100 | 1000 |
| MUSK | MUSK | 100 | 1000 |
| MYLK4 | MYLK4 | 100 | 1000 |
| NEK1 | NEK1 | 100 | 1000 |
| NEK4 | NEK4 | 100 | 1000 |
| NEK5 | NEK5 | 100 | 1000 |
| NEK7 | NEK7 | 100 | 1000 |
| NIK | MAP3K14 | 100 | 1000 |

| | | | |
|---|---|---|---|
| OSR1 | OXSR1 | 100 | 1000 |
| p38-alpha | MAPK14 | 100 | 1000 |
| p38-delta | MAPK13 | 100 | 1000 |
| p38-gamma | MAPK12 | 100 | 1000 |
| PAK3 | PAK3 | 100 | 1000 |
| PCTK1 | CDK16 | 100 | 1000 |
| PCTK3 | CDK18 | 100 | 1000 |
| PFTK1 | CDK14 | 100 | 1000 |
| PHKG2 | PHKG2 | 100 | 1000 |
| PIK3C2G | PIK3C2G | 100 | 1000 |
| PIK3CA(C420R) | PIK3CA | 100 | 1000 |
| PIK3CA(E545A) | PIK3CA | 100 | 1000 |
| PIK3CA(H1047L) | PIK3CA | 100 | 1000 |
| PIK3CA(H1047Y) | PIK3CA | 100 | 1000 |
| PIK3CA(M1043I) | PIK3CA | 100 | 1000 |
| PIKFYVE | PIKFYVE | 100 | 1000 |
| PIM1 | PIM1 | 100 | 1000 |
| PIM2 | PIM2 | 100 | 1000 |
| PIM3 | PIM3 | 100 | 1000 |
| PKMYT1 | PKMYT1 | 100 | 1000 |
| PRKCD | PRKCD | 100 | 1000 |
| PRKG1 | PRKG1 | 100 | 1000 |
| PRKR | EIF2AK2 | 100 | 1000 |
| RET(M918T) | RET | 100 | 1000 |
| RET(V804L) | RET | 100 | 1000 |
| RIOK1 | RIOK1 | 100 | 1000 |
| RIOK2 | RIOK2 | 100 | 1000 |

| | | | |
|---|---|---|---|
| RIPK1 | RIPK1 | 100 | 1000 |
| ROS1 | ROS1 | 100 | 1000 |
| RPS6KA4(Kin.Dom.1-N-terminal) | RPS6KA4 | 100 | 1000 |
| SBK1 | SBK1 | 100 | 1000 |
| SRMS | SRMS | 100 | 1000 |
| SRPK1 | SRPK1 | 100 | 1000 |
| SRPK3 | SRPK3 | 100 | 1000 |
| STK35 | STK35 | 100 | 1000 |
| TAK1 | MAP3K7 | 100 | 1000 |
| TAOK2 | TAOK2 | 100 | 1000 |
| TEC | TEC | 100 | 1000 |
| TGFBR1 | TGFBR1 | 100 | 1000 |
| TLK2 | TLK2 | 100 | 1000 |
| TNK1 | TNK1 | 100 | 1000 |
| TRPM6 | TRPM6 | 100 | 1000 |
| TSSK1B | TSSK1B | 100 | 1000 |
| TTK | TTK | 100 | 1000 |
| TXK | TXK | 100 | 1000 |
| TYK2(JH2domain-pseudokinase) | TYK2 | 100 | 1000 |
| VEGFR2 | KDR | 100 | 1000 |
| WEE1 | WEE1 | 100 | 1000 |
| YES | YES1 | 100 | 1000 |
| ZAK | ZAK | 100 | 1000 |
| ZAP70 | ZAP70 | 100 | 1000 |

**Table S5**. Selectivity screening results compound **10** (MRIA9) at 1µM.

| Kinase Name | Mutant/ Wild type | Residual activity | Compound Concentration (nM) |
|---|---|---|---|
| SIK2 | Wild type | 1 | 1000 |
| SIK3 | Wild type | 2 | 1000 |
| SIK1 | Wild type | 4 | 1000 |
| MAP4K5 | Wild type | 8 | 1000 |
| PAK3 | Wild type | 9 | 1000 |
| PAK2 | Wild type | 10 | 1000 |
| NLK | Wild type | 13 | 1000 |
| PKN3 | Wild type | 35 | 1000 |
| PAK1 | Wild type | 36 | 1000 |
| MKK4 | Wild type | 37 | 1000 |
| TIE2 | Wild type | 39 | 1000 |
| KIT D816V | Mutant | 41 | 1000 |
| MST4 | Wild type | 45 | 1000 |
| RAF1 YDYD | Const. active variant | 47 | 1000 |
| MELK | Wild type | 48 | 1000 |
| PRKD2 | Wild type | 51 | 1000 |
| RET S891A | Mutant | 52 | 1000 |
| PKCmu | Wild type | 53 | 1000 |
| EPHA1 | Wild type | 53 | 1000 |
| EPHA6 | Wild type | 54 | 1000 |
| STK25 | Wild type | 56 | 1000 |
| TYRO3 | Wild type | 56 | 1000 |
| ERK2 | Wild type | 58 | 1000 |
| TIE2 Y1108F | Mutant | 58 | 1000 |

| | | | |
|---|---|---|---|
| BMX | Wild type | 60 | 1000 |
| LCK | Wild type | 61 | 1000 |
| CAMK1D | Wild type | 62 | 1000 |
| RON | Wild type | 63 | 1000 |
| MAP3K10 | Wild type | 64 | 1000 |
| WNK2 | Wild type | 65 | 1000 |
| TIE2 R849W | Mutant | 65 | 1000 |
| MET D1228H | Mutant | 67 | 1000 |
| MKK6 SDTD | Const. active variant | 68 | 1000 |
| MEK1 F53L | Mutant | 68 | 1000 |
| KIT V560G | Mutant | 68 | 1000 |
| RIPK2 | Wild type | 69 | 1000 |
| RPS6KA4 | Wild type | 69 | 1000 |
| RET R749T | Mutant | 69 | 1000 |
| KIT D816H | Mutant | 70 | 1000 |
| KIT V559D/T670I | Mutant | 71 | 1000 |
| TRKA G667C | Mutant | 71 | 1000 |
| AuroraB | Wild type | 72 | 1000 |
| FGFR3 K650M | Mutant | 72 | 1000 |
| EGFR G719C | Mutant | 72 | 1000 |
| MINK1 | Wild type | 75 | 1000 |
| BRK | Wild type | 75 | 1000 |
| MET F1200I | Mutant | 75 | 1000 |
| KIT V559D | Mutant | 75 | 1000 |
| MYLK | Wild type | 76 | 1000 |
| RPS6KA5 | Wild type | 77 | 1000 |
| MET D1228N | Mutant | 77 | 1000 |

| | | | |
|---|---|---|---|
| MET Y1230D | Mutant | 77 | 1000 |
| KIT V654A | Mutant | 77 | 1000 |
| MEK1 | Wild type | 77 | 1000 |
| CSK | Wild type | 77 | 1000 |
| SLK | Wild type | 77 | 1000 |
| DDR2 | Wild type | 77 | 1000 |
| JNK2 | Wild type | 78 | 1000 |
| WNK1 | Wild type | 78 | 1000 |
| MEK2 | Wild type | 78 | 1000 |
| RET R813Q | Mutant | 78 | 1000 |
| EGFR d747-752/P753S | Mutant | 78 | 1000 |
| ALK L1196M | Mutant | 78 | 1000 |
| RET E762Q | Mutant | 78 | 1000 |
| MET Y1230H | Mutant | 79 | 1000 |
| RET V804L | Mutant | 79 | 1000 |
| TSF1 | Wild type | 79 | 1000 |
| p38beta | Wild type | 79 | 1000 |
| ACVR1B | Wild type | 79 | 1000 |
| MAP3K11 | Wild type | 79 | 1000 |
| RET Y791F | Mutant | 79 | 1000 |
| TGFBR2 | Wild type | 79 | 1000 |
| CAMKK1 | Wild type | 79 | 1000 |
| KIT | Wild type | 80 | 1000 |
| AMPKalpha1 | Wild type | 80 | 1000 |
| MET Y1230C | Mutant | 80 | 1000 |
| EGFR d747-749/A750P | Mutant | 81 | 1000 |
| VEGFR2 | Wild type | 81 | 1000 |

| | | | |
|---|---|---|---|
| PDGFRalpha D842V | Mutant | 81 | 1000 |
| ACK1 | Wild type | 82 | 1000 |
| MET Y1230A | Mutant | 82 | 1000 |
| MLK4 | Wild type | 82 | 1000 |
| FGFR2 | Wild type | 82 | 1000 |
| MAP3K9 | Wild type | 82 | 1000 |
| MAP4K4 | Wild type | 83 | 1000 |
| MET M1250T | Mutant | 83 | 1000 |
| PKCnu | Wild type | 83 | 1000 |
| PYK2 | Wild type | 83 | 1000 |
| EGFR L858R | Mutant | 84 | 1000 |
| EPHA3 | Wild type | 84 | 1000 |
| FGFR3 K650E | Mutant | 84 | 1000 |
| EGFR C797S/L858R | Mutant | 84 | 1000 |
| ACVR2B | Wild type | 84 | 1000 |
| EIF2AK2 | Wild type | 84 | 1000 |
| TAOK3 | Wild type | 84 | 1000 |
| PDGFRalpha V561D | Mutant | 84 | 1000 |
| RPS6KA6 | Wild type | 84 | 1000 |
| RET M918T | Mutant | 84 | 1000 |
| FLT3 ITD | Mutant | 84 | 1000 |
| MST1 | Wild type | 84 | 1000 |
| MYLK2 | Wild type | 84 | 1000 |
| ABL1 H396P | Mutant | 85 | 1000 |
| MET | Wild type | 85 | 1000 |
| KIT V559D/V654A | Mutant | 85 | 1000 |
| VEGFR3 | Wild type | 85 | 1000 |

| | | | |
|---|---|---|---|
| EPHA4 | Wild type | 85 | 1000 |
| PKCdelta | Wild type | 85 | 1000 |
| RET G691S | Mutant | 85 | 1000 |
| AKT2 | Wild type | 85 | 1000 |
| TIE2 Y897S | Mutant | 85 | 1000 |
| EGFR G719S | Mutant | 86 | 1000 |
| FGFR1 V561M | Mutant | 86 | 1000 |
| FGFR1 | Wild type | 86 | 1000 |
| PAK7 | Wild type | 86 | 1000 |
| RET V804E | Mutant | 86 | 1000 |
| MET Y1235D | Mutant | 86 | 1000 |
| EPHA5 | Wild type | 87 | 1000 |
| PKCbeta2 | Wild type | 87 | 1000 |
| COT | Wild type | 87 | 1000 |
| NPM1 ALK F1174L | Mutant | 87 | 1000 |
| CDK3/CycC | Wild type | 87 | 1000 |
| RET V804M | Mutant | 87 | 1000 |
| EGFR C797S | Mutant | 87 | 1000 |
| MAP3K7/MAP3K7IP1 | Wild type | 87 | 1000 |
| TAOK2 | Wild type | 87 | 1000 |
| AuroraA | Wild type | 88 | 1000 |
| PIM1 | Wild type | 88 | 1000 |
| RET G810C | Mutant | 88 | 1000 |
| FYN Y531F | Mutant | 88 | 1000 |
| CDK4/CycD3 | Wild type | 88 | 1000 |
| RPS6KA2 | Wild type | 88 | 1000 |
| KIT T670I | Mutant | 88 | 1000 |

| | | | |
|---|---|---|---|
| ERK1 | Wild type | 88 | 1000 |
| EGFR L861Q | Mutant | 88 | 1000 |
| RPS6KA1 | Wild type | 89 | 1000 |
| STK33 | Wild type | 89 | 1000 |
| BLK | Wild type | 89 | 1000 |
| BMPR1B | Wild type | 89 | 1000 |
| CHK2 | Wild type | 89 | 1000 |
| PI4K2A | Wild type | 89 | 1000 |
| SGK1 | Wild type | 89 | 1000 |
| NEK9 | Wild type | 89 | 1000 |
| p38gamma | Wild type | 89 | 1000 |
| CDK19/CycC | Wild type | 89 | 1000 |
| EPHB2 | Wild type | 89 | 1000 |
| CDK2/CycA2 | Wild type | 89 | 1000 |
| BRAF | Wild type | 90 | 1000 |
| p38alpha | Wild type | 90 | 1000 |
| MARK1 | Wild type | 90 | 1000 |
| NEK2 | Wild type | 90 | 1000 |
| ALK R1275Q | Mutant | 90 | 1000 |
| ERBB2 775YVMA776 | Mutant | 90 | 1000 |
| RET Y806H | Mutant | 90 | 1000 |
| TGFBR1 | Wild type | 91 | 1000 |
| FGFR3 G697C | Mutant | 91 | 1000 |
| IRAK1 | Wild type | 91 | 1000 |
| ALK F1174S | Mutant | 91 | 1000 |
| RET G810S | Mutant | 91 | 1000 |
| EIF2AK3 | Wild type | 91 | 1000 |

| | | | |
|---|---|---|---|
| CAMKK2 | Wild type | 91 | 1000 |
| PLK1 | Wild type | 91 | 1000 |
| PKA | Wild type | 91 | 1000 |
| DYRK2 | Wild type | 91 | 1000 |
| ROS | Wild type | 91 | 1000 |
| IRAK4 | Wild type | 92 | 1000 |
| EGFR d752-759 | Mutant | 92 | 1000 |
| PKMzeta | Wild type | 92 | 1000 |
| KIT A829P | Mutant | 92 | 1000 |
| MUSK | Wild type | 92 | 1000 |
| RPS6KA3 | Wild type | 92 | 1000 |
| CDK1/CycA2 | Wild type | 92 | 1000 |
| PKCepsilon | Wild type | 92 | 1000 |
| PHKG1 | Wild type | 92 | 1000 |
| ASK1 | Wild type | 93 | 1000 |
| PKCgamma | Wild type | 93 | 1000 |
| S6Kbeta | Wild type | 93 | 1000 |
| STK23 | Wild type | 93 | 1000 |
| ARK5 | Wild type | 93 | 1000 |
| RET | Wild type | 93 | 1000 |
| FLT3 D835Y | Mutant | 93 | 1000 |
| MTOR | Wild type | 93 | 1000 |
| STK39 | Wild type | 93 | 1000 |
| CDK1/CycE1 | Wild type | 93 | 1000 |
| MAP4K2 | Wild type | 93 | 1000 |
| MYLK3 | Wild type | 93 | 1000 |
| PDGFRbeta | Wild type | 93 | 1000 |

| | | | |
|---|---|---|---|
| ABL1 T315I | Mutant | 94 | 1000 |
| MET L1195V | Mutant | 94 | 1000 |
| PAK4 | Wild type | 94 | 1000 |
| SNARK | Wild type | 94 | 1000 |
| DDR2 N456S | Mutant | 94 | 1000 |
| JAK1 S729C | Mutant | 94 | 1000 |
| ABL1 F317I | Mutant | 94 | 1000 |
| JNK1 | Wild type | 94 | 1000 |
| CCDC6 RET | Mutant | 94 | 1000 |
| CAMK4 | Wild type | 94 | 1000 |
| CK1epsilon | Wild type | 94 | 1000 |
| NPM1 ALK | Mutant | 94 | 1000 |
| PKMYT1 | Wild type | 94 | 1000 |
| CSF1R | Wild type | 94 | 1000 |
| AuroraC | Wild type | 95 | 1000 |
| TRKA | Wild type | 95 | 1000 |
| MKNK1 | Wild type | 95 | 1000 |
| CDK1/CycB1 | Wild type | 95 | 1000 |
| DYRK3 | Wild type | 95 | 1000 |
| MST2 | Wild type | 95 | 1000 |
| TNK1 | Wild type | 95 | 1000 |
| SRPK1 | Wild type | 95 | 1000 |
| p38delta | Wild type | 95 | 1000 |
| RIPK5 | Wild type | 95 | 1000 |
| ROCK2 | Wild type | 95 | 1000 |
| ZAK | Wild type | 96 | 1000 |
| EPHB3 | Wild type | 96 | 1000 |

| | | | |
|---|---|---|---|
| CDK4/CycD1 | Wild type | 96 | 1000 |
| JAK1 | Wild type | 96 | 1000 |
| AXL | Wild type | 96 | 1000 |
| ACVRL1 | Wild type | 96 | 1000 |
| IKKepsilon | Wild type | 96 | 1000 |
| EEF2K | Wild type | 96 | 1000 |
| PBK | Wild type | 96 | 1000 |
| CDK2/CycD1 | Wild type | 96 | 1000 |
| CDK6/CycD2 | Wild type | 96 | 1000 |
| GSK3beta | Wild type | 96 | 1000 |
| EGFR T790M/L858R | Mutant | 96 | 1000 |
| PDGFRalpha T674I | Mutant | 96 | 1000 |
| PAK6 | Wild type | 96 | 1000 |
| PIP5K1C | Wild type | 96 | 1000 |
| FYN | Wild type | 97 | 1000 |
| CDK9/CycK | Wild type | 97 | 1000 |
| EML4 ALK F1174L | Mutant | 97 | 1000 |
| CK1alpha1 | Wild type | 97 | 1000 |
| HIPK4 | Wild type | 97 | 1000 |
| CDK3/CycE1 | Wild type | 97 | 1000 |
| IKKbeta | Wild type | 97 | 1000 |
| ALK | Wild type | 97 | 1000 |
| IKKalpha | Wild type | 97 | 1000 |
| ERK7 | Wild type | 97 | 1000 |
| VRK1 | Wild type | 97 | 1000 |
| ITK | Wild type | 97 | 1000 |
| ABL1 Q252H | Mutant | 97 | 1000 |

| | | | |
|---|---|---|---|
| NEK1 | Wild type | 98 | 1000 |
| DNAPK | Wild type | 98 | 1000 |
| PKCiota | Wild type | 98 | 1000 |
| ZAP70 | Wild type | 98 | 1000 |
| EGFR T790M/C797S/L858R | Mutant | 98 | 1000 |
| S6K | Wild type | 98 | 1000 |
| MERTK | Wild type | 98 | 1000 |
| PI4KB | Wild type | 98 | 1000 |
| GSG2 | Wild type | 98 | 1000 |
| EPHA2 | Wild type | 98 | 1000 |
| TRKC | Wild type | 98 | 1000 |
| SYK | Wild type | 98 | 1000 |
| FER | Wild type | 98 | 1000 |
| CDK8/CycC | Wild type | 98 | 1000 |
| ERBB2 | Wild type | 98 | 1000 |
| INSR | Wild type | 98 | 1000 |
| DYRK1B | Wild type | 98 | 1000 |
| MATK | Wild type | 98 | 1000 |
| ULK2 | Wild type | 98 | 1000 |
| WNK3 | Wild type | 99 | 1000 |
| LRRK2 I2020T | Mutant | 99 | 1000 |
| LRRK2 G2019S | Mutant | 99 | 1000 |
| ABL1 M351T | Mutant | 99 | 1000 |
| SGK3 | Wild type | 99 | 1000 |
| TEC | Wild type | 99 | 1000 |
| ABL1 G250E | Mutant | 99 | 1000 |
| CDK5/p25NCK | Wild type | 99 | 1000 |

| | | | |
|---|---|---|---|
| CDK20/CycT1 | Wild type | 99 | 1000 |
| EML4 ALK | Mutant | 99 | 1000 |
| AKT3 | Wild type | 99 | 1000 |
| EPHB1 | Wild type | 99 | 1000 |
| ABL2 | Wild type | 99 | 1000 |
| MEK5 | Wild type | 99 | 1000 |
| PIK3CA/PIK3R1 | Wild type | 99 | 1000 |
| ABL1 E255K | Mutant | 99 | 1000 |
| TYK2 | Wild type | 99 | 1000 |
| HIPK1 | Wild type | 99 | 1000 |
| LYN | Wild type | 100 | 1000 |
| PRKG2 | Wild type | 100 | 1000 |
| ALK F1174L | Mutant | 100 | 1000 |
| PRKG1 | Wild type | 100 | 1000 |
| MAPKAPK3 | Wild type | 100 | 1000 |
| ABL1 Y253F | Mutant | 100 | 1000 |
| NEK4 | Wild type | 100 | 1000 |
| ALK C1156Y | Mutant | 100 | 1000 |
| CDK7/CycH/MAT1 | Wild type | 100 | 1000 |
| MKNK2 | Wild type | 100 | 1000 |
| CDK16/CycY | Wild type | 100 | 1000 |
| ERBB4 | Wild type | 100 | 1000 |
| CDK5/p35NCK | Wild type | 100 | 1000 |
| FGFR3 | Wild type | 100 | 1000 |
| PIP5K1B | Wild type | 100 | 1000 |
| PIP5K1A | Wild type | 100 | 1000 |
| ACVR2A | Wild type | 101 | 1000 |

| | | | |
|---|---|---|---|
| CAMK2D | Wild type | 101 | 1000 |
| PIK3CG | Wild type | 101 | 1000 |
| TBK1 | Wild type | 101 | 1000 |
| PIK3C2A | Wild type | 101 | 1000 |
| BRSK1 | Wild type | 101 | 1000 |
| CDK6/CycD3 | Wild type | 101 | 1000 |
| EGFR d746-750/T790M/C797S | Mutant | 101 | 1000 |
| CDK9/CycT1 | Wild type | 101 | 1000 |
| EPHA8 | Wild type | 101 | 1000 |
| CDK20/CycH | Wild type | 101 | 1000 |
| HIPK3 | Wild type | 101 | 1000 |
| TSK2 | Wild type | 101 | 1000 |
| PIK3CB/PIK3R1 | Wild type | 102 | 1000 |
| ACVR1 | Wild type | 102 | 1000 |
| SRPK2 | Wild type | 102 | 1000 |
| AKT1 | Wild type | 102 | 1000 |
| BMPR1A | Wild type | 102 | 1000 |
| PKCzeta | Wild type | 102 | 1000 |
| ABL1 | Wild type | 102 | 1000 |
| EGFR T790M | Mutant | 102 | 1000 |
| TTK | Wild type | 102 | 1000 |
| MET G1163R | Mutant | 102 | 1000 |
| CHK1 | Wild type | 103 | 1000 |
| INSRR | Wild type | 103 | 1000 |
| CDC7/DBF4 | Wild type | 103 | 1000 |
| GRK7 | Wild type | 103 | 1000 |
| MARK2 | Wild type | 103 | 1000 |

| | | | |
|---|---|---|---|
| PDK1 | Wild type | 103 | 1000 |
| PIK3C3 | Wild type | 103 | 1000 |
| EPHA7 | Wild type | 103 | 1000 |
| FES | Wild type | 103 | 1000 |
| ROCK1 | Wild type | 103 | 1000 |
| CAMK2A | Wild type | 103 | 1000 |
| CDK13/CycK | Wild type | 103 | 1000 |
| MST3 | Wild type | 103 | 1000 |
| CAMK2G | Wild type | 103 | 1000 |
| PIK3CD/PIK3R1 | Wild type | 104 | 1000 |
| FRK | Wild type | 104 | 1000 |
| CDC42BPA | Wild type | 104 | 1000 |
| SNK | Wild type | 104 | 1000 |
| CDK12/CycK | Wild type | 104 | 1000 |
| NEK11 | Wild type | 104 | 1000 |
| MAP3K1 | Wild type | 104 | 1000 |
| PRK2 | Wild type | 104 | 1000 |
| LIMK2 | Wild type | 104 | 1000 |
| DMPK | Wild type | 104 | 1000 |
| JNK3 | Wild type | 104 | 1000 |
| DYRK1A | Wild type | 104 | 1000 |
| CDK6/CycD1 | Wild type | 104 | 1000 |
| CDK18/CycY | Wild type | 105 | 1000 |
| LTK | Wild type | 105 | 1000 |
| TTBK2 | Wild type | 105 | 1000 |
| PIM2 | Wild type | 105 | 1000 |
| EGFR d746-750 | Mutant | 105 | 1000 |

| | | | |
|---|---|---|---|
| HIPK2 | Wild type | 105 | 1000 |
| EPHB4 | Wild type | 105 | 1000 |
| EGFR | Wild type | 105 | 1000 |
| EGFR d746-750/C797S | Mutant | 105 | 1000 |
| DYRK4 | Wild type | 105 | 1000 |
| CK1delta | Wild type | 105 | 1000 |
| CDK17/p35NCK | Wild type | 105 | 1000 |
| PDGFRalpha | Wild type | 105 | 1000 |
| FGR | Wild type | 106 | 1000 |
| PI4K2B | Wild type | 106 | 1000 |
| CDK12 R722C/CycK | Mutant | 106 | 1000 |
| PKCbeta1 | Wild type | 106 | 1000 |
| VEGFR1 | Wild type | 106 | 1000 |
| WEE1 | Wild type | 106 | 1000 |
| PASK | Wild type | 106 | 1000 |
| PIM3 | Wild type | 106 | 1000 |
| LRRK2 | Wild type | 106 | 1000 |
| JAK2 | Wild type | 106 | 1000 |
| MARK4 | Wild type | 107 | 1000 |
| SGK2 | Wild type | 107 | 1000 |
| DDR2 T654M | Mutant | 107 | 1000 |
| GRK3 | Wild type | 107 | 1000 |
| TLK1 | Wild type | 107 | 1000 |
| PRK1 | Wild type | 107 | 1000 |
| TRKB | Wild type | 107 | 1000 |
| MEKK3 | Wild type | 107 | 1000 |
| CDC42BPB | Wild type | 107 | 1000 |

| | | | |
|---|---|---|---|
| NEK6 | Wild type | 107 | 1000 |
| TXK | Wild type | 107 | 1000 |
| CDK2/CycE1 | Wild type | 107 | 1000 |
| PKCalpha | Wild type | 107 | 1000 |
| BRAF V600E | Mutant | 107 | 1000 |
| RET G810R | Mutant | 108 | 1000 |
| SRC | Wild type | 108 | 1000 |
| YES | Wild type | 108 | 1000 |
| NEK3 | Wild type | 108 | 1000 |
| PHKG2 | Wild type | 108 | 1000 |
| JAK3 | Wild type | 108 | 1000 |
| FAK | Wild type | 108 | 1000 |
| GSK3alpha | Wild type | 108 | 1000 |
| ALK G1202R | Mutant | 109 | 1000 |
| HRI | Wild type | 109 | 1000 |
| CDK4/CycD2 | Wild type | 109 | 1000 |
| FGFR4 | Wild type | 109 | 1000 |
| BTK | Wild type | 109 | 1000 |
| IGF1R | Wild type | 109 | 1000 |
| EGFR d746-750/T790M/C797S/L858R | Mutant | 109 | 1000 |
| EGFR L718Q | Mutant | 109 | 1000 |
| CK2alpha2 | Wild type | 110 | 1000 |
| CLK2 | Wild type | 110 | 1000 |
| PIK3C2G | Wild type | 110 | 1000 |
| PKCtheta | Wild type | 110 | 1000 |
| GRK2 | Wild type | 111 | 1000 |

| | | | |
|---|---|---|---|
| MKK7 | Wild type | 111 | 1000 |
| BRSK2 | Wild type | 111 | 1000 |
| STK17A | Wild type | 111 | 1000 |
| GRK6 | Wild type | 111 | 1000 |
| ERK5 | Wild type | 111 | 1000 |
| HCK | Wild type | 111 | 1000 |
| MAPKAPK5 | Wild type | 112 | 1000 |
| DAPK2 | Wild type | 112 | 1000 |
| MARK3 | Wild type | 112 | 1000 |
| DCAMKL2 | Wild type | 112 | 1000 |
| CK1gamma3 | Wild type | 113 | 1000 |
| LIMK1 | Wild type | 113 | 1000 |
| FLT3 | Wild type | 113 | 1000 |
| SRMS | Wild type | 114 | 1000 |
| MASTL | Wild type | 114 | 1000 |
| PIK3C2B | Wild type | 115 | 1000 |
| PLK3 | Wild type | 116 | 1000 |
| CK2alpha1 | Wild type | 116 | 1000 |
| DAPK3 | Wild type | 116 | 1000 |
| VRK2 | Wild type | 117 | 1000 |
| SAK | Wild type | 117 | 1000 |
| PRKX | Wild type | 117 | 1000 |
| CAMK2B | Wild type | 117 | 1000 |
| CLK1 | Wild type | 118 | 1000 |
| PKCeta | Wild type | 119 | 1000 |
| CLK4 | Wild type | 119 | 1000 |
| RIPK4 | Wild type | 120 | 1000 |

| | | | |
|---|---|---|---|
| MEKK2 | Wild type | 120 | 1000 |
| DAPK1 | Wild type | 121 | 1000 |
| GRK5 | Wild type | 122 | 1000 |
| CK1gamma2 | Wild type | 123 | 1000 |
| BUB1B | Wild type | 123 | 1000 |
| TLK2 | Wild type | 123 | 1000 |
| MAPKAPK2 | Wild type | 124 | 1000 |
| LRRK2 R1441C | Mutant | 125 | 1000 |
| CK1gamma1 | Wild type | 127 | 1000 |
| GRK4 | Wild type | 127 | 1000 |
| TSSK1 | Wild type | 127 | 1000 |
| NEK7 | Wild type | 131 | 1000 |
| TTBK1 | Wild type | 131 | 1000 |
| CLK3 | Wild type | 132 | 1000 |
| NIK | Wild type | 140 | 1000 |

**Table S6**. NanoBRET™ assay information.

| Protein Kinase | Alias | Catalog #/ CAS # | NanoLuc orientation | Tracer K10 $IC_{50}$, [nM] | Tracer K10, used [nM] |
|---|---|---|---|---|---|
| SIK1 | | NV2031 | N | 120 | 150 |
| SNF1LK2 | SIK2 | NV2061 | N | 30 | 50 |
| SIK3 | | NV2041 | N | 250 | 250 |
| STK24 | MST3 | CS1810C243 | C | 260 | 300 |
| STK26 | MST4 | CS1810C245 | C | 270 | 300 |
| MAP4K5 | | CS1810C270 | C | 2000 | 1000 |
| NLK | | CS1810C222 | C | 750 | 500 |
| PKN3 | | CS181023 | C | 260 | 300 |
| TIE2 | | NV2151 | C | 145 | 150 |

**Table S7**. Data collection and refinement statistics.

| **Complex** | MST3- **6** (G-5555) | MST3-**10** (MRIA9) | MST3- **7** (MRIA7) | MST3-**16** (MRIA11) | MST3-**9** (MRIA12) | MST3-**14** (MRIA13) | MST4- **6** (G-5555) |
|---|---|---|---|---|---|---|---|
| PDB code | 7B30 | 7B31 | 7B32 | 7B33 | 7B34 | 7B35 | 7B36 |
| ***Data Collection*** | | | | | | | |
| Resolution range (Å)[a] | 39.64 - 2.10 (2.16 - 2.10) | 39.86 - 1.80 (1.84 - 1.80) | 49.90 - 1.75 (1.78 - 1.75) | 49.41 - 1.90 (1.94 - 1.90) | 39.73 - 2.10 (2.16 - 2.10) | 47.89 - 2.40 (2.49 - 2.40) | 46-47 - 2.11 (2.17 - 2.11) |
| Space group | *C*2 | *C*2 | *C*2 | *C*2 | *C*2 | $P2_1$ | $P2_1$ |
| No. molecules/asymmetric unit | 1 | 1 | 1 | 1 | 1 | 2 | 2 |
| Cell dimensions a, b, c (Å) | 99.00, 58.85, 61.84 | 99.43, 59.07, 61.73 | 100.04, 59.21, 61.55 | 98.97, 58.95, 61.44 | 99.26, 59.07, 61.79 | 55.88, 93.21, 61.30 | 63.75, 53.19, 93.52 |
| α, β, γ (°C) | 89.96, 92.92, 89.98 | 90.00, 93.52, 90.00 | 90.00, 93.95, 90.00 | 90.00, 93.13, 90.00 | 90.00, 93.05, 90.00 | 90.00, 92.19, 90.00 | 90.00, 96.36, 90.00 |
| No. unique observations [a] | 20236 (1341) | 33013 (1926) | 36097 (1989) | 27657 (1776) | 20791 (1732) | 24436 (2563) | 35855 (2756) |
| Completeness (%)[a] | 97.0 (79.3) | 99.4 (99.6) | 99.5 (99.9) | 98.9 (99.5) | 99.1 (99.9) | 99.2 (99.1) | 98.7 (94.0) |
| Mean (($I$)/σ($I$)) [a] | 12.6 (3.6) | 16.7 (2.2) | 18.4 (3.6) | 10.5 (2.6) | 12.1 (2.1) | 9.2 (2.3) | 13.3 (2.1) |
| $R_{pim}$ [a] | 0.030 (0.190) | 0.024 (0.352) | 0.024 (0.209) | 0.045 (0.307) | 0.042 (0.477) | 0.059 (0.353) | 0.029 (0.348) |
| CC(1/2) [a] | 0.997 (0.881) | 0.999 (0.742) | 0.999 (0.905) | 0.997 (0.813) | 0.997 (0.518) | 0.995 (0.823) | 0.999 (0.886) |
| Multiplicity [a] | 6.0 (4.7) | 6.1 (6.3) | 6.9 (7.1) | 5.0 (5.3) | 5.4 (5.6) | 5.6 (5.4) | 5.0 (4.5) |
| ***Refinement*** | | | | | | | |
| No. atoms in refinement (P/L/O) | 2035 / 35 / 108 | 2109 / 35 / 116 | 2070 / 42 / 125 | 2069 / 37 / 119 | 2056 / 36 / 102 | 4009 / 37 / 49 | 4157 / 35 / 96 |
| B factor ($Å^2$) (P/L/O) | 46.0 / 39.2 / 45.4 | 42.1 / 27.3 / 37.1 | 32.2 / 20.9 / 29.4 | 43.1 / 30.0 / 40.8 | 51.5 / 34.0 / 50.4 | 49.9 / 37.3 / 40.8 | 54.4 / 43.5 / 48.5 |
| $R_{work}$ (%) | 17.7 | 17.7 | 17.4 | 18.1 | 18.9 | 20.5 | 20.1 |
| $R_{free}$ (%) | 19.6 | 21.2 | 20.3 | 20.8 | 21.5 | 26.3 | 24.4 |
| r.m.s. deviation bond length (Å) | 0.007 | 0.006 | 0.006 | 0.007 | 0.008 | 0.004 | 0.008 |
| r.m.s deviation bond angle (°) | 0.99 | 0.82 | 0.79 | 0.85 | 0.88 | 0.67 | 0.85 |
| ***Molprobity Ramachandran*** | | | | | | | |
| Favoured (%) | 96 | 96 | 96 | 97 | 95 | 96 | 96 |
| Outliers (%) | 0 | 0 | 0 | 0 | 0 | 0 | 0 |
| Crystallization conditions | 10% PEG 6000, 0.1M HEPES pH 7.0 | 16% PEG 6000, 0.1M HEPES pH 7.0 | 20% PEG 3350, 0.1M Bis-tris pH 6.5 | 19% PEG 3350, 0.1M Bis-tris pH 6.8 | 14% PEG 6000, 0.1M HEPES pH 7.2 | 24% PEG 3350, 0.1M citrate pH 5.6 | 28% PEG 6000, 0.1M HEPES pH 7.5 |

[a] Numbers in parenthesis refer to the highest-resolution shell.

**Analytical data for compounds 7-38.**

$^1$H NMR compound **21**:

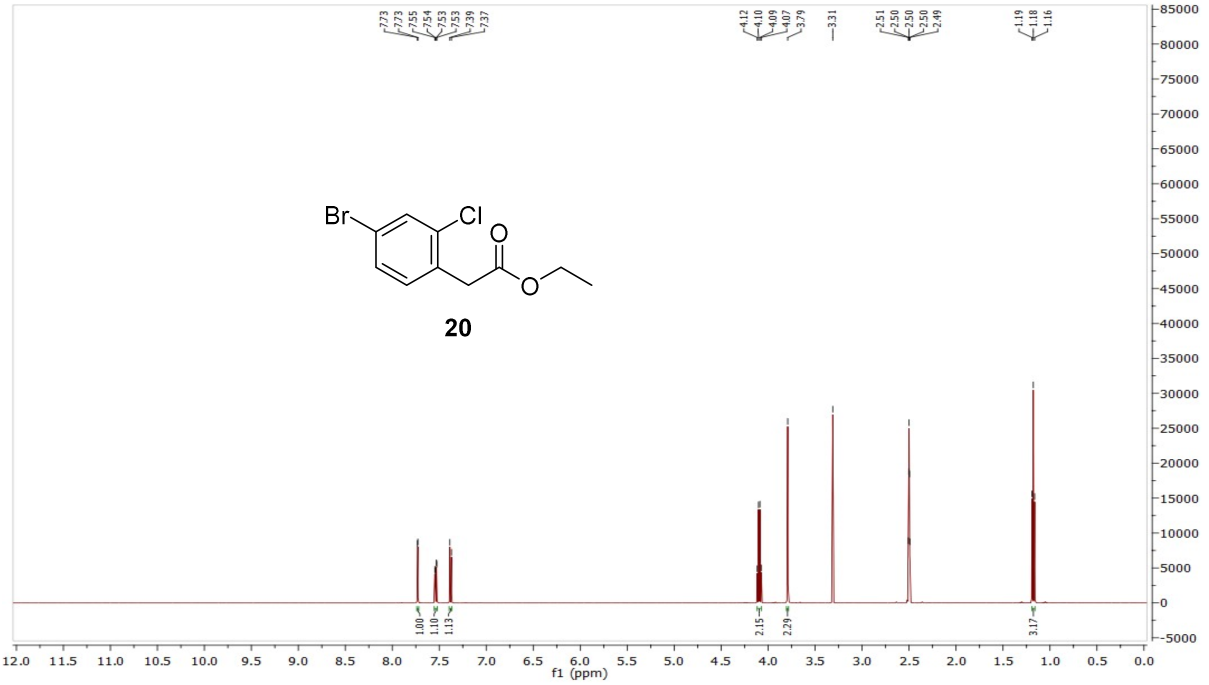

$^{13}$C NMR compound **21**:

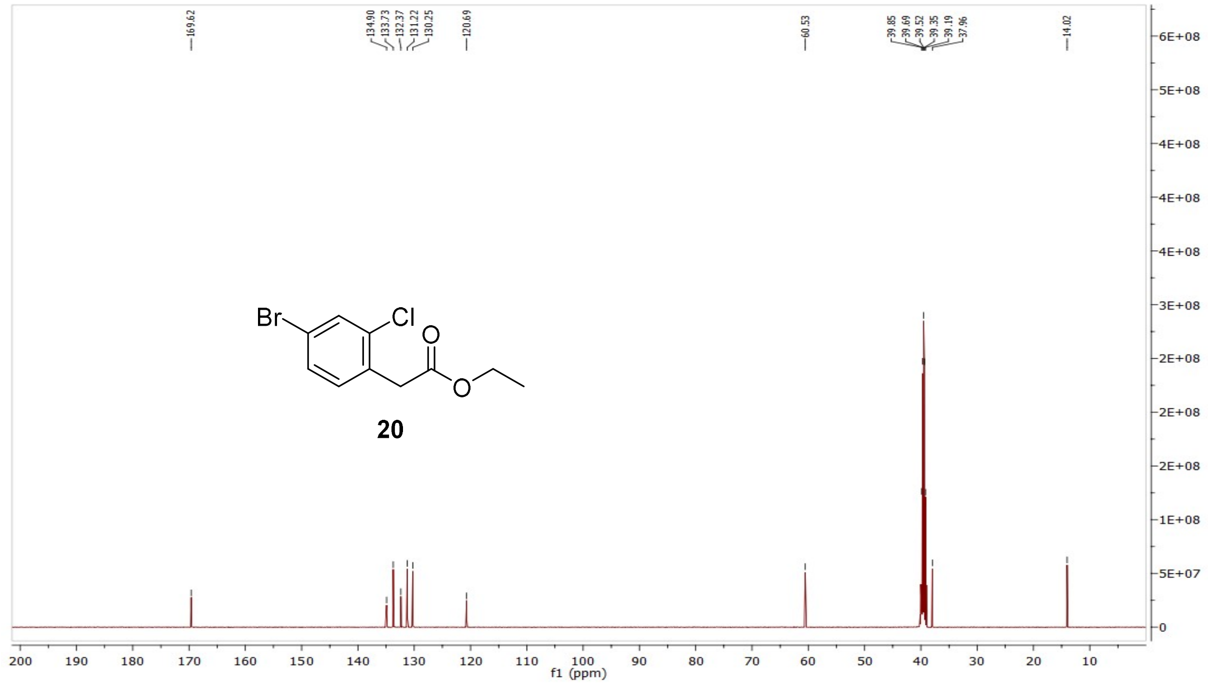

MS (ESI+) compound **21**:

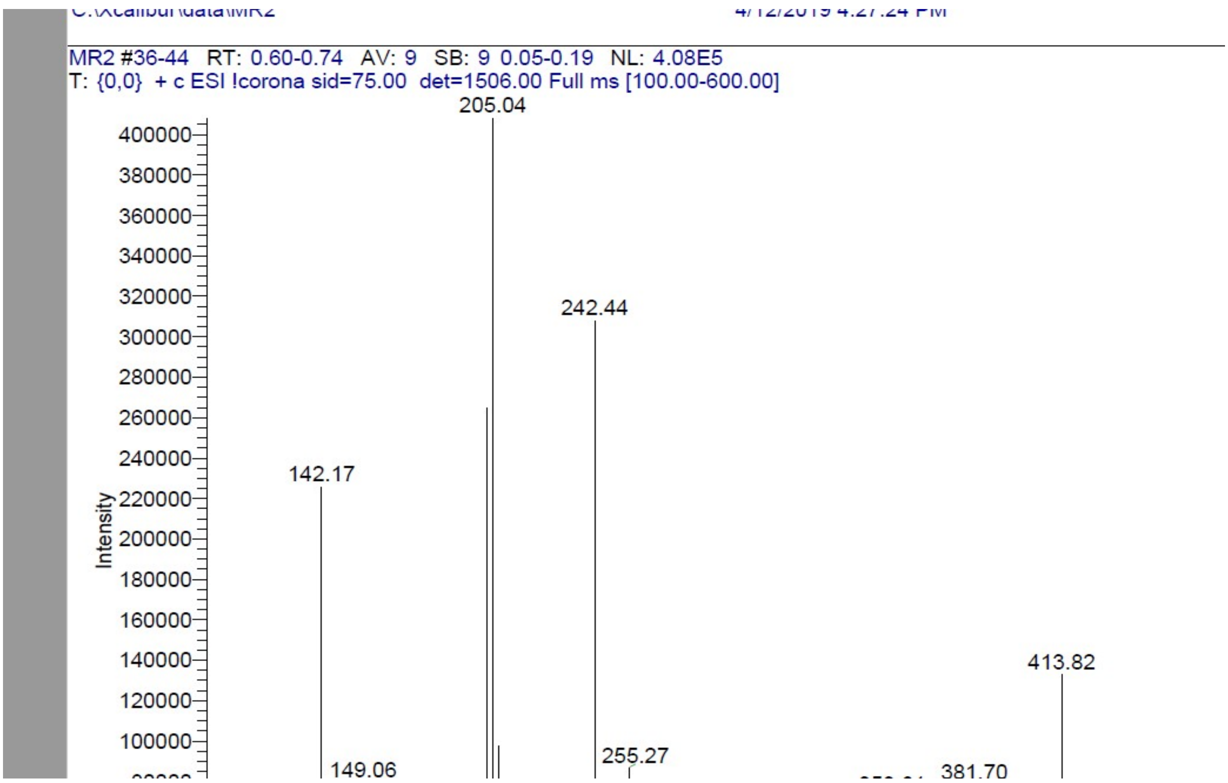


$^{1}$H NMR compound **22**:

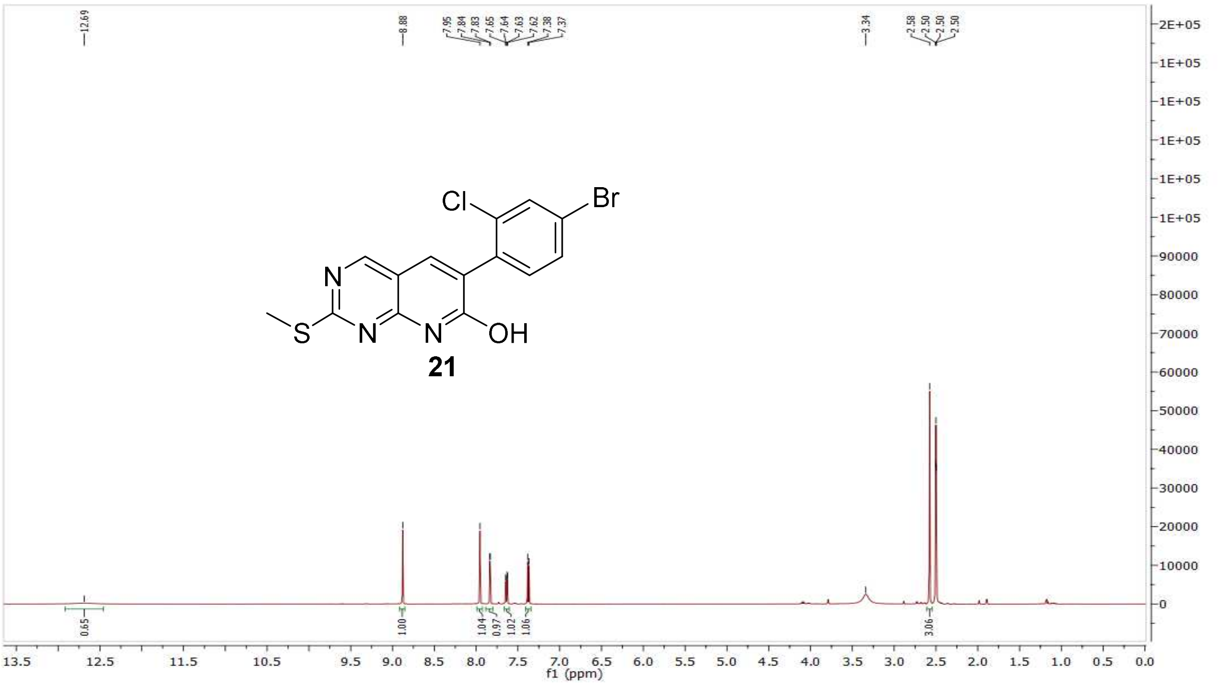

$^{13}C$ NMR compound **22**:

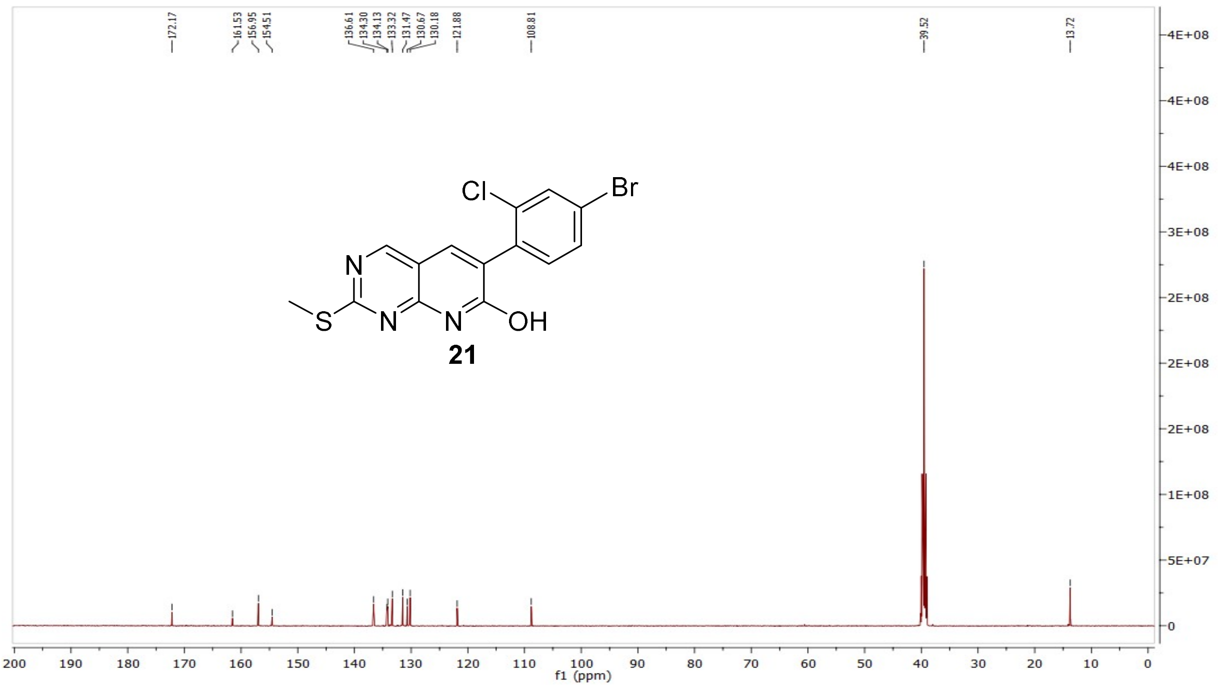


$^{1}H$ NMR compound **23**:

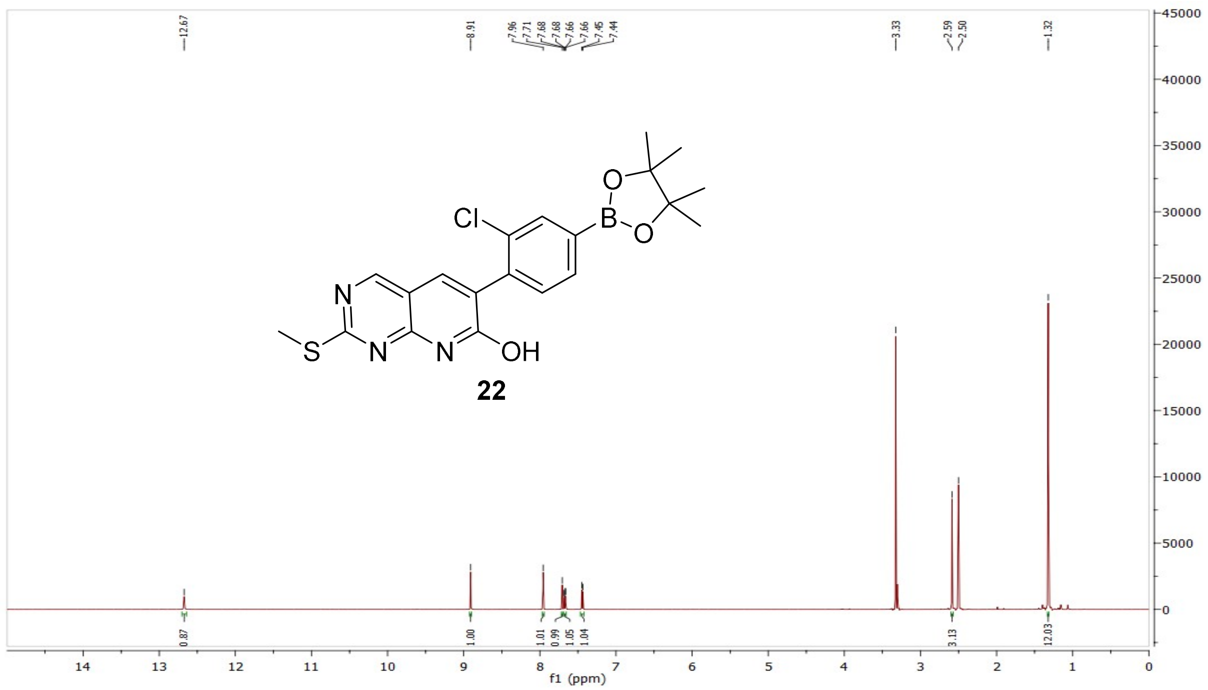

[13]C NMR compound **23**:

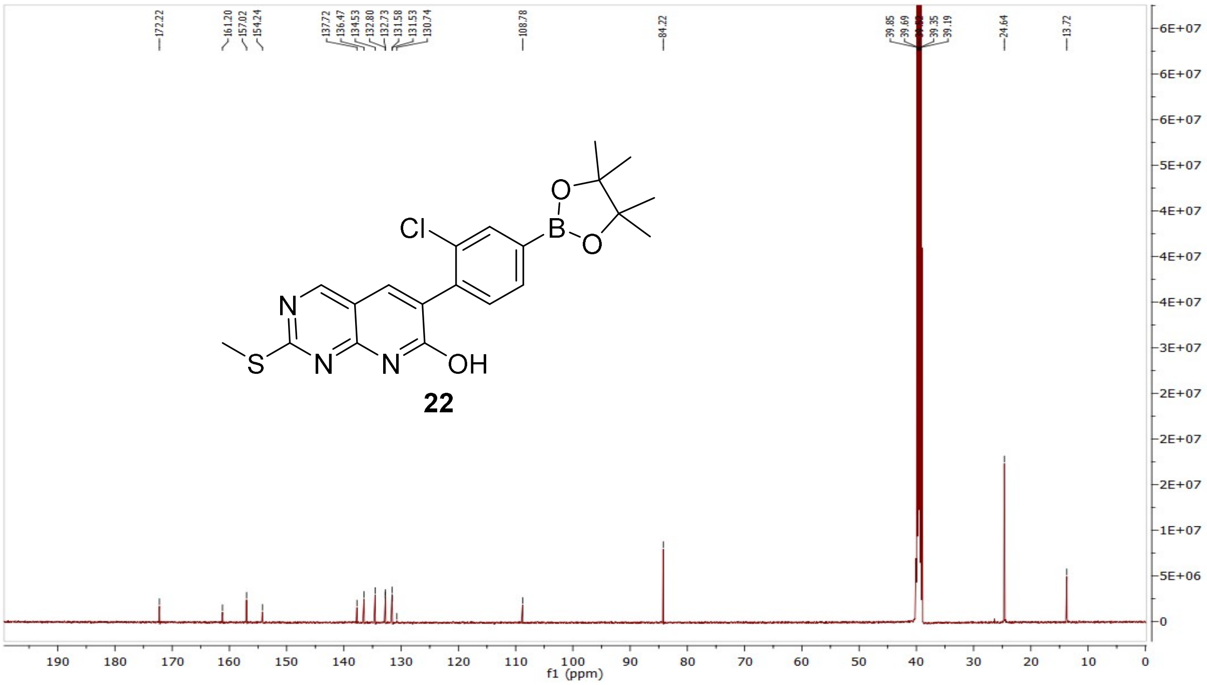


MS (ESI+) compound **23**:

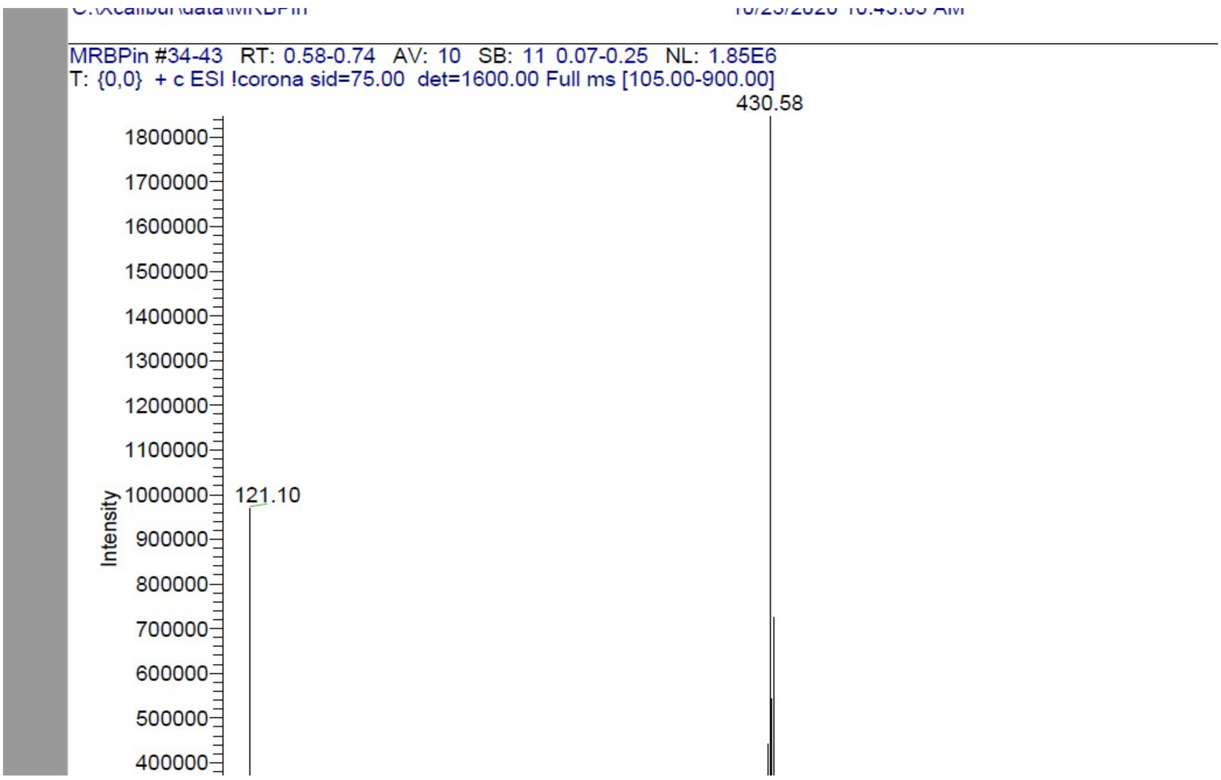

$^{1}$H NMR compound **25**:

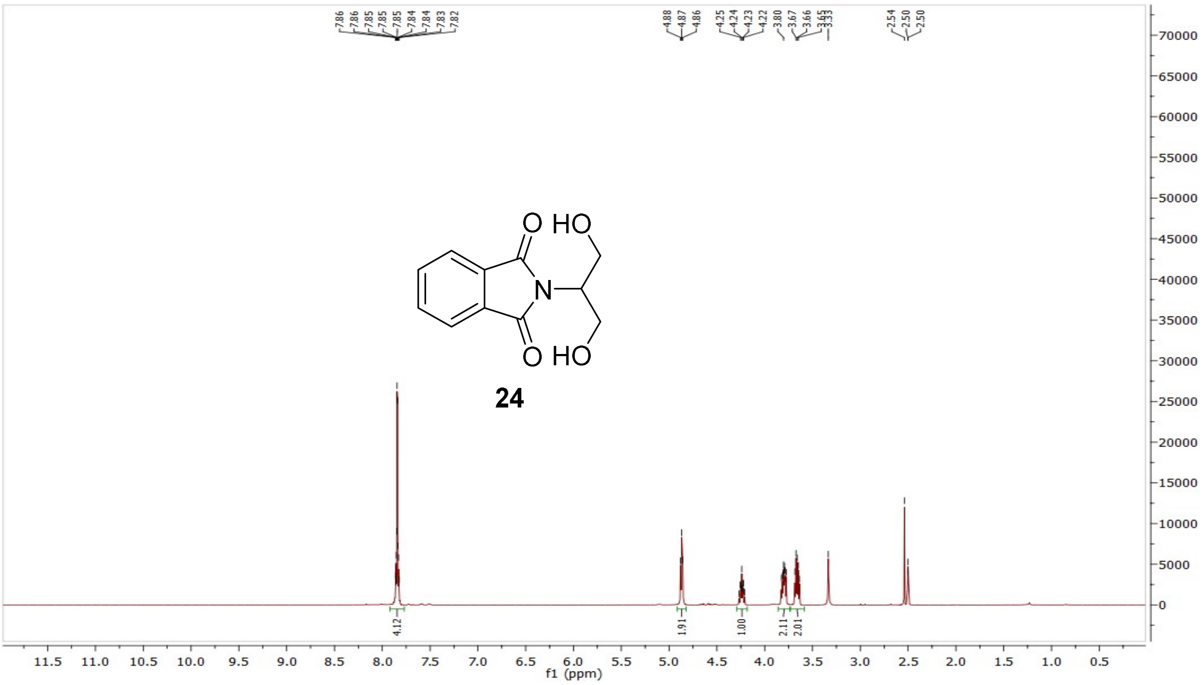


$^{13}$C NMR compound **25**:

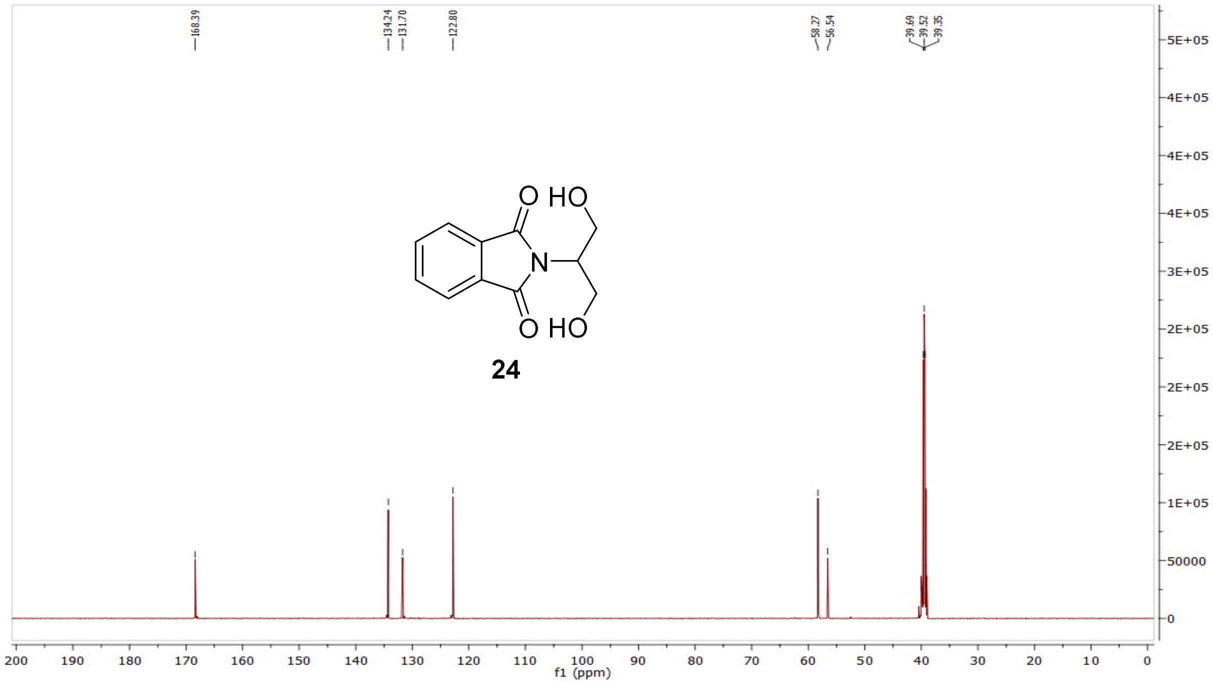

MS (ESI+) compound **25**:

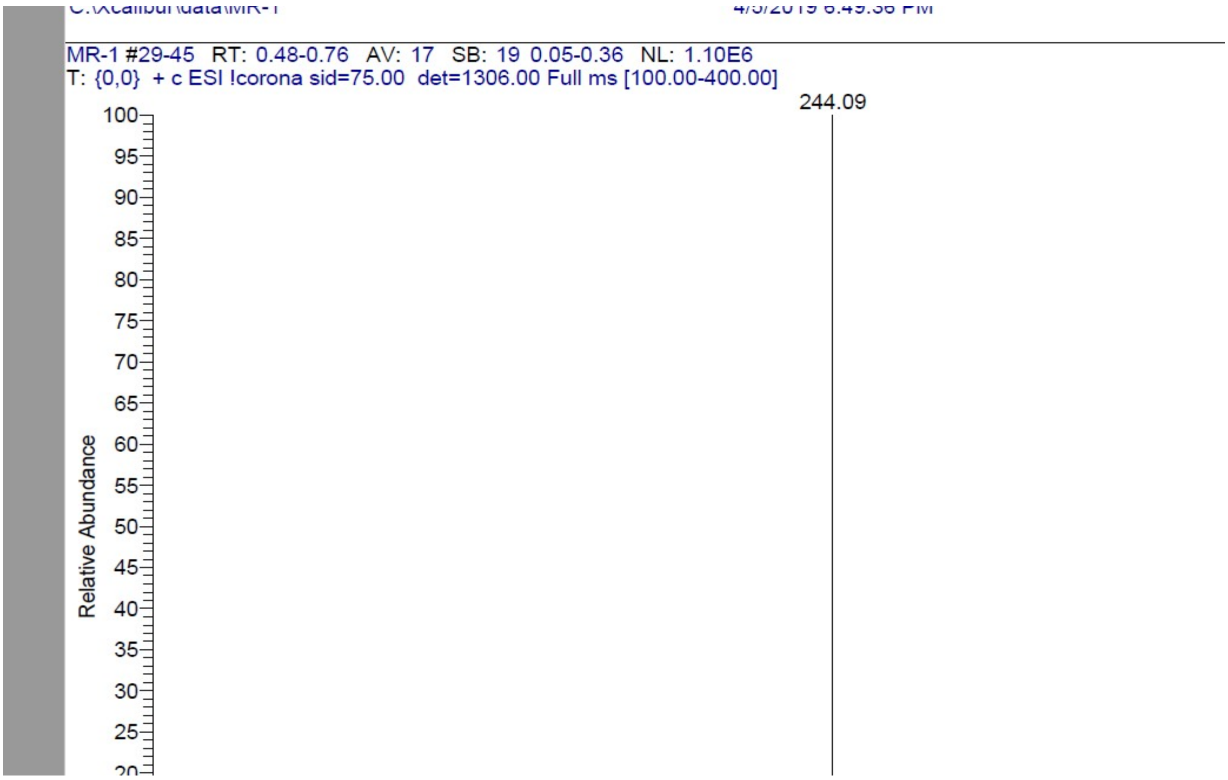


[1]H NMR compound **26**:

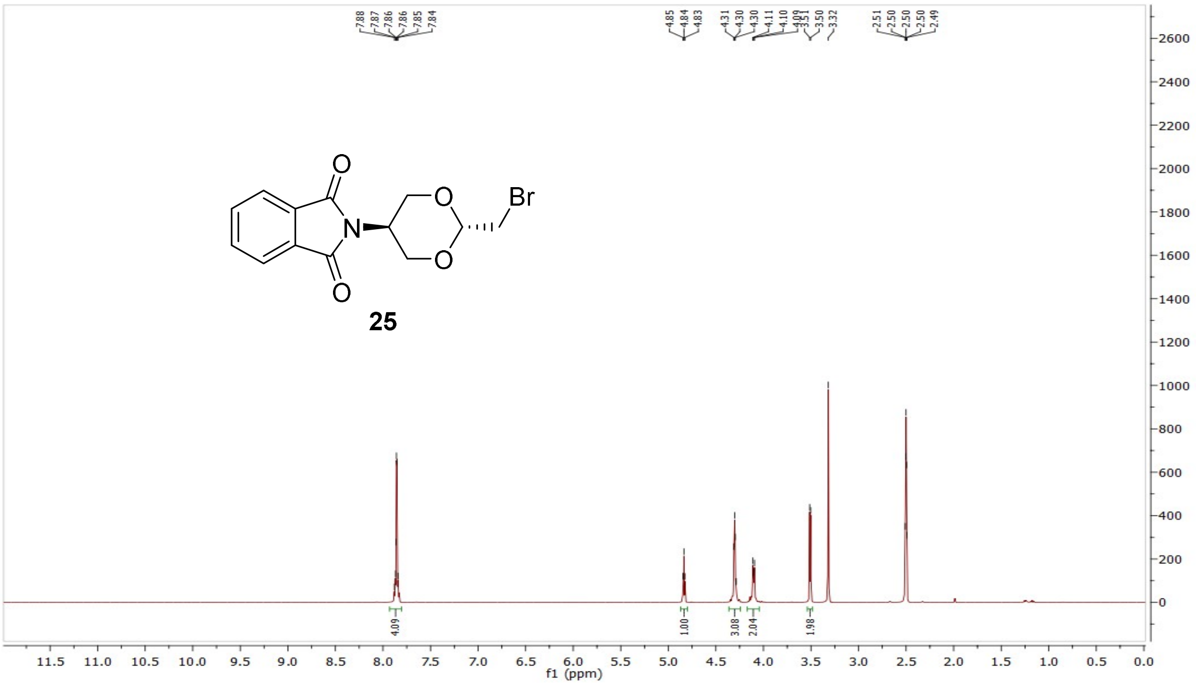

$^{13}$C NMR compound **26**:

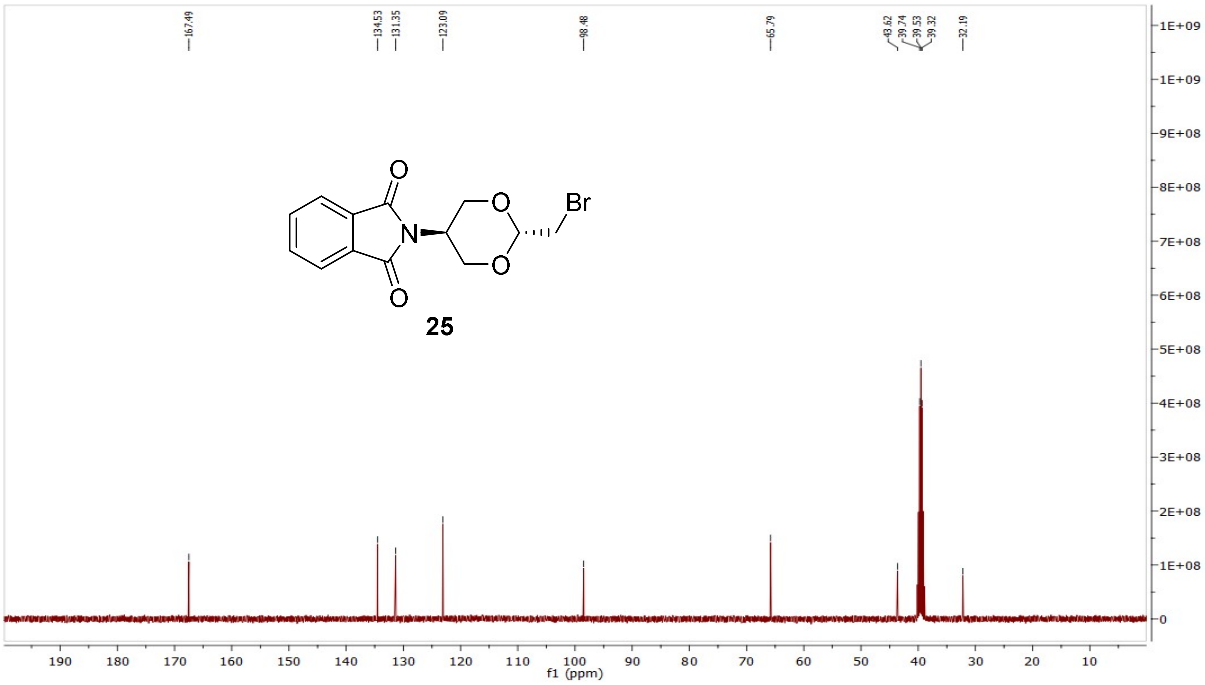


MS (ESI+) compound **26**:

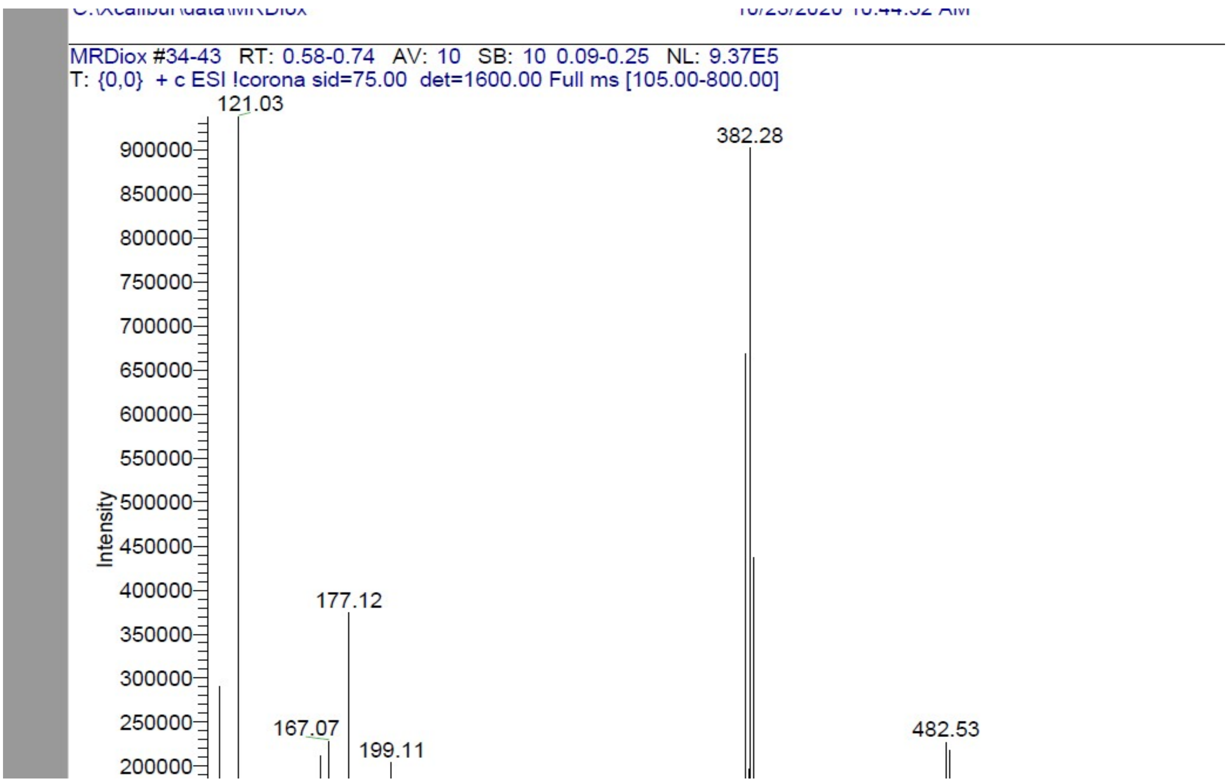

[1]H NMR compound **27**:

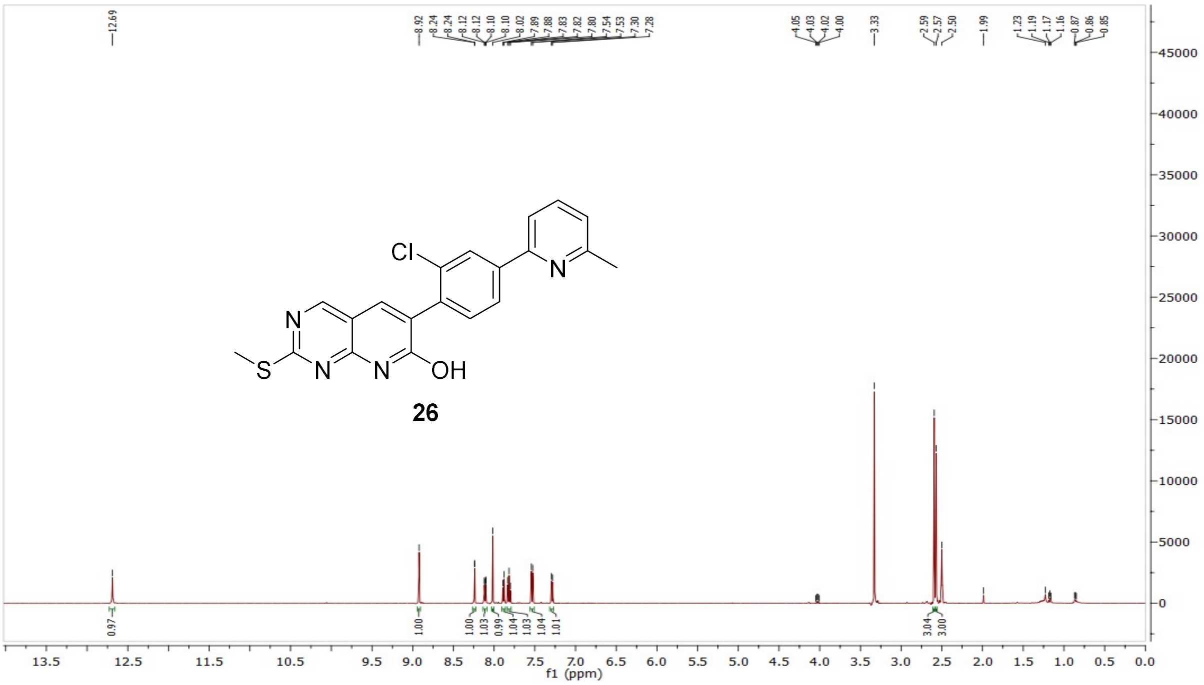


[13]C NMR compound **27**:

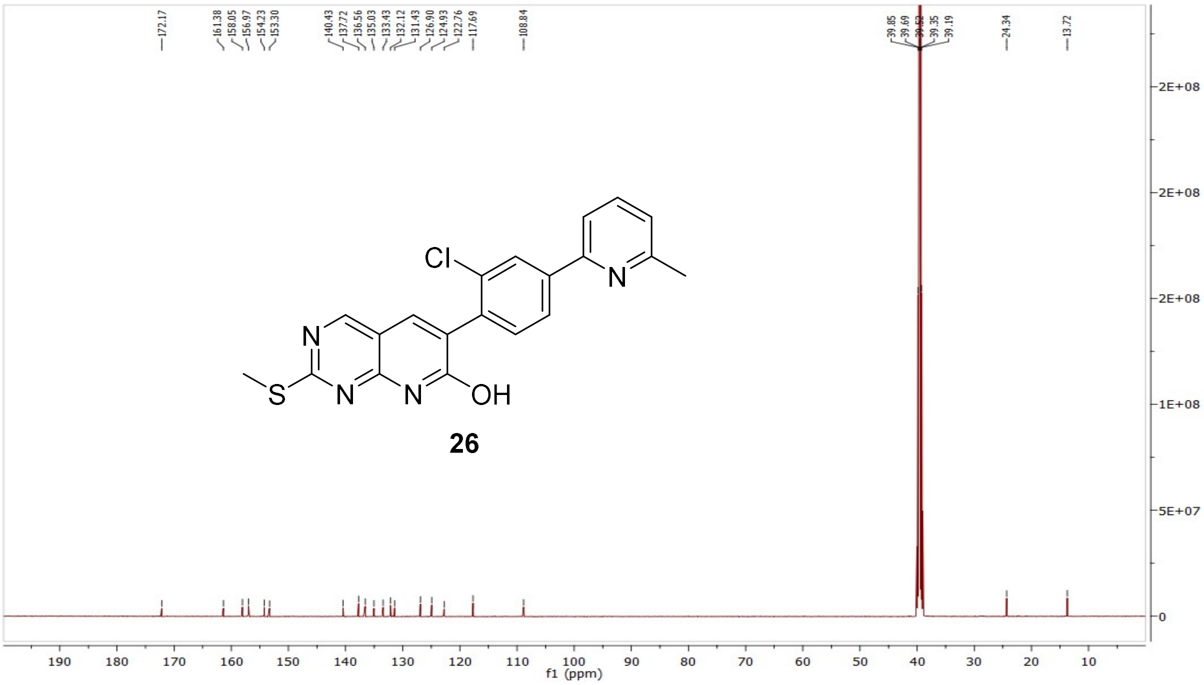

MS (ESI+) compound **27**:

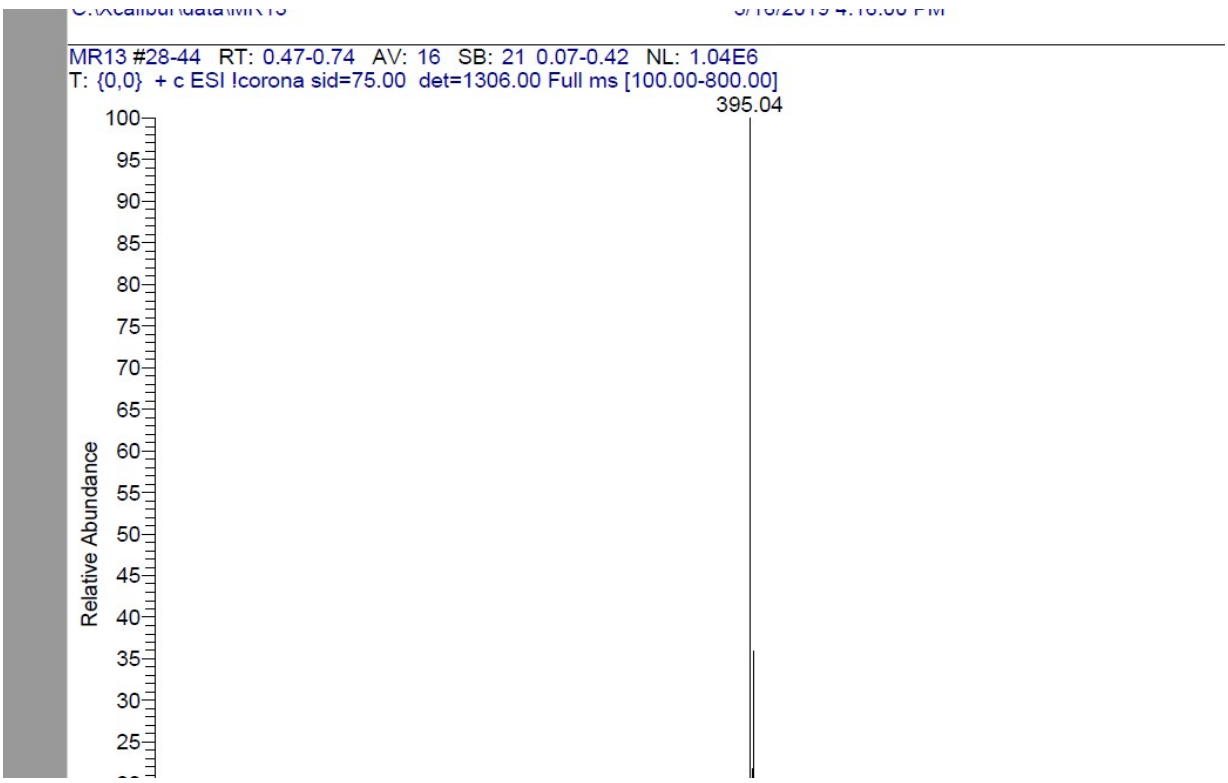


$^1$H NMR compound **28**:

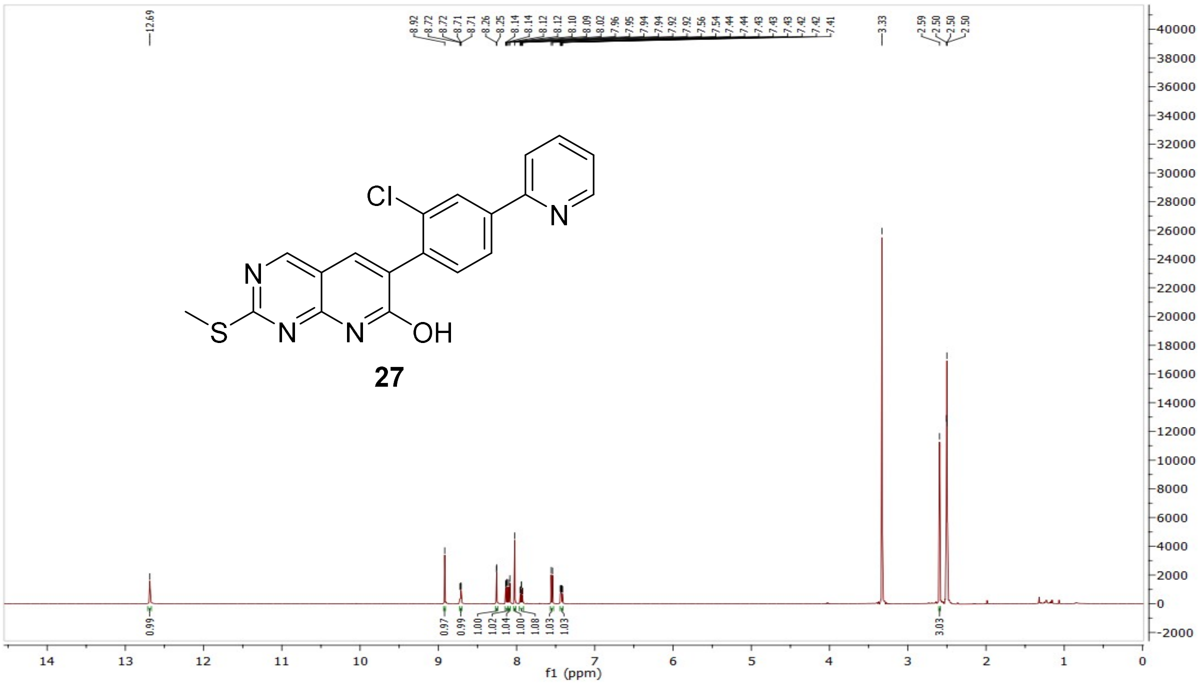

[13]C NMR compound **28**:

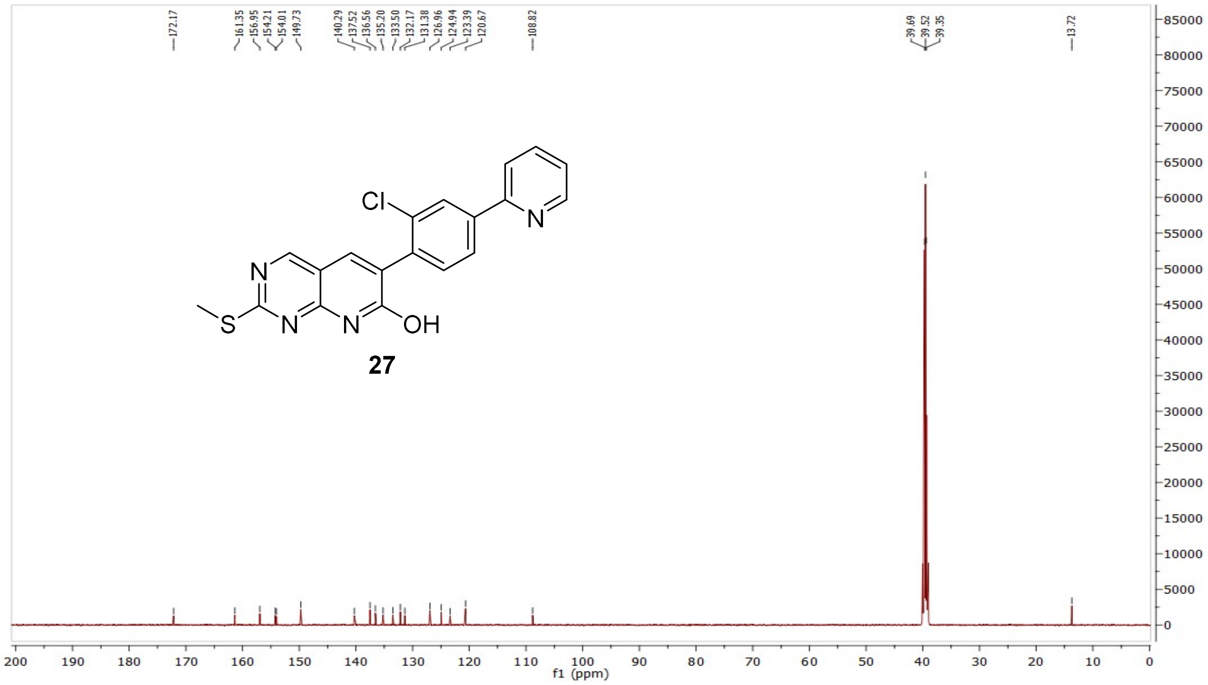


MS (ESI+) compound **28**:

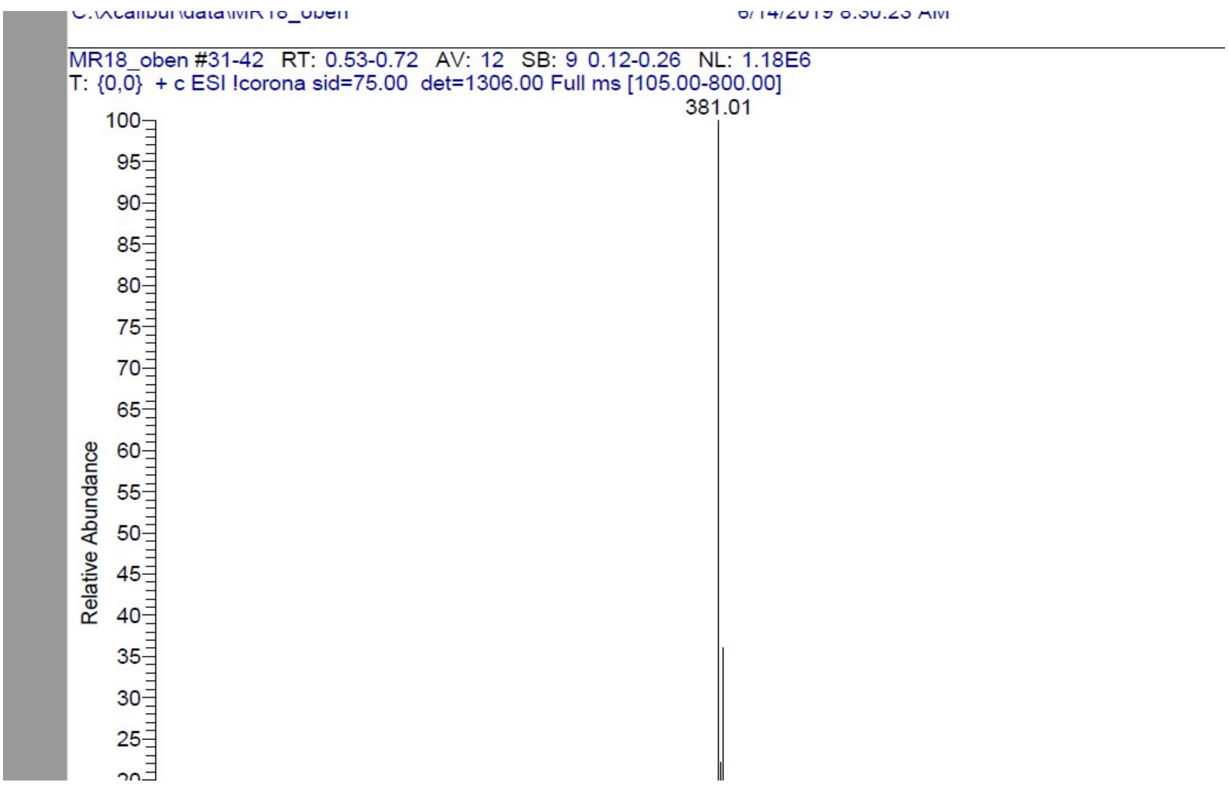

$^{1}$H NMR compound **29**:

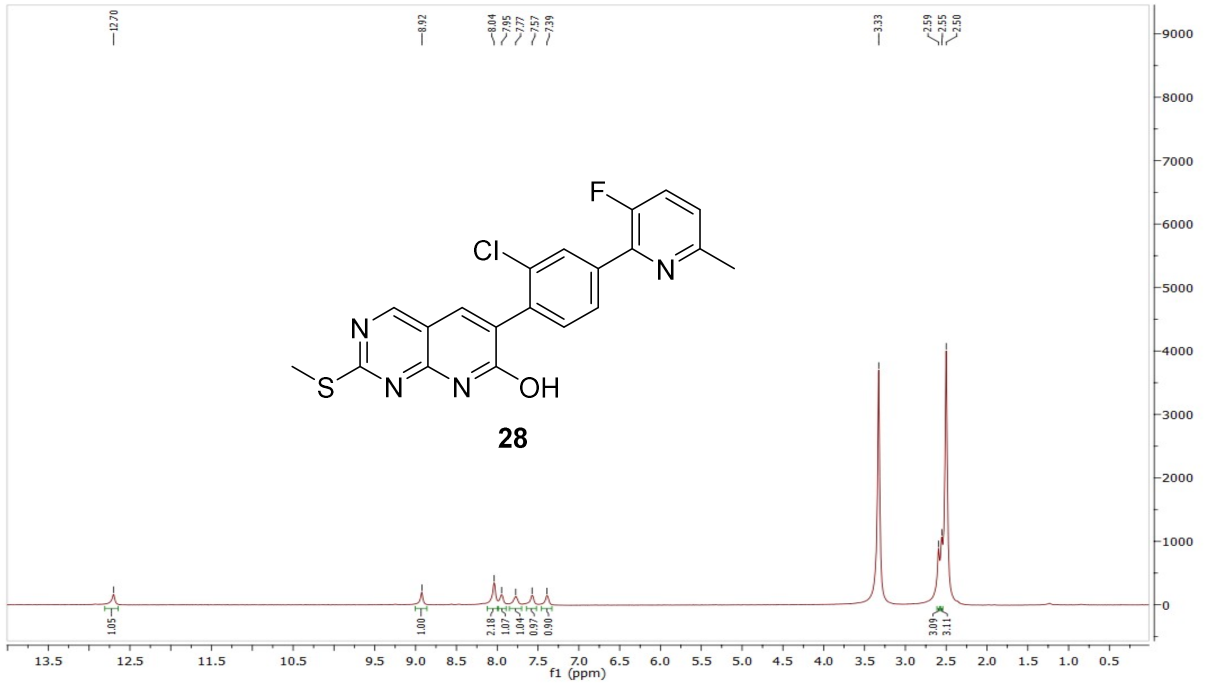


$^{13}$C NMR compound **29**:

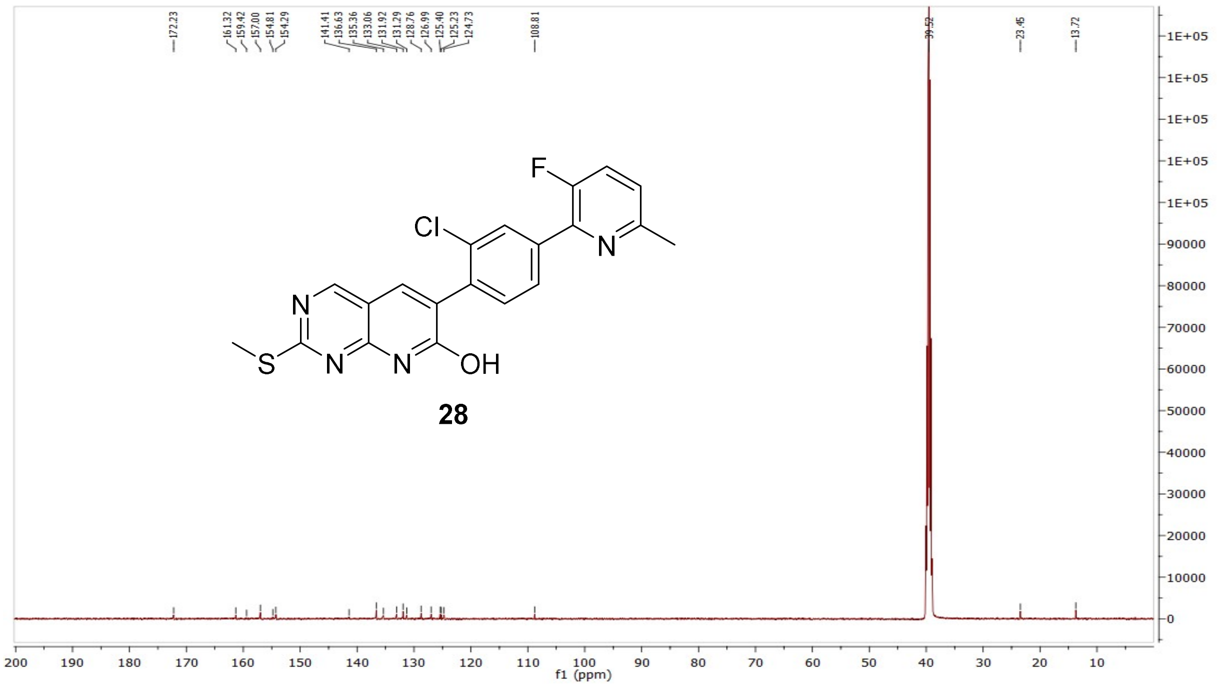

MS (ESI+) compound **29**:

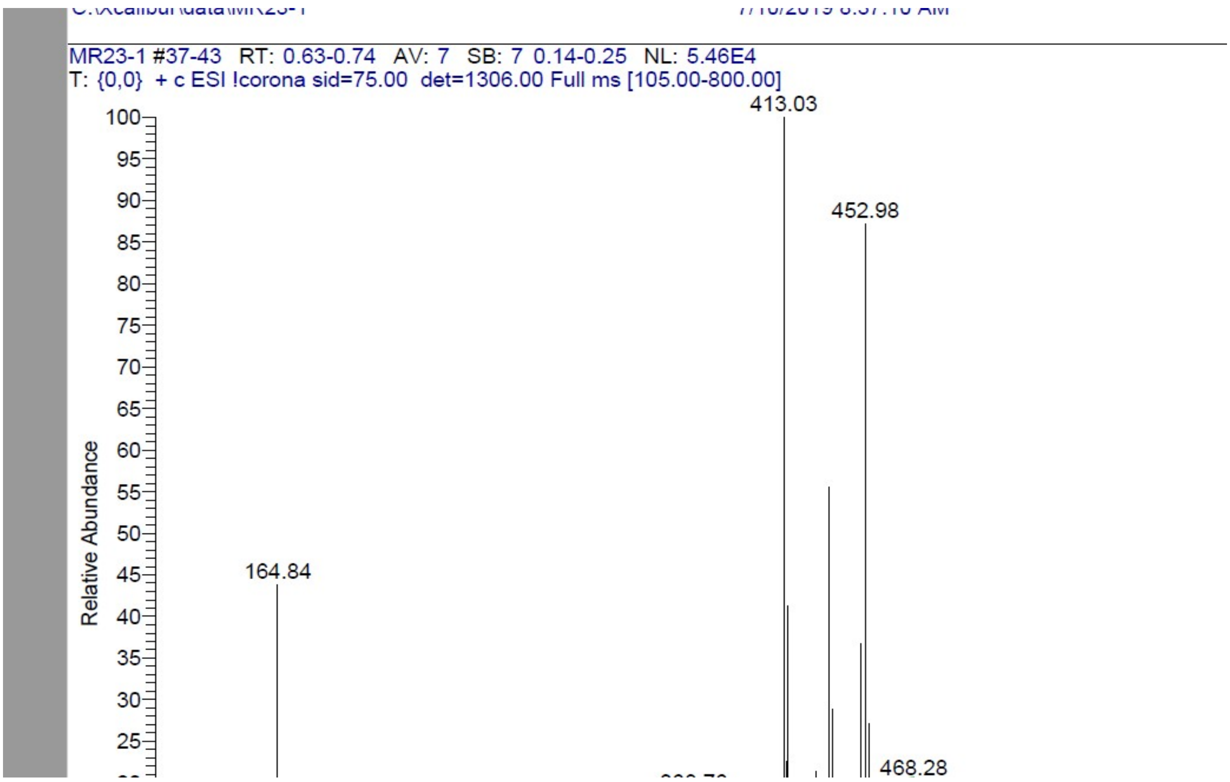


[1]H NMR compound **30**:

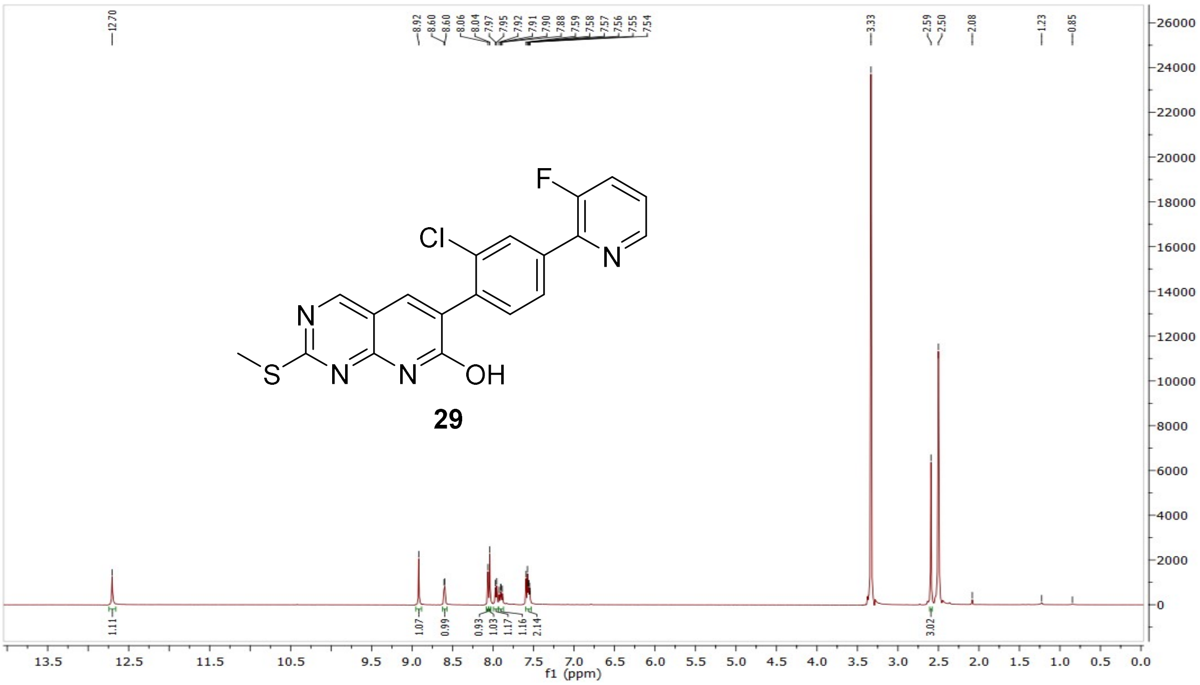

$^{13}C$ NMR compound **30**:

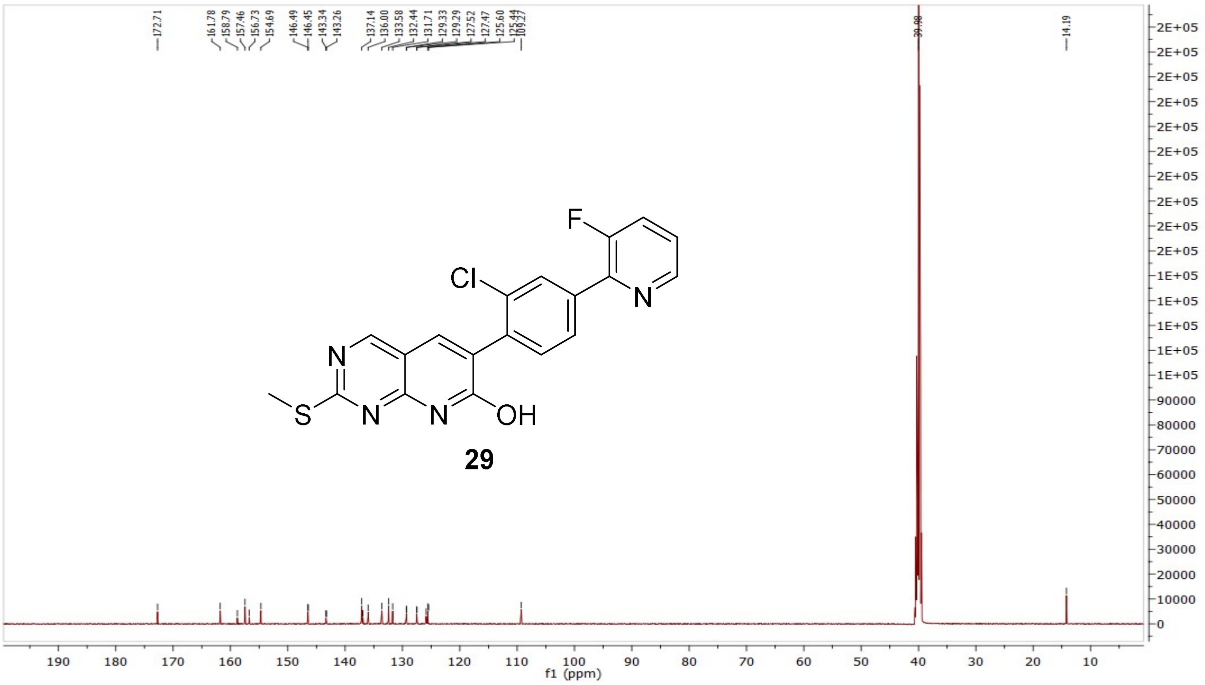

MS (ESI-) compound **30**:

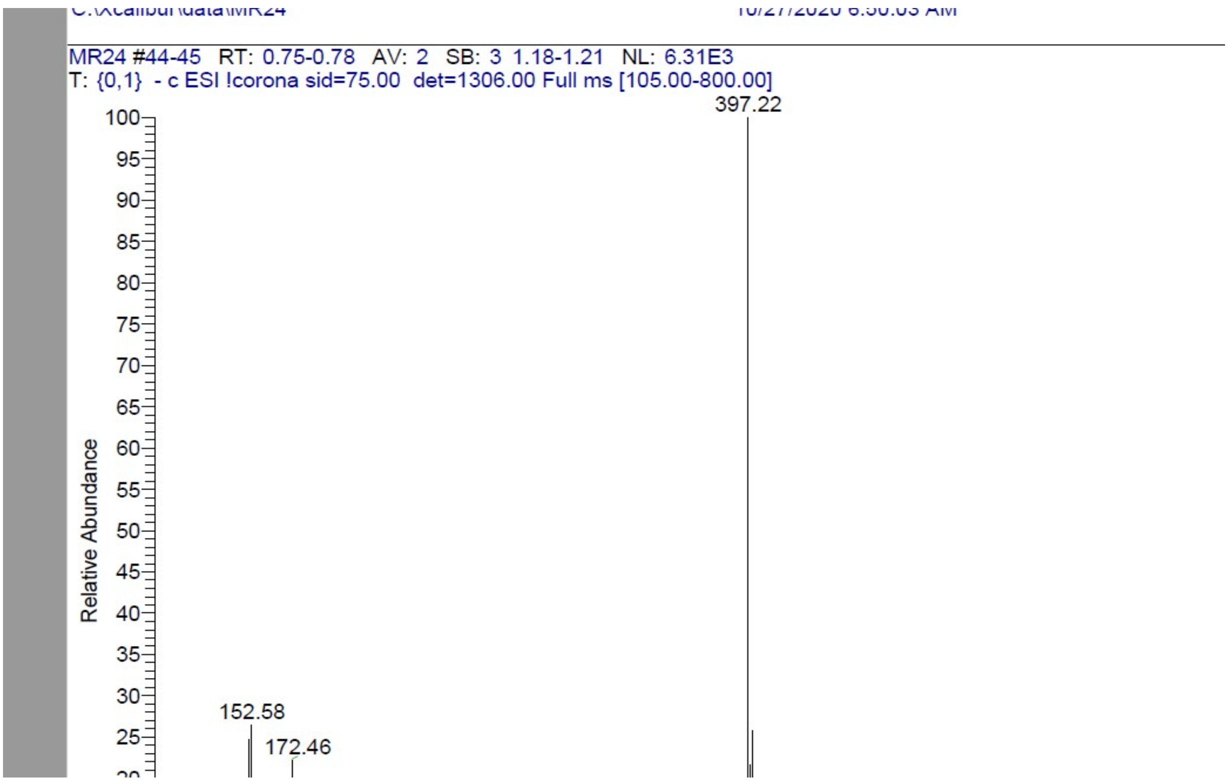

$^{1}$H NMR compound **31**:

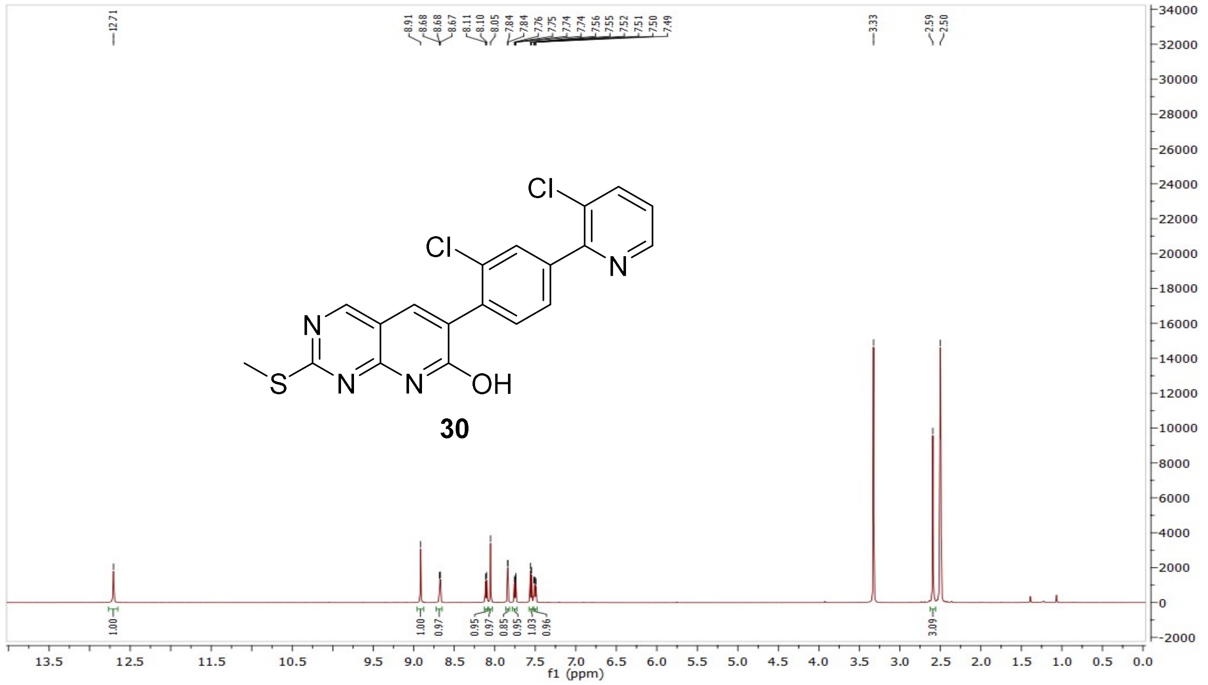


$^{13}$C NMR compound **31**:

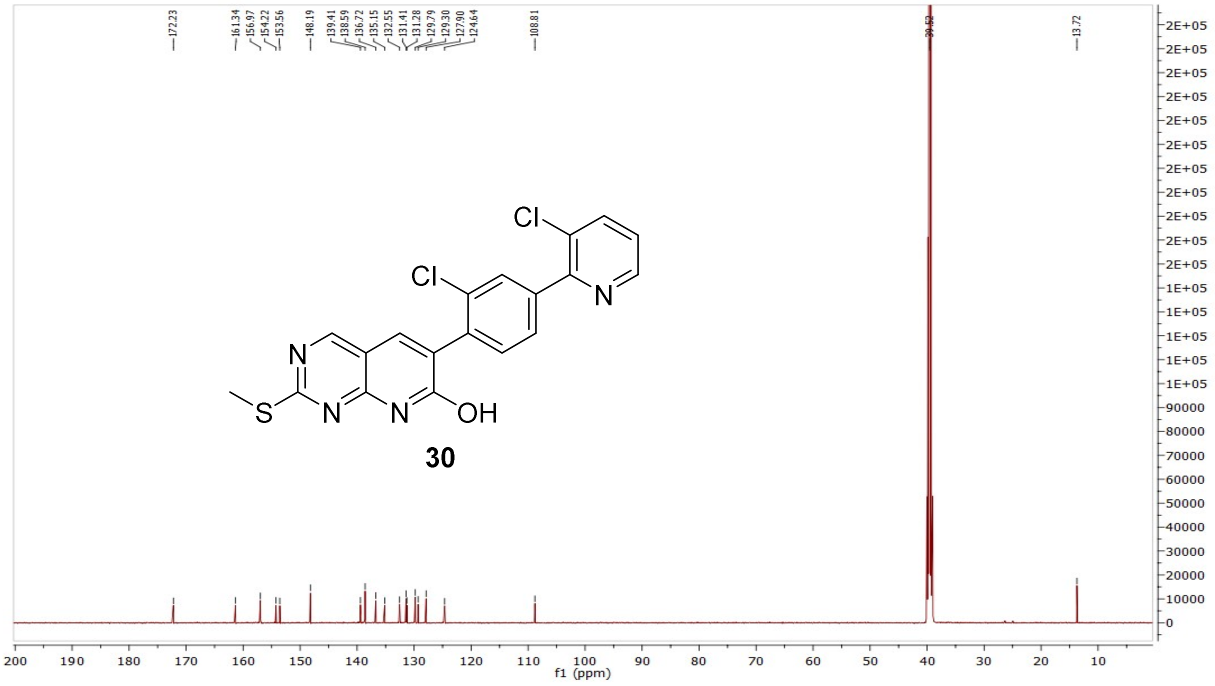

MS (ESI+) compound **31**:

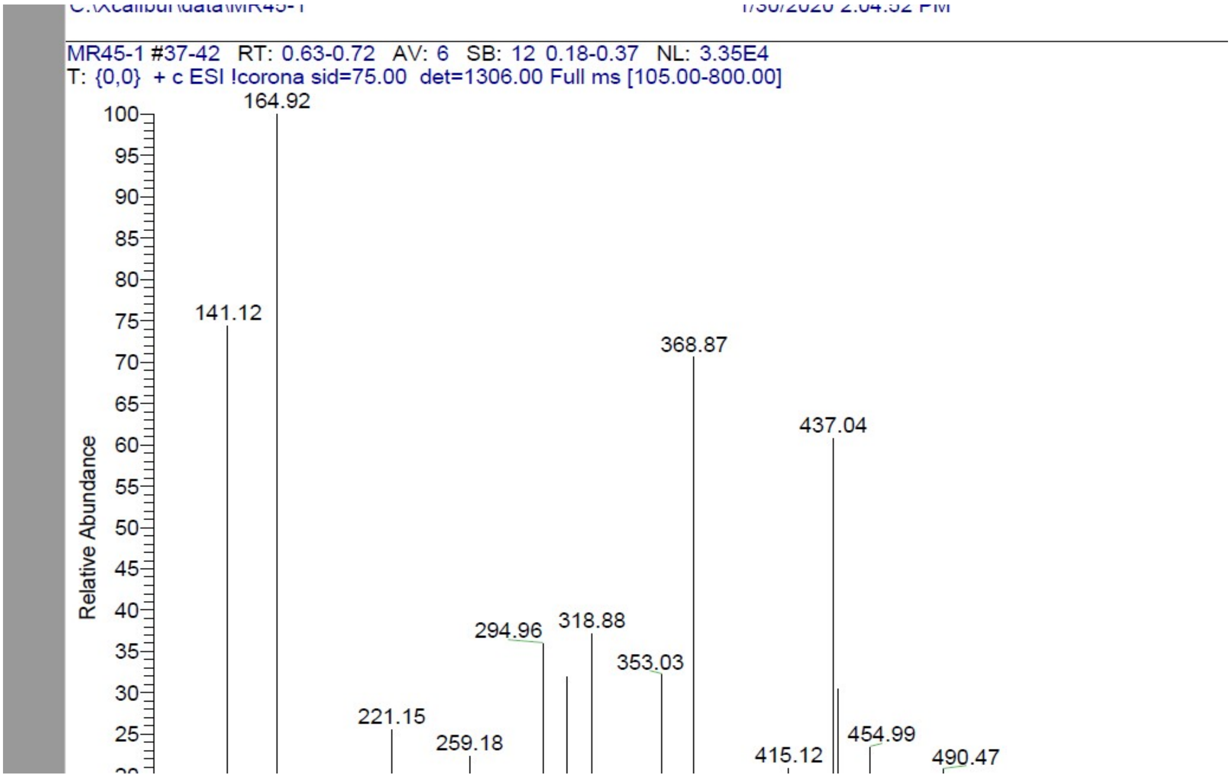


[1]H NMR compound **32**:

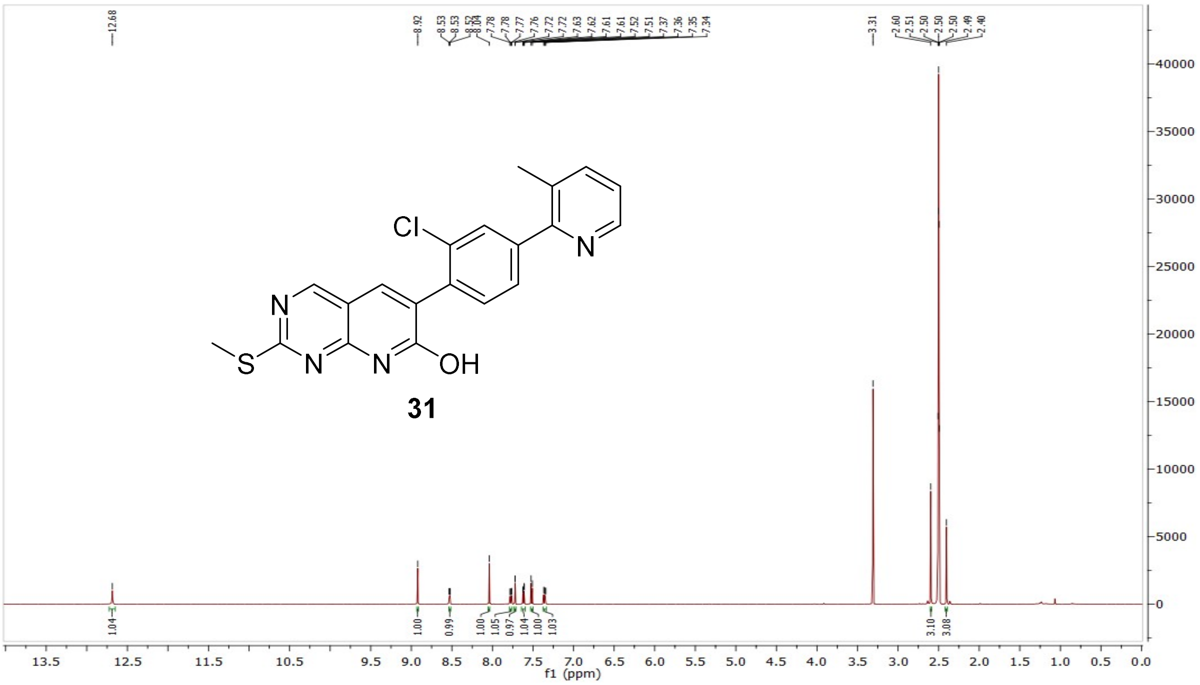

$^{13}$C NMR compound **32**:

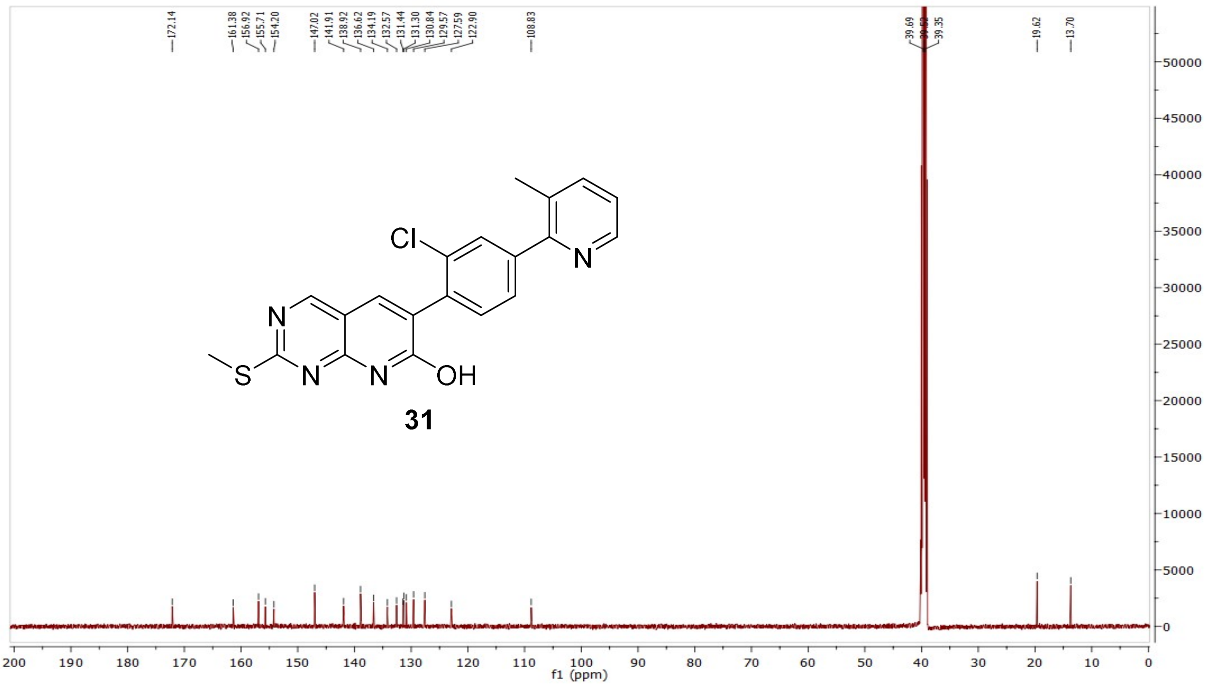


MS (ESI+) compound **32**:

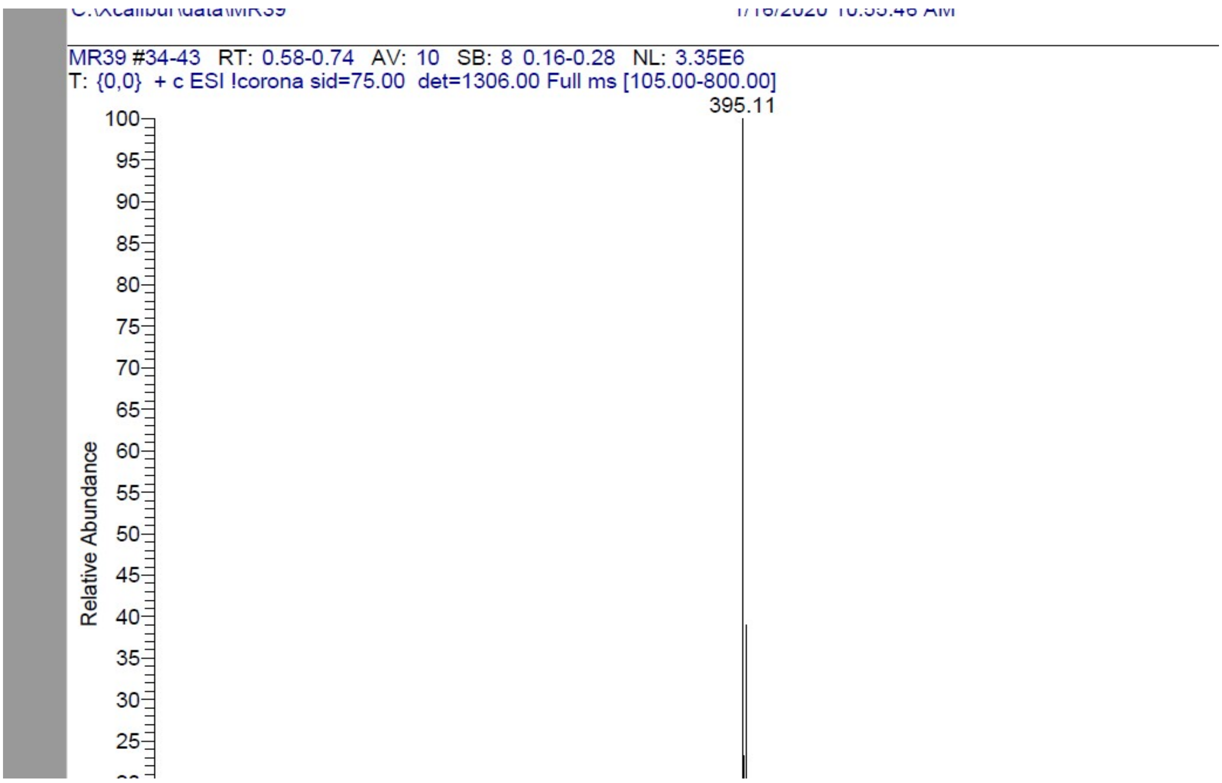

$^{1}$H NMR compound **33**:

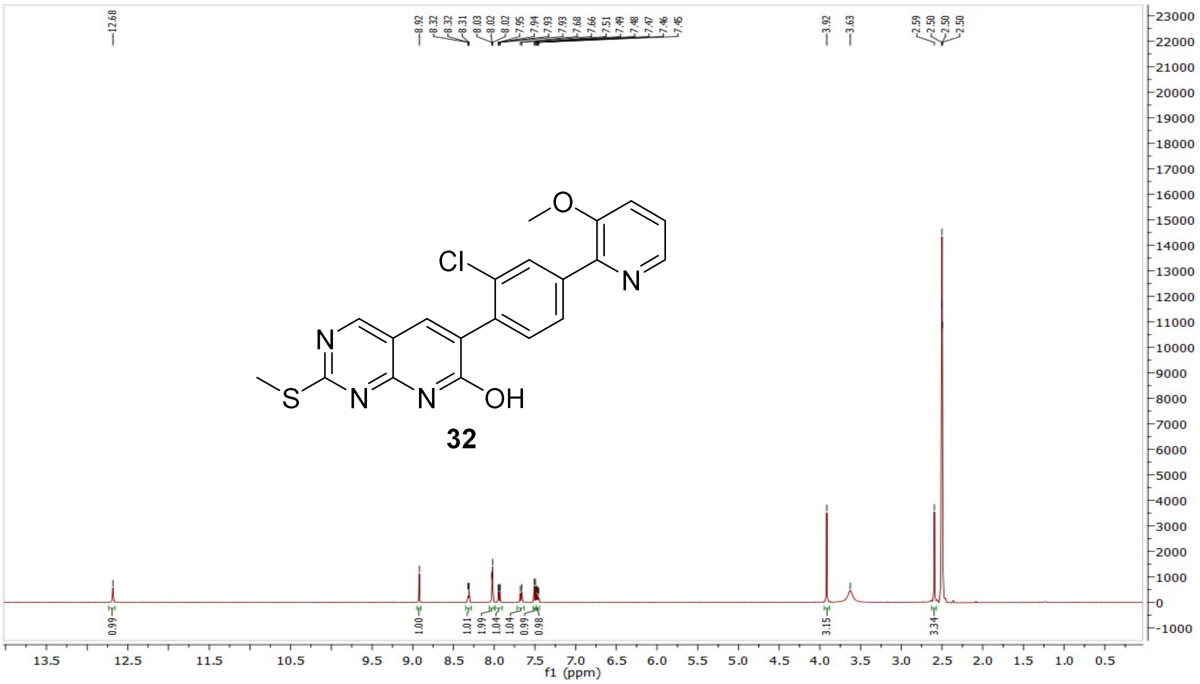


$^{13}$C NMR compound **33**:

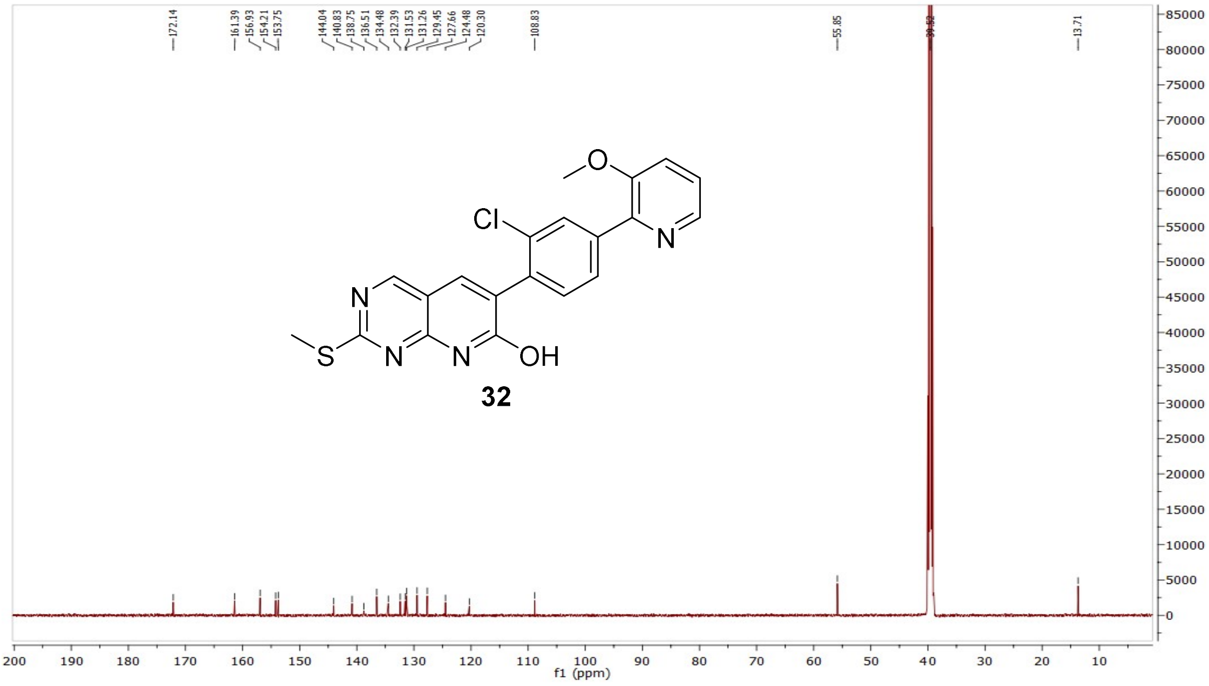

MS (ESI+) compound **33**:

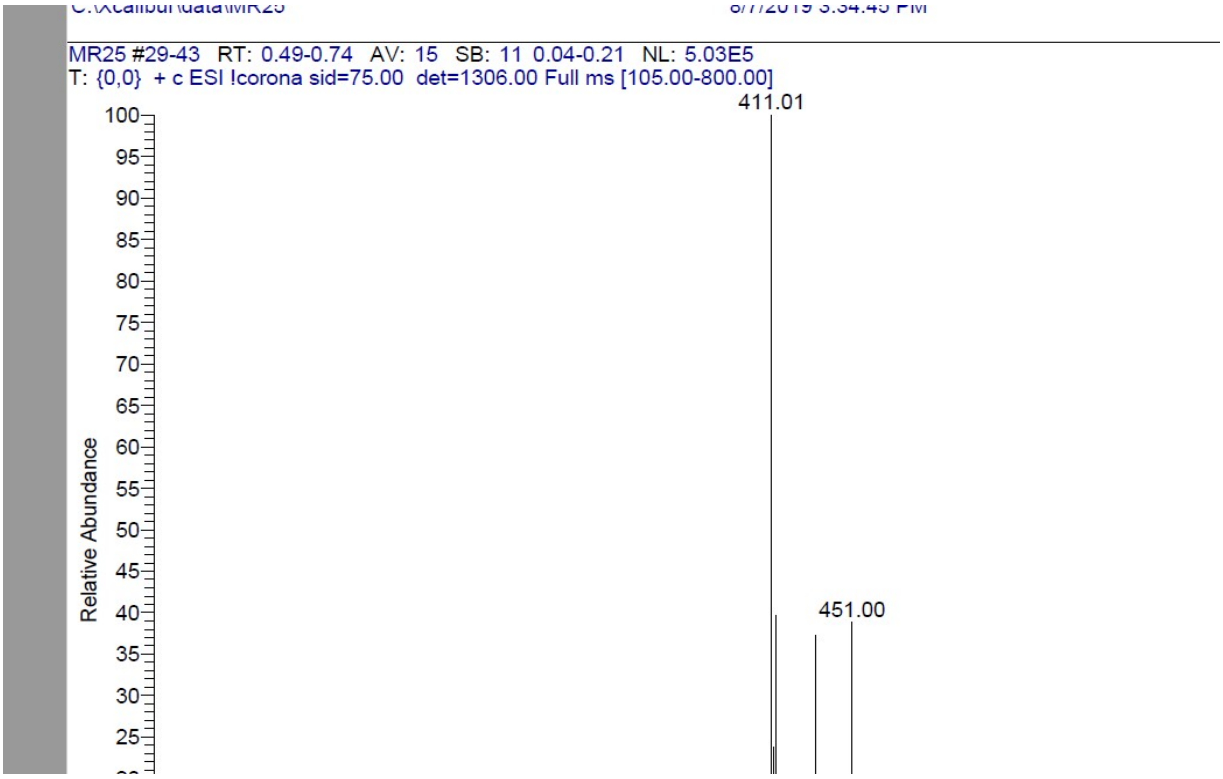


[1]H NMR compound **34**:

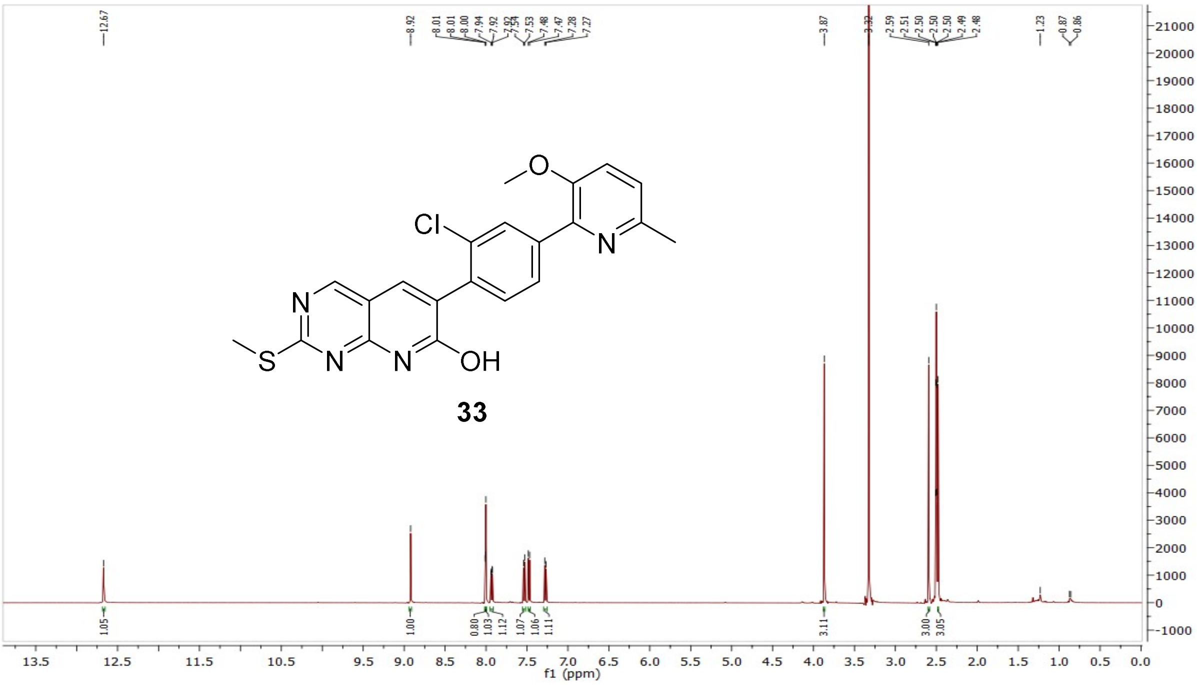

$^{13}$C NMR compound **34**:

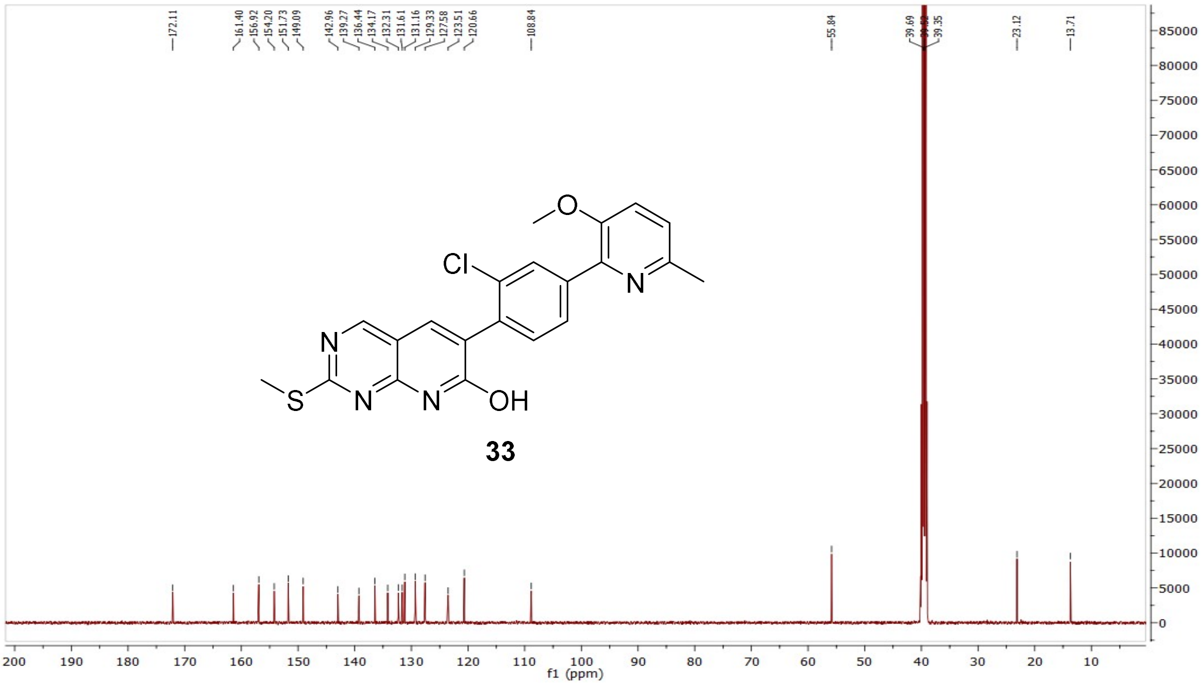


MS (ESI+) compound **34**:

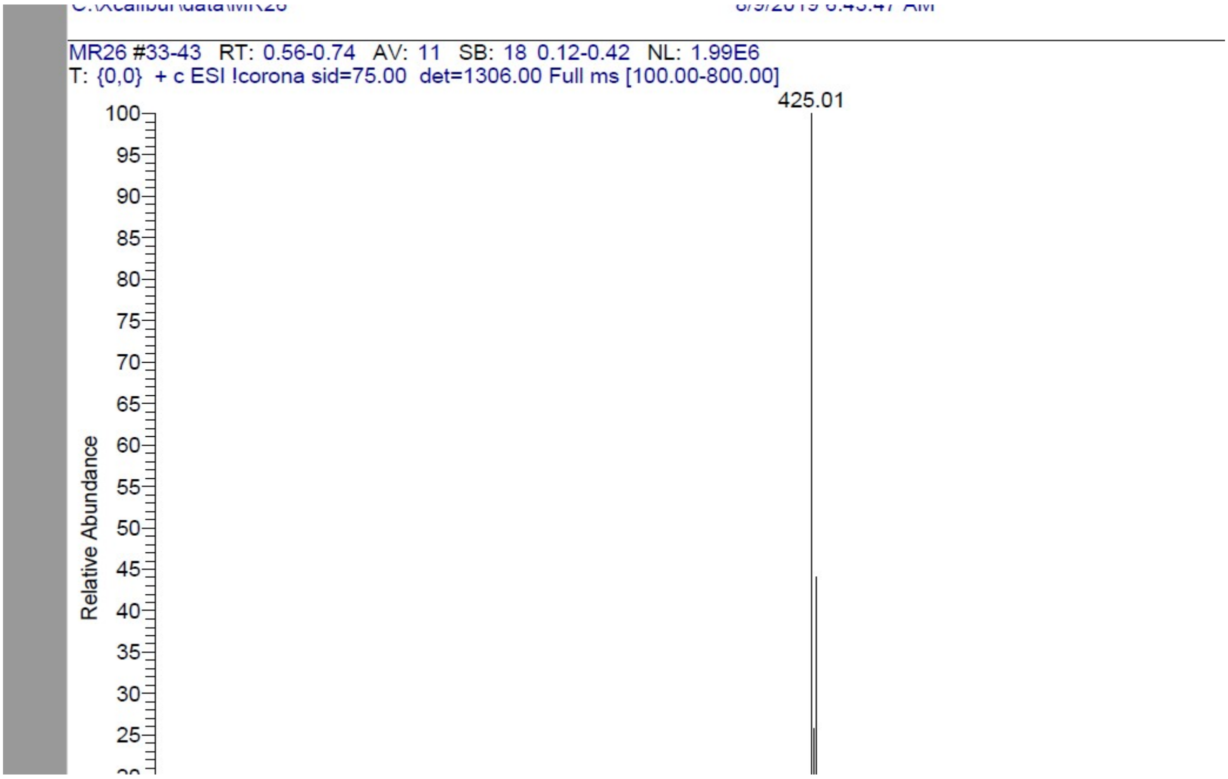

$^{1}$H NMR compound **35**:

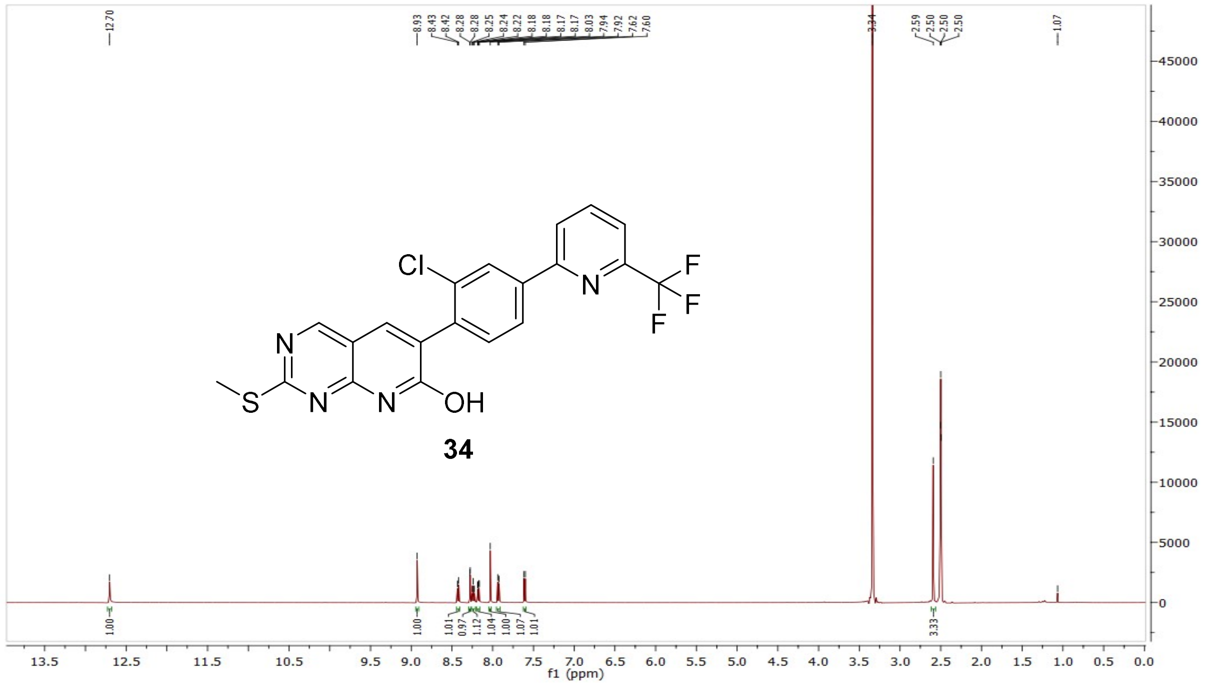


$^{13}$C NMR compound **35**:

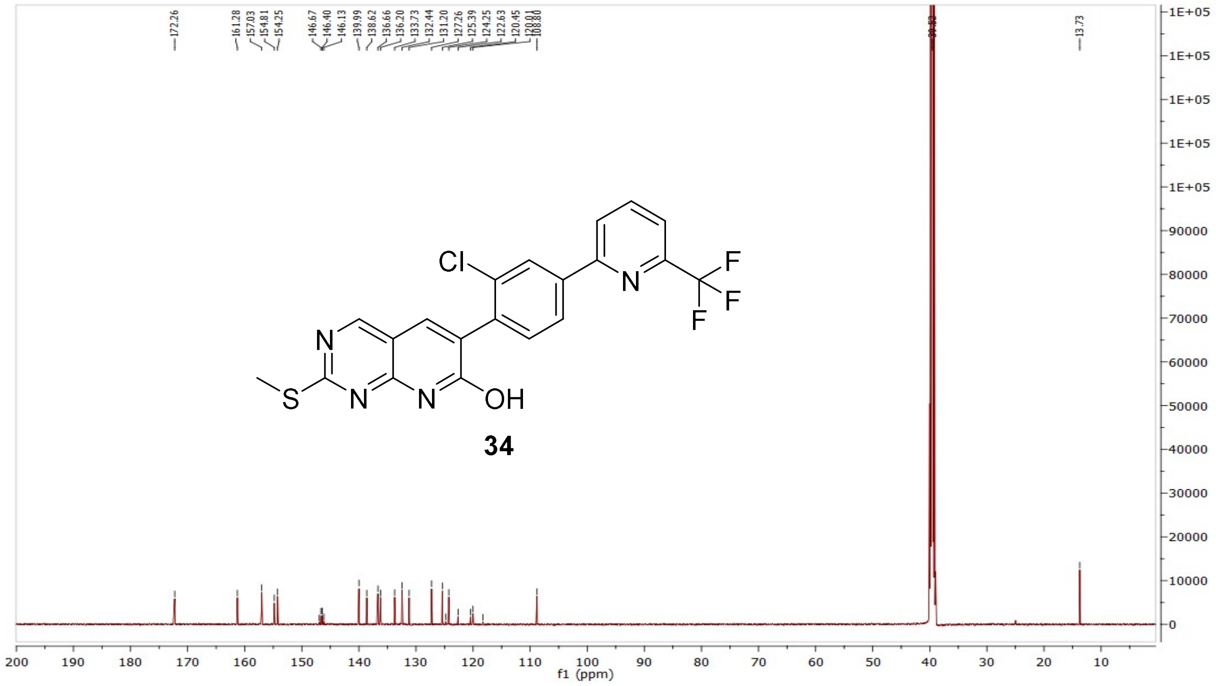

MS (ESI-) compound **35**:

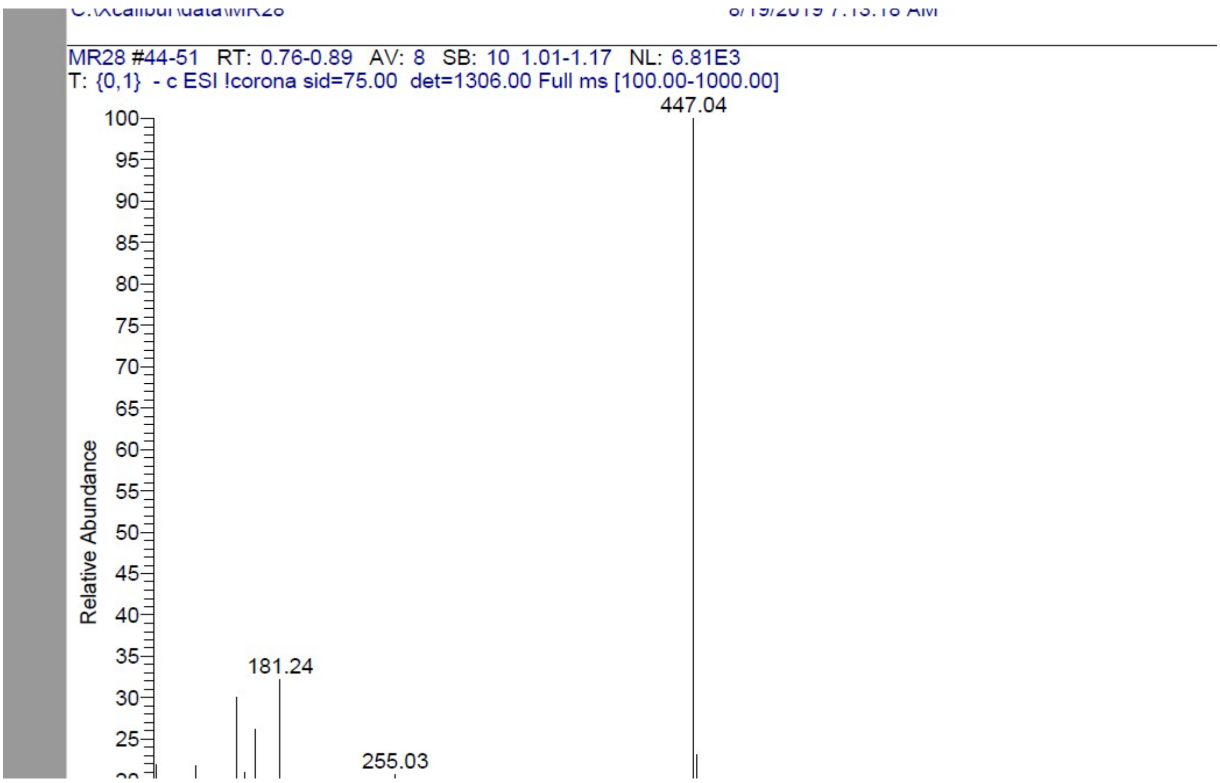


[1]H NMR compound **36**:

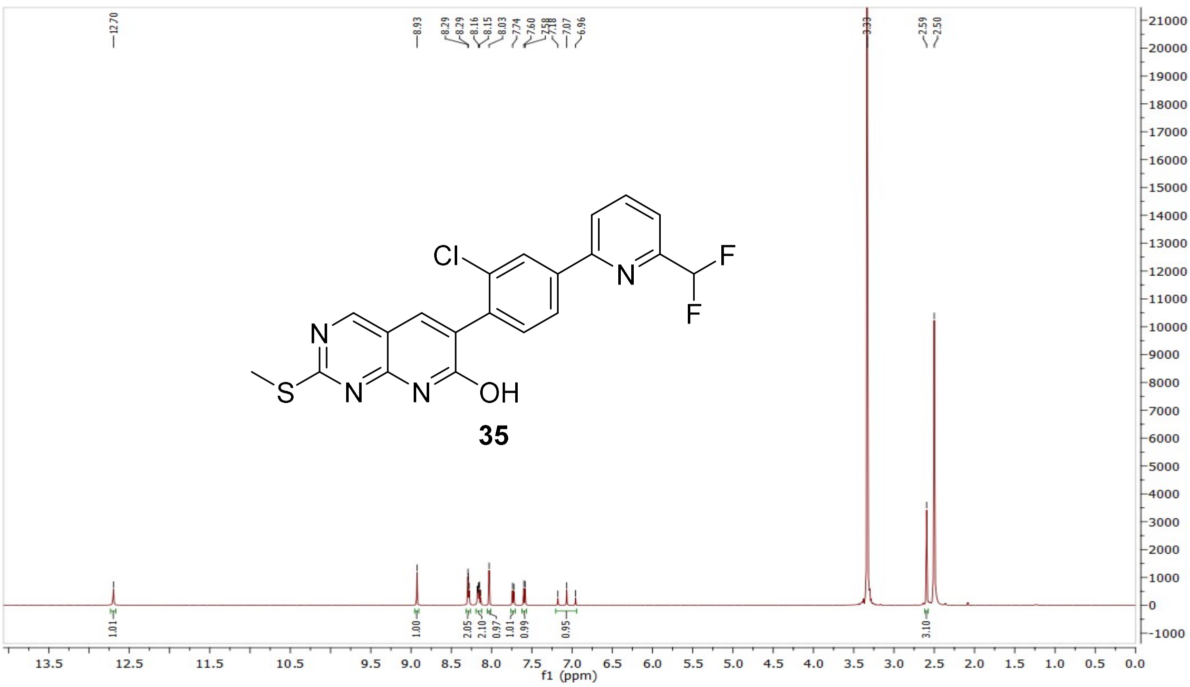

[13]C NMR compound **36**:

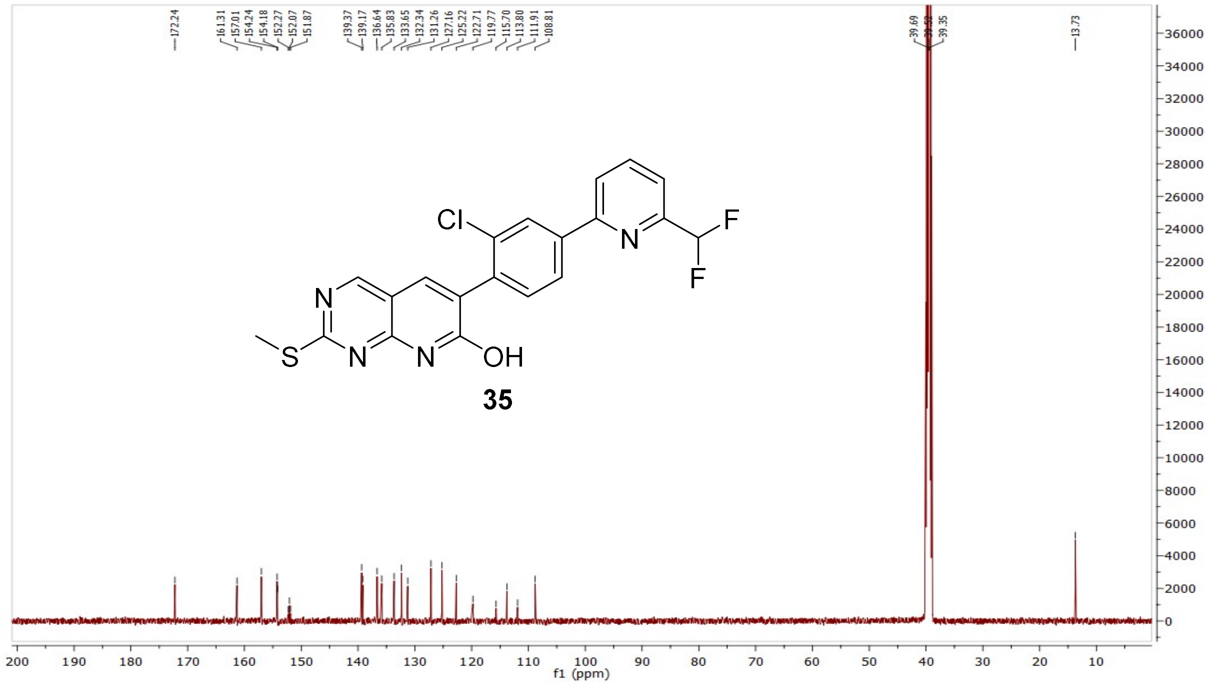


MS (ESI+) compound **36**:

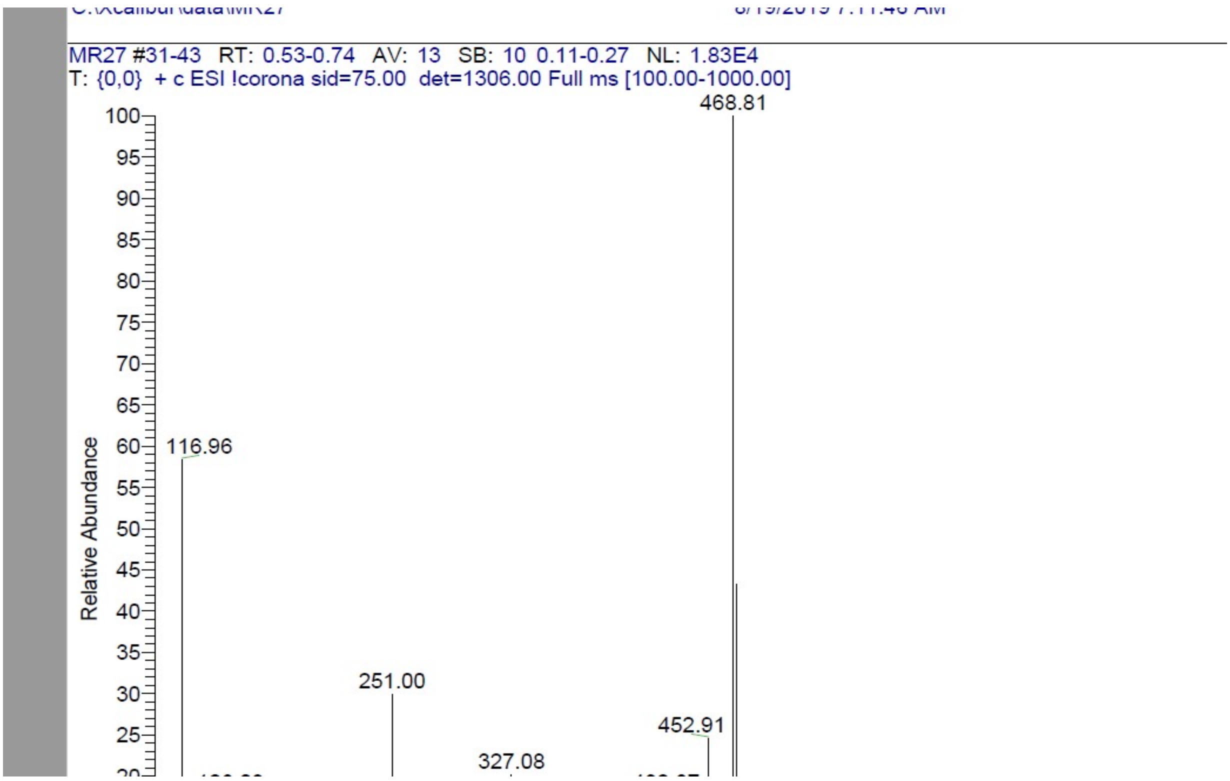

$^1$H NMR compound **37**:

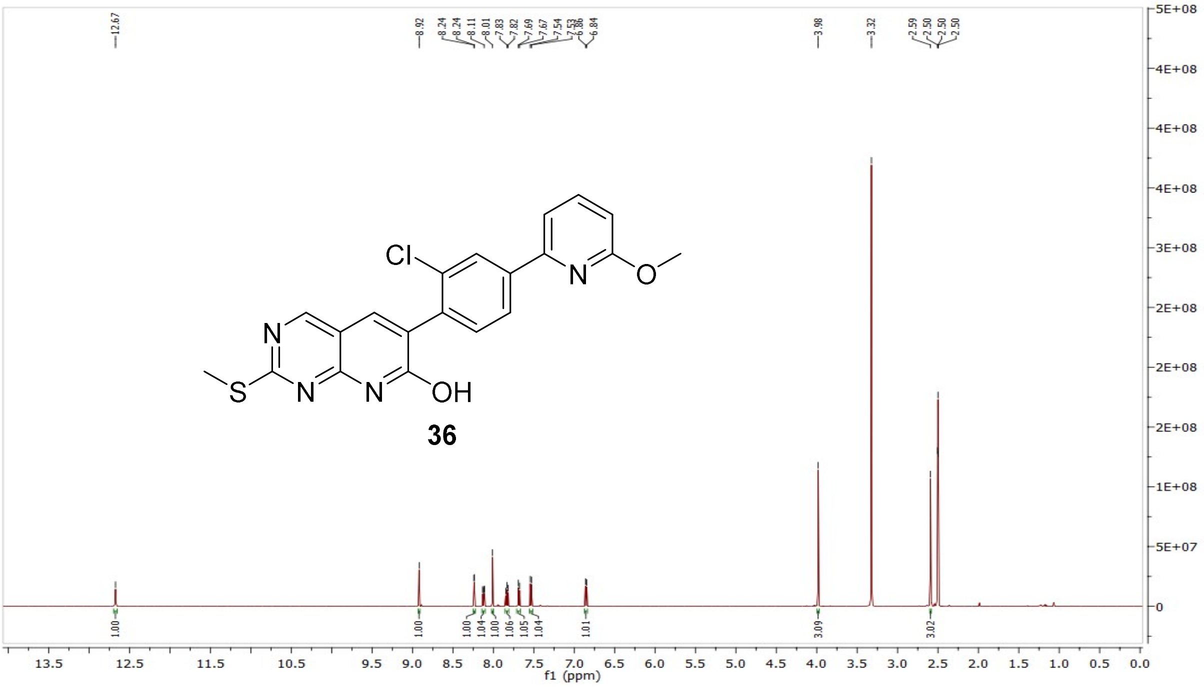


$^{13}$C NMR compound **37**:

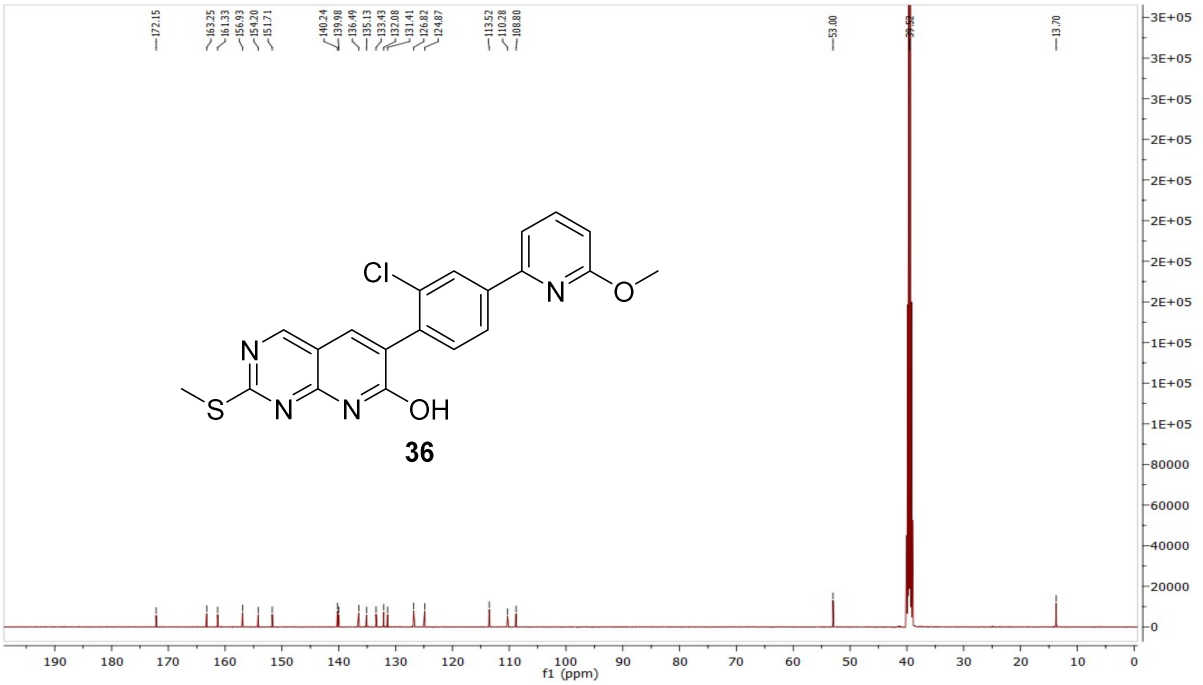

MS (ESI+) compound **37**:

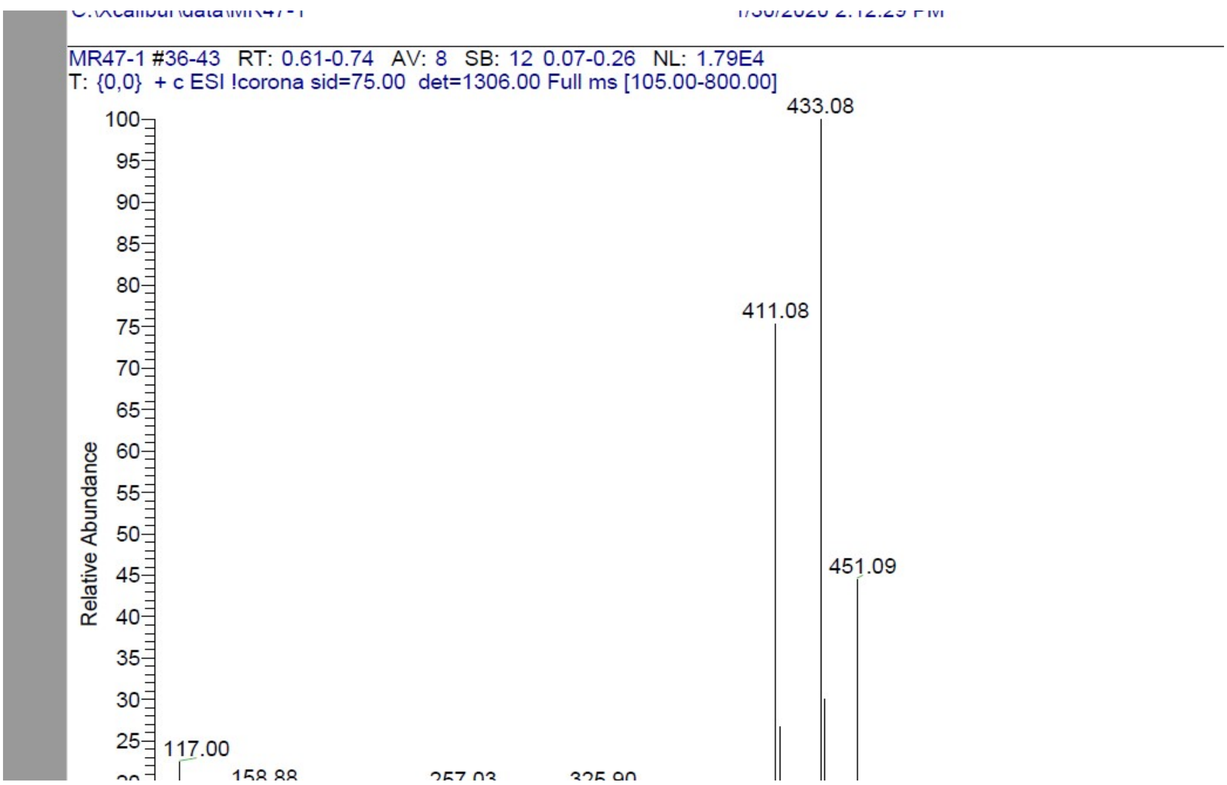


$^{1}$H NMR compound **38**:

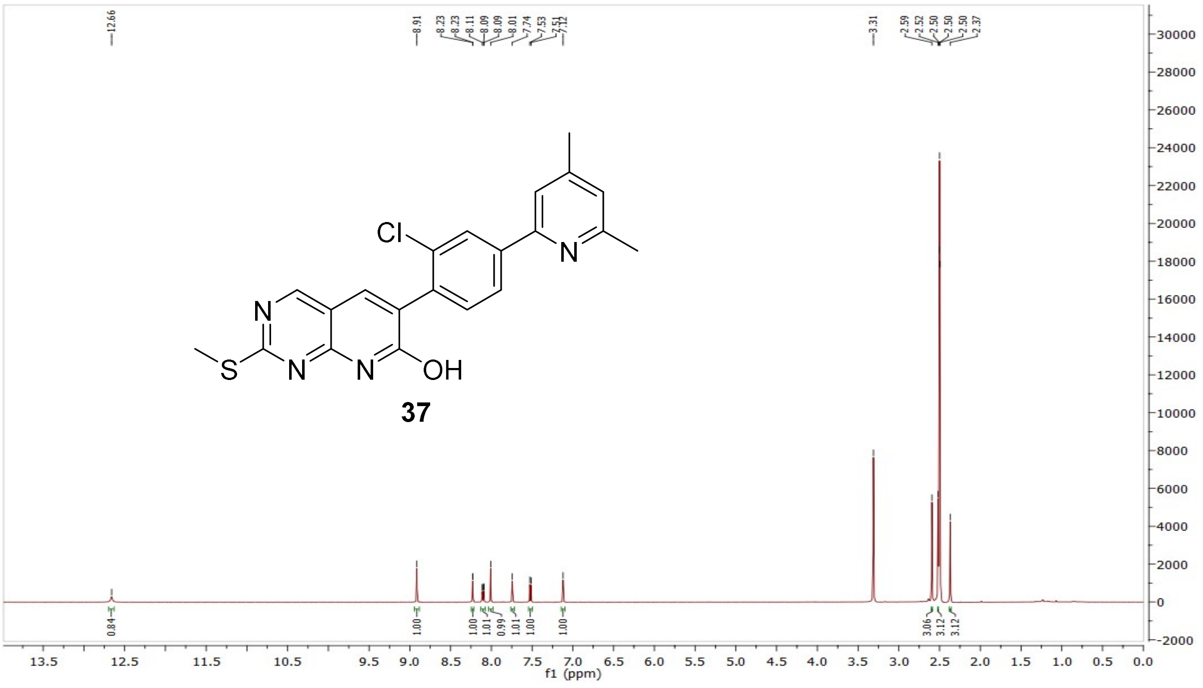

$^{13}C$ NMR compound **38**:

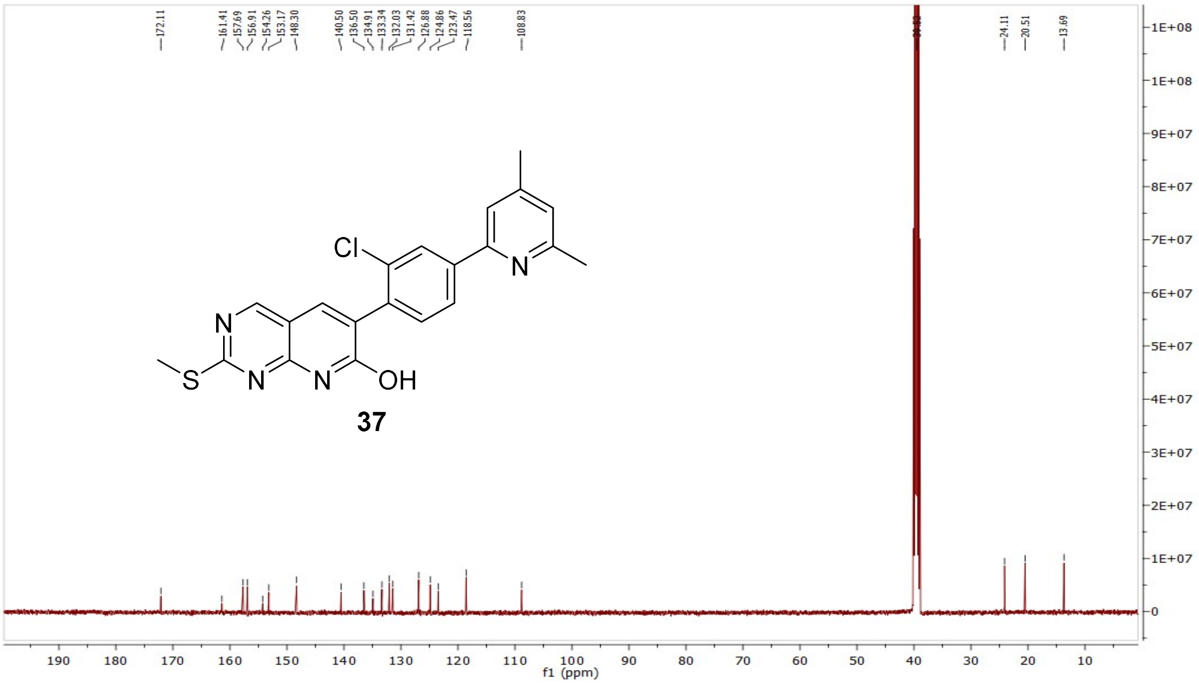


MS (ESI+) compound **38**:

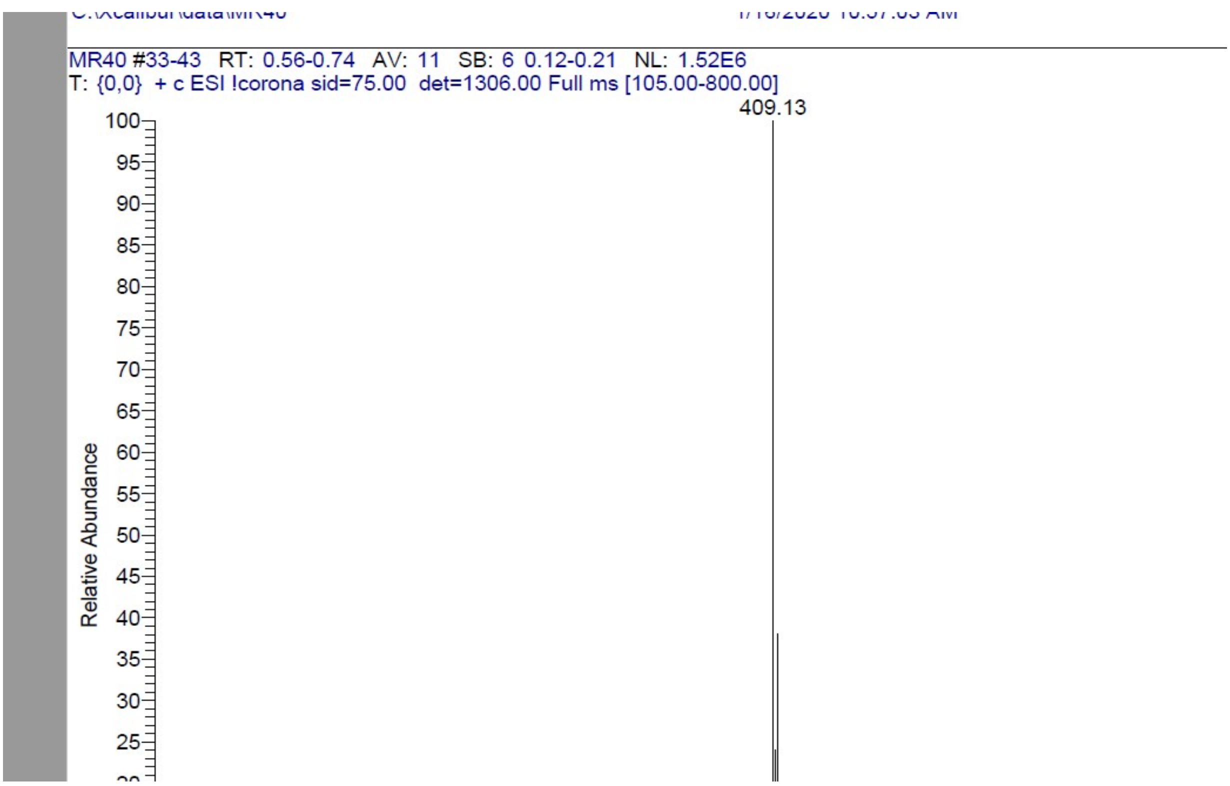

$^{1}H$ NMR compound **7**:

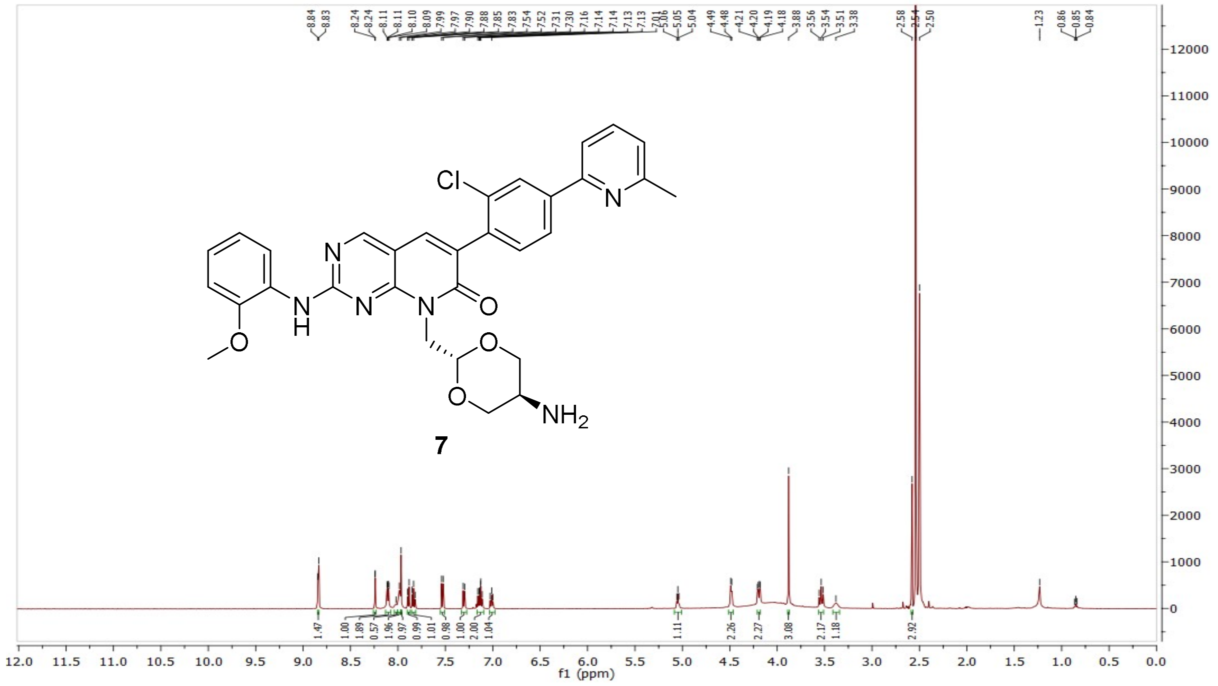

$^{13}C$ NMR compound **7**:

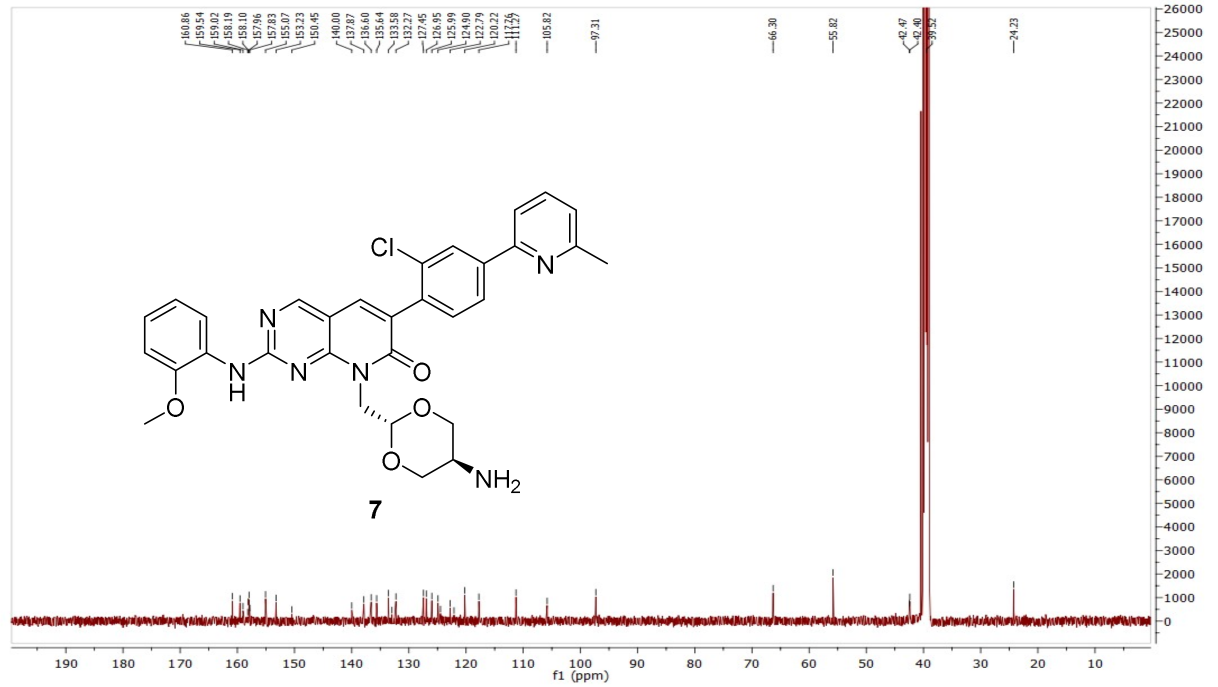

HRMS (FTMS +p MALDI) compound **7**:

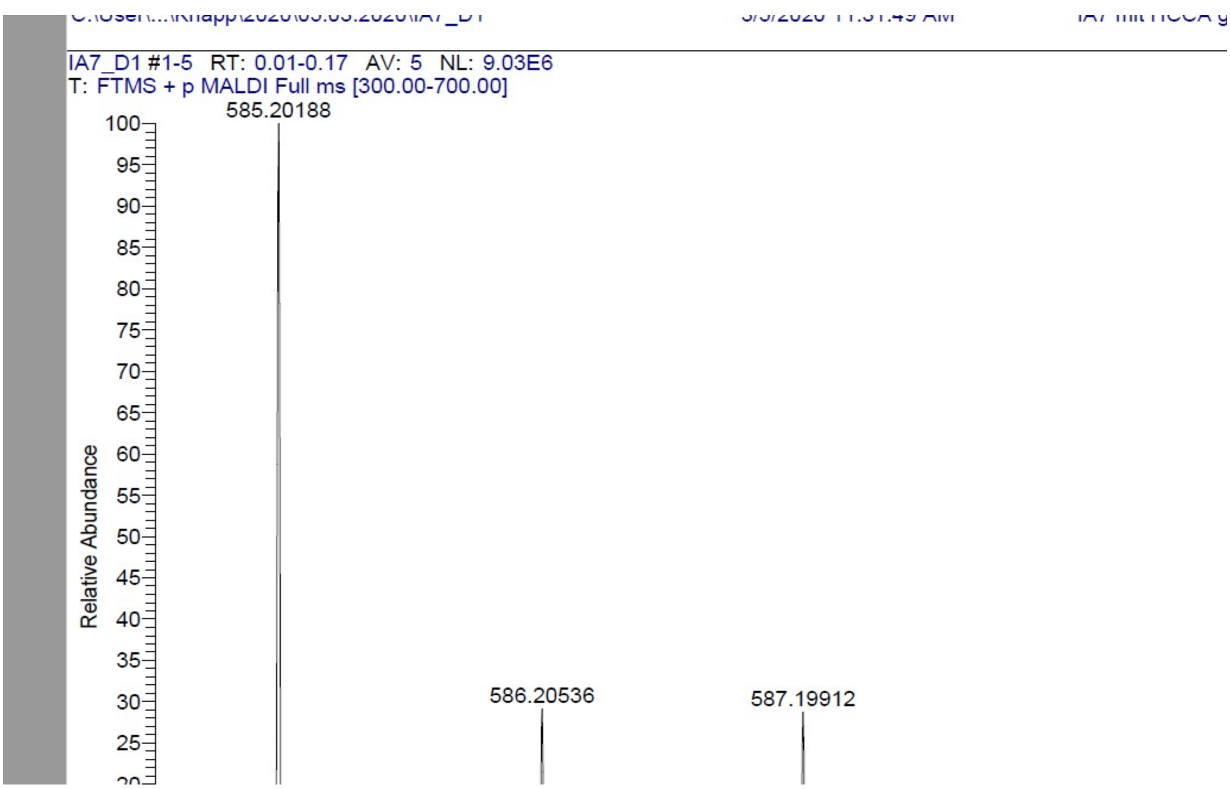

HPLC-chromatograms compound **7**:

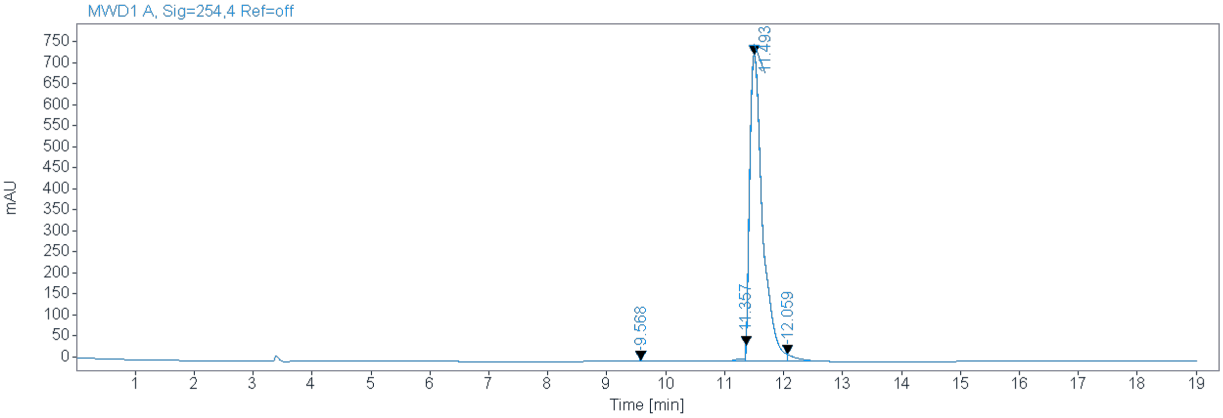


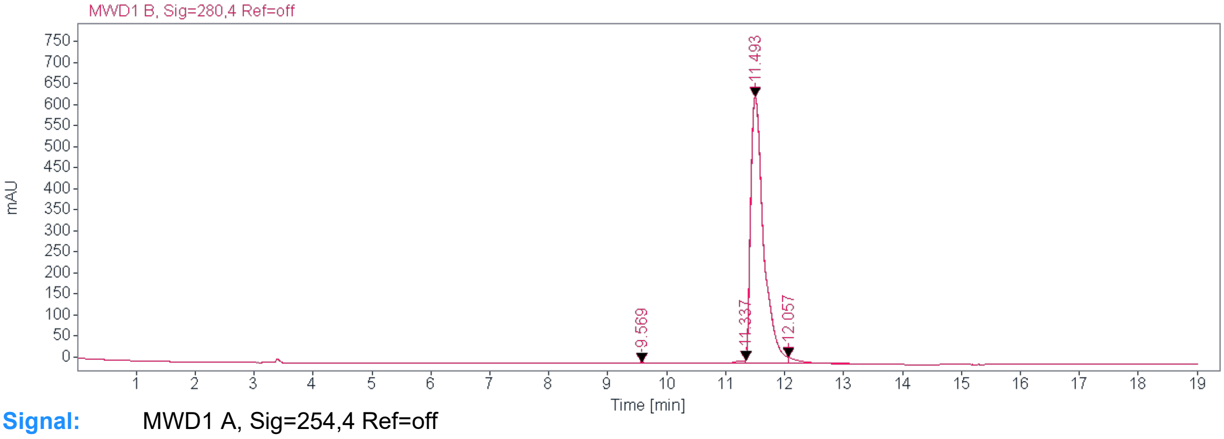


**Signal:** MWD1 A, Sig=254,4 Ref=off

| RT [min] | Type | Width [min] | Area | Height | Area% | Name |
|---|---|---|---|---|---|---|
| 9.568 | MM | 0.8038 | 110.8189 | 2.2979 | 0.9661 | |
| 11.357 | MM | 0.0392 | 89.1243 | 37.8772 | 0.7769 | |
| 11.493 | MM | 0.2533 | 11085.0762 | 729.4694 | 96.6346 | |
| 12.059 | MM | 0.1912 | 186.1067 | 16.2195 | 1.6224 | |
| | | Sum | 11471.1261 | | | |

**Signal:** MWD1 B, Sig=280,4 Ref=off

| RT [min] | Type | Width [min] | Area | Height | Area% | Name |
|---|---|---|---|---|---|---|
| 9.569 | MM | 0.7332 | 109.5441 | 2.4901 | 1.0867 | |
| 11.337 | MM | 0.2213 | 93.9797 | 7.0787 | 0.9323 | |
| 11.493 | MM | 0.2543 | 9672.0059 | 633.9951 | 95.9459 | |
| 12.057 | MM | 0.1576 | 205.1585 | 15.2968 | 2.0352 | |
| | | Sum | 0080.882 | | | |

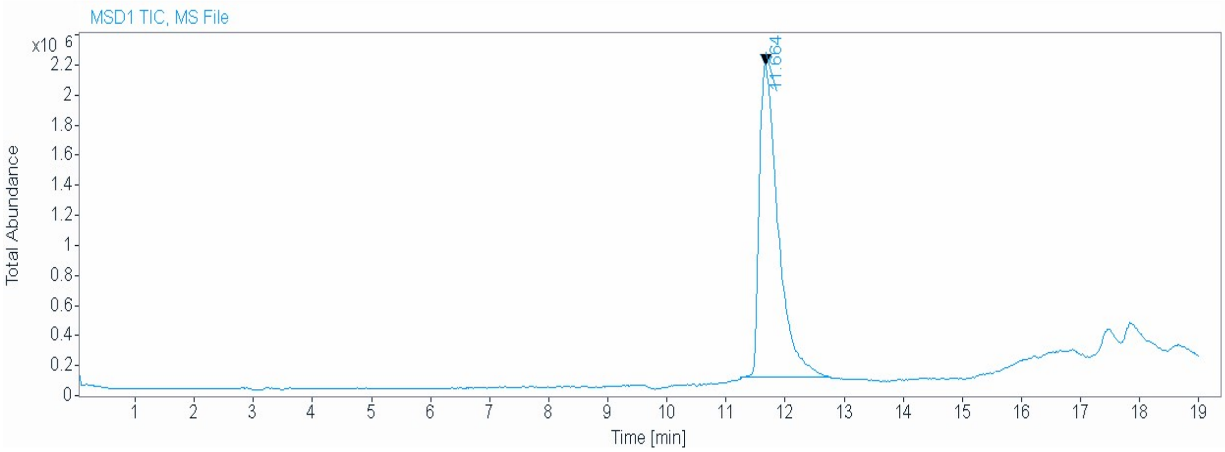


| | |
|---|---|
| Signal | MSD1 TIC, MS File |
| Peak RT | 11.664 |

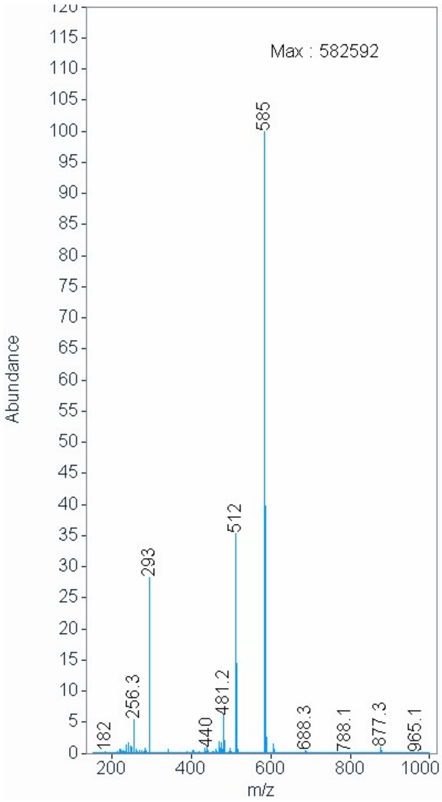

[1]H NMR compound **8**:

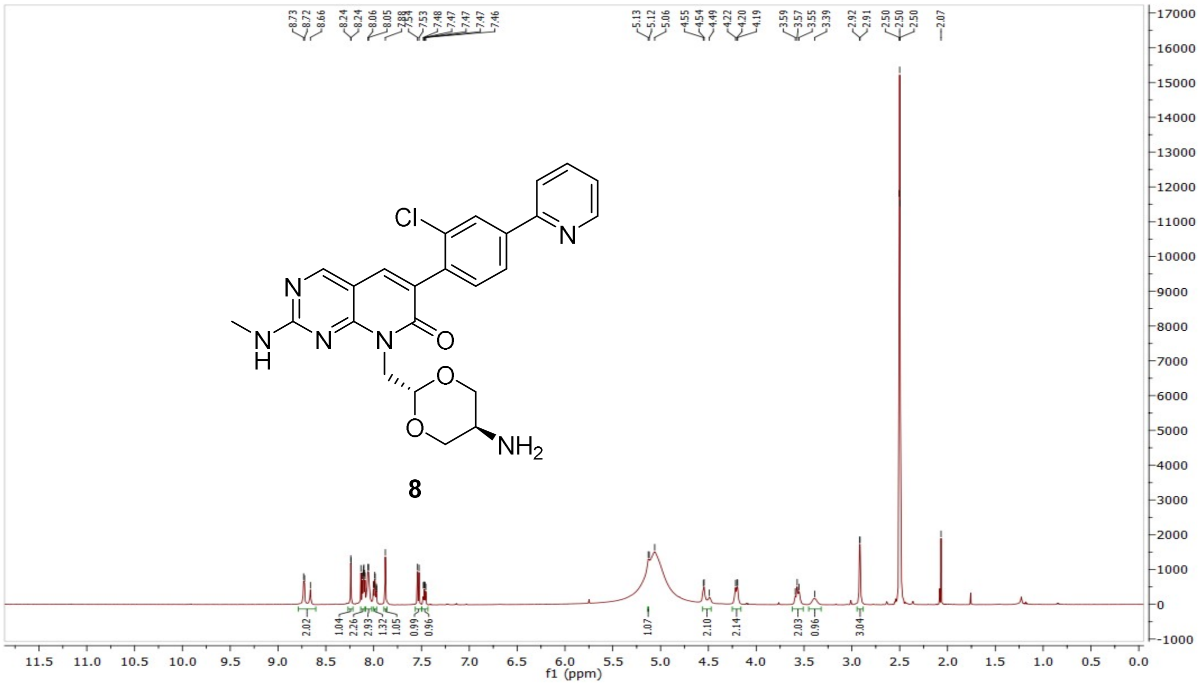


[13]C NMR compound **8**:

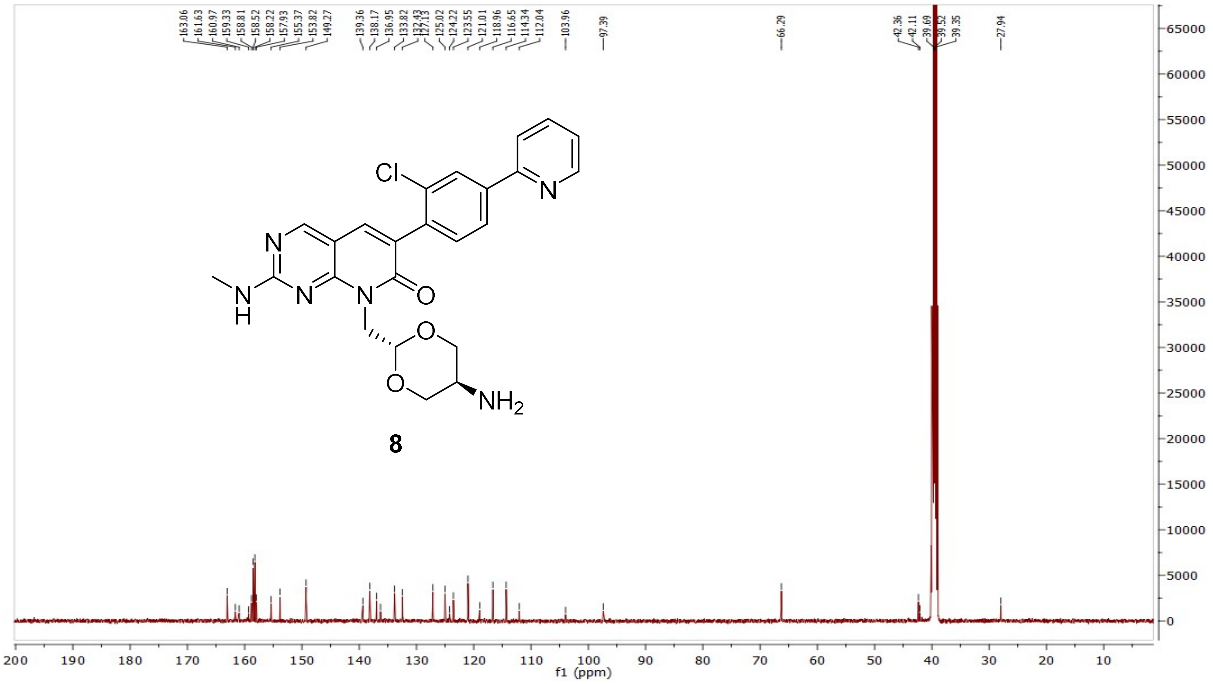

HRMS (FTMS +p MALDI) compound **8**:

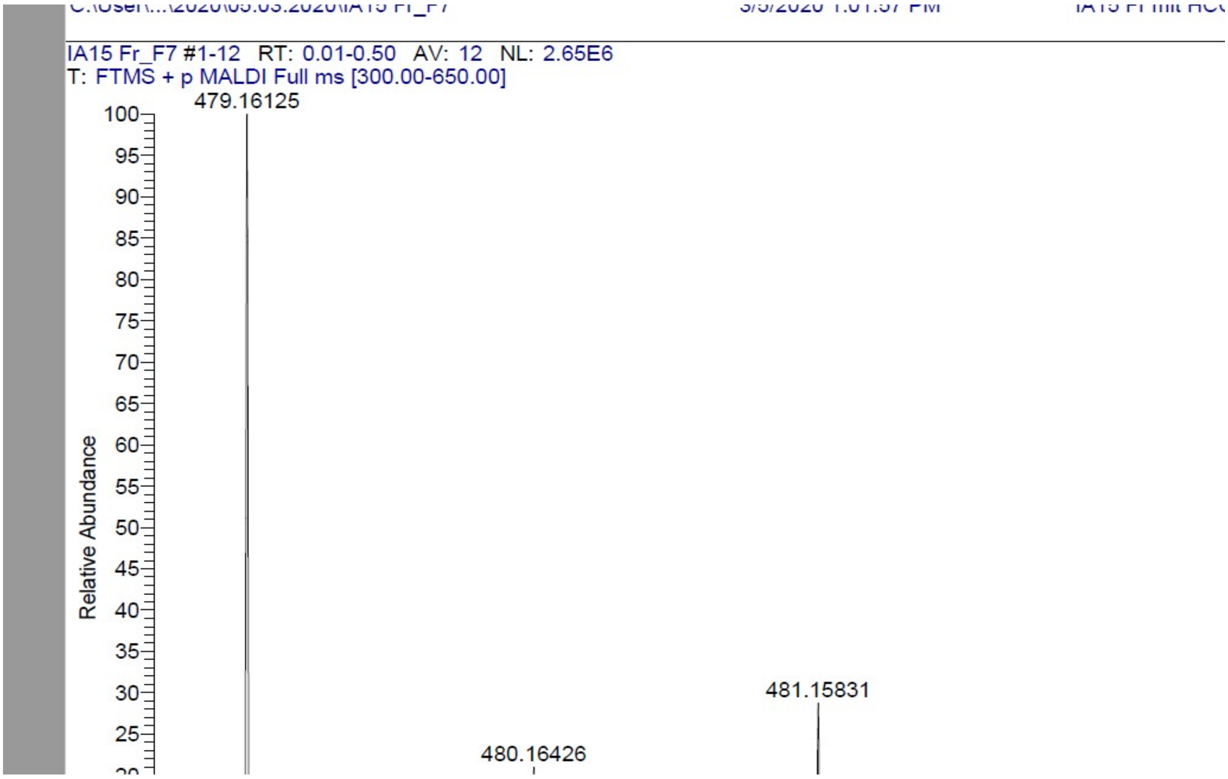

HPLC-chromatograms compound **8**:

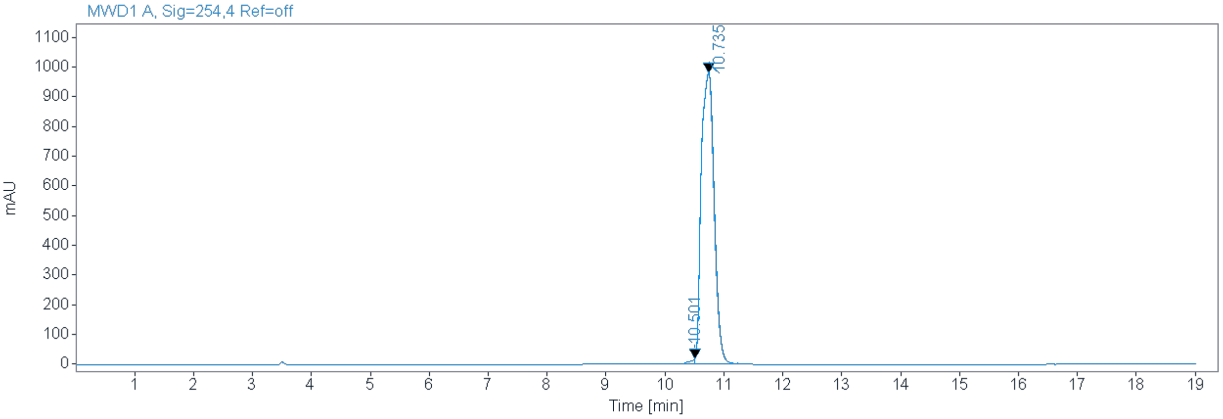


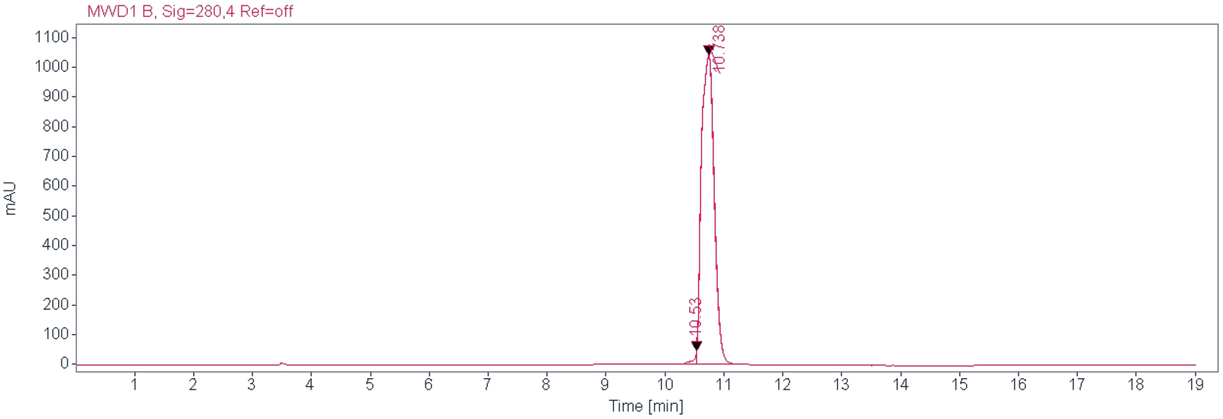


**Signal:** MWD1 A, Sig=254,4 Ref=off

| RT [min] | Type | Width [min] | Area | Height | Area% | Name |
|---|---|---|---|---|---|---|
| 10.501 | MM | 0.1003 | 120.8871 | 20.0778 | 0.8036 | |
| 10.735 | MM | 0.2533 | 14922.7129 | 981.7443 | 99.1964 | |
| | | Sum | 15043.5999 | | | |

**Signal:** MWD1 B, Sig=280,4 Ref=off

| RT [min] | Type | Width [min] | Area | Height | Area% | Name |
|---|---|---|---|---|---|---|
| 10.530 | MM | 0.0641 | 177.3867 | 46.1568 | 1.1163 | |
| 10.738 | MM | 0.2515 | 15713.6553 | 1041.1525 | 98.8837 | |
| | | Sum | 15891.0419 | | | |

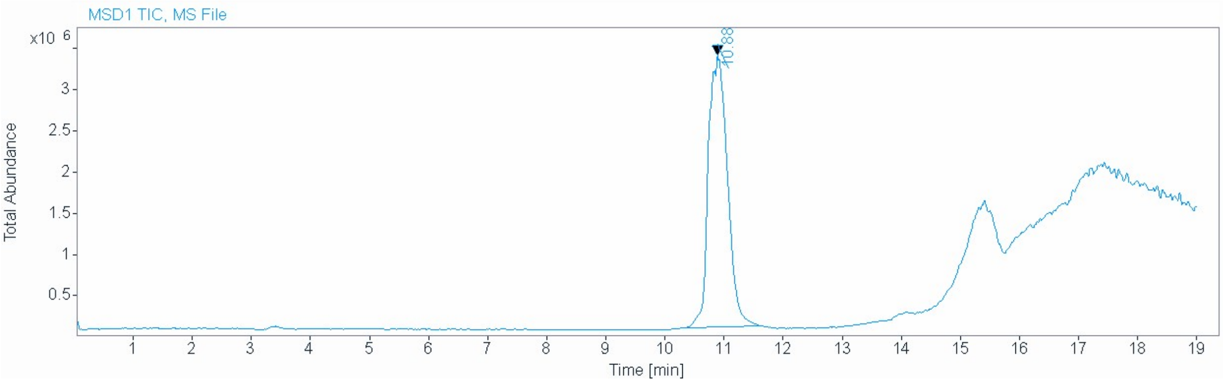


**Signal** MSD1 TIC, MS File

**Peak RT** 10.880

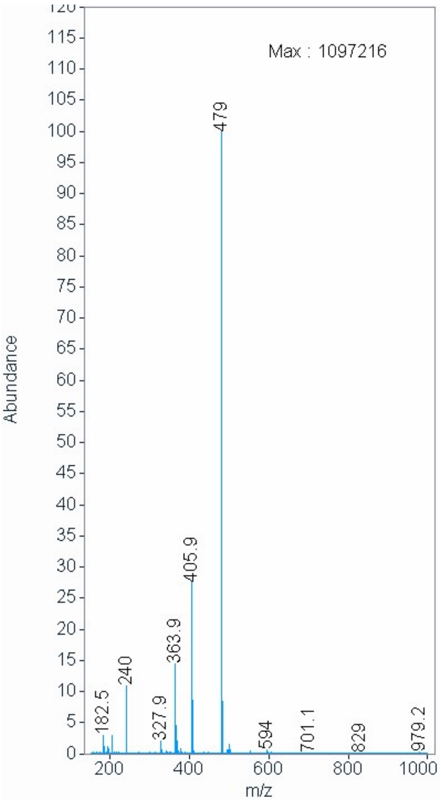

$^{1}H$ NMR compound **9**:

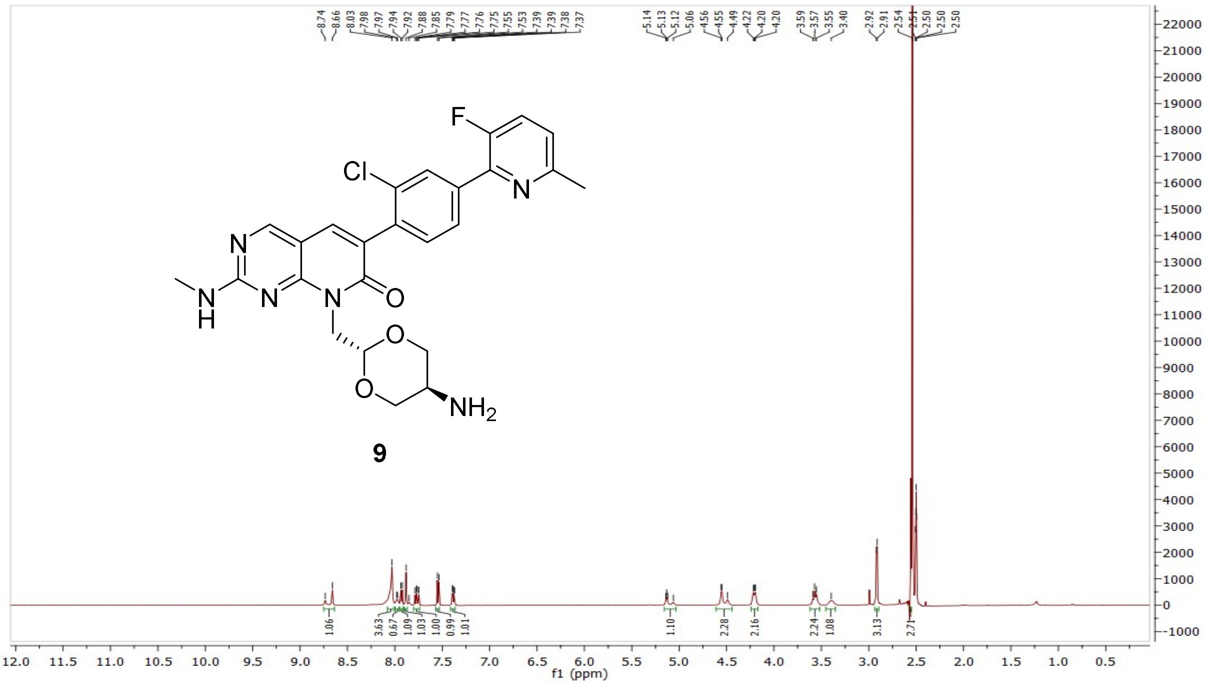


$^{13}C$ NMR compound **9**:

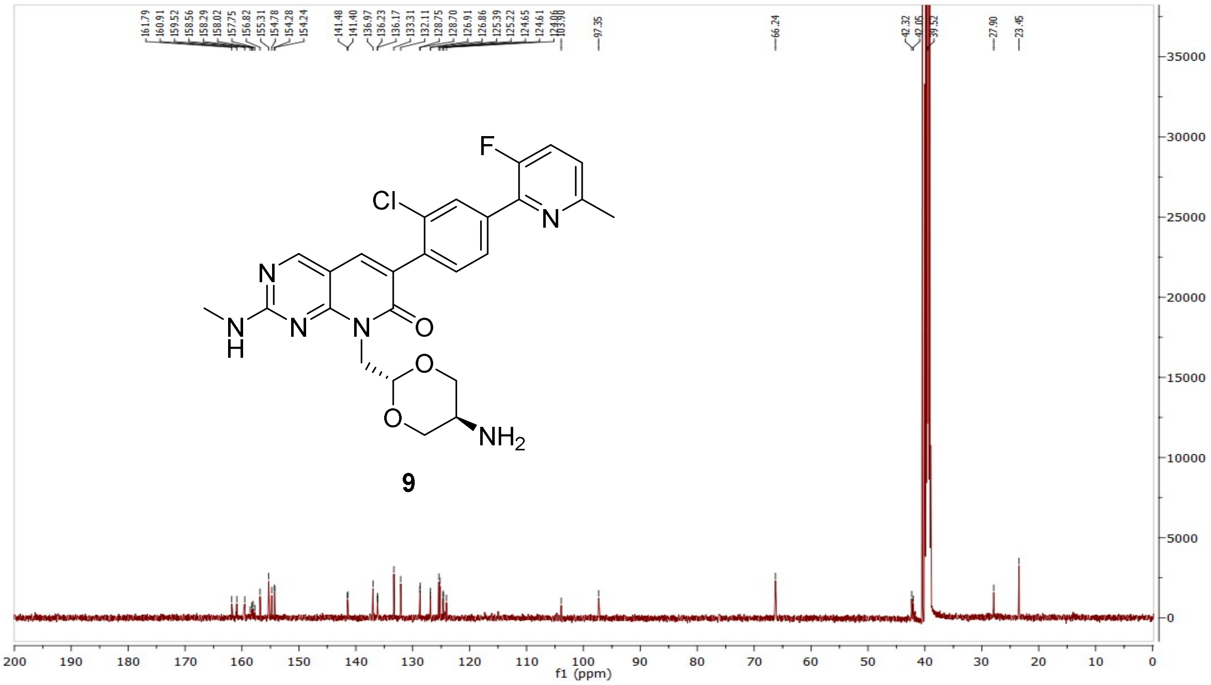

HRMS (FTMS +p MALDI) compound **9**:

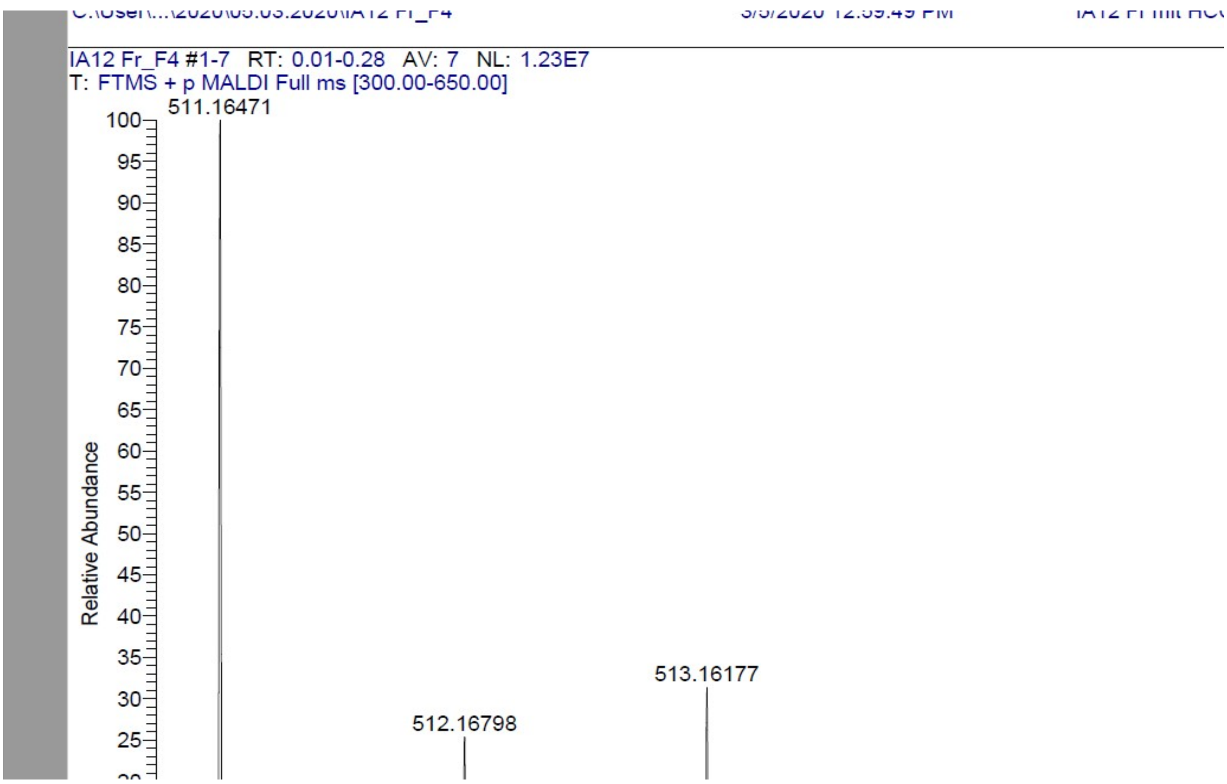

HPLC-chromatograms compound **9**:

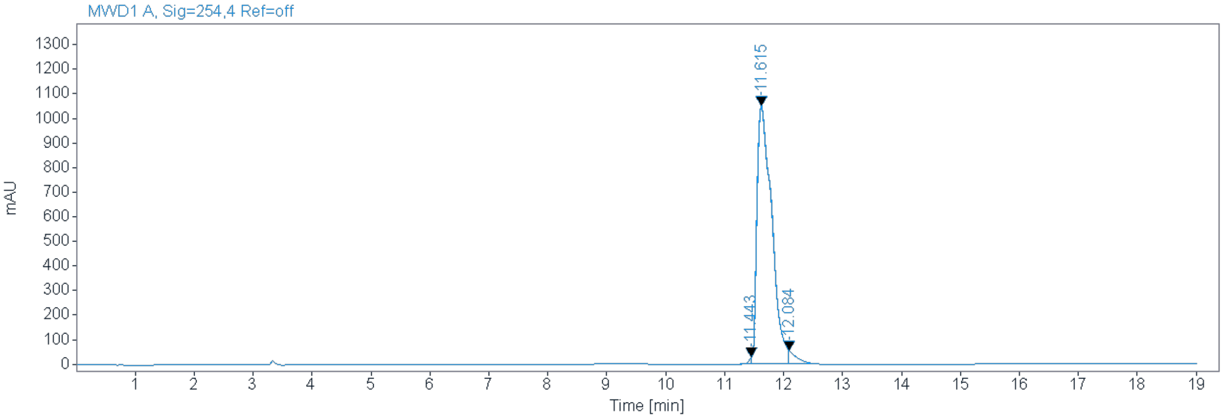


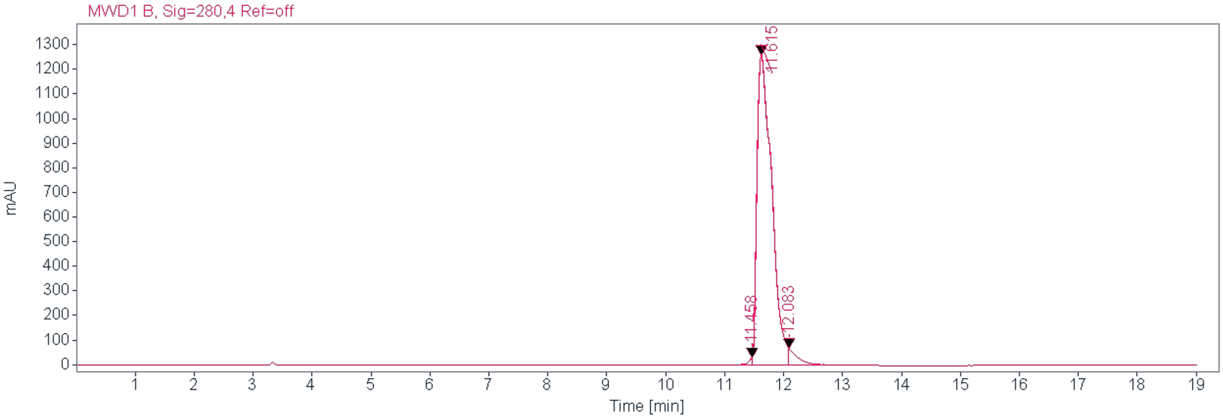


**Signal:** MWD1 B, Sig=280,4 Ref=off

| RT [min] Type | Width [min] | Area | Height | Area% Name |
|---|---|---|---|---|
| 11.443 MM | 0.0502 | 85.6657 | 28.4422 | 0.4551 |
| 11.615 MM | 0.2890 | 18216.6621 | 1050.6274 | 96.7666 |
| 12.084 MM | 0.1218 | 523.0377 | 55.6911 | 2.7784 |
| | Sum | 18825.3655 | | |

**Signal:** MWD1 B, Sig=280,4 Ref=off

| RT [min] Type | Width [min] | Area | Height | Area% Name |
|---|---|---|---|---|
| 11.458 MM | 0.0683 | 123.5502 | 30.1602 | 0.5462 |
| 11.615 MM | 0.2885 | 21807.9980 | 1259.8616 | 96.4134 |
| 12.083 MM | 0.1291 | 687.7219 | 68.4209 | 3.0404 |
| | Sum | 22619.2702 | | |

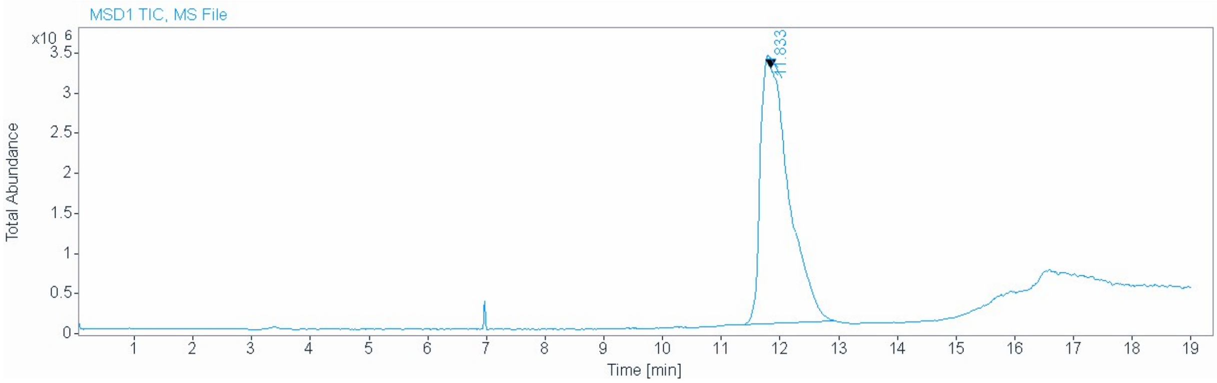


**Signal** MSD1 TIC, MS File

**Peak RT** 11.833

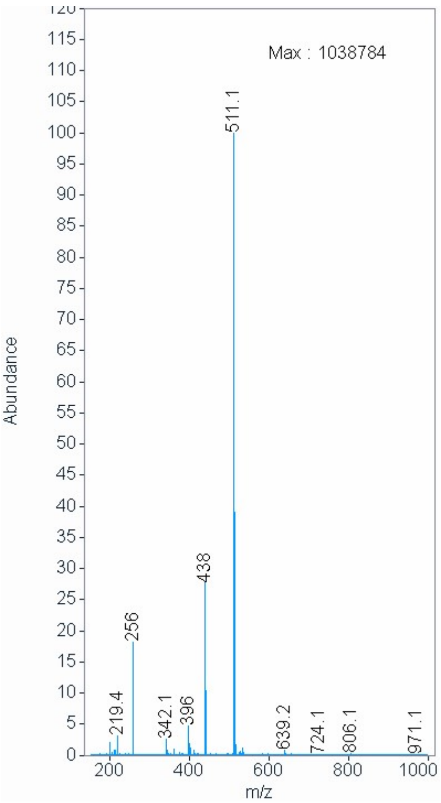

[1]H NMR compound **10**:

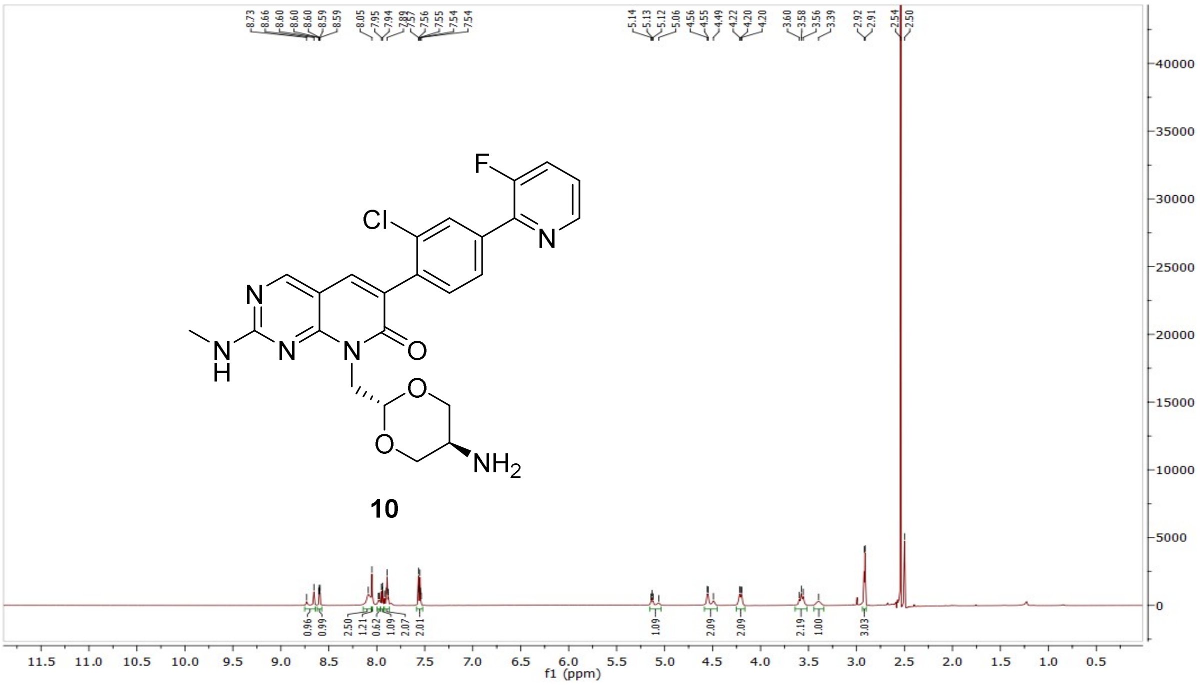


[13]C NMR compound **10**:

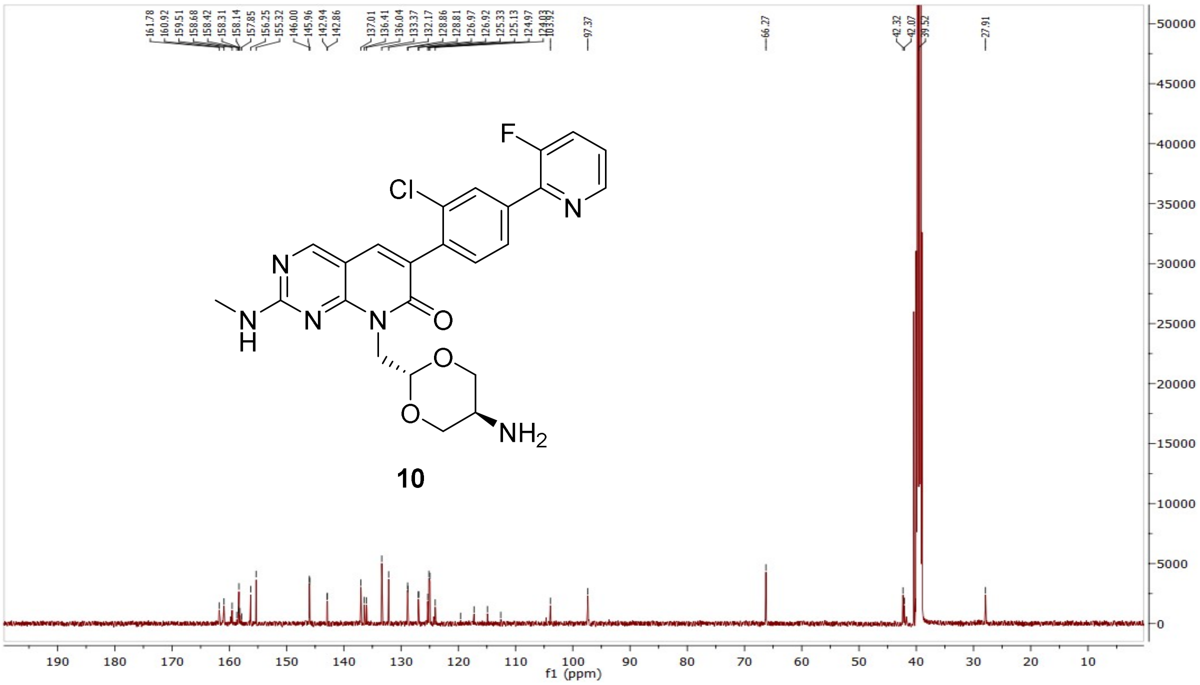

HRMS (FTMS +p MALDI) compound **10**:

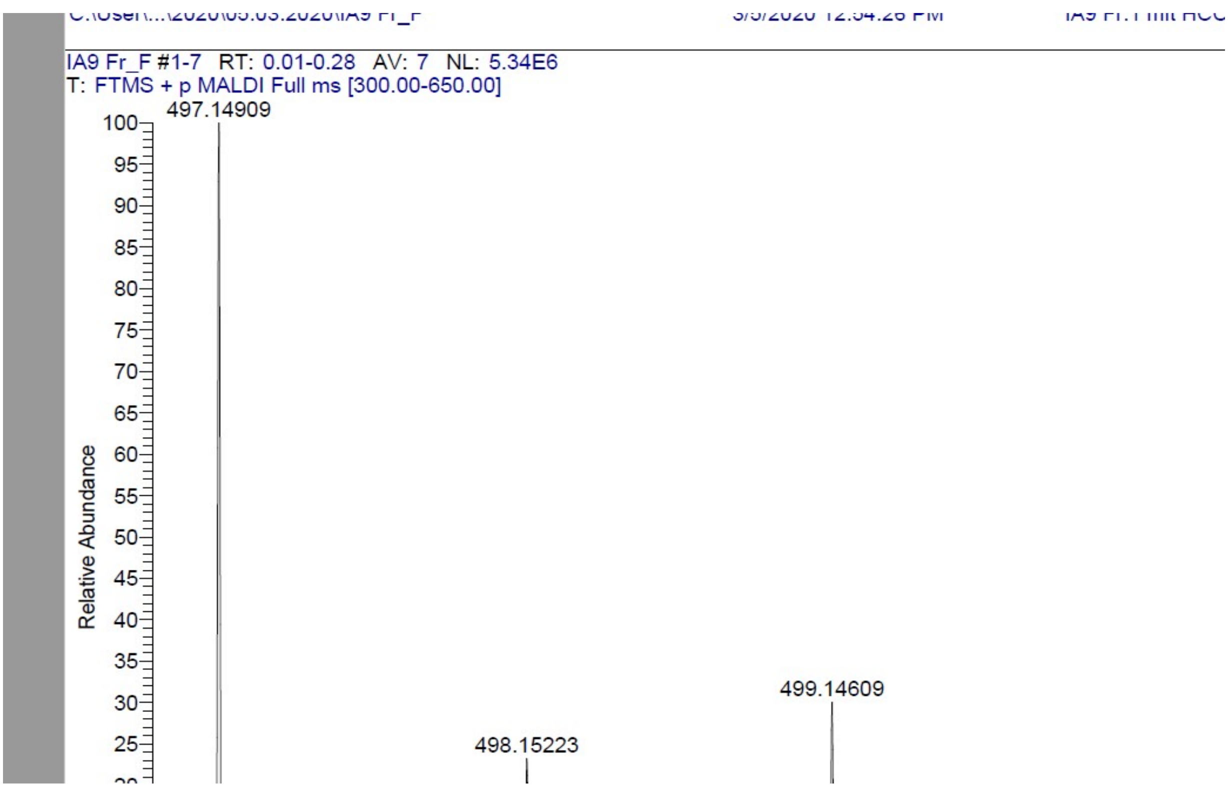

HPLC-chromatograms compound **10**:

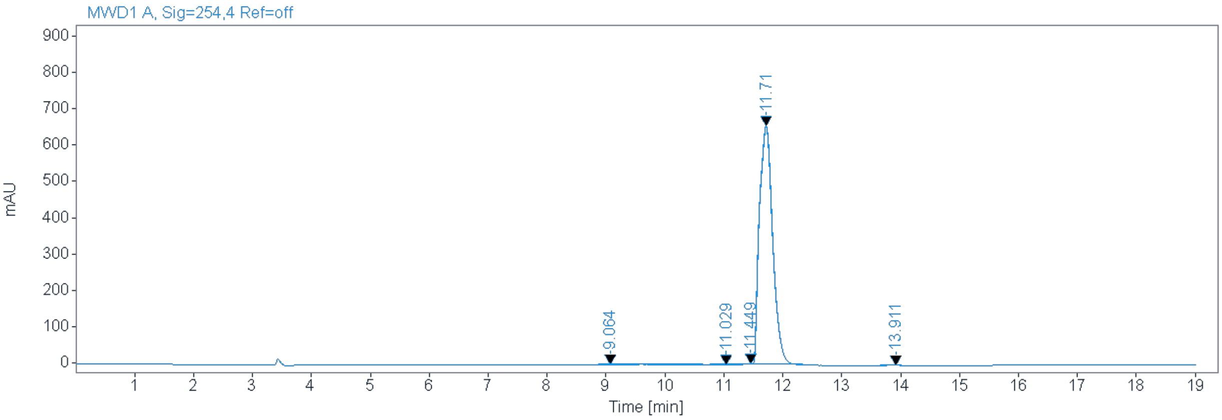


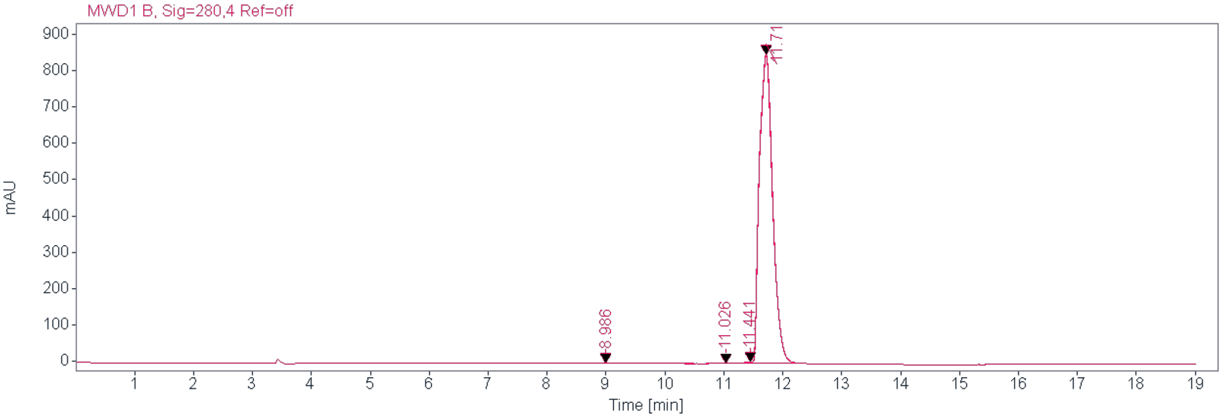


**Signal:** MWD1 B, Sig=254,4 Ref=off

| RT [min] | Type | Width [min] | Area | Height | Area% | Name |
|---|---|---|---|---|---|---|
| 9.064 | MM | 1.0205 | 122.5904 | 2.0021 | 1.1764 | |
| 11.029 | MM | 0.2613 | 7.2988 | 0.4655 | 0.0700 | |
| 11.449 | MM | 0.1052 | 21.5046 | 3.4072 | 0.2064 | |
| 11.710 | MM | 0.2597 | 10239.5293 | 657.1000 | 98.2597 | |
| 13.911 | MM | 0.2819 | 29.9574 | 1.7710 | 0.2875 | |
| | | Sum | 10420.8806 | | | |

**Signal:** MWD1 B, Sig=280,4 Ref=off

| RT [min] | Type | Width [min] | Area | Height | Area% | Name |
|---|---|---|---|---|---|---|
| 8.986 | MM | 0.9499 | 74.7159 | 1.3109 | 0.5594 | |
| 11.026 | MM | 0.2749 | 7.0301 | 0.4262 | 0.0526 | |
| 11.441 | MM | 0.1003 | 27.4708 | 4.5632 | 0.2057 | |
| 11.710 | MM | 0.2600 | 13247.0850 | 849.1813 | 99.1823 | |
| | | Sum | 3356.3018 | | | |

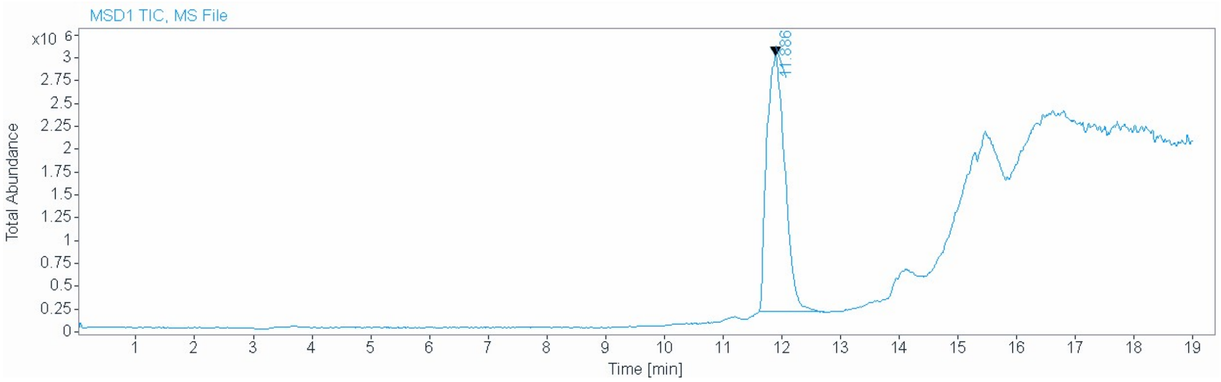


**Signal** MSD1 TIC, MS File

**Peak RT** 11.886

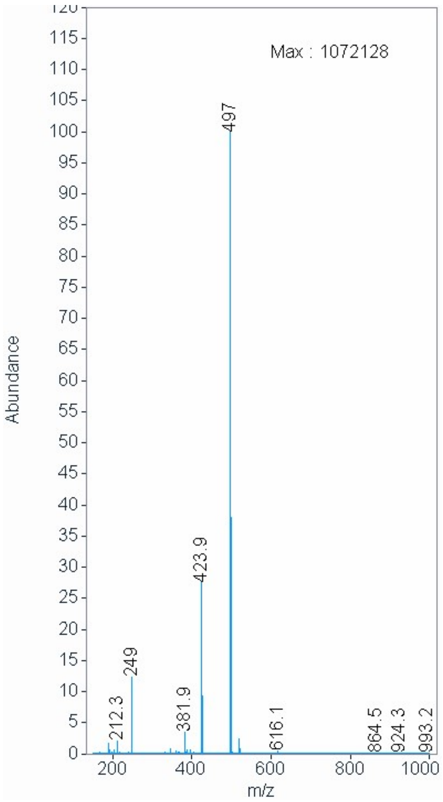

$^{1}H$ NMR compound **11**:

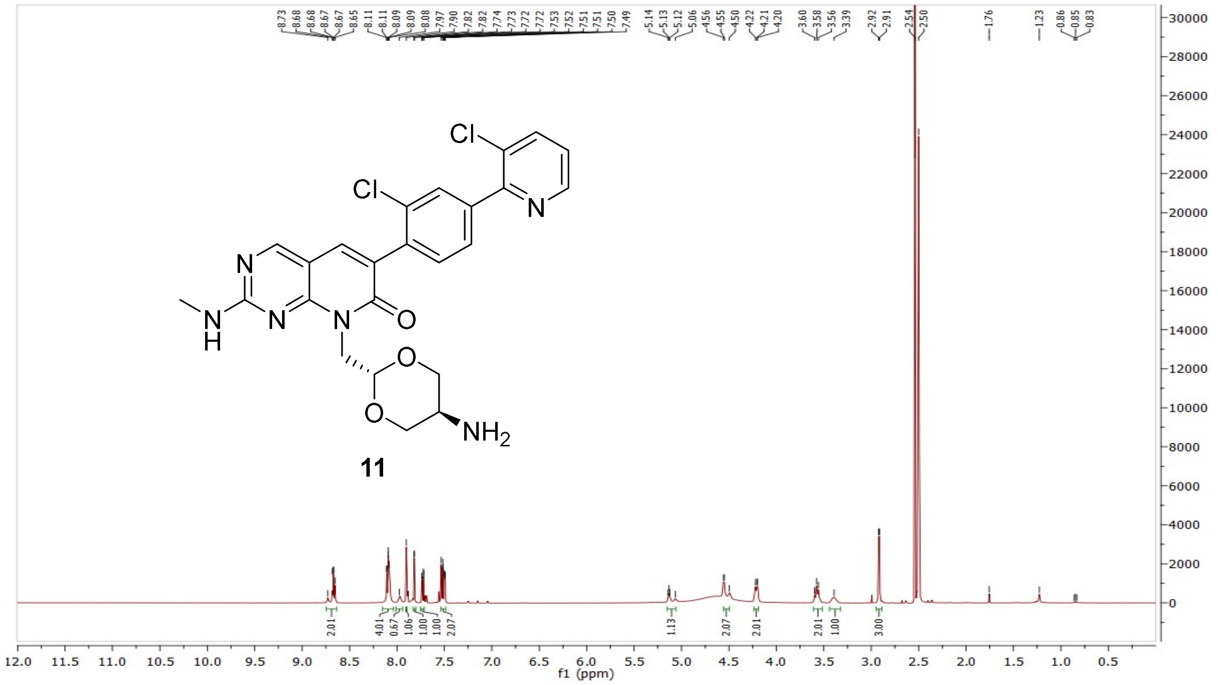

$^{13}C$ NMR compound **11**:

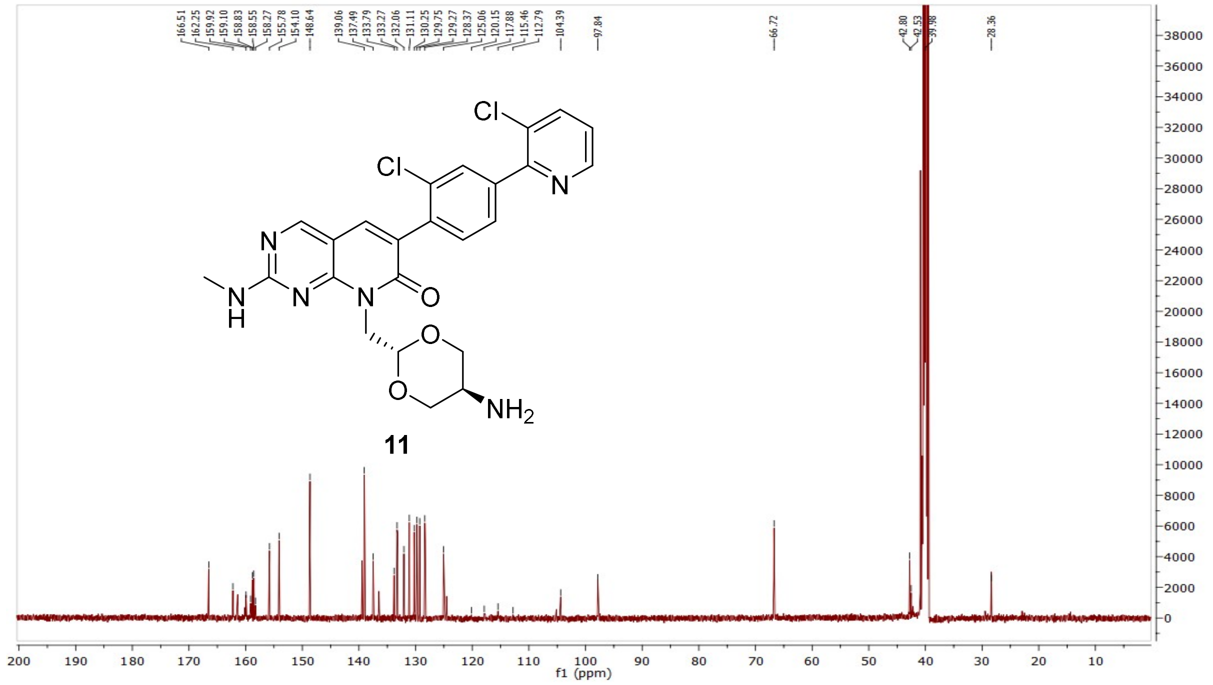

HRMS (FTMS +p MALDI) compound **11**:

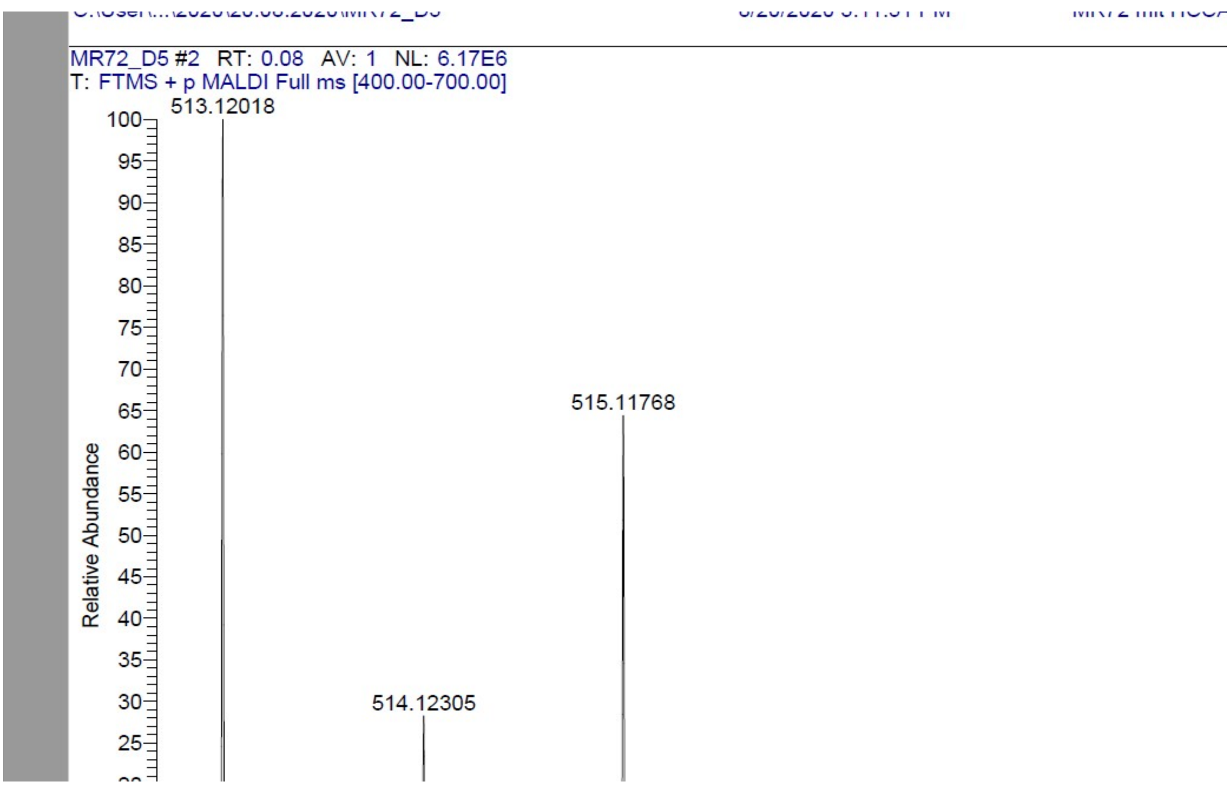

HPLC-chromatograms compound **11**:

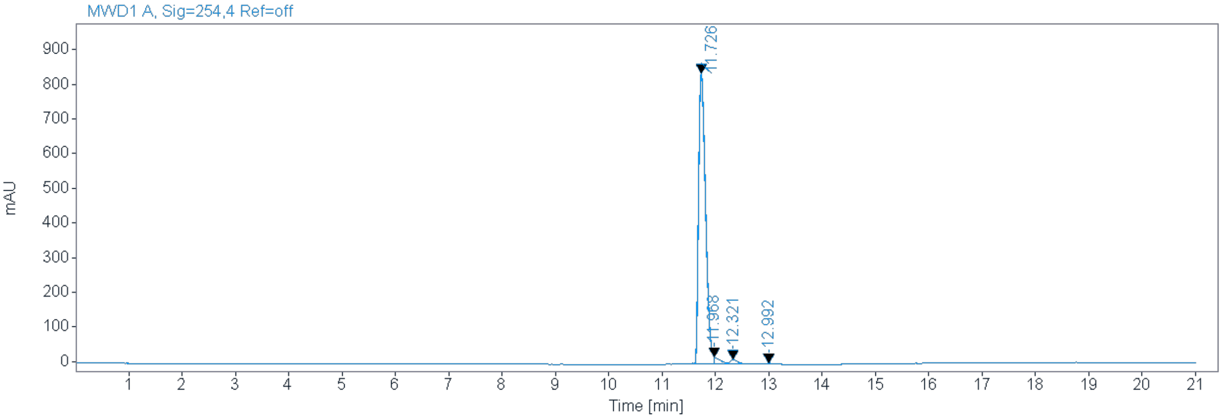


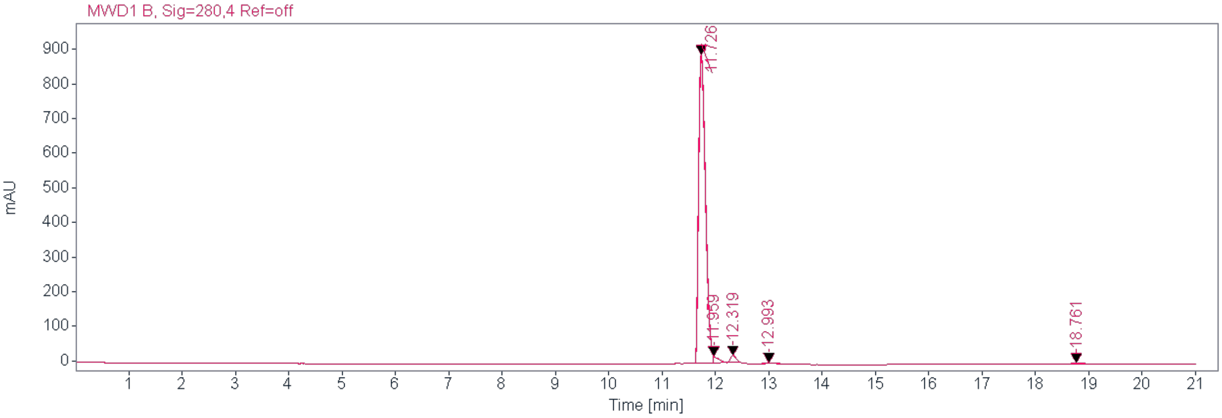


**Signal:** MWD1 A, Sig=254,4 Ref=off

| RT [min] | Type | Width [min] | Area | Height | Area% | Name |
|---|---|---|---|---|---|---|
| 11.726 | MM | 0.1512 | 7596.8838 | 837.4711 | 96.2666 | |
| 11.968 | MM | 0.0944 | 137.5973 | 18.7323 | 1.7436 | |
| 12.321 | MM | 0.1568 | 114.3046 | 12.1498 | 1.4485 | |
| 12.992 | MM | 0.2028 | 42.7236 | 3.5116 | 0.5414 | |
| | | Sum | 7891.5093 | | | |

**Signal:** MWD1 B, Sig=280,4 Ref=off

| RT [min] | Type | Width [min] | Area | Height | Area% | Name |
|---|---|---|---|---|---|---|
| 11.726 | MM | 0.1506 | 8061.8169 | 892.2730 | 95.0987 | |
| 11.959 | MM | 0.0843 | 136.8278 | 21.4267 | 1.6140 | |
| 12.319 | MM | 0.1287 | 165.3623 | 21.4181 | 1.9506 | |
| 12.993 | MM | 0.1497 | 38.6249 | 4.2991 | 0.4556 | |
| 18.761 | MM | 0.2741 | 74.6817 | 4.5410 | 0.8810 | |
| | | Sum | 8477.3136 | | | |

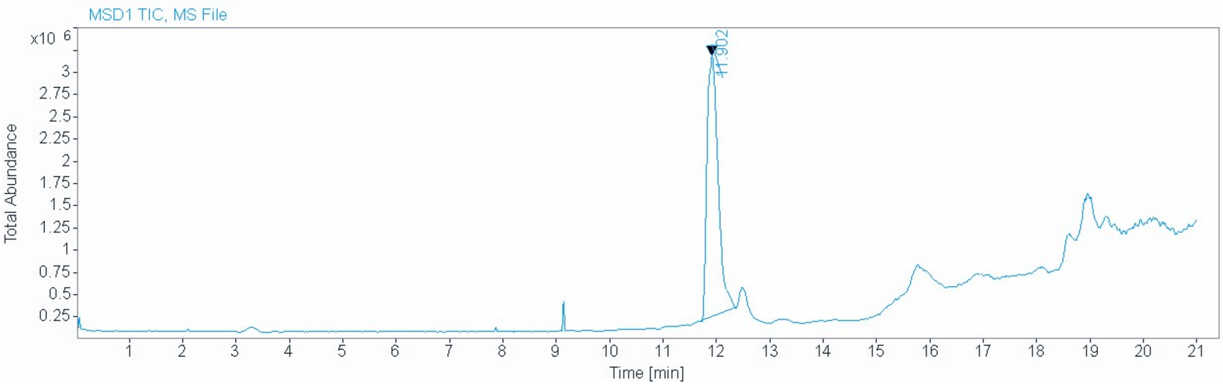


| | |
|---|---|
| **Signal** | MSD1 TIC, MS File |
| **Peak RT** | 11.902 |

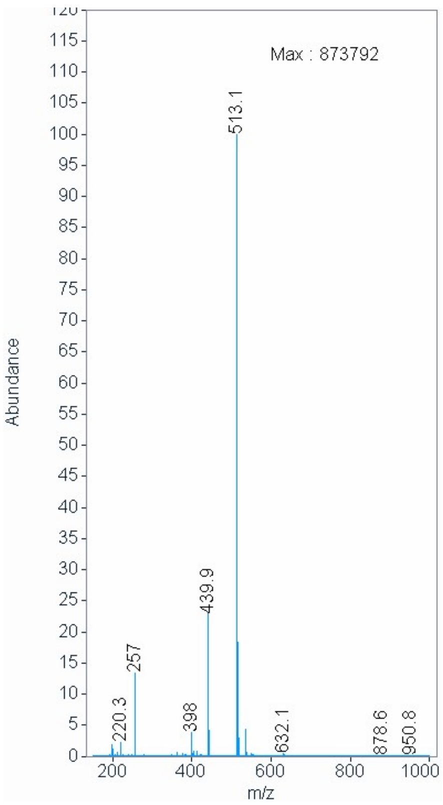

[1]H NMR compound **12**:

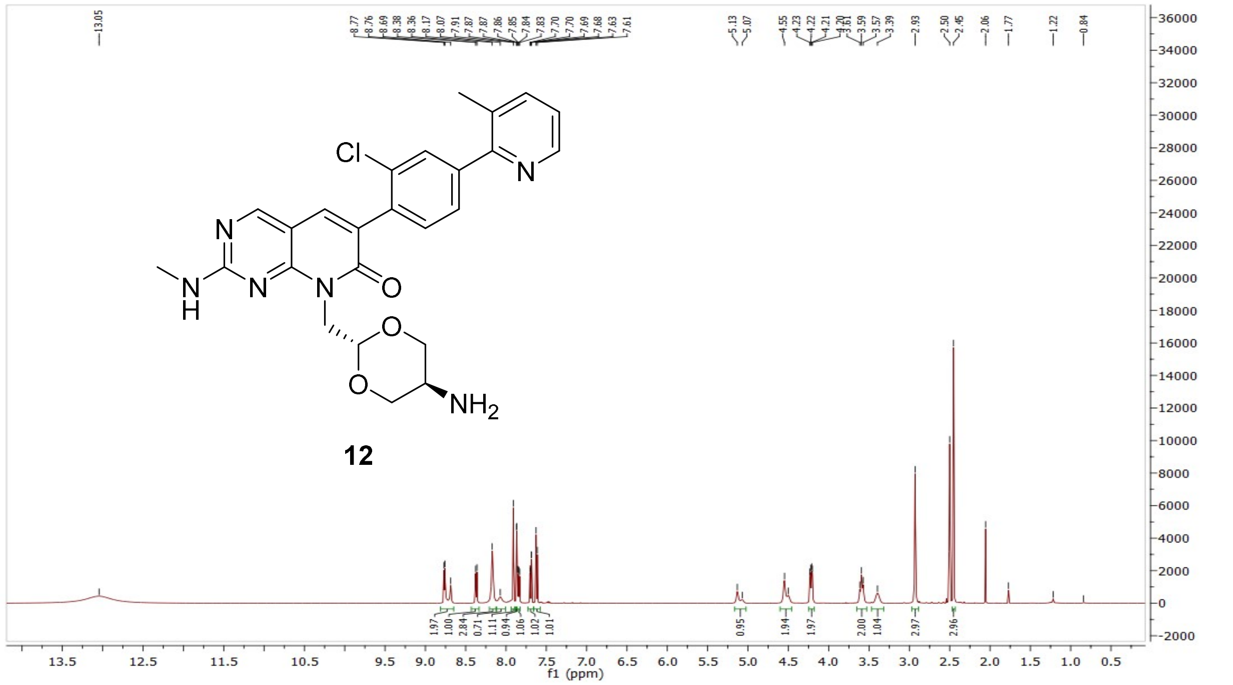

[13]C NMR compound **12**:

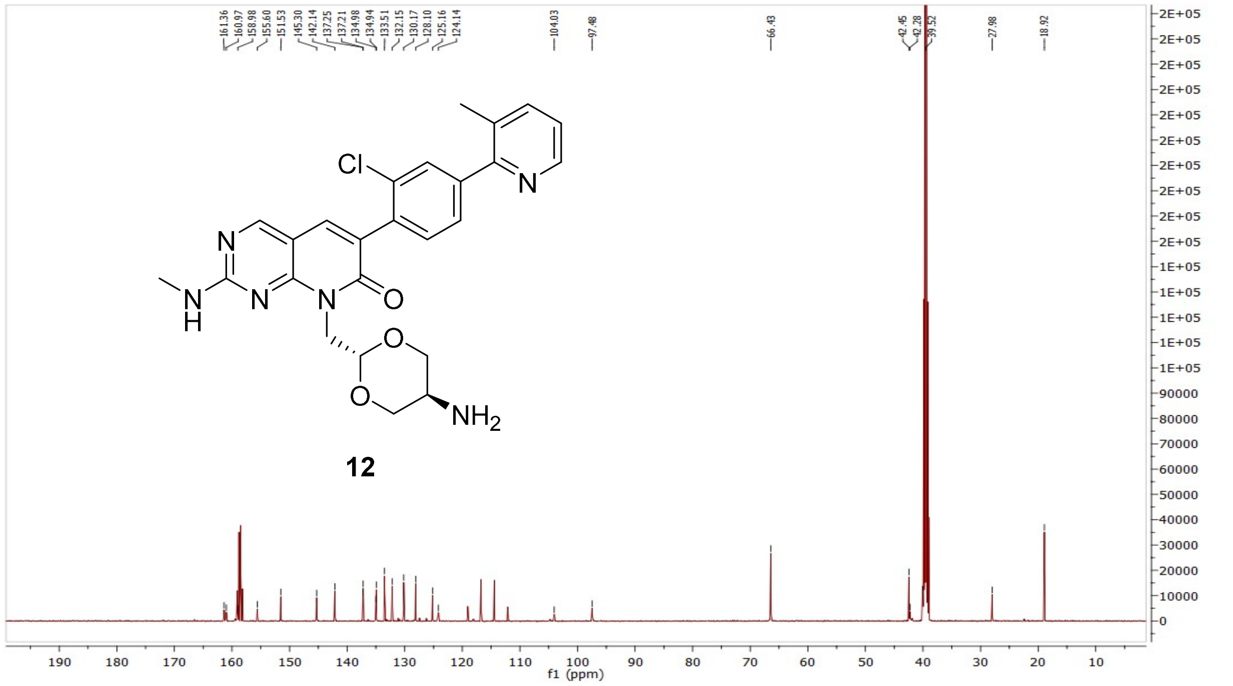

HRMS (FTMS +p MALDI) compound **12**:

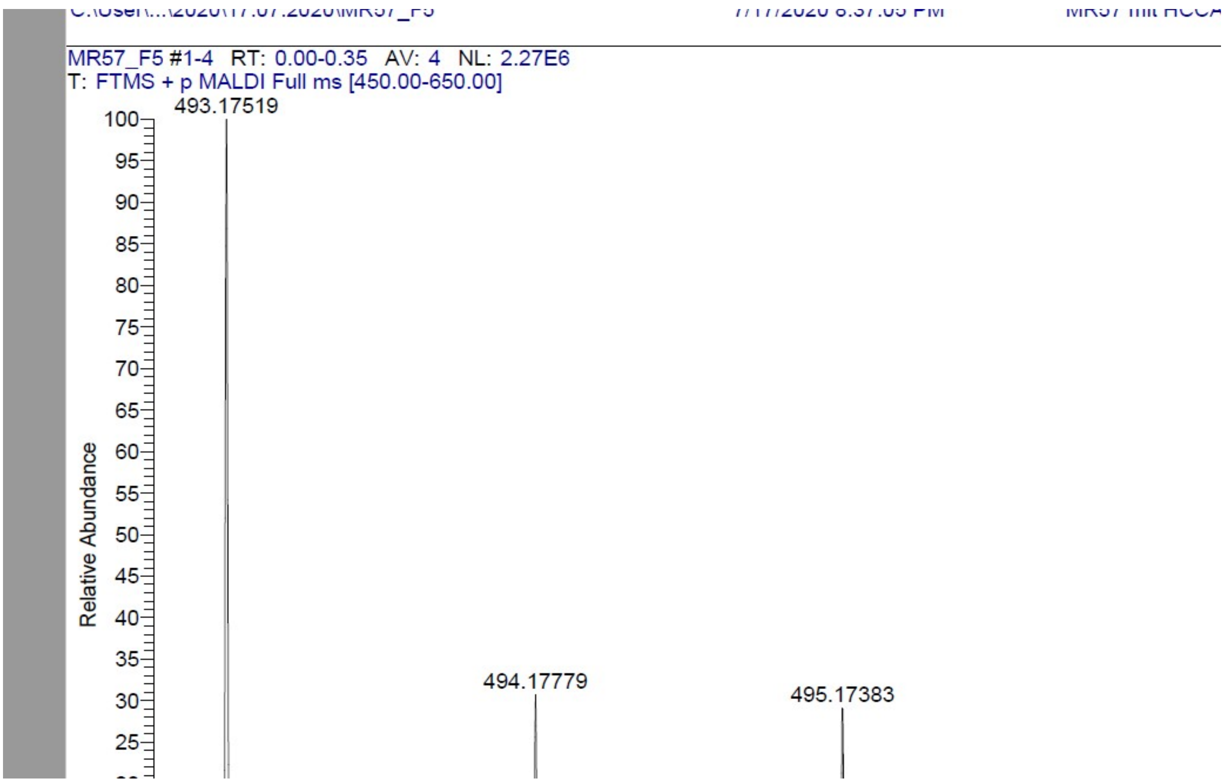

HPLC-chromatograms compound **12**:

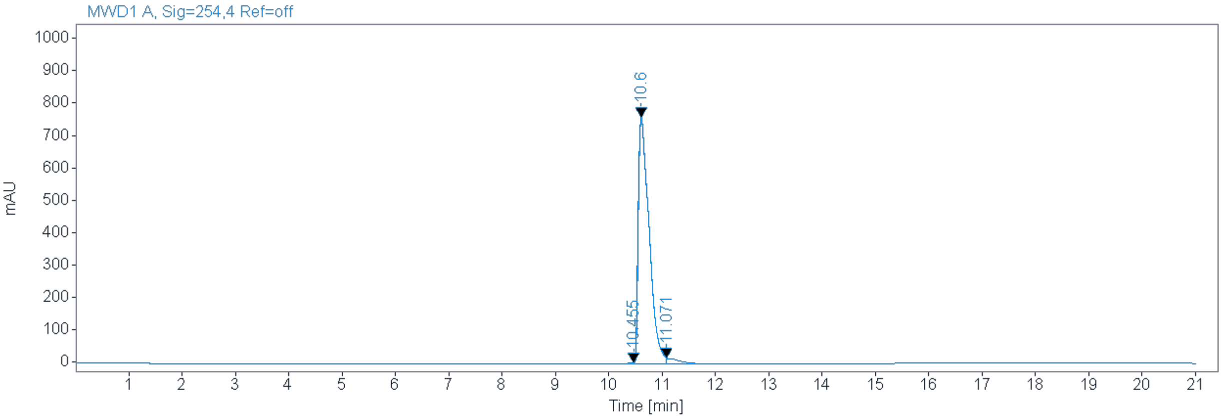


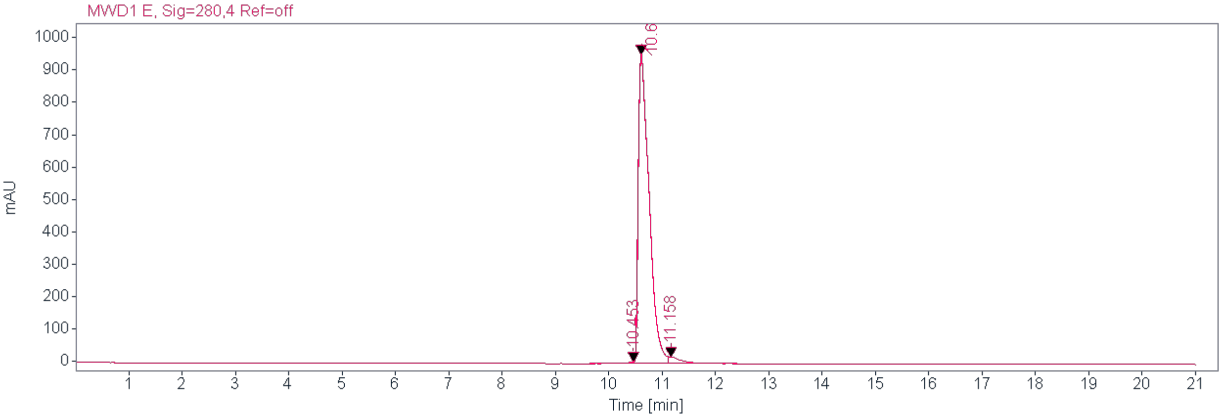


**Signal:** MWD1 A, Sig=254,4 Ref=off

| RT [min] | Type | Width [min] | Area | Height | Area% | Name |
|---|---|---|---|---|---|---|
| 10.455 | MM | 0.1863 | 51.0844 | 4.5689 | 0.4567 | |
| 10.600 | MM | 0.2358 | 10794.9180 | 762.9228 | 96.5067 | |
| 11.071 | MM | 0.2201 | 339.6660 | 19.0116 | 3.0366 | |
| | | Sum | 11185.6684 | | | |

**Signal:** MWD1 E, Sig=280,4 Ref=off

| RT [min] | Type | Width [min] | Area | Height | Area% | Name |
|---|---|---|---|---|---|---|
| 10.453 | MM | 0.2230 | 75.3987 | 5.6364 | 0.5394 | |
| 10.600 | MM | 0.2364 | 13542.0771 | 954.7060 | 96.8774 | |
| 11.158 | MM | 0.2917 | 361.0904 | 20.6307 | 2.5832 | |
| | | Sum | 13978.5663 | | | |

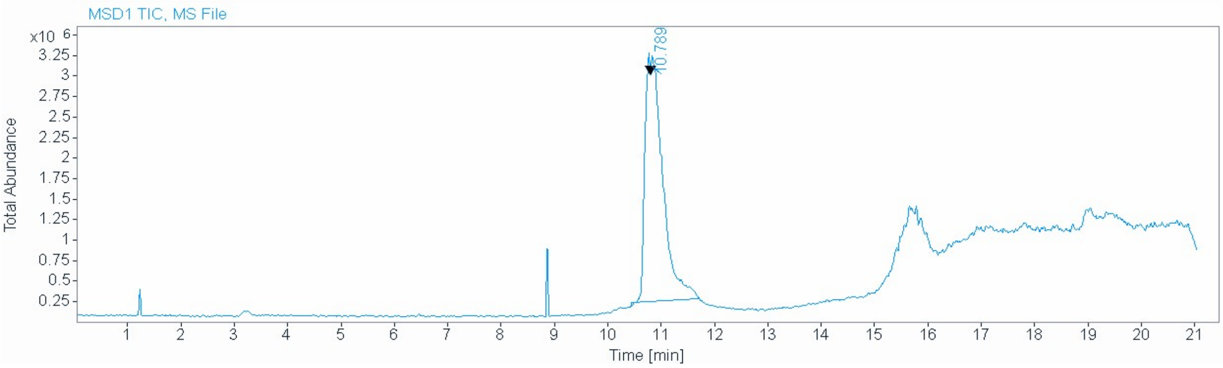


**Signal** MSD1 TIC, MS File

**Peak RT** 10.789

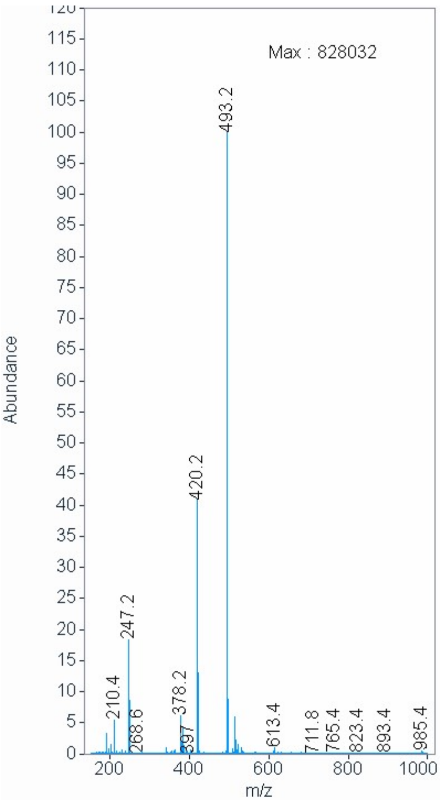

[1]H NMR compound **13**:

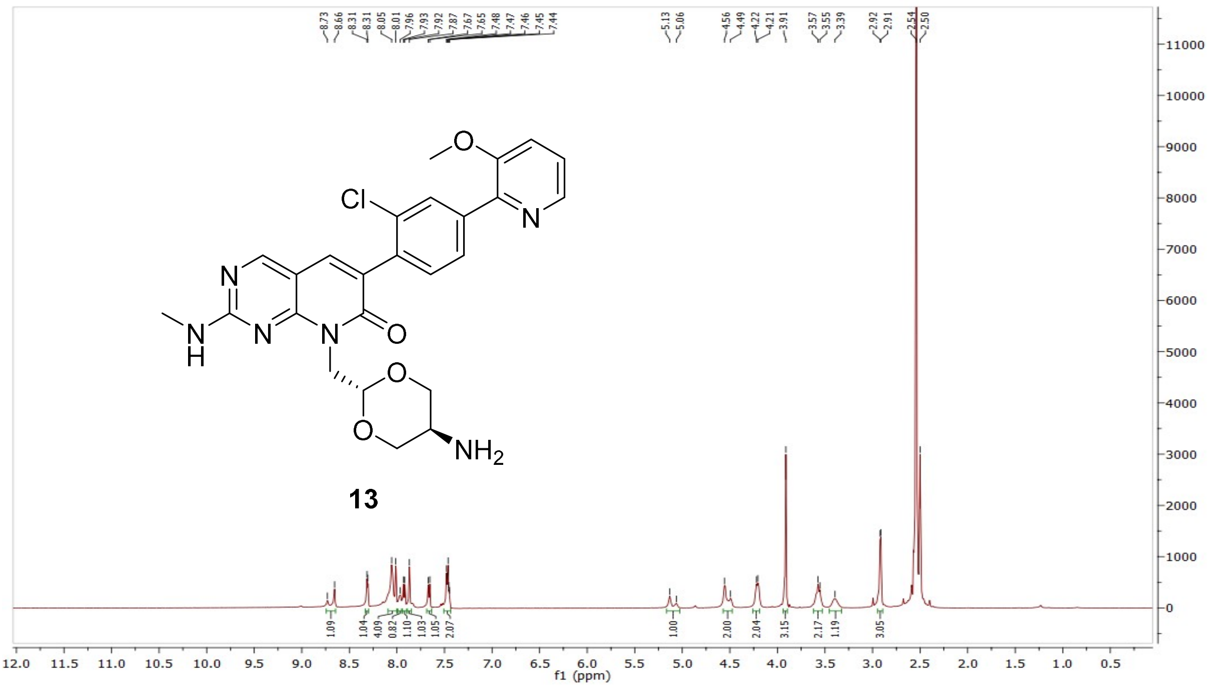


[13]C NMR compound **13**:

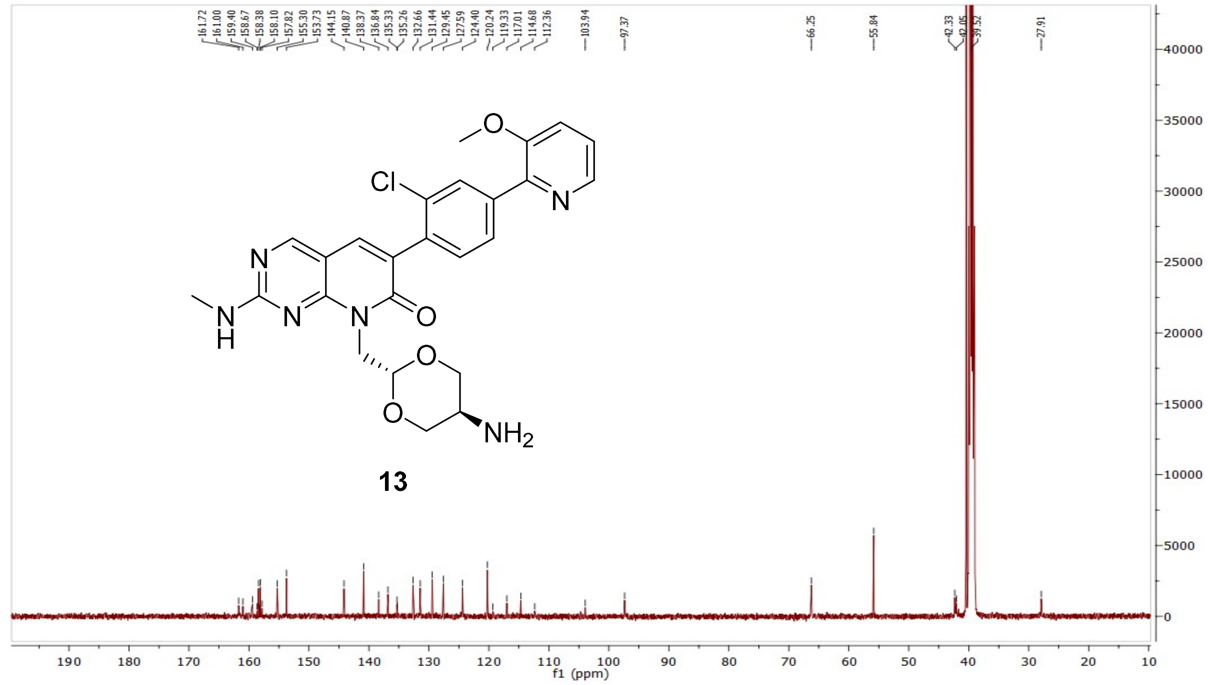

HRMS (FTMS +p MALDI) compound **13**:

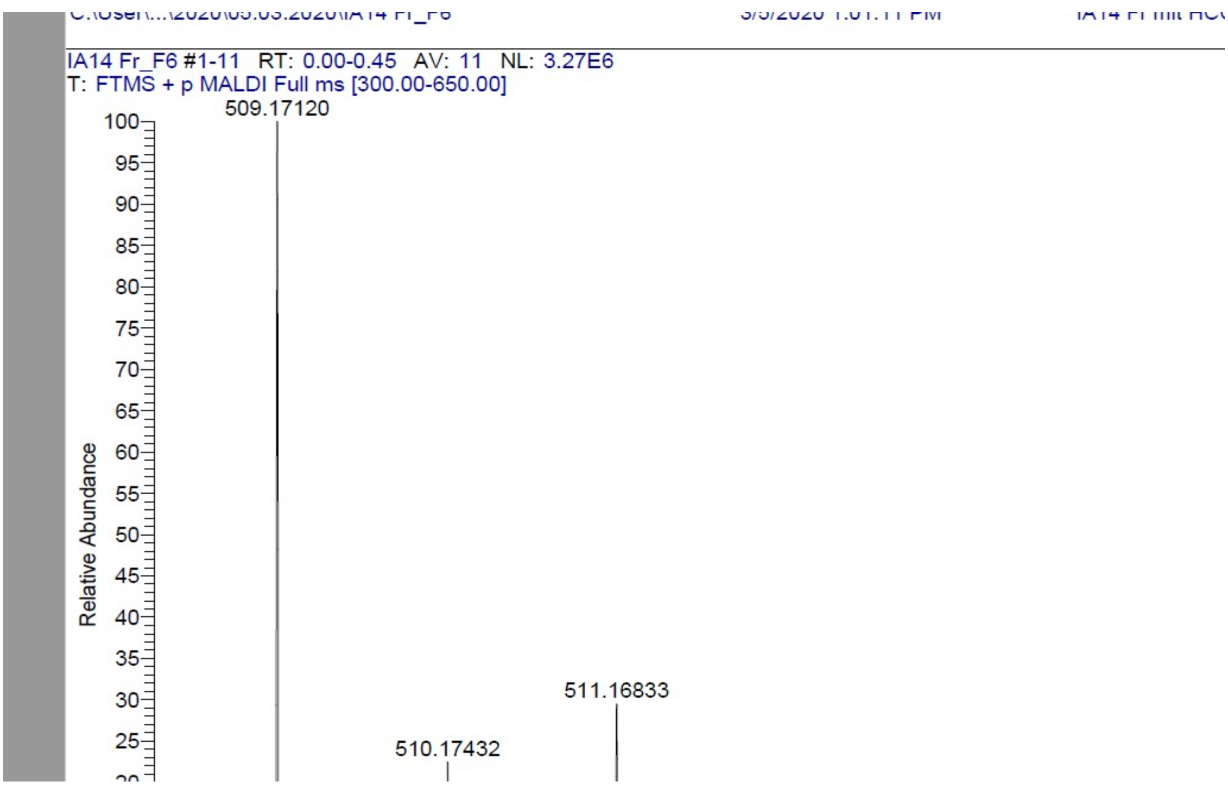

HPLC-chromatograms compound **13**:

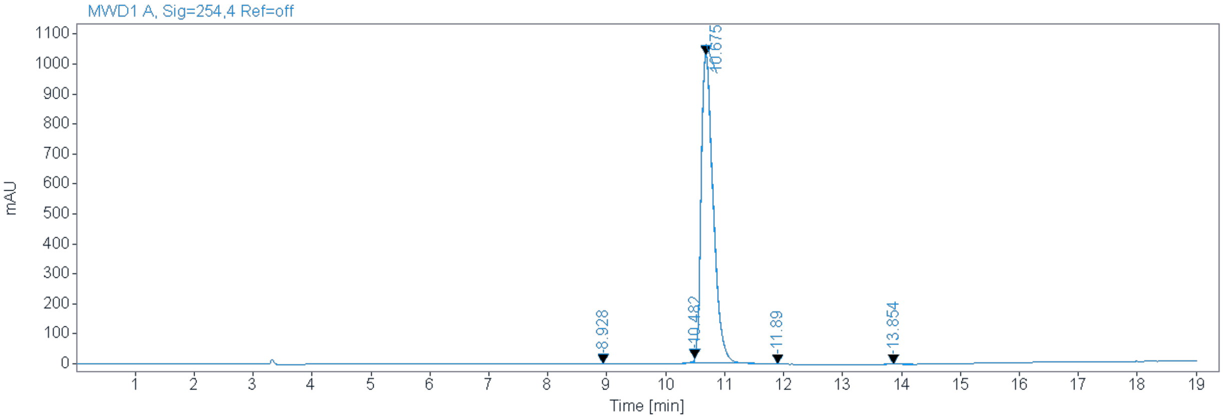


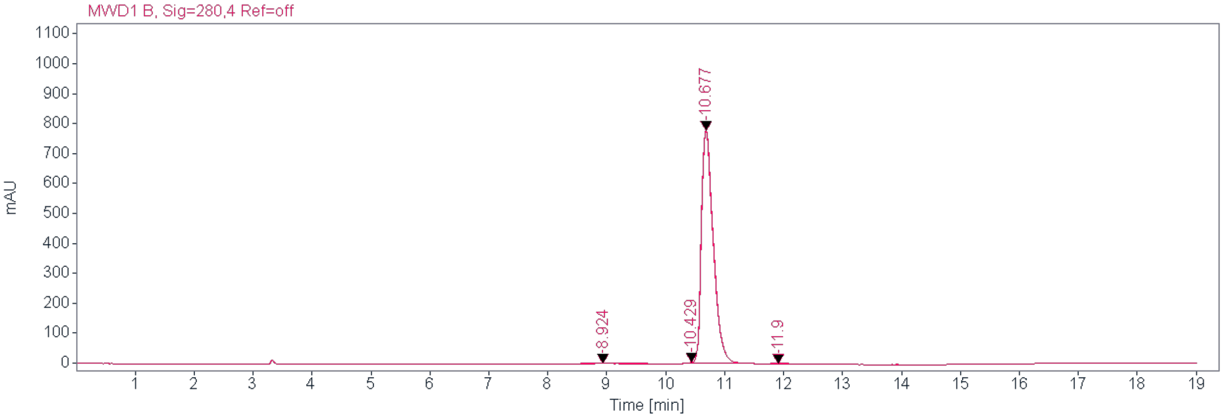


**Signal:** MWD1 B, Sig=254,4 Ref=off

| RT [min] Type | Width [min] | Area | Height | Area% Name |
|---|---|---|---|---|
| 8.928 MM | 0.7858 | 78.3781 | 1.6623 | 0.5283 |
| 0.482 MM | 0.0530 | 54.7126 | 17.2084 | 0.3688 |
| 10.675 MM | 0.237 | 114668.0869 | 1030.8914 | 98.8728 |
| 11.890 MM | 0.1731 | 11.7460 | 1.1311 | 0.0792 |
| 13.8541 MM | 0.2388 | 22.3856 | 1.5621 | 0.1509 |
| | Sum | 14835.3092 | | |

**Signal:** MWD1 B, Sig=280,4 Ref=off

| RT [min] Type | Width [min] | Area | Height | Area% Name |
|---|---|---|---|---|
| 8.924 MM | 0.5921 | 35.1620 | 0.9898 | 0.3194 |
| 10.429 MM | 0.0679 | 16.9301 | 4.1544 | 0.1538 |
| 10.677 MM | 0.2343 | 10946.5010 | 778.5373 | 99.4392 |
| 11.900 MM | 0.1744 | 9.6435 | 0.9218 | 0.0876 |
| | Sum | 11008.2365 | | |

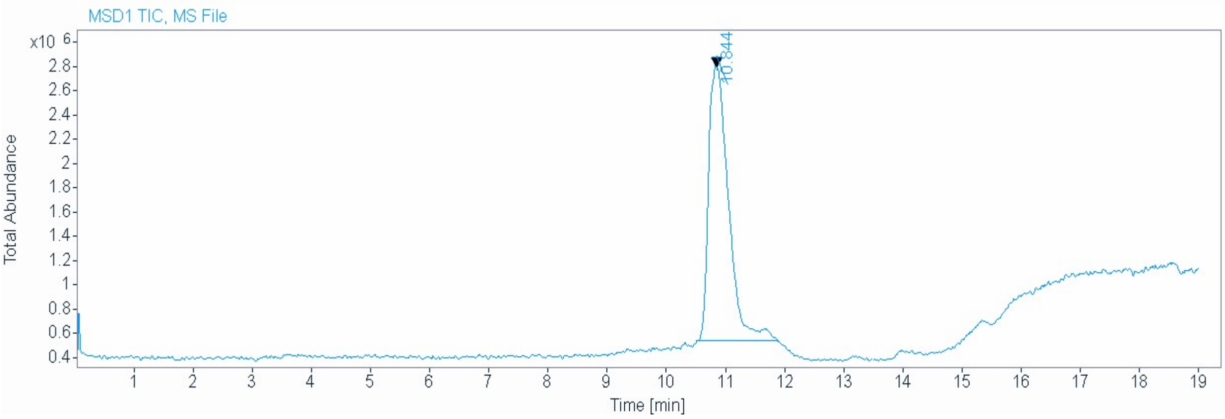


| | |
|---|---|
| **Signal** | MSD1 TIC, MS File |
| **Peak RT** | 10.844 |

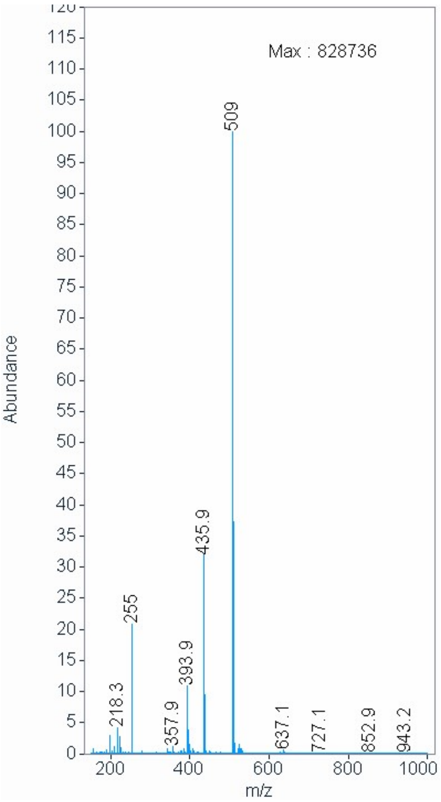

$^{1}H$ NMR compound **14**:

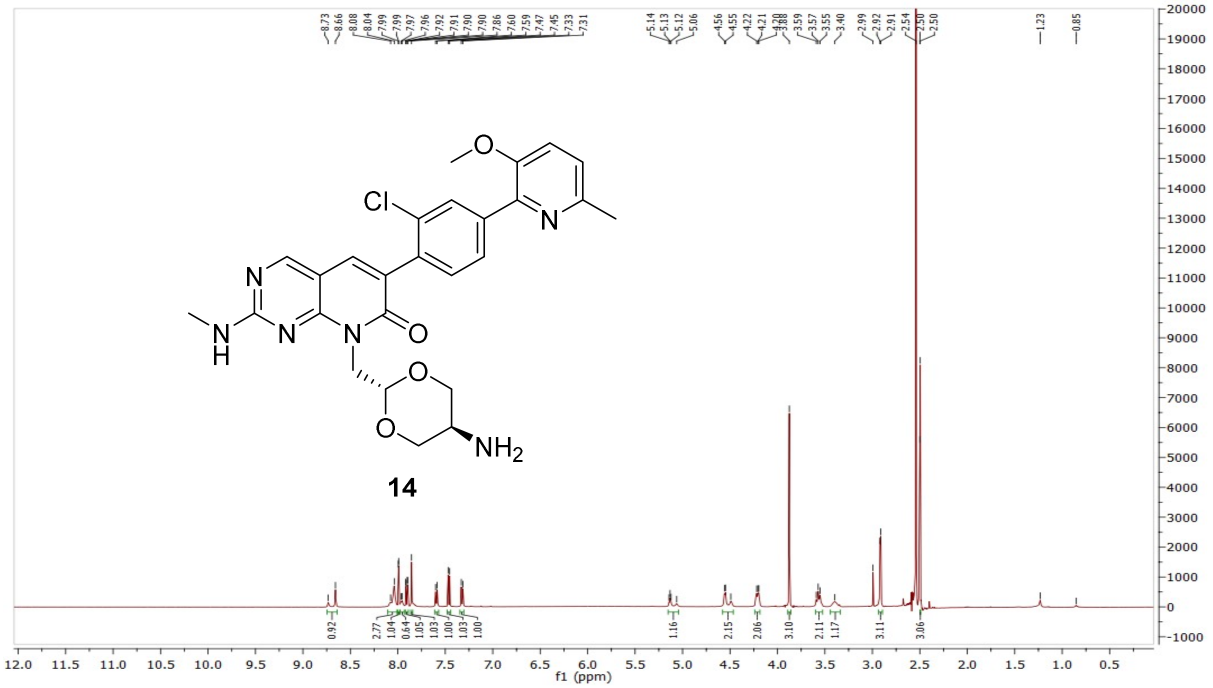

$^{13}C$ NMR compound **14**:

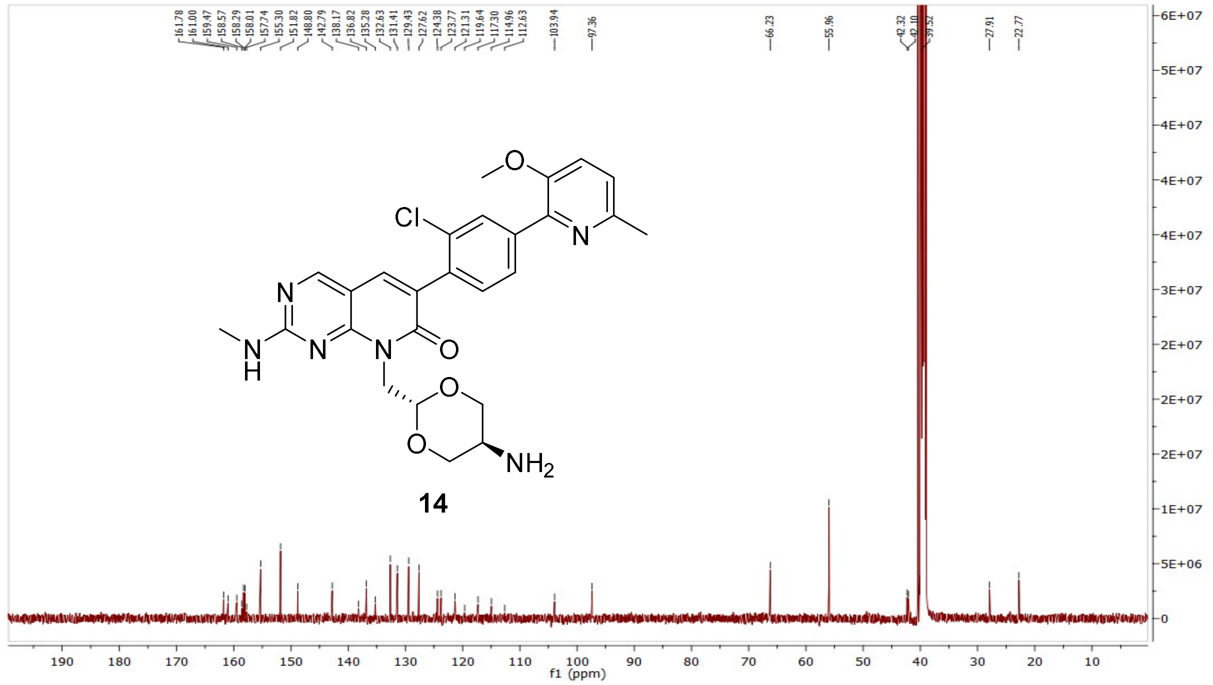

HRMS (FTMS +p MALDI) compound **14**:

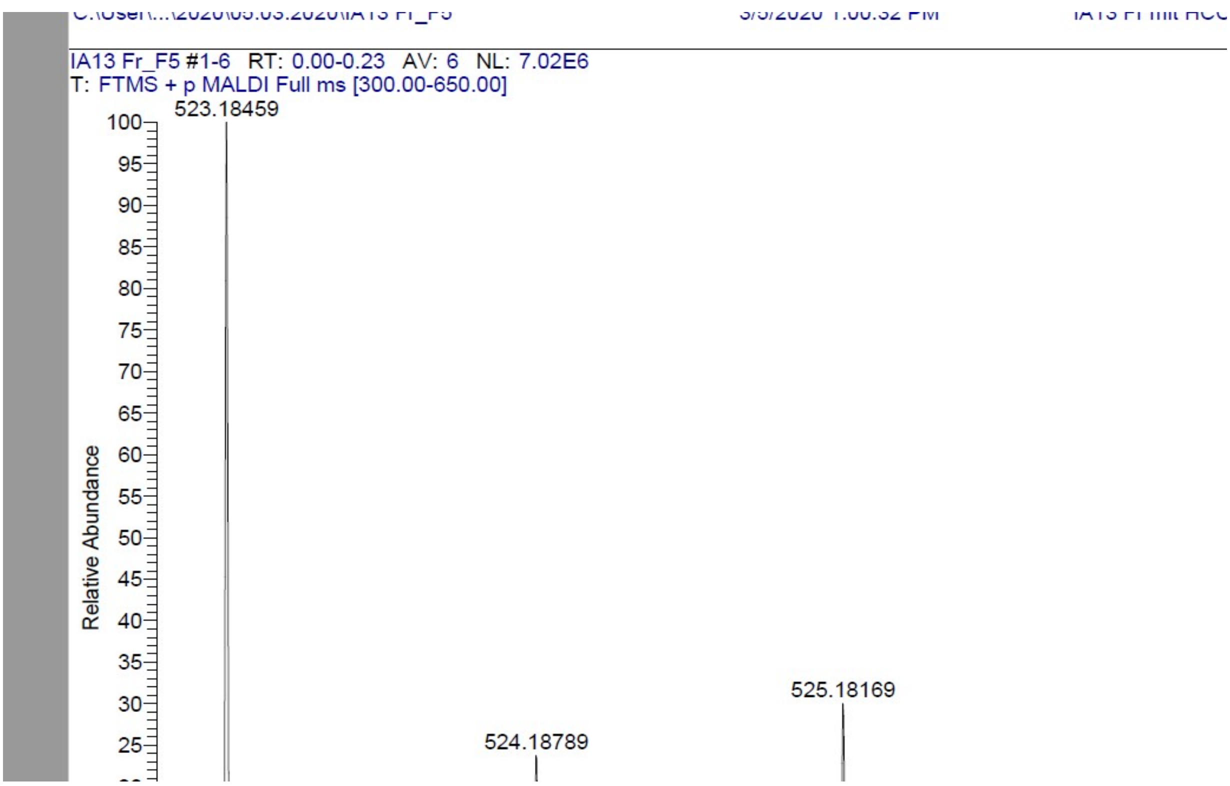

HPLC-chromatograms compound **14**:

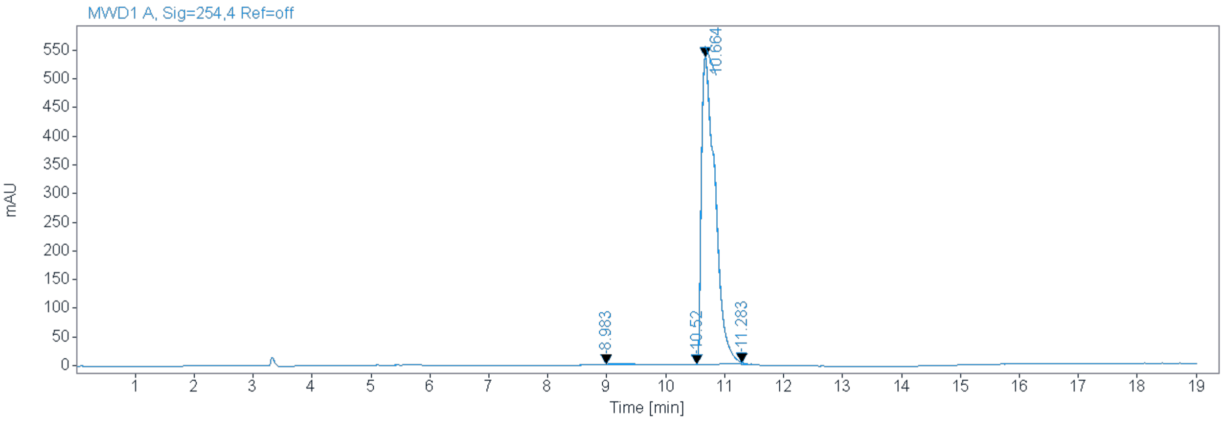


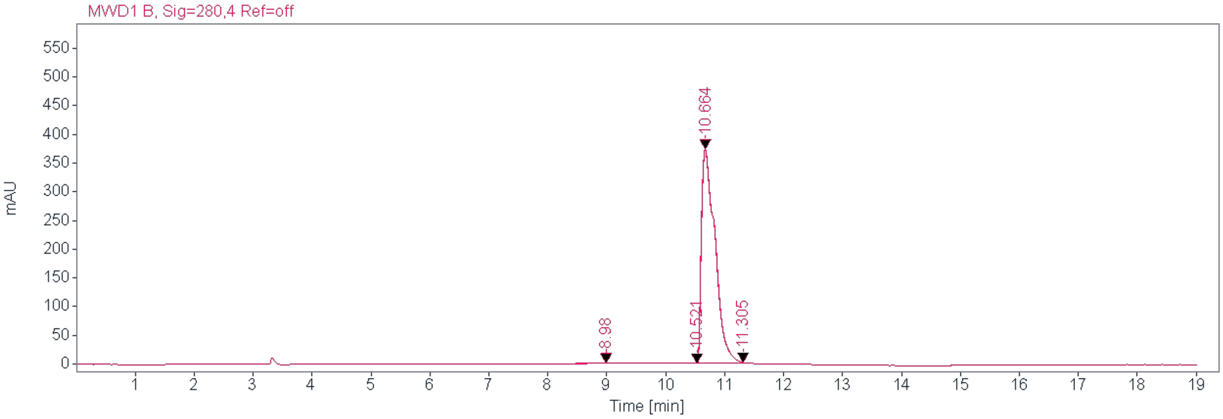


**Signal:** MWD1 B, Sig=254,4 Ref=off

| RT [min] Type | Width [min] | Area | Height | Area% Name |
|---|---|---|---|---|
| 8.983 MM | 0.5270 | 40.0456 | 1.2665 | 0.4618 |
| 10.520 MM | 0.0872 | 4.0174 | 0.7682 | 0.0463 |
| 10.664 MM | 0.2675 | 8613.9375 | 536.6498 | 99.3355 |
| 11.283 MM | 0.0454 | 13.5567 | 3.5621 | 0.1563 |
| | Sum | 8671.5573 | | |

**Signal:** MWD1 B, Sig=280,4 Ref=off

| RT [min] Type | Width [min] | Area | Height | Area% Name |
|---|---|---|---|---|
| 8.980 MM | 0.4528 | 23.3658 | 0.8601 | 0.3876 |
| 10.521 MM | 0.1583 | 8.0296 | 0.8453 | 0.1332 |
| 10.664 MM | 0.2669 | 5988.4331 | 373.9075 | 99.3269 |
| 11.305 MM | 0.0718 | 9.1867 | 2.1336 | 0.1524 |
| | Sum | 6029.0152 | | |

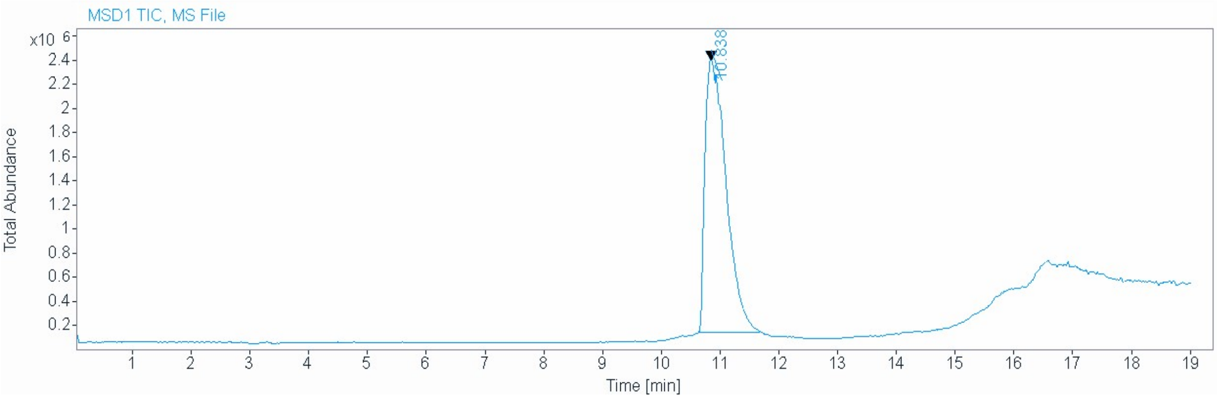


**Signal** MSD1 TIC, MS File

**Peak RT** 10.838

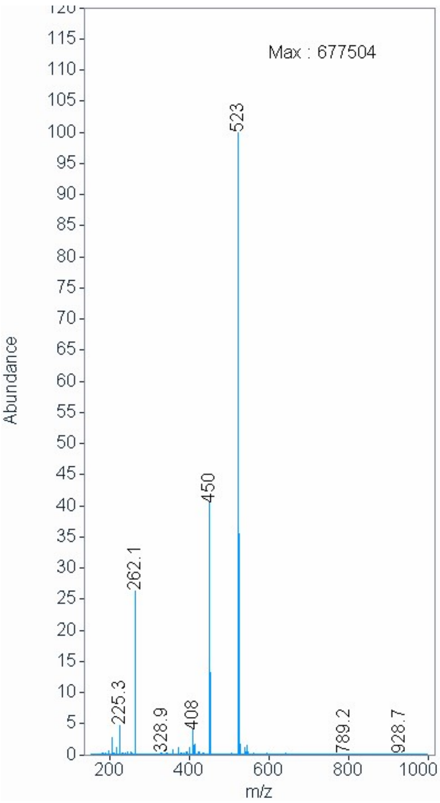

$^{1}$H NMR compound **15**:

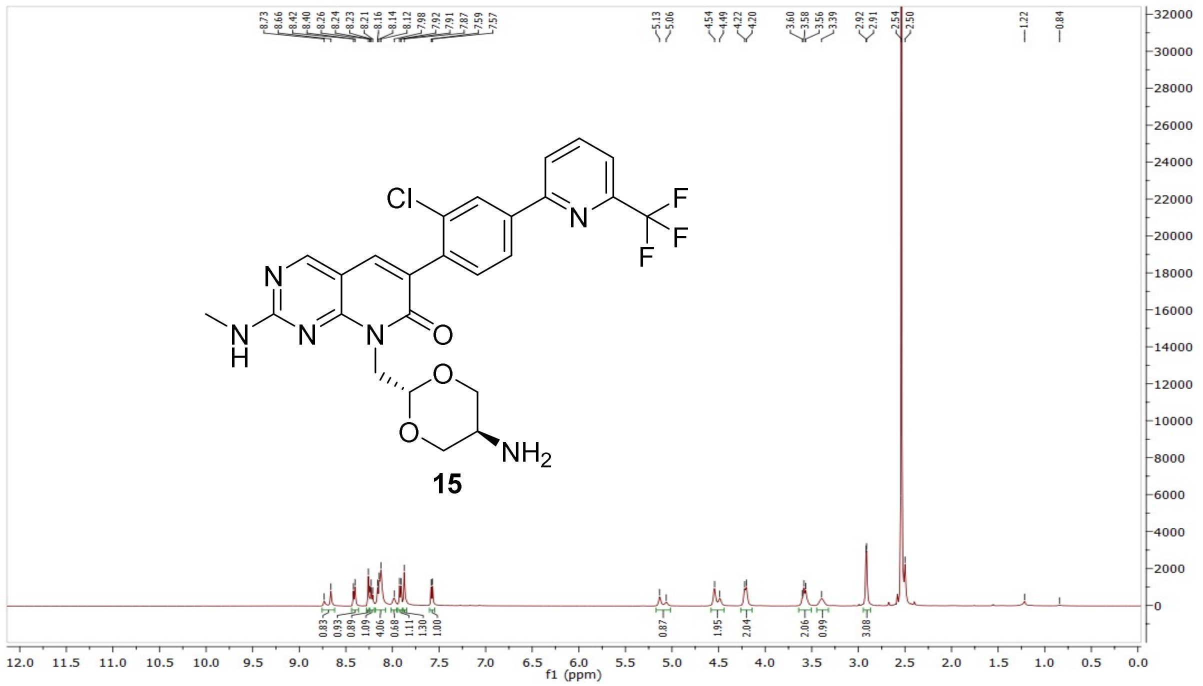


$^{13}$C NMR compound **15**:

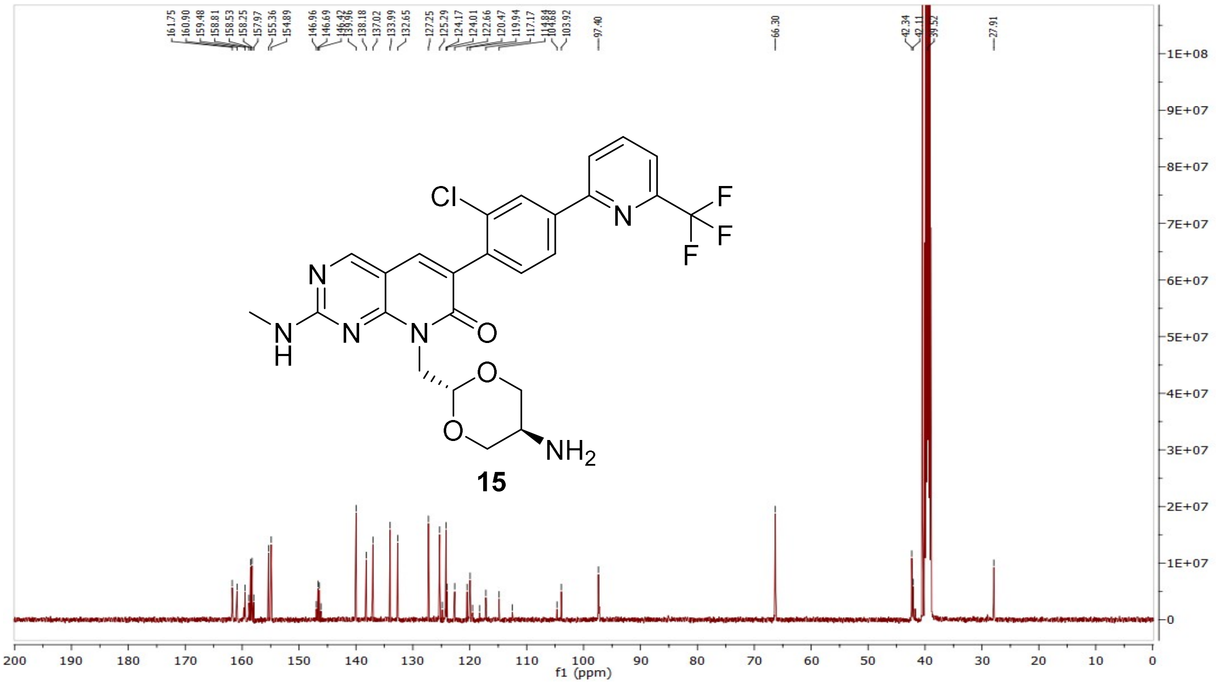

HRMS (FTMS +p MALDI) compound **15**:

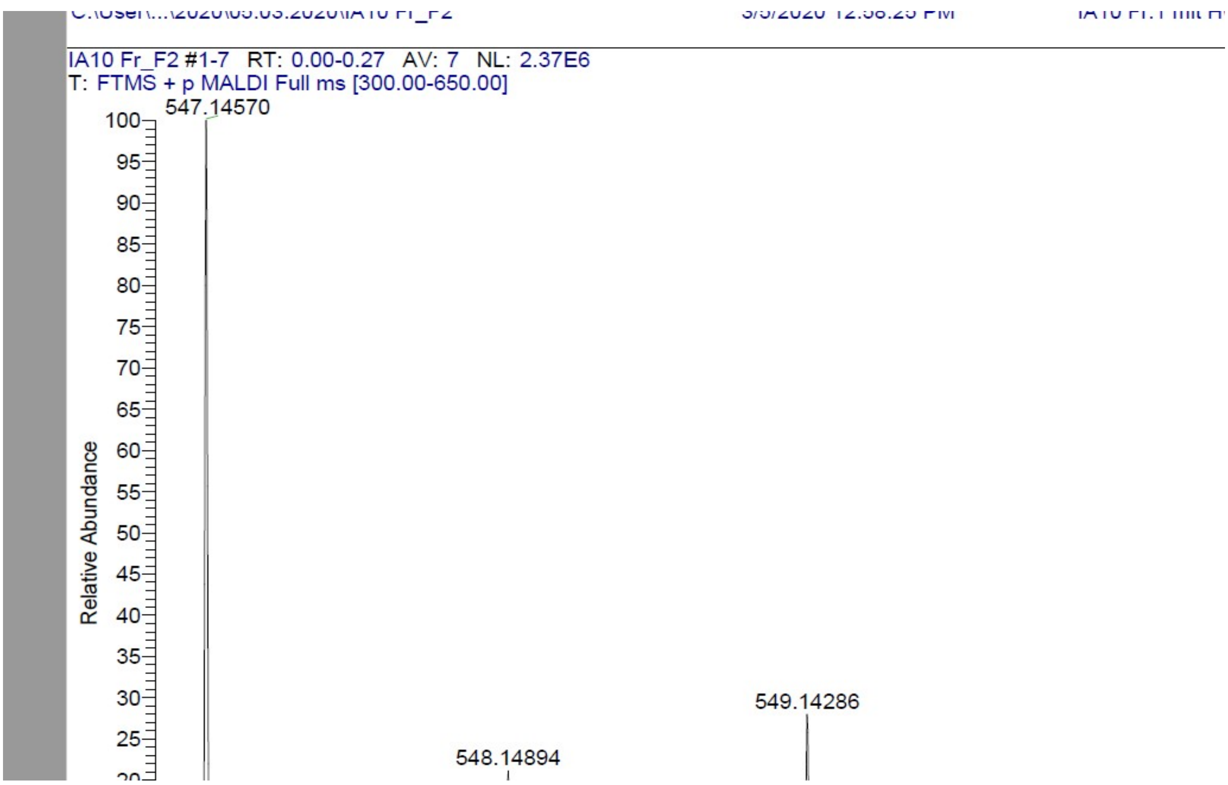

HPLC-chromatograms compound **15**:

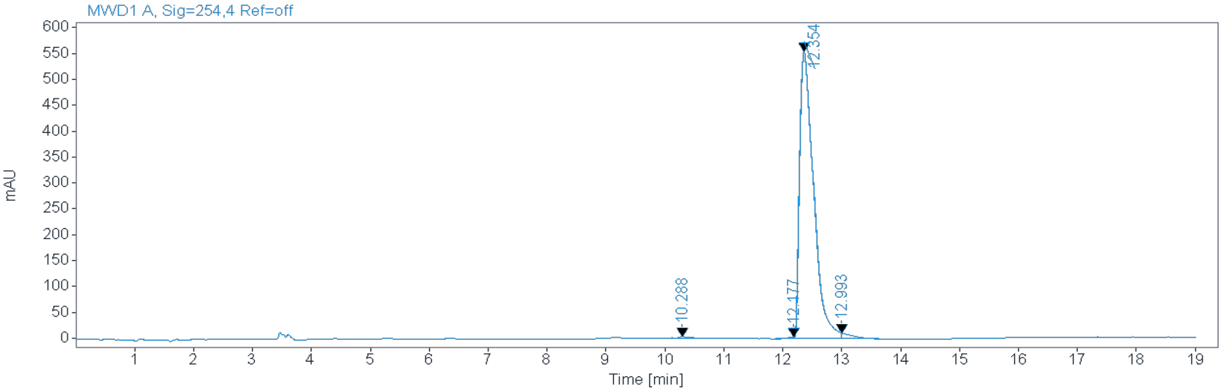


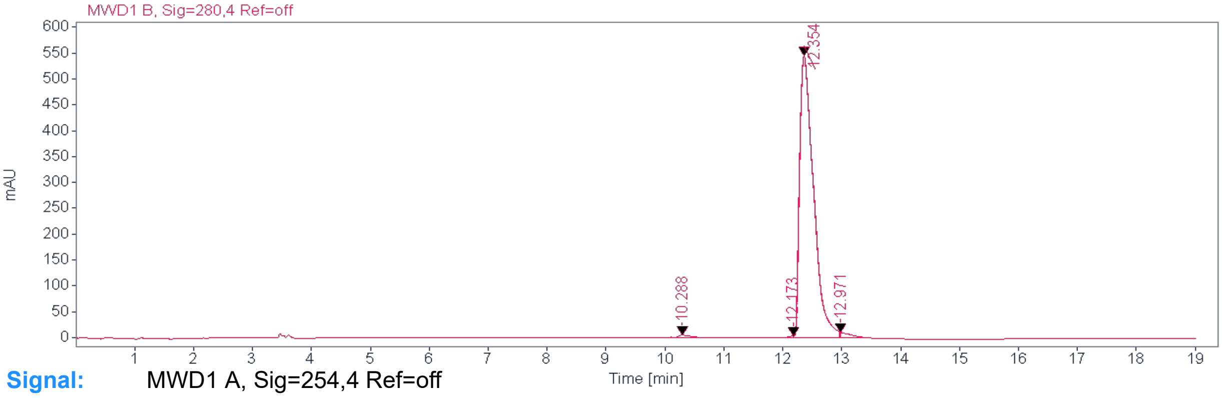


**Signal:** MWD1 A, Sig=254,4 Ref=off

| RT [min] | Type | Width [min] | Area | Height | Area% | Name |
|---|---|---|---|---|---|---|
| 10.288 | MM | 0.2572 | 47.9928 | 3.1098 | 0.5090 | |
| 12.177 | MM | 0.1312 | 31.8189 | 4.0426 | 0.3375 | |
| 12.354 | MM | 0.2757 | 9188.1260 | 555.4003 | 97.4454 | |
| 12.993 | MM | 0.1682 | 161.0629 | 11.3575 | 1.7082 | |
| | | Sum | 9429.0006 | | | |

**Signal:** MWD1 B, Sig=280,4 Ref=off

| RT [min] | Type | Width [min] | Area | Height | Area% | Name |
|---|---|---|---|---|---|---|
| 10.288 | MM | 0.2357 | 68.5422 | 4.8457 | 0.7445 | |
| 12.173 | MM | 0.1164 | 27.3388 | 3.9133 | 0.2969 | |
| 12.354 | MM | 0.2739 | 8973.6221 | 545.9641 | 97.4655 | |
| 12.971 | MM | 0.1468 | 137.4744 | 11.1369 | 1.4932 | |
| | | Sum | 9206.9775 | | | |

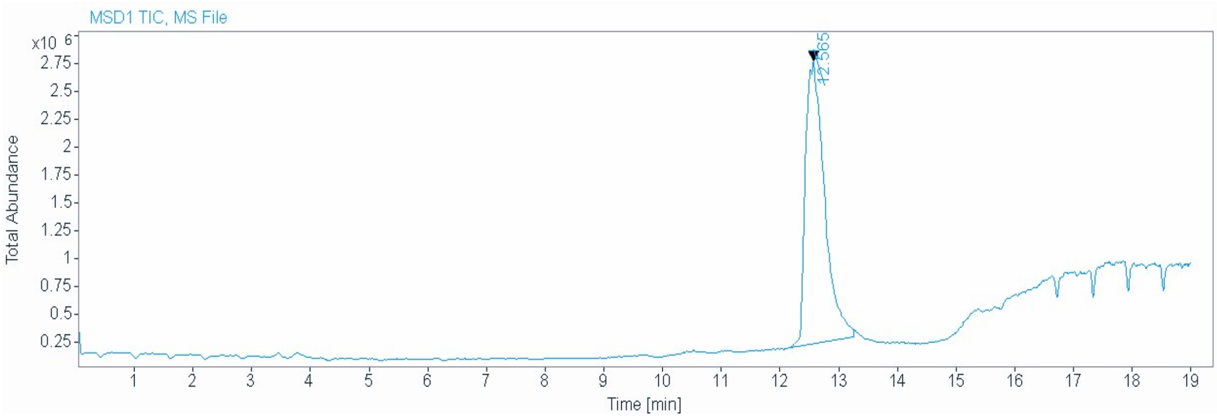


**Signal** MSD1 TIC, MS File

**Peak RT** 12.565

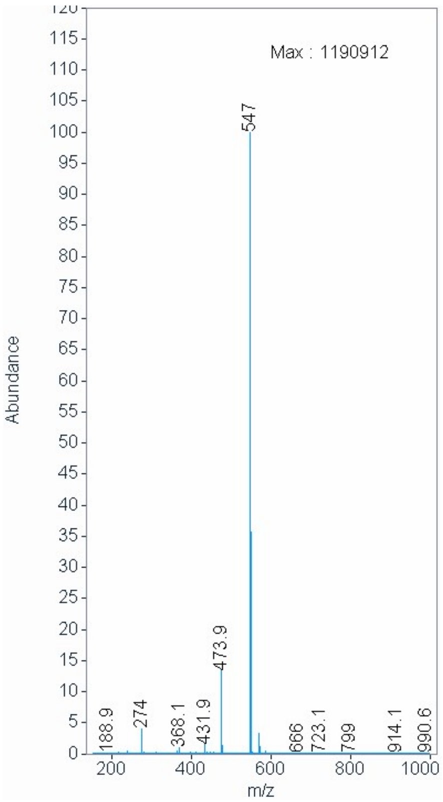

$^{1}$H NMR compound **16**:

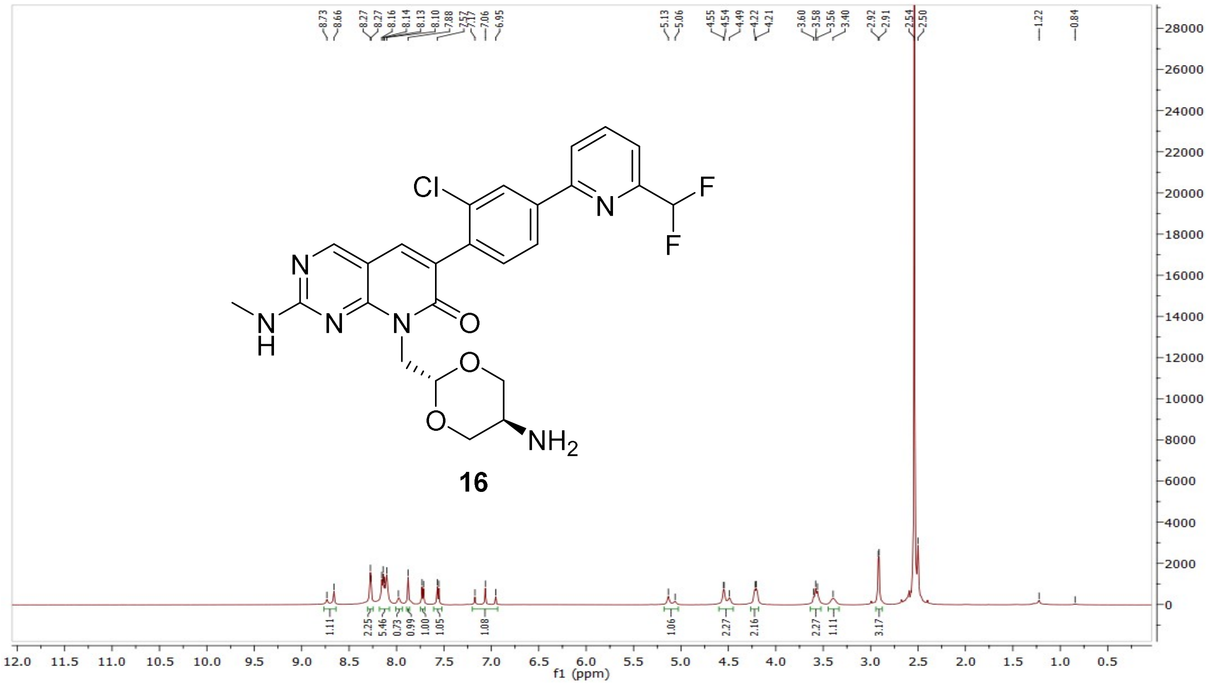


$^{13}$C NMR compound **16**:

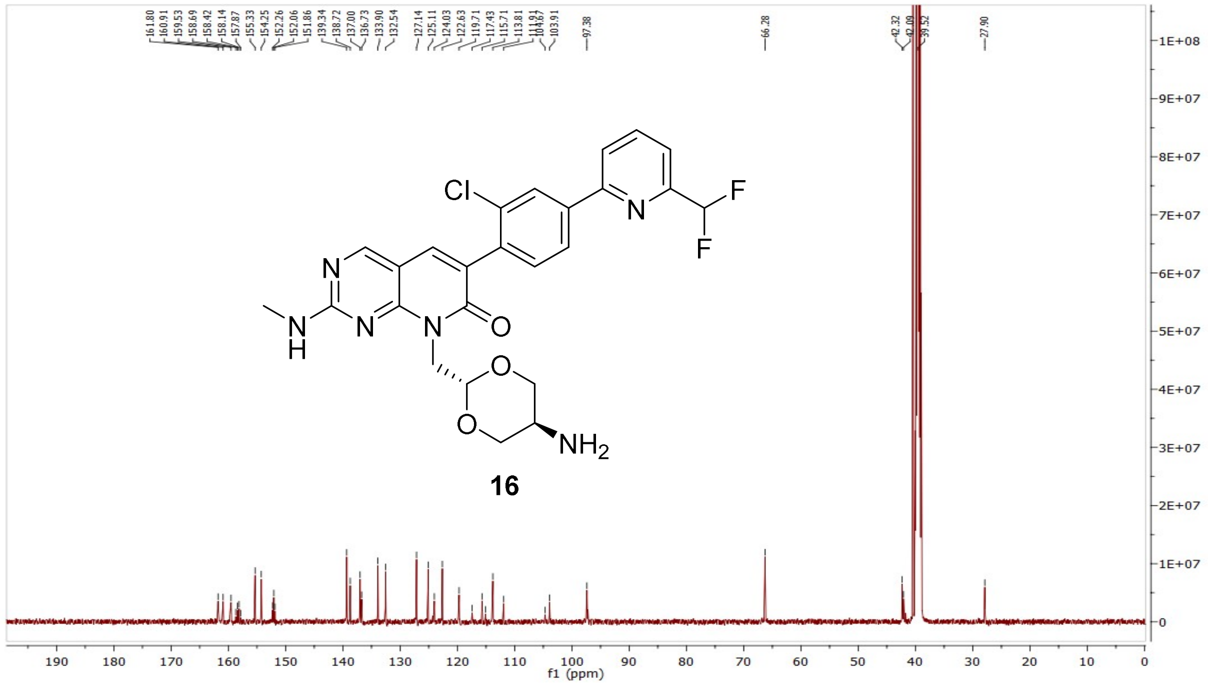

HRMS (FTMS +p MALDI) compound **16**:

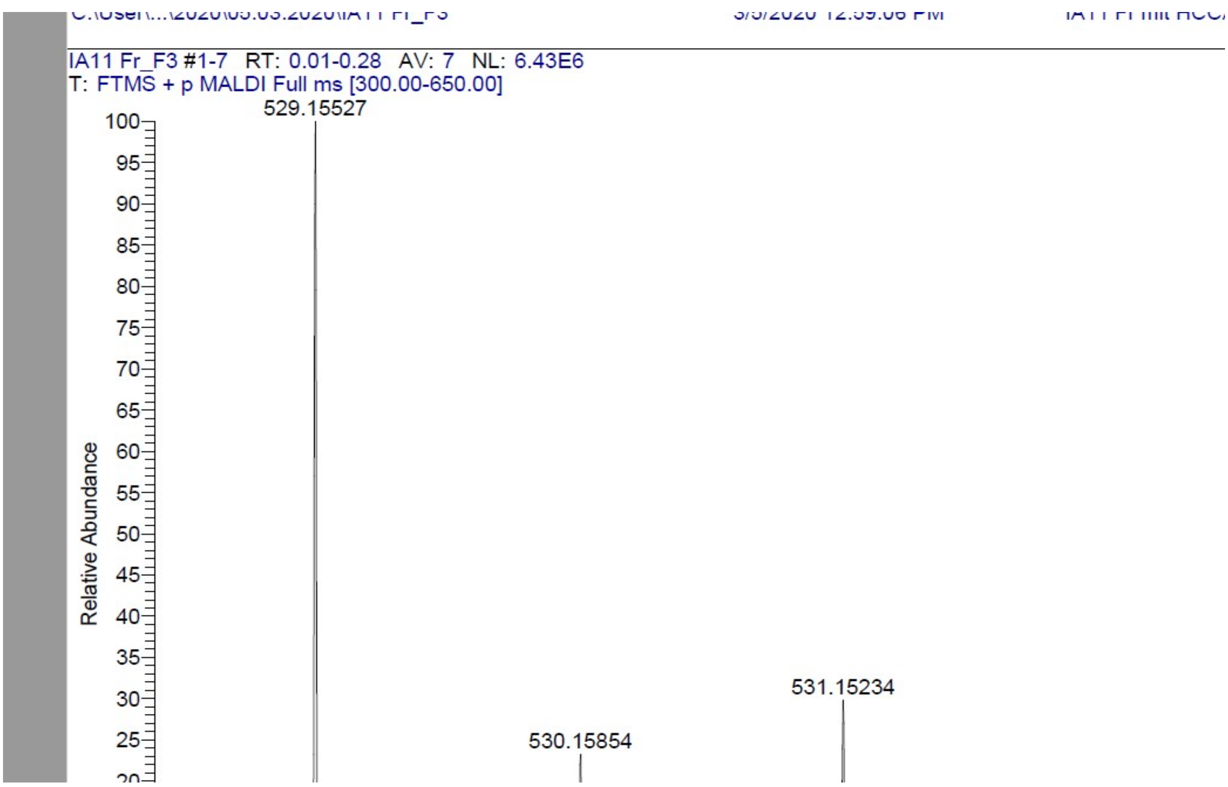

HPLC-chromatograms compound **16**:

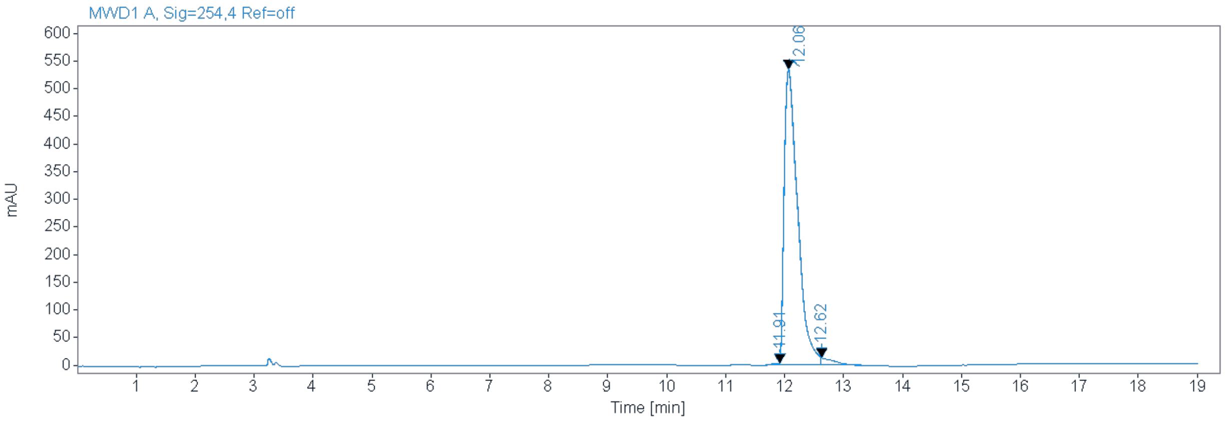


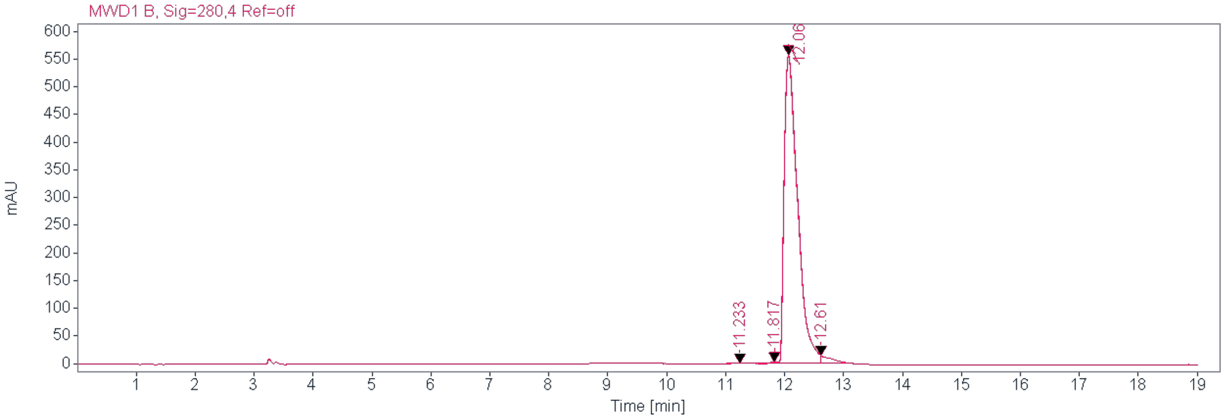


**Signal:** MWD1 B, Sig=254,4 Ref=off

| RT [min] Type | Width [min] | Area | Height | Area% Name |
|---|---|---|---|---|
| 11.910 MM | 0.1333 | 31.7624 | 3.9714 | 0.3618 |
| 12.060 MM | 0.2657 | 8540.8691 | 535.7968 | 97.2979 |
| 12.620 MM | 0.2482 | 205.4298 | 13.7930 | 2.3403 |
| | Sum | 8778.0613 | | |

**Signal:** MWD1 B, Sig=280,4 Ref=off

| RT [min] Type | Width [min] | Area | Height | Area% Name |
|---|---|---|---|---|
| 11.233 MM | 0.3910 | 17.4154 | 0.7424 | 0.1915 |
| 11.817 MM | 0.1473 | 25.8467 | 2.9238 | 0.2843 |
| 12.060 MM | 0.2646 | 8852.8828 | 557.6185 | 97.3696 |
| 12.610 MM | 0.1574 | 195.8993 | 14.5508 | 2.1546 |
| | Sum | 9092.0442 | | |

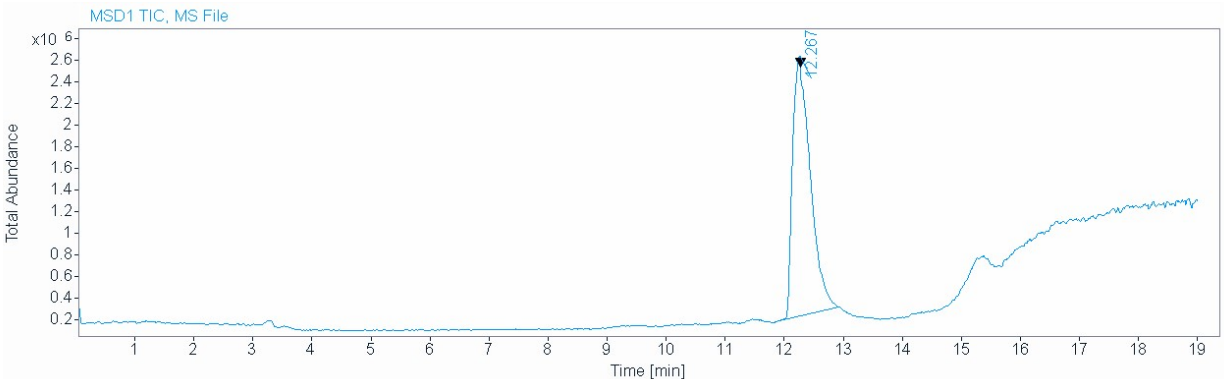


| | |
|---|---|
| **Signal** | MSD1 TIC, MS File |
| **Peak RT** | 12.267 |

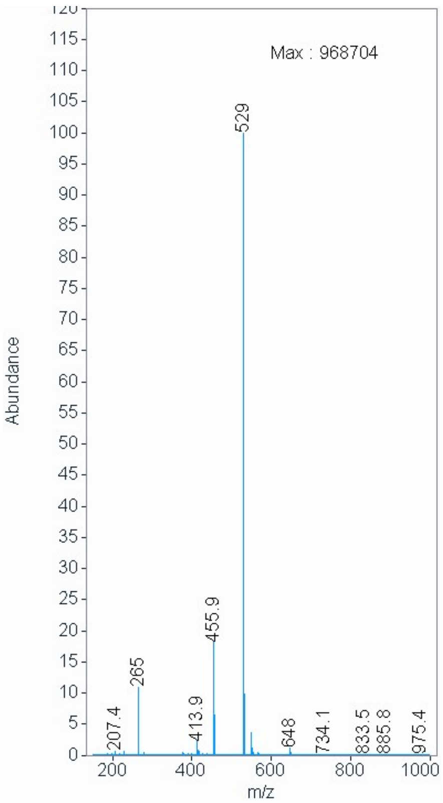

[1]H NMR compound **17**:

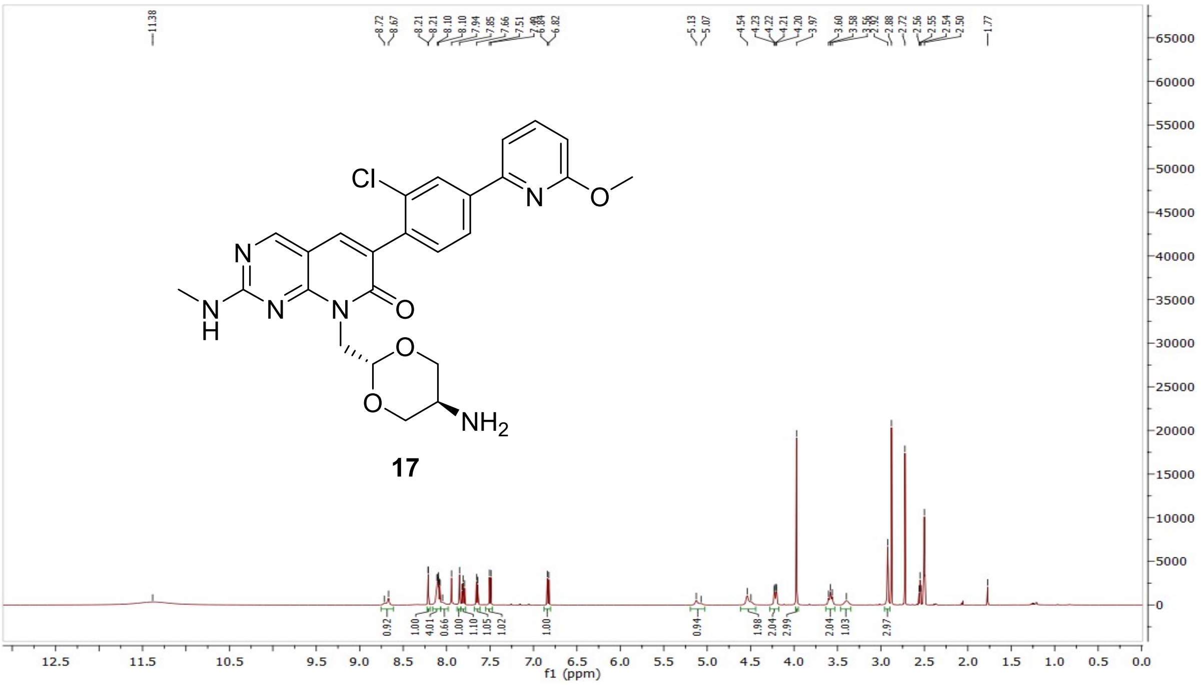


[13]C NMR compound **17**:

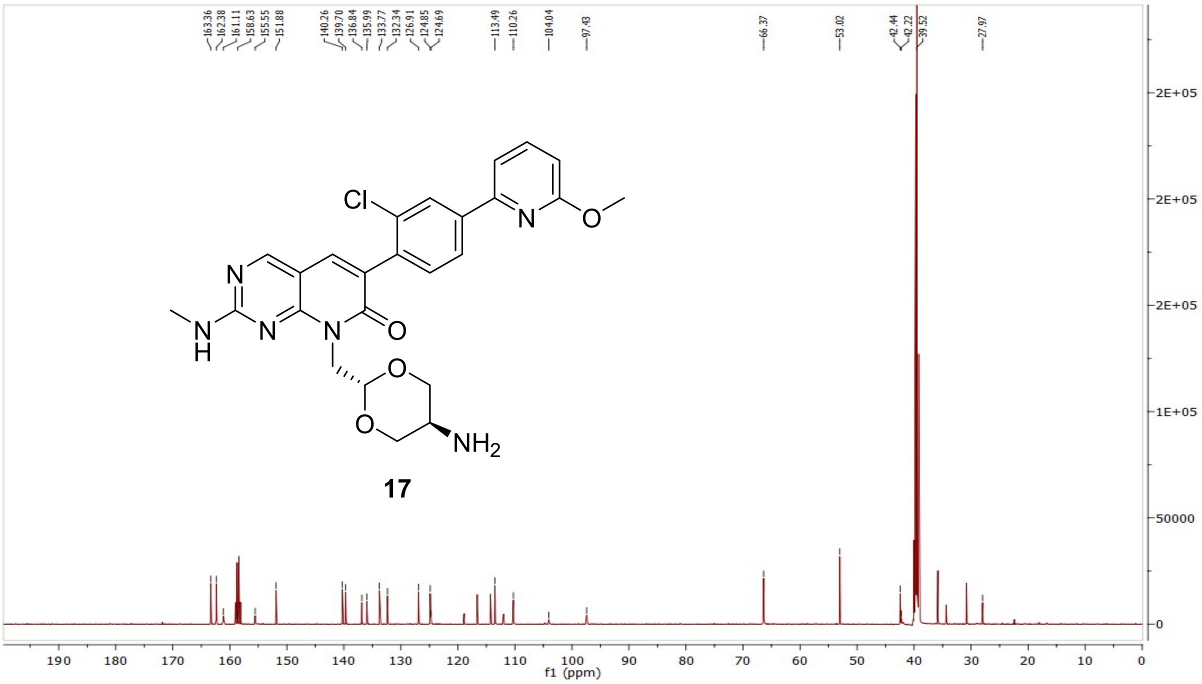

HRMS (FTMS +p MALDI) compound **17**:

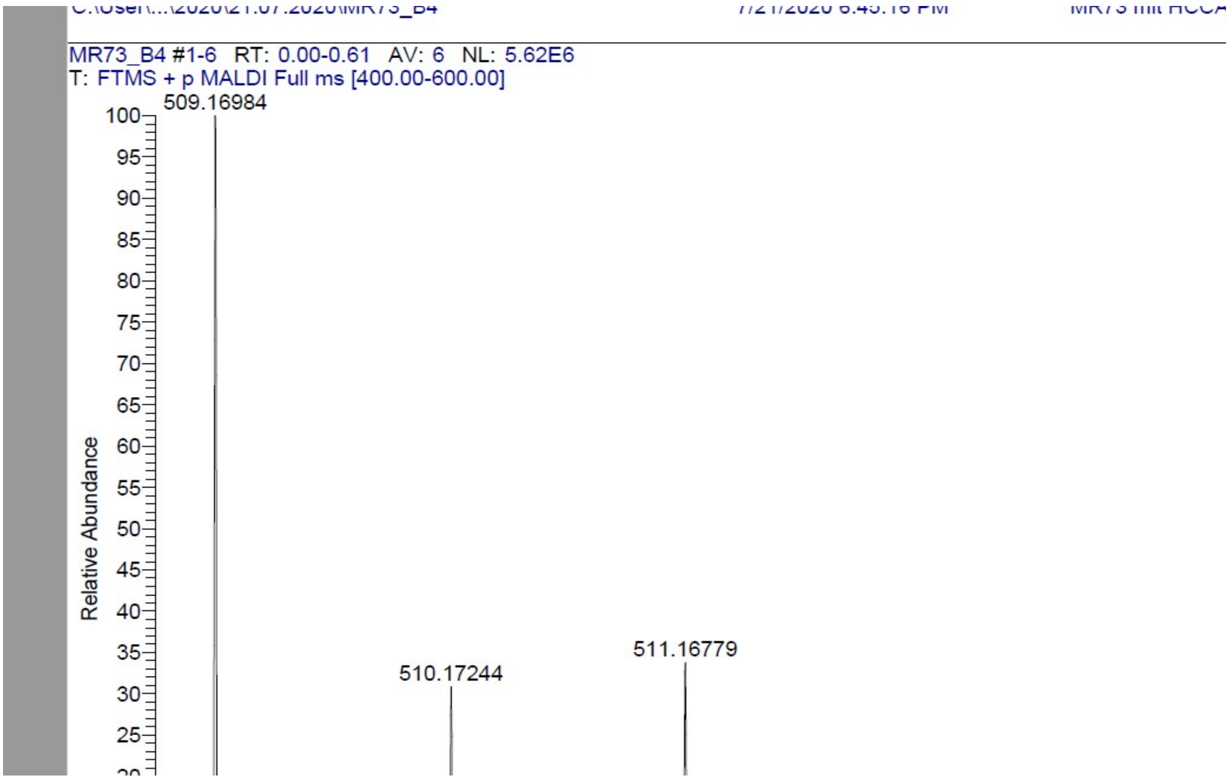

HPLC-chromatograms compound **17**:

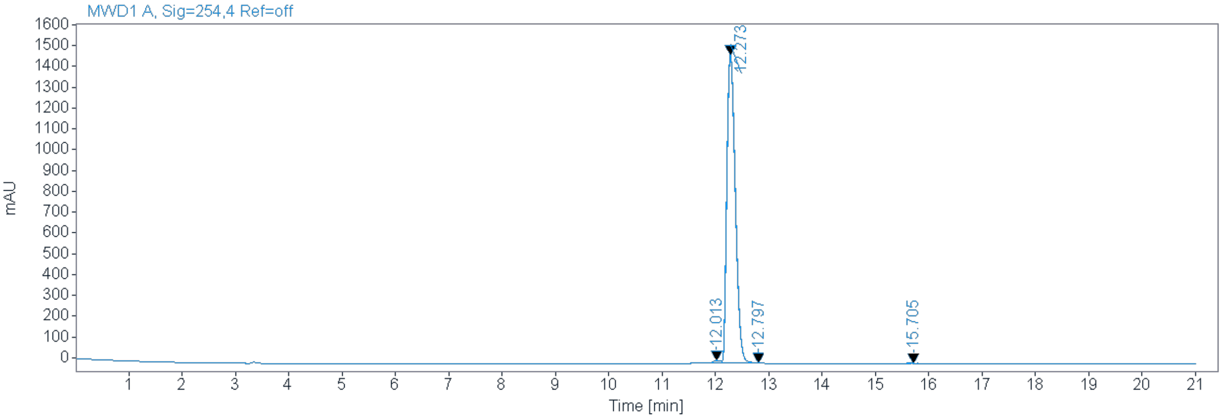


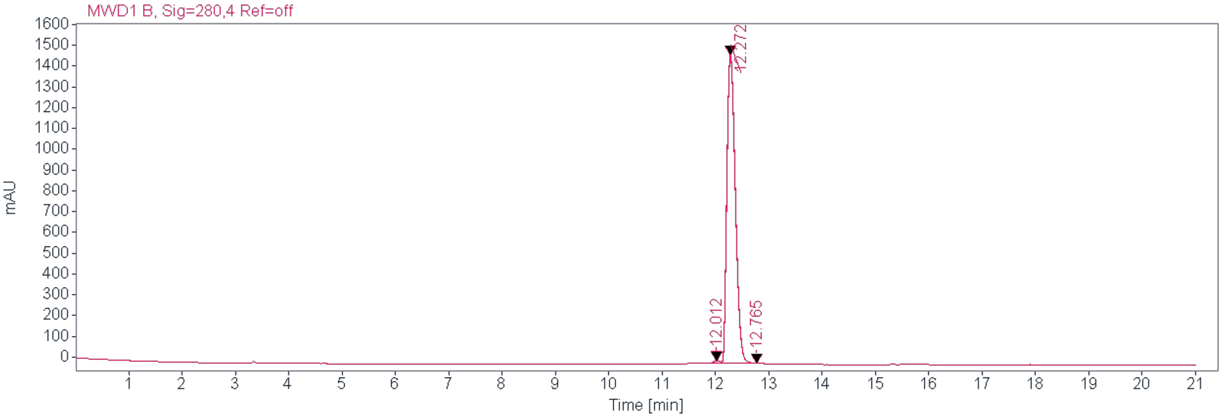


**Signal:** MWD1 A, Sig=254,4 Ref=off

| RT [min] | Type | Width [min] | Area | Height | Area% | Name |
|---|---|---|---|---|---|---|
| 12.013 | MM | 0.2048 | 179.2532 | 14.5895 | 1.0797 | |
| 12.273 | MM | 0.1842 | 16380.1162 | 1482.1260 | 98.6652 | |
| 12.797 | MM | 0.0800 | 17.5503 | 2.6820 | 0.1057 | |
| 15.705 | MM | 0.1394 | 24.7923 | 2.9645 | 0.1493 | |
| | | Sum | 16601.7119 | | | |

**Signal:** MWD1 B, Sig=280,4 Ref=off

| RT [min] | Type | Width [min] | Area | Height | Area% | Name |
|---|---|---|---|---|---|---|
| 12.012 | MM | 0.2251 | 178.3978 | 13.2059 | 1.0702 | |
| 12.272 | MM | 0.1848 | 16472.2617 | 1485.4857 | 98.8137 | |
| 12.765 | MM | 0.0750 | 19.3637 | 3.1213 | 0.1162 | |
| | | Sum | 16670.0232 | | | |

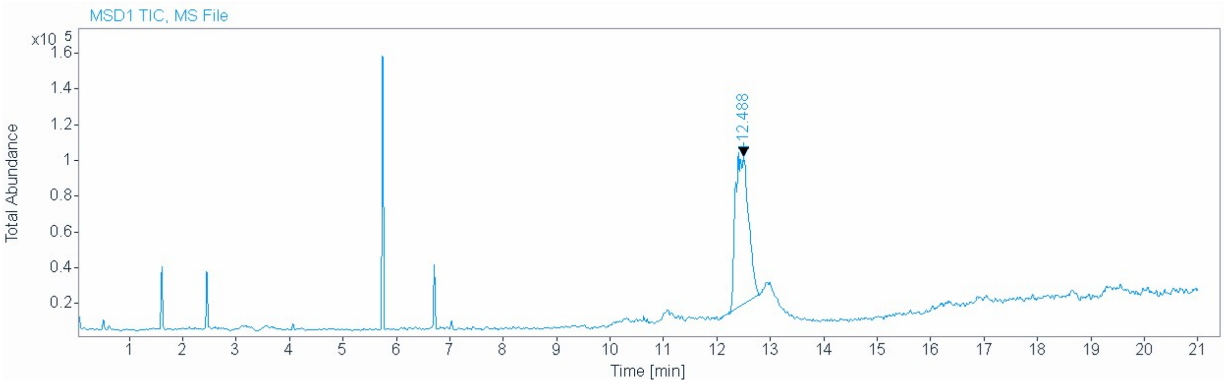


**Signal** MSD1 TIC, MS File

**Peak RT** 12.488

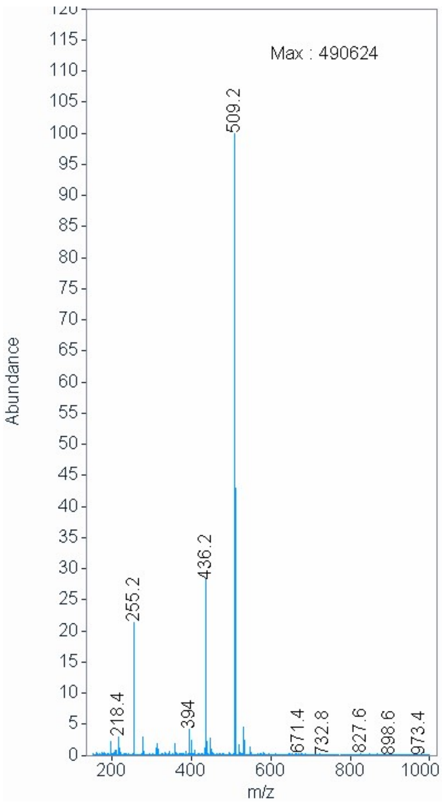

[1]H NMR compound **18**:

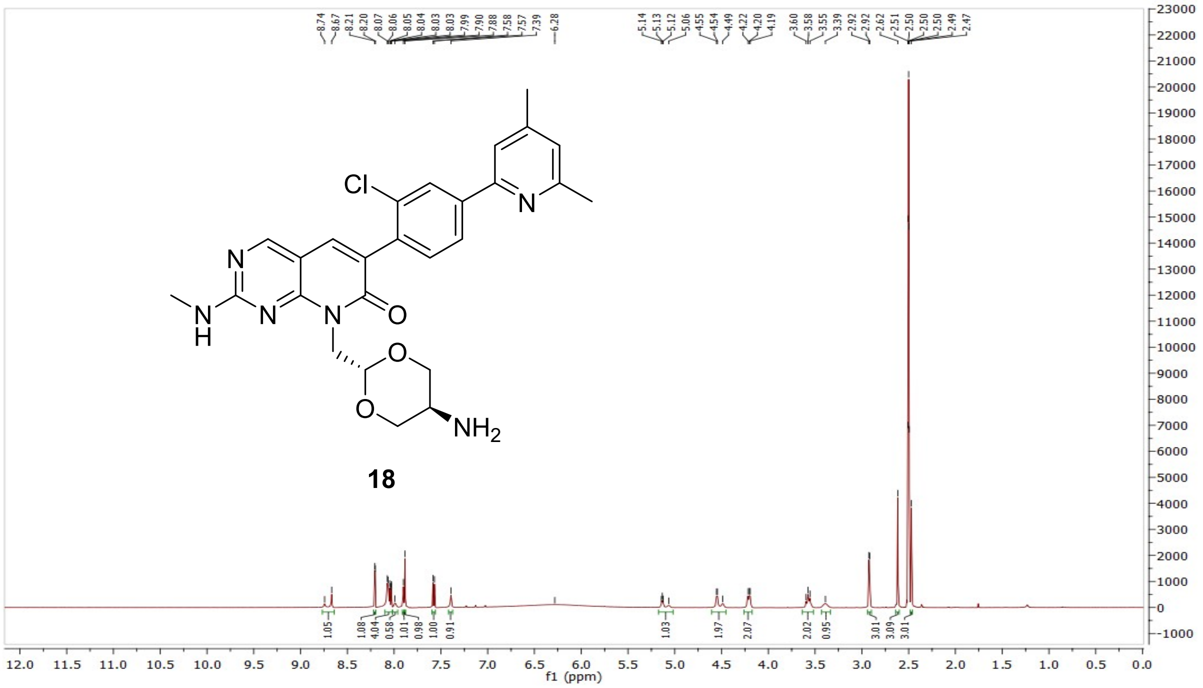


[13]C NMR compound **18**:

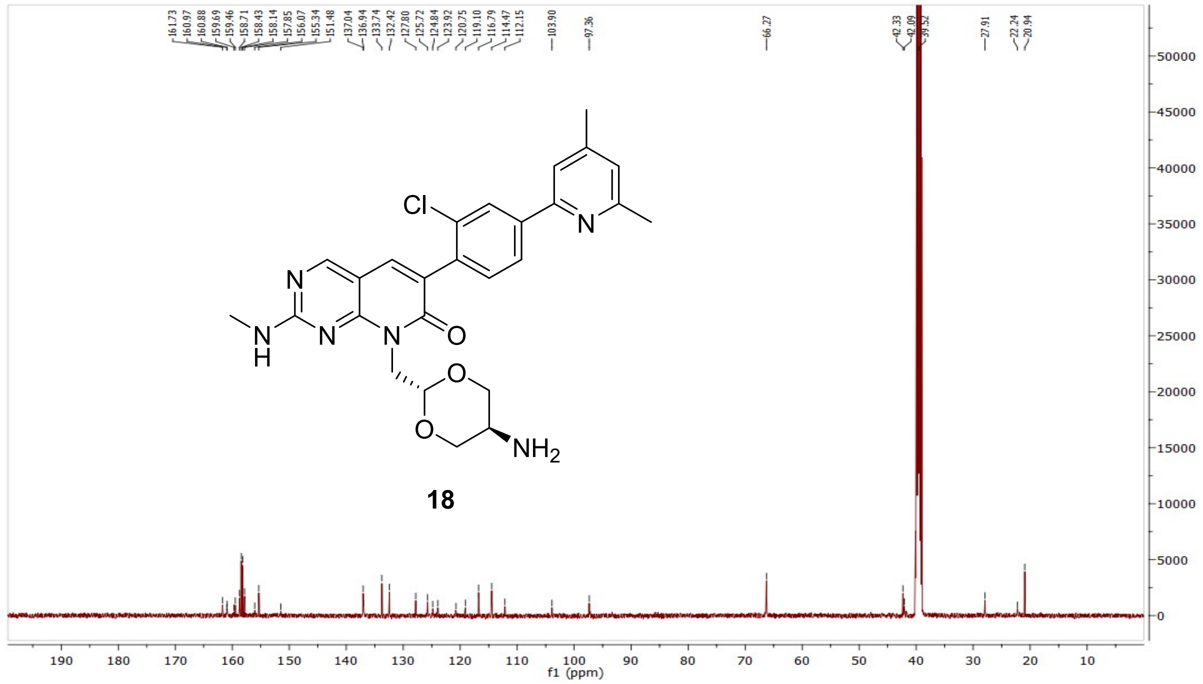

HRMS (FTMS +p MALDI) compound **18**:

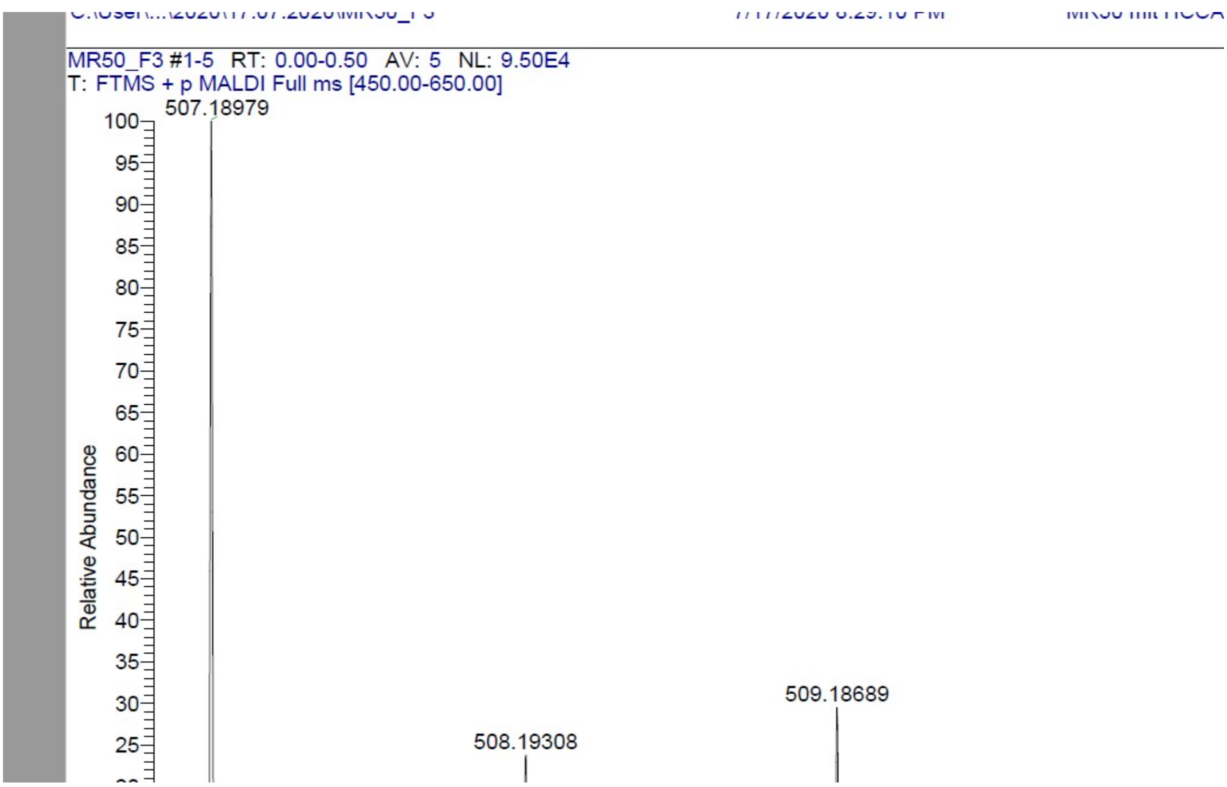

HPLC-chromatograms compound **18**:

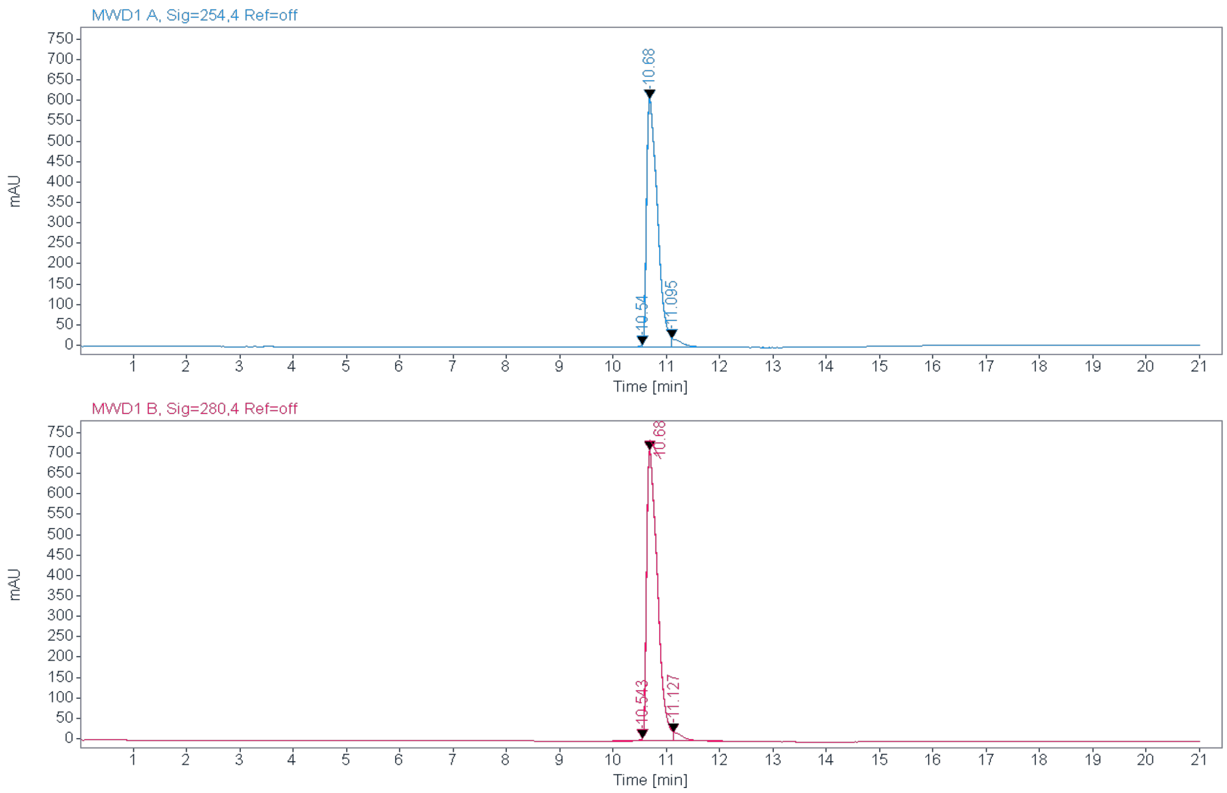


**Signal:** MWD1 A, Sig=254,4 Ref=off

| RT [min] | Type | Width [min] | Area | Height | Area% | Name |
|---|---|---|---|---|---|---|
| 10.540 | MM | 0.0698 | 24.7134 | 5.9042 | 0.2805 | |
| 10.680 | MM | 0.2329 | 8514.1914 | 609.2924 | 96.6284 | |
| 11.095 | MM | 0.1652 | 272.3693 | 21.4063 | 3.0911 | |
| | | Sum | 8811.2740 | | | |

**Signal:** MWD1 B, Sig=280,4 Ref=off

| RT [min] | Type | Width [min] | Area | Height | Area% | Name |
|---|---|---|---|---|---|---|
| 10.543 | MM | 0.0942 | 37.4948 | 6.6360 | 0.3629 | |
| 10.680 | MM | 0.2338 | 10014.0166 | 713.7814 | 96.9262 | |
| 11.127 | MM | 0.1574 | 280.0792 | 21.8666 | 2.7109 | |
| | | Sum | 10331.5906 | | | |

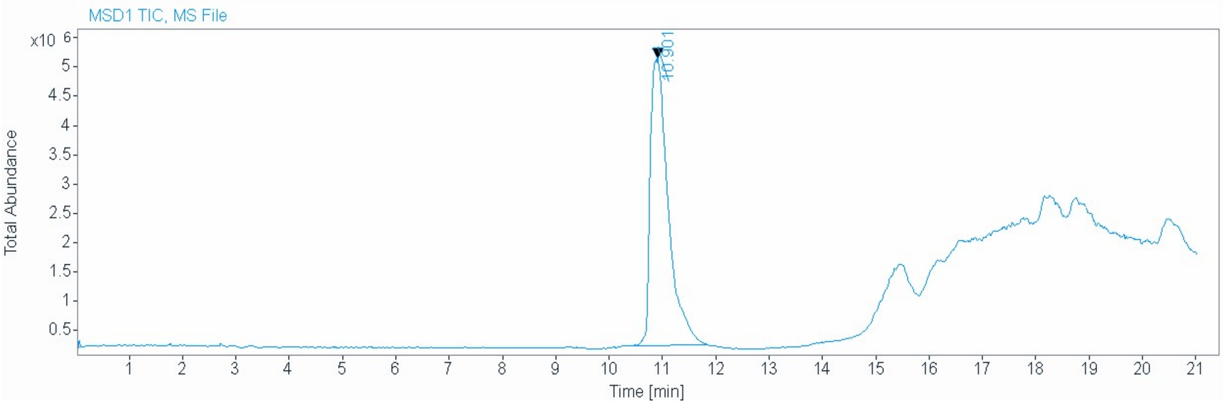


| | |
|---|---|
| **Signal** | MSD1 TIC, MS File |
| **Peak RT** | 10.901 |

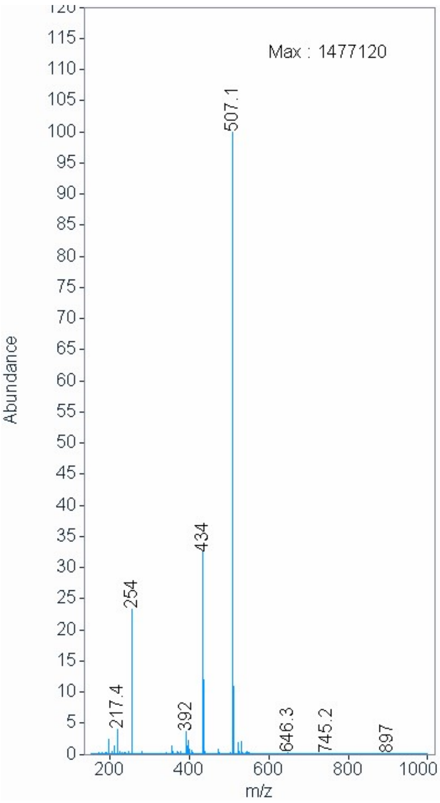

$^{1}H$ NMR compound **19**:

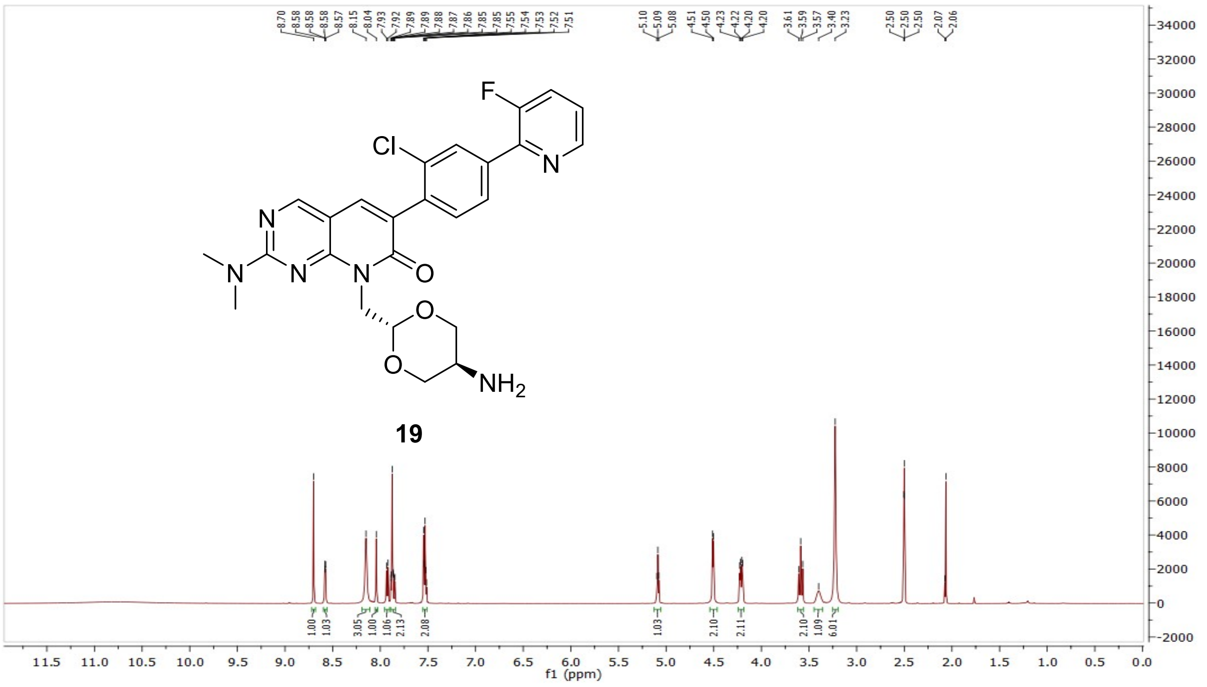


$^{13}C$ NMR compound **19**:

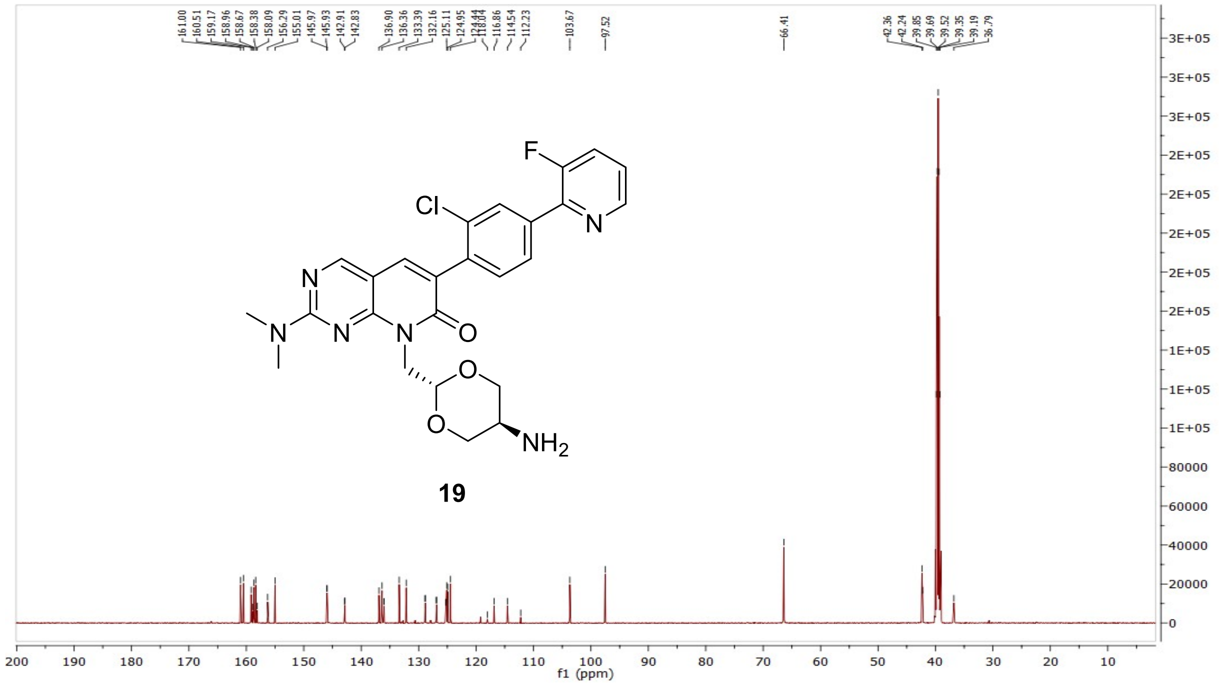

HRMS (FTMS +p MALDI) compound **19**:

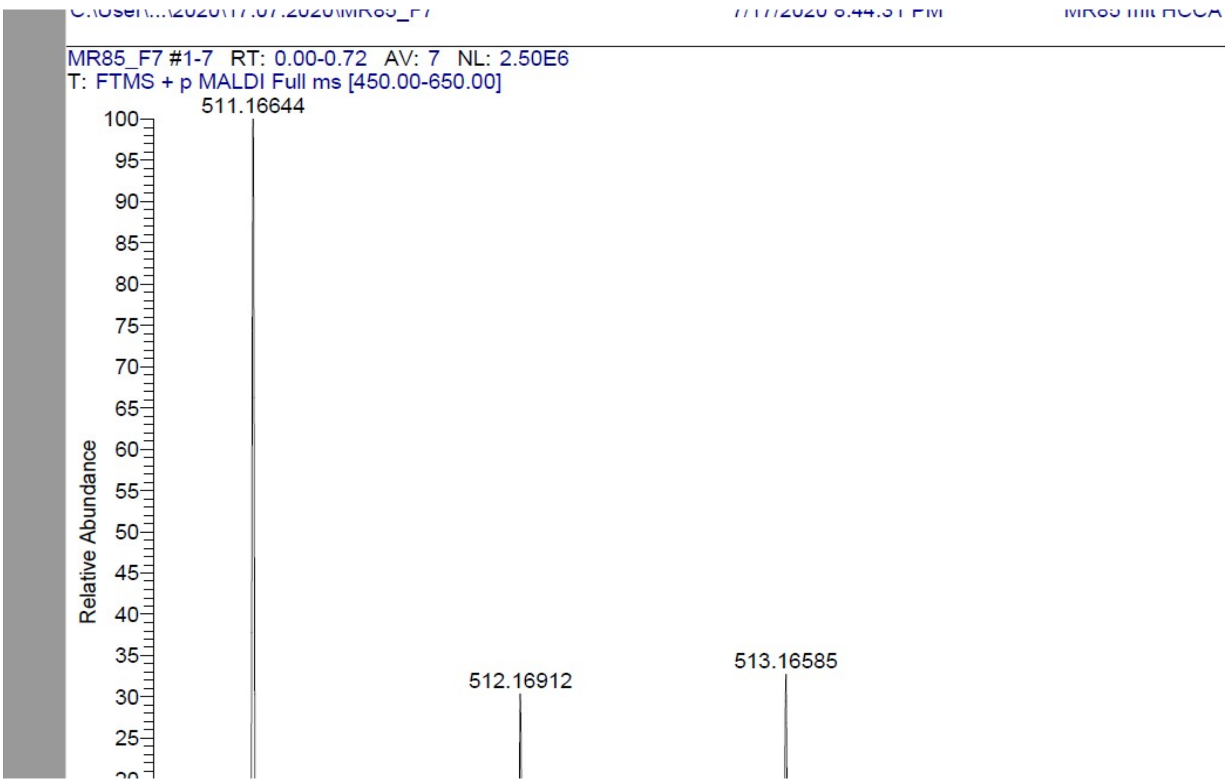

HPLC-chromatograms compound **19**:

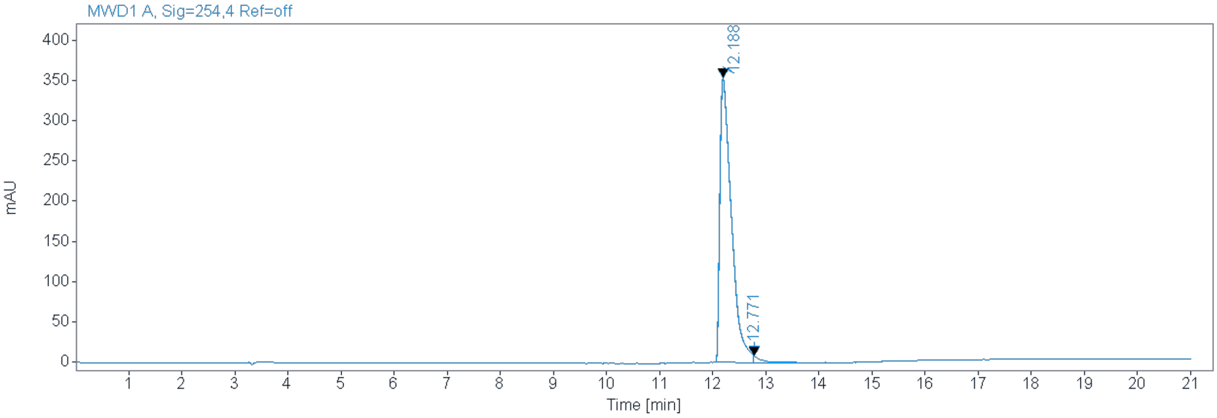


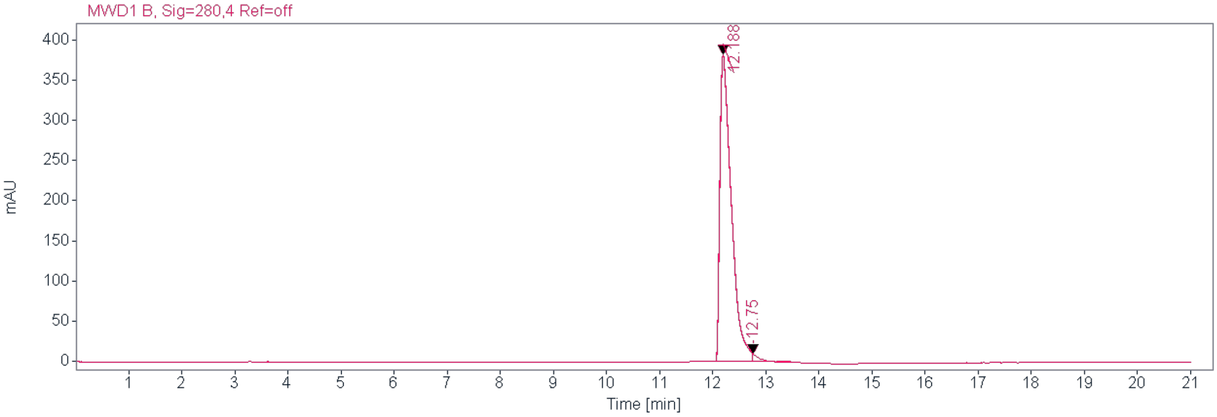


**Signal:** MWD1 A, Sig=254,4 Ref=off

| RT [min] | Type | Width [min] | Area | Height | Area% | Name |
|---|---|---|---|---|---|---|
| 12.188 | MM | 0.2522 | 5368.8589 | 354.7678 | 97.7154 | |
| 12.771 | MM | 0.1685 | 125.5265 | 8.9348 | 2.2846 | |
| | | Sum | 5494.3854 | | | |

**Signal:** MWD1 B, Sig=280,4 Ref=off

| RT [min] | Type | Width [min] | Area | Height | Area% | Name |
|---|---|---|---|---|---|---|
| 12.188 | MM | 0.2509 | 5768.1577 | 383.2370 | 98.0186 | |
| 12.750 | MM | 0.1898 | 116.5990 | 10.2369 | 1.9814 | |
| | | Sum | 5884.7567 | | | |

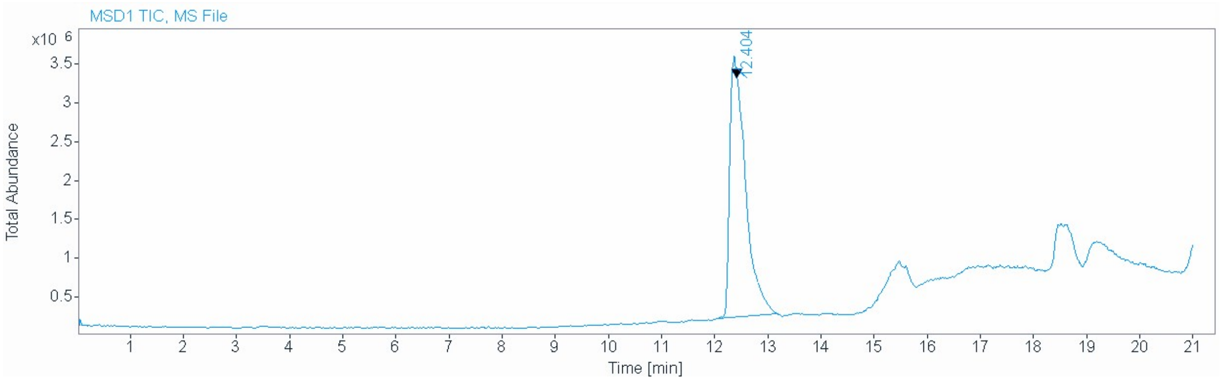


**Signal** MSD1 TIC, MS File

**Peak RT** 12.404

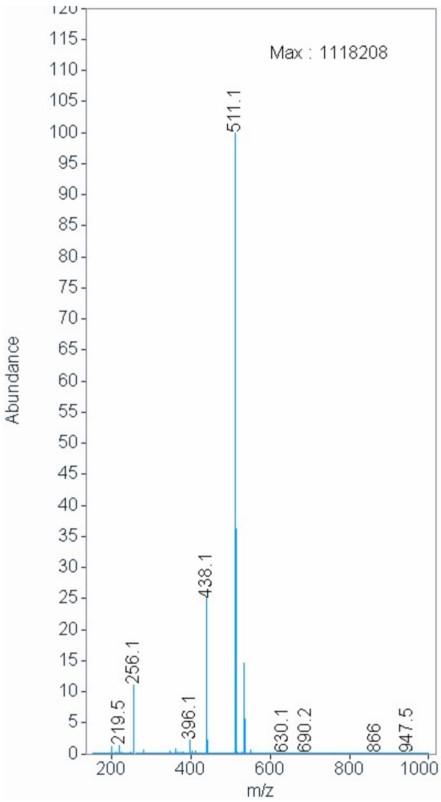
